# Subcellular localization and mitotic interactome analyses identify SIRT4 as a centrosomally localized and microtubule associated protein

**DOI:** 10.1101/2020.02.17.940692

**Authors:** Laura Bergmann, Alexander Lang, Christoph Bross, Simone Altinoluk-Hambüchen, Iris Fey, Nina Overbeck, Anja Stefanski, Constanze Wiek, Andreas Kefalas, Patrick Verhülsdonk, Christian Mielke, Dennis Sohn, Kai Stühler, Helmut Hanenberg, Reiner U. Jänicke, Jürgen Scheller, Andreas S. Reichert, M. Reza Ahmadian, Roland P. Piekorz

**Author notes:** Corresponding author: Roland P. Piekorz, Institute for Biochemistry and Molecular Biology II, Medical Faculty, Heinrich Heine-University Düsseldorf, D-40225 Düsseldorf, Germany.

## Abstract

The stress-inducible and senescence-associated tumor suppressor SIRT4, a member of the family of mitochondrial sirtuins (SIRT3, SIRT4, and SIRT5), regulates bioenergetics and metabolism *via* NAD^+^-dependent enzymatic activities. Next to the known mitochondrial location, we found that a fraction of endogenous or ectopically expressed SIRT4, but not SIRT3, is located at the mitotic spindle apparatus in the cytosol. Confocal spinning disk microscopy revealed that SIRT4 localizes during the cell cycle dynamically at centrosomes with an intensity peak in G_2_ and early mitosis. Moreover, SIRT4 binds to microtubules and interacts with structural (α,β-tubulin, γ-tubulin, TUBGCP2, TUBGCP3) and regulatory (HDAC6) microtubule components as detected by co-immunoprecipitation and mass spectrometric analyses of the mitotic SIRT4 interactome. Overexpression of SIRT4 resulted in a pronounced decrease of acetylated α-tubulin (K40) associated with altered microtubule dynamics in mitotic cells. SIRT4 or the N-terminally truncated variant SIRT4(ΔN28), which is unable to translocate into mitochondria, delayed mitotic progression and reduced cell proliferation. This study extends the functional roles of SIRT4 beyond mitochondrial metabolism, and suggests that SIRT4 acts as a novel centrosomal / microtubule-associated protein in the regulation of cell cycle progression. Thus, stress-induced SIRT4 may exert its role as tumor suppressor through mitochondrial as well as extramitochondrial functions, the latter associated with its localization at the mitotic spindle apparatus.

## Introduction

Mitotic cell division represents a complex and highly regulated process that allows the equal partitioning of duplicated DNA content from a mother cell into two daughter cells. The mitotic spindle apparatus is comprised of two centrosomes, one at each spindle pole, astral and spindle microtubules, and microtubule-associated protein (MAP) complexes ^1–3^. Centrosomes are the main microtubule organizing centers in animal cells comprising a pair of centrioles that are surrounded by pericentriolar material (PCM) components with pericentrin as the major anchoring factor ^4^. During the G_2_/M-phase of the cell cycle, cells undergo a massive microtubule rearrangement that functions as an important regulatory switch and consists of microtubule nucleation, elongation, polymerization, and depolymerization ^5^. The length of microtubules and their “dynamic instability” depend on an equilibrium shift between “catastrophe” (microtubule shrinkage) and “rescue” (microtubule growth) that is primarily regulated by several MAPs ^3^. The dynamic status of microtubules and their (in)stability are critically regulated by post-translational modifications, including (de)acetylation of α-tubulin (in particular lysine 40 [K40]) ^6,7^. The sirtuin SIRT2 and histone deacetylase 6 (HDAC6) are known deacetylases which target K40-acetylated α-tubulin in a NAD^+^-dependent manner ^8^, thereby altering microtubule dynamics by decreasing microtubule stability ^9,10^. The mammalian protein family of NAD^+^-dependent sirtuins (SIRT) comprises seven members which function in different cellular compartments mainly as deacetylases, deacylases, or ADP-ribosyltransferases. SIRT proteins are implicated in multiple pathways involved in epigenetic regulation and gene expression in the nucleus (SIRT1, 2, 6, and 7), proliferation/cell survival, aging, and life-span regulation (*e.g.* SIRT6) ^11–16^, as well as mitochondrial metabolism and bioenergetics (SIRT3, 4, 5) ^15–18^. SIRT3, a major mitochondrial deacetylase, and SIRT5 promote mitochondrial energy production, whereas SIRT4 exerts the opposite effect ^19–22^. In particular, the metabolic gatekeepers glutamate dehydrogenase (GDH) and pyruvate dehydrogenase (PDH) ^23,24^ are inhibited by SIRT4 by its ADP-ribosyltransferase and Lipoamidase activities, respectively ^25^. Furthermore, SIRT4 displays target-specific deacetylase ^26,27^ and deacylase ^25,28^ enzymatic activities.

Possible extra-mitochondrial roles of SIRT3 and SIRT5 are most likely due to their not well understood nuclear and cytosolic localization and their corresponding protein targets ^29–31^. SIRT3 regulates, in a direct or indirect (i.e. mitochondria-dependent) manner, microtubule dynamics and chromosomal alignment during mitosis by currently unknown mechanism(s) ^32,33^. SIRT4 could also play an extramitochondrial role in microtubule dynamics, given that SIRT4 interacts with Leucine-rich protein 130 (LRP130) ^24,34^, a multi-domain and dual-function protein that binds to the microtubule-associated protein MAP1S and integrates mitochondrial transport and the microtubule cytoskeleton in interphase ^35^. Moreover, recent work visualized a partial localization of SIRT4 into the nucleus that is even increased upon mitochondrial stress ^36^.

Consistent with a role for SIRT3 and SIRT4 as tumor suppressor proteins, knock-out mouse lines for SIRT3 and SIRT4 develop mammary and lung tumors, respectively ^37,38^. The tumorigenic phenotype of SIRT4 knock-out mice is associated with an increased chromosomal missegregation and aneuploidy/polyploidy that was also detected in primary SIRT4^−/−^mouse embryonic fibroblasts ^37^. Compared to wild-type cells, SIRT4^−/−^cells show increased DNA damage and sensitivity towards chromosomal instability upon treatment with stressors like UV radiation ^37^. It is unknown, whether the tumor phenotypes of mice lacking SIRT3 or SIRT4 are primarily based on mitochondria-dependent and/or -independent (i.e., mitotic/microtubule-associated) mechanisms. Moreover, SIRT4 was recently identified at the meiotic spindle apparatus during oocyte maturation. Oocytes from aged mice display higher SIRT4 levels leading to increased meiotic defects ^39^, which can be ameliorated by SIRT4 depletion. Consistent with its accumulation in aged oocytes, expression of SIRT4 is upregulated during replicative and stress-induced senescence, the latter triggered by different DNA-damaging stressors ^37^ as well as *in vivo* by UV radiation in photo-aged human skin ^40^.

Here, we performed subcellular localization and mitotic interactome analyses of SIRT4. Our findings indicate that besides its role in mitochondrial metabolism, SIRT4 functions also as a new centrosome- and microtubule-associated protein possibly involved in the regulation of mitotic cell cycle progression. In particular, ectopically expressed SIRT4 binds to α-Tubulin, interacts with HDAC6, and downregulates the levels of acetyl α-tubulin (K40) in G_2_-synchronized cells. Thus, both mitochondrial localized and extra-mitochondrial SIRT4 (presumably *via* metabolic inhibition/ROS generation ^41^ and alteration of mitotic regulation and/or microtubule dynamics, respectively) may trigger the anti-proliferative tumor suppressor function(s) of SIRT4 upon replicative/mitotic stress.

## Results

### SIRT4 localizes at interphase and mitotic centrosomes and in part at microtubules of the mitotic spindle

During our studies on the expression and mitochondrial function of SIRT4 ^40,42,43^ we noticed an extra-mitochondrial localization of endogenous SIRT4 at centrosomes and in part at the mitotic spindle in confocal laser scanning and spinning disk microscopy-based analyses of various human cell lines. We employed two independent anti-human SIRT4 antibodies, SAB1407208 (Sigma-Aldrich) raised against full-length human SIRT4 (a.a.1-314) and H-234 (sc-135053, Santa Cruz Biotechnology) raised against a N-terminally truncated version of human SIRT4 (a.a. 81-314). As depicted in Fig. S1, SIRT4 was detected at interphase/G_2_ centrosomes in HT1080 fibrosarcoma cells and progressively colocalized with the PCM component Pericentrin and MTOCs (microtubule organizing centers) as indicated by α-tubulin staining. A comparable staining of SIRT4 at interphase centrosomes was observed in Colo-680N human esophageal and in HEK293 human embryonic kidney cells (Figs. S2 and S3). Moreover, we detected SIRT4 at the mitotic spindle colocalized with the spindle marker TACC3 (Transforming Acidic Coiled-Coil 3) ^44^ and in reduced levels at the central spindle (Fig. S3). Besides its extramitochondrial/centrosomal localization, SIRT4 was observed as described ^23,42^ in mitochondria using a co-staining against the mitochondrial marker MTC02 (Fig. S4). Similar to endogenously expressed SIRT4, C-terminal eGFP fusion proteins of SIRT4 and SIRT4(ΔN28), the latter representing an N-terminally (a.a. 1-28) truncated SIRT4 mutant unable to translocate into mitochondria ^42,45^, were also detected at interphase centrosomes of HeLa cervix carcinoma and HT1080 fibrosarcoma cells (Fig. S6 and suppl. Movie 5). In clear contrast to SIRT4, SIRT3 was not detectable at interphase or mitotic centrosomes or at the spindle apparatus, but displayed solely a mitochondrial localization (Fig. S5) as previously described ^46^.

### Centrosomal localization kinetics of SIRT4 during cell cycle progression

Next, we quantitatively analyzed SIRT4 at centrosomes during G_2_ and the course of mitotic cell division using Pericentrin as centrosomal marker. In HeLa cervix carcinoma cells, SIRT4 showed a dynamic centrosomal localization pattern where it displayed the highest signals in centrosomal staining during G_2_ and early mitosis, followed by a significant drop in signal intensity from prophase onwards until late mitosis/cytokinesis (Fig. 1a and b). At the same time, centrosomal Pericentrin levels were comparable between G_2_ and metaphase, but significantly dropped thereafter in the second half of mitosis (Fig. 1a and b). Similarly, SIRT4-eGFP that was transiently expressed in HeLa cells localized at centrosomes in interphase cells (suppl. Movie 1), prominently decorated Pericentrin at spindle poles in metaphase cells (suppl. Movie 2), and lastly disappeared from centrosomes during telophase (suppl. Movie 4). Parallel control imaging experiments failed to detect eGFP localization at centrosomes (suppl. Movie 3). Thus, centrosomal localization of SIRT4 seems to be dynamically regulated during cell cycle progression with a peak in G_2_ and early mitosis.

**Figure 1.**
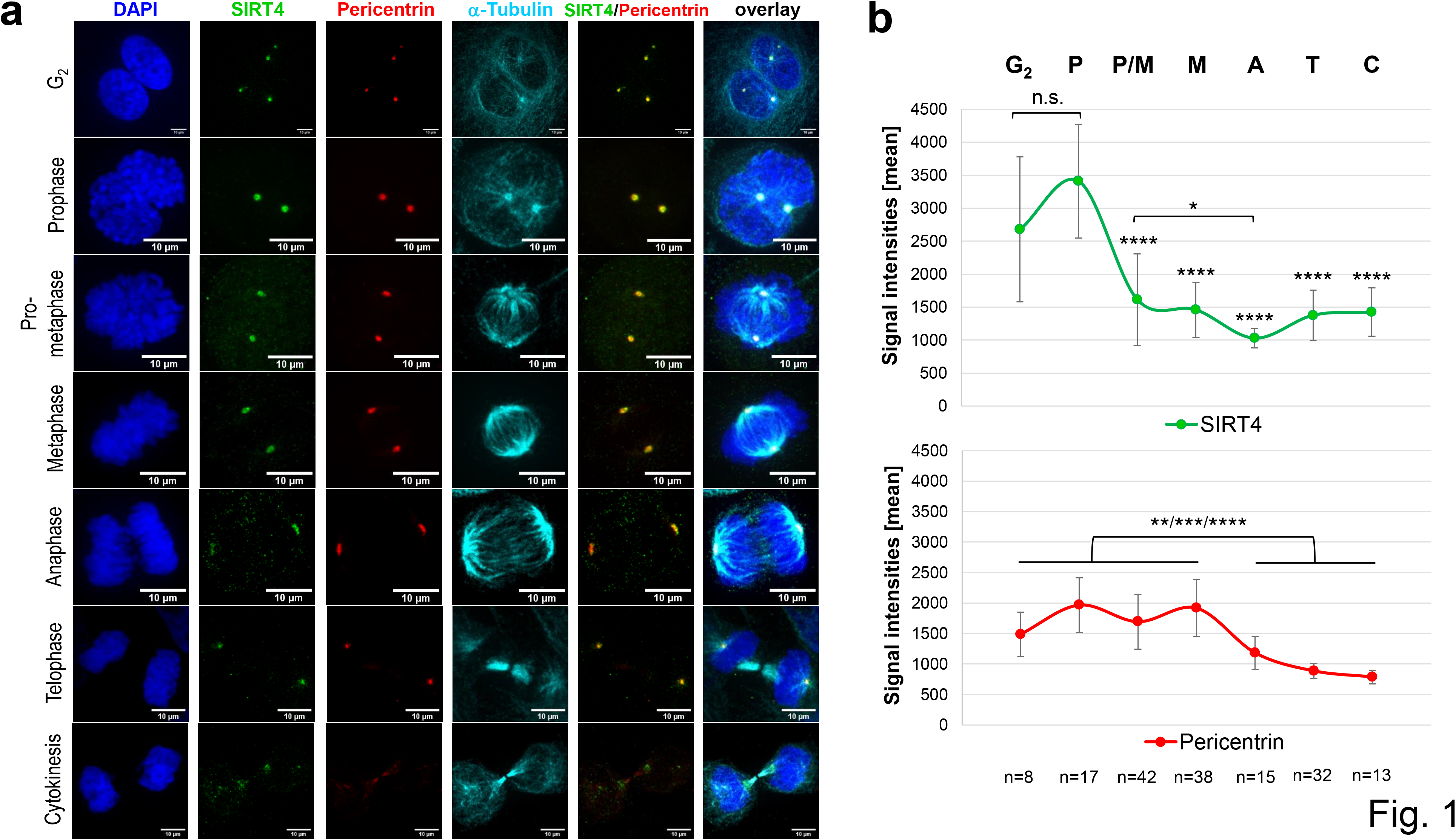
Centrosomal localization pattern of SIRT4 during G_2_/M progression. (**a**) Endogenous SIRT4 was detected in HeLa cells in G_2_ and subsequent mitotic stages using a polyclonal antibody against SIRT4 (sc-135053, Santa Cruz Biotechnology) and spinning disk microscopy. Antibody staining against Pericentrin and α-tubulin was employed to visualize centrosomes and microtubules, respectively. DAPI was used to detect DNA. Bar: 10 μm. (**b**) Quantification of centrosomal SIRT4 and Pericentrin levels during G_2_/M progression. Endogenous SIRT4 and Pericentrin were detected in HeLa cells as indicated above. Shown are mean signal intensities (± S.D.) which were analysed using ImageJ software (Materials & Methods). Numbers of cells analysed per mitotic phase are indicated. P, prophase; P/M, prometaphase; M, metaphase; A, anaphase; T, telophase; C, cytokinesis. To evaluate statistical significance (comparison of SIRT4 intensities between Prophase and G_2_ or mitotic phases) one-way ANOVA followed-up by Tukey’s test was performed (*p<0.05; **p<0.01; ***p<0.001; ****p<0.0001; n.s., not significant).

### Subcellular fractionation reveals a cytosolic, extra-mitochondrial pool of SIRT4, but not SIRT3

Given the extramitochondrial localization of SIRT4 at centrosomes, we next analysed the intracellular SIRT4 protein distribution by subjecting total cell lysates to a subcellular fractionation protocol *via* differential centrifugation steps (Material & Methods). The method yielded in a highly cleared cytosolic fraction together with a mitochondrial enriched particulate fraction as controlled by marker proteins specific for subcellular compartments. Interestingly, in addition to their mitochondrial localization, both endogenous SIRT4 (Fig. 2) as well as ectopically expressed SIRT4-eGFP (Fig. S7) were also found at substantial levels in the cytosolic fraction. In the case of SIRT5, we observed also a clear cytosolic fraction of ectopically expressed, C-terminally Flag-tagged SIRT5 (Fig. S8), consistent with previous findings ^31^. In contrast, endogenous SIRT3 or C-terminally Flag-tagged SIRT3 were almost exclusively found in the mitochondrial enriched fraction (Fig. 2 and Fig. S8, respectively). Taken together, our findings obtained from confocal microscopic imaging and subcellular fractionation analyses both indicate that substantial levels of SIRT4, but not SIRT3, localize outside mitochondria in the cytoplasm.

**Figure 2.**
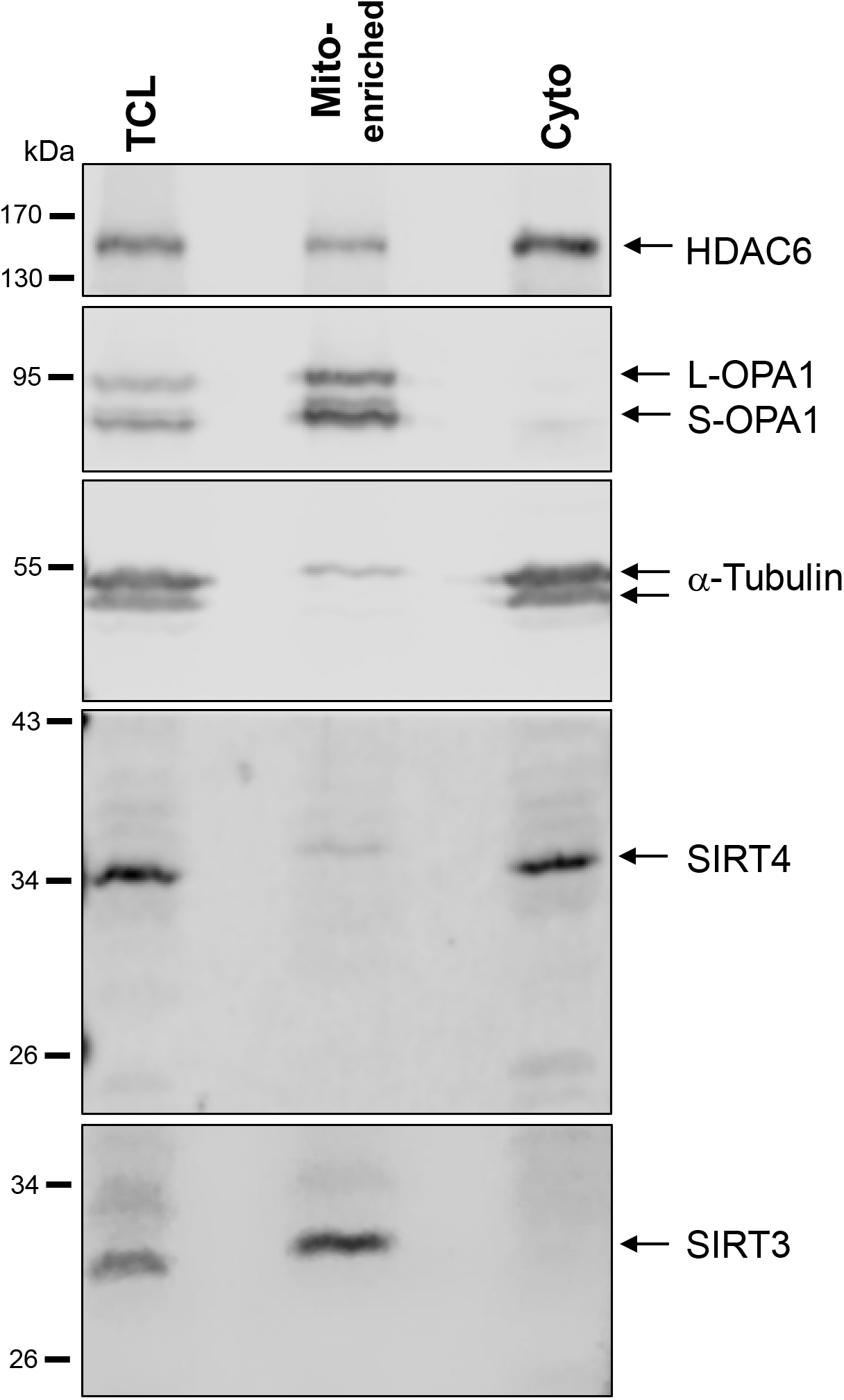
Subcellular fractionation analysis of endogenous SIRT4 and SIRT3 protein levels in HEK293 cells. Total cell lysates (TCL; 80 μg) and the respective mitochondria enriched (Mito-enriched; 50 μg) and cytosolic (Cyto; 80 μg) fractions were subjected to immunoblot analysis. Subcellular marker proteins detected were OPA1 (long and short forms of OPA1; mitochondria), α-tubulin (cytoplasm), and HDAC6 (predominantly cytoplasmatic localization).

### Ectopic overexpression of SIRT4 or the extra-mitochondrial localized deletion mutant SIRT4(ΔN28) inhibits mitotic progression and cell proliferation

Given the stress induced/DNA damage-associated upregulation of SIRT4 and its anti-proliferative role ^37,40^, we next aimed to link SIRT4 localized at centrosomes and the mitotic spindle to a possible inhibitory function on cell cycle progression and proliferation. HEK293 cells stably expressing SIRT4-eGFP, SIRT4(ΔN28)-eGFP, the catalytically inactive mutant SIRT4(H161Y)-eGFP, or eGFP as control, were subjected to continuous live cell imaging analyses during cell division. As depicted and quantitatively analyzed in Fig. 3, expression of all three SIRT4 variants led to a significant prolongation of mitosis with strongest impacts of SIRT4-eGFP and the exclusively outside mitochondria localized SIRT4(ΔN28)-eGFP fusion protein. In accordance with these findings, cellular proliferation was significantly reduced by all three SIRT4 variants as compared to eGFP-expressing cells (Fig. 3c). Of note, expression of SIRT4(ΔN28)-eGFP was associated with an almost three-fold increase in bi- or multinucleated cells (Fig. S9). The observation that SIRT4(H161Y)-eGFP, albeit catalytically inactive, still delays mitosis and inhibits proliferation indicates that SIRT4 possibly targets structural or regulatory factors in cell cycle progression (and/or mitochondrial function that then impacts on mitosis) through both catalytically-dependent and -independent mechanisms.

**Figure 3.**
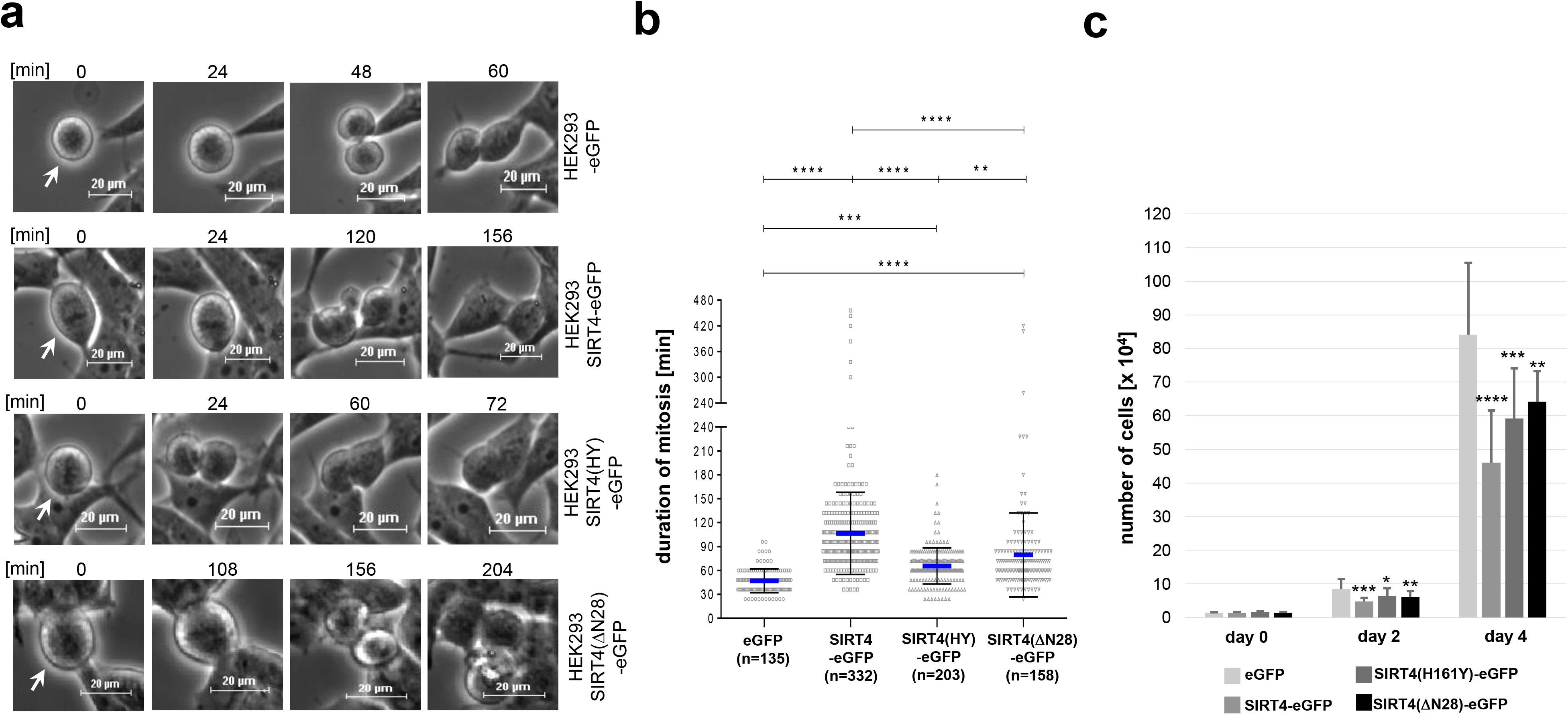
Ectopic expression of SIRT4 prolongs mitotic progression and inhibits cell proliferation. (**a**) HEK293 cell lines stably expressing SIRT4-eGFP, the enzymatically inactive mutant SIRT4(H161Y)-eGFP, or SIRT4(ΔN28)-eGFP lacking the N-terminal mitochondrial targeting signal were analyzed by live cell imaging. Cell lines were cultured in CO_2_-independent, HEPES containing media. Pictures of cells undergoing cell division were taken in intervals of 12 min. (**b**) Duration of mitosis was calculated from cell rounding/early mitosis to cytokinesis/cell reattachment. To evaluate statistical significance one-way ANOVA followed-up by Tukey’s test was performed (**p<0.01; ***p<0.001; ****p<0.0001). (**c**) Proliferation kinetics of HEK293 cells expressing SIRT4-eGFP or the indicated SIRT4 mutants. Cells were seeded at day 0 (15.000 cells/well) in triplicates and total cell numbers were counted at the indicated time points (N=3-5 independent experiments). To evaluate statistical significance one-way ANOVA followed-up by Tukey’s test was performed (*p<0.05; **p<0.01; ***p<0.001; ****p<0.0001).

### The mitotic SIRT4 interactome comprises microtubule-associated structural and regulatory proteins

To better understand the mechanism(s) through which SIRT4 impacts on mitosis, we next analysed the SIRT4 interactome in mitotic SIRT4-eGFP expressing HEK293 cells as compared to eGFP expressing control cells. Cells were synchronized in G_2_ by RO3306-mediated, reversible inhibition of cyclin dependent kinase 1 (CDK1) followed by release into mitosis for 45 min. Native SIRT4 containing protein complexes were isolated by anti-eGFP nanobody-based co-immunoprecipitation from total cell lysates followed by mass spectrometric characterization of SIRT4 interacting proteins (Table S1). Protein network analyses revealed several known (*e.g*. DNA damage response ^37^; mitochondrial respiratory chain components and glutamate metabolism regulators ^47^; regulation of mitochondrial organization ^42,43^) as well as novel functions and components associated with the mitotic SIRT4 interactome (*e.g* tRNA aminoacylation and mitochondrial translation; cell cycle regulation; microtubule regulation) (Fig. S11). In particular, as depicted in Fig. 4, we identified several mitochondrial SIRT4-binding proteins and potential substrates (OPA1 ^42^; ATP5F1A ^34^; ANT2 ^21^; IDE ^48^) as well as mostly novel, extra-mitochondrial localized SIRT4 interactors. The latter comprise α- and β-tubulin as subunits of microtubules, components of the centrosomally localized γTURC complex (γ-tubulin, TUBGCP2, TUBGCP3) ^49,50^ that nucleates microtubules at their minus poles, the microtubule deacetylase HDAC6 that is critically involved in the regulation of microtubule stability and dynamics ^9^, and the G_2_/M cell cycle regulator CDK1 ^51^. These SIRT4 interactions were confirmed by nanobody-mediated co-immunoprecipitation analyses (Fig. 4b). The observed mitotic interaction pattern of SIRT4 appears specific, given that SIRT4 failed to co-immunoprecipitate with other centrosomal or mitotic spindle localized proteins like Pericentrin (Fig. S10) or the TACC protein family member TACC3 ^42^, respectively.

**Figure 4.**
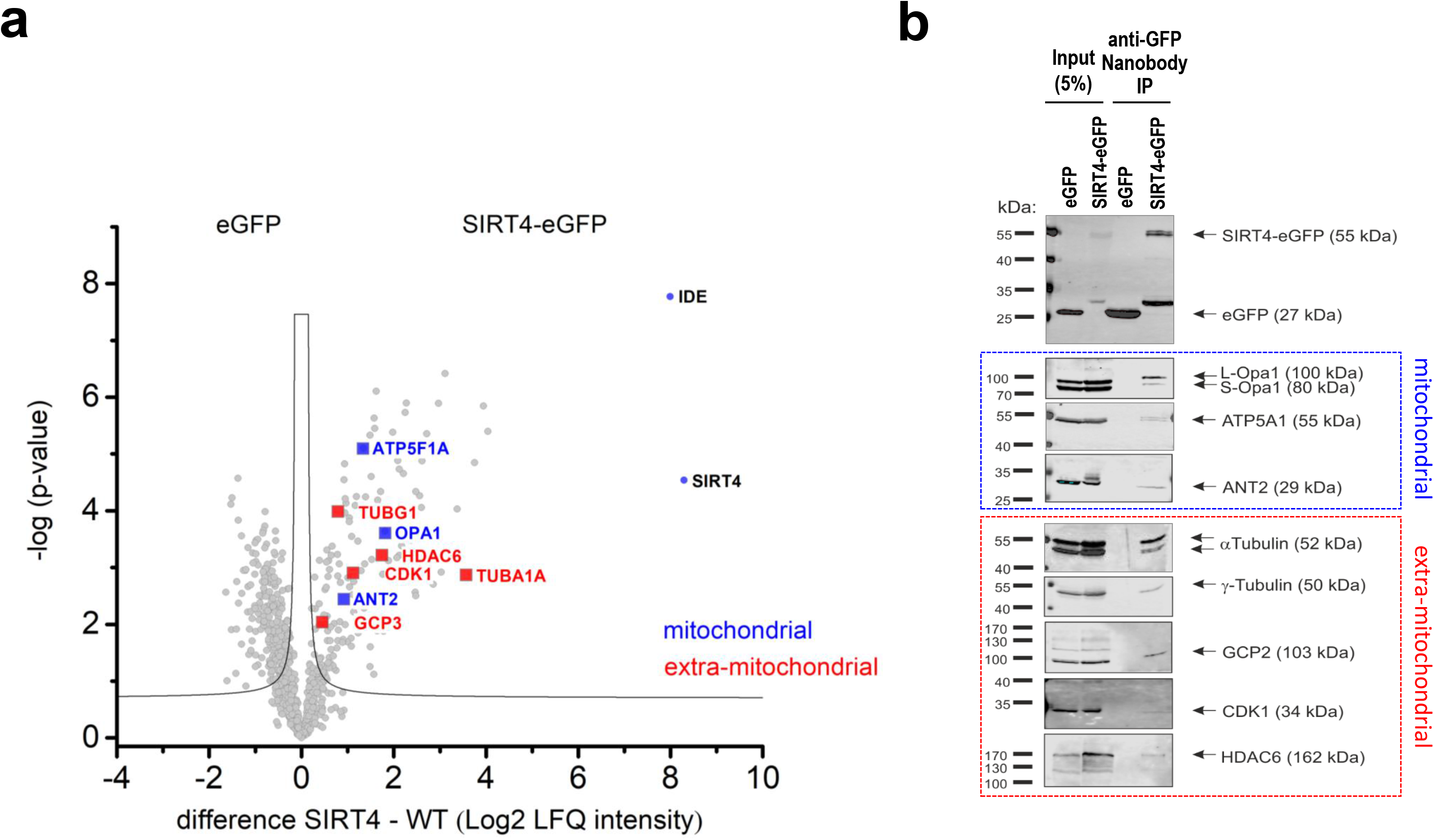
Analysis of the mitotic SIRT4 interactome by mass spectrometry. (**a**) Volcano plot analysis of mitochondrial and extra-mitochondrial proteins interacting with SIRT4. HEK293 cells stably expressing eGFP or SIRT4-eGFP (n=4 replicates each) were arrested in G_2_ by the reversible CDK1 inhibitor RO3306 followed by release into mitosis (45 min after RO3306 wash out). Total cell lysates were subjected to anti-eGFP nanobody co-immunoprecipitations followed by their mass spectrometric analysis. (**b**) Anti-eGFP nanobody co-immunoprecipitations of selected mitochondrial and extra-mitochondrial SIRT4 interacting proteins as compared to eGFP controls. Total cell lysates analysed in (a) were employed.

### SIRT4 binds to microtubules and negatively regulates acetyl-α-tubulin (K40) levels

Given the link between SIRT4 and proteins of the microtubule network, we next addressed the interaction of SIRT4 with microtubules and its role in regulation of microtubule dynamics. We performed microtubule pulldown assays and observed SIRT4-eGFP in the pelleted fraction of Taxol-stabilized microtubules (Fig. 5a). In contrast to this, eGFP as control was almost exclusively detected in the soluble fraction of Taxol-stabilized microtubules (Fig. 5a). Consistent with these findings, (i) an α-tubulin specific antibody coimmunoprecipitated α-tubulin and SIRT4-eGFP, but not eGFP (Fig. 5b), and (ii) α-tubulin co-localized with endogenous SIRT4 at MTOCs in mitotic cells as detected by spinning disk microscopy (Fig. 5c). Given the interaction (Fig. 4) and partial mitotic colocalization (suppl. Movie 6) of SIRT4 with the microtubule deacetylase HDAC6, we next analyzed the levels of K40-acetylated α-tubulin upon ectopic expression of SIRT4-eGFP or mutants thereof. Interestingly, as indicated in Fig. 6, SIRT4-eGFP, but not the catalytically inactive mutant SIRT4(H161Y)-eGFP or SIRT4(ΔN28)-eGFP, led to a profound decrease in the ratio of K40-acetylated α-tubulin *vs*. total α-tubulin levels in G_2_ synchronized HEK293 cells as compared to asynchronously growing cells. Thus, our findings indicate that full-length SIRT4 impacts in an enzymatically dependent manner on microtubule dynamics in G_2_/M by decreasing microtubule stability, which presumably translates into inhibition of mitotic progression and proliferation (Fig. 3).

**Figure 5.**
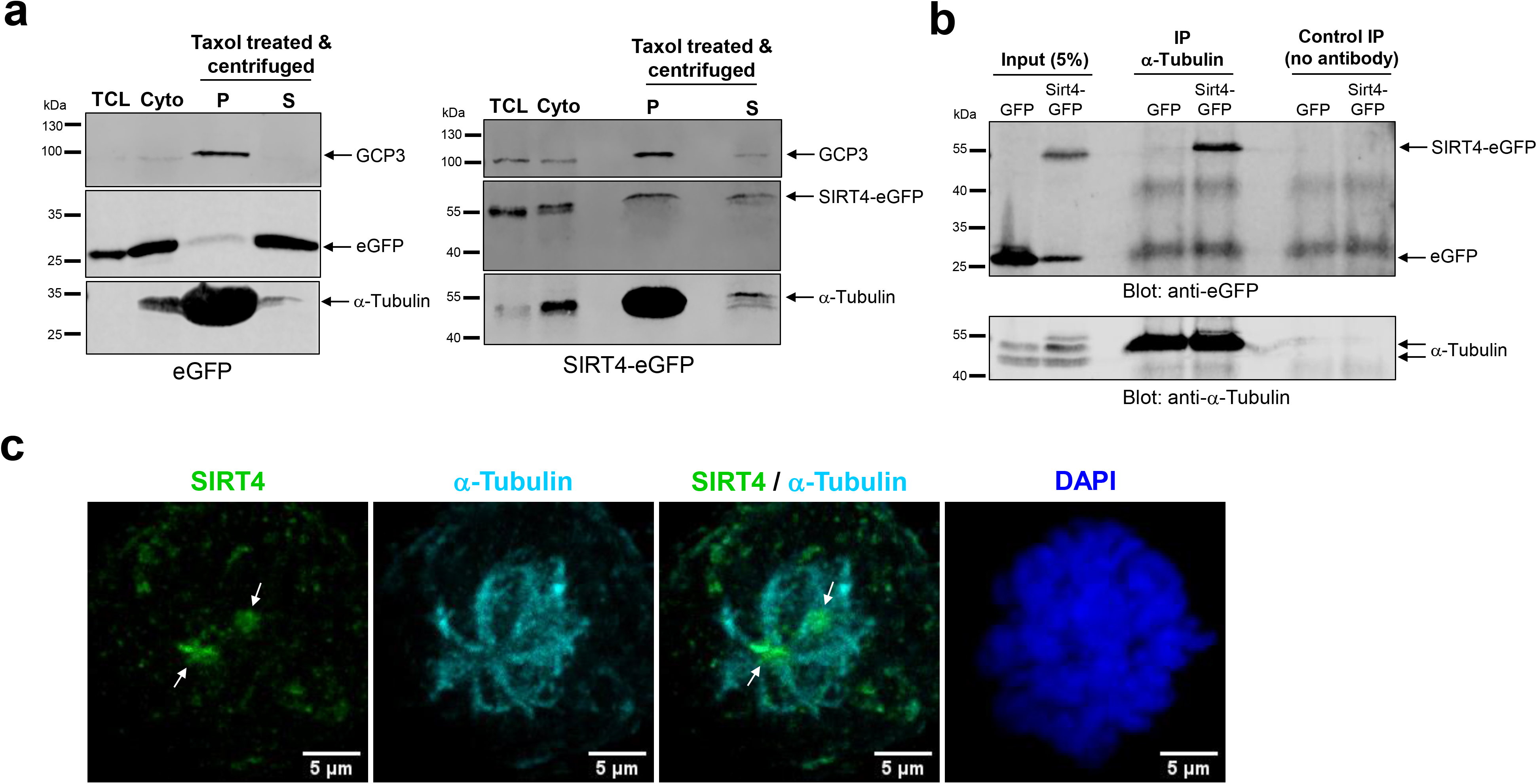
SIRT4 binds to microtubules and co-immunoprecipitates with α-tubulin in HEK293 cells. (**a**) SIRT4-eGFP, but not eGFP, is present in the pelleted fraction (P) of microtubules which were Taxol-stabilized in the cytosolic fraction (Cyto) followed by pelleting *via* centrifugation through a sucrose cushion. TCL, Total cell lysate; S, supernatant. Tubulin Gamma Complex Associated Protein 3 (TUBGCP3 or GCP3) was detected as co-marker for microtubules (**b**). An α-tubulin specific antibody co-immunoprecipitates SIRT4-eGFP, but not eGFP, from total cell lysates of stably transfected HEK293 cells. As control, immunoprecipitation without α-tubulin antibody was performed. (**c**) Localization of SIRT4 at spindle poles/Microtubule Organizing Centers (MTOCs) of mitotic HeLa cells using a polyclonal antibody against SIRT4 (sc-135053, Santa Cruz Biotechnology) and analysis by spinning disk microscopy. Antibodies against α-Tubulin were employed to visualize microtubules. DAPI was used to detect DNA. Bar: 5 μm.

**Figure 6.**
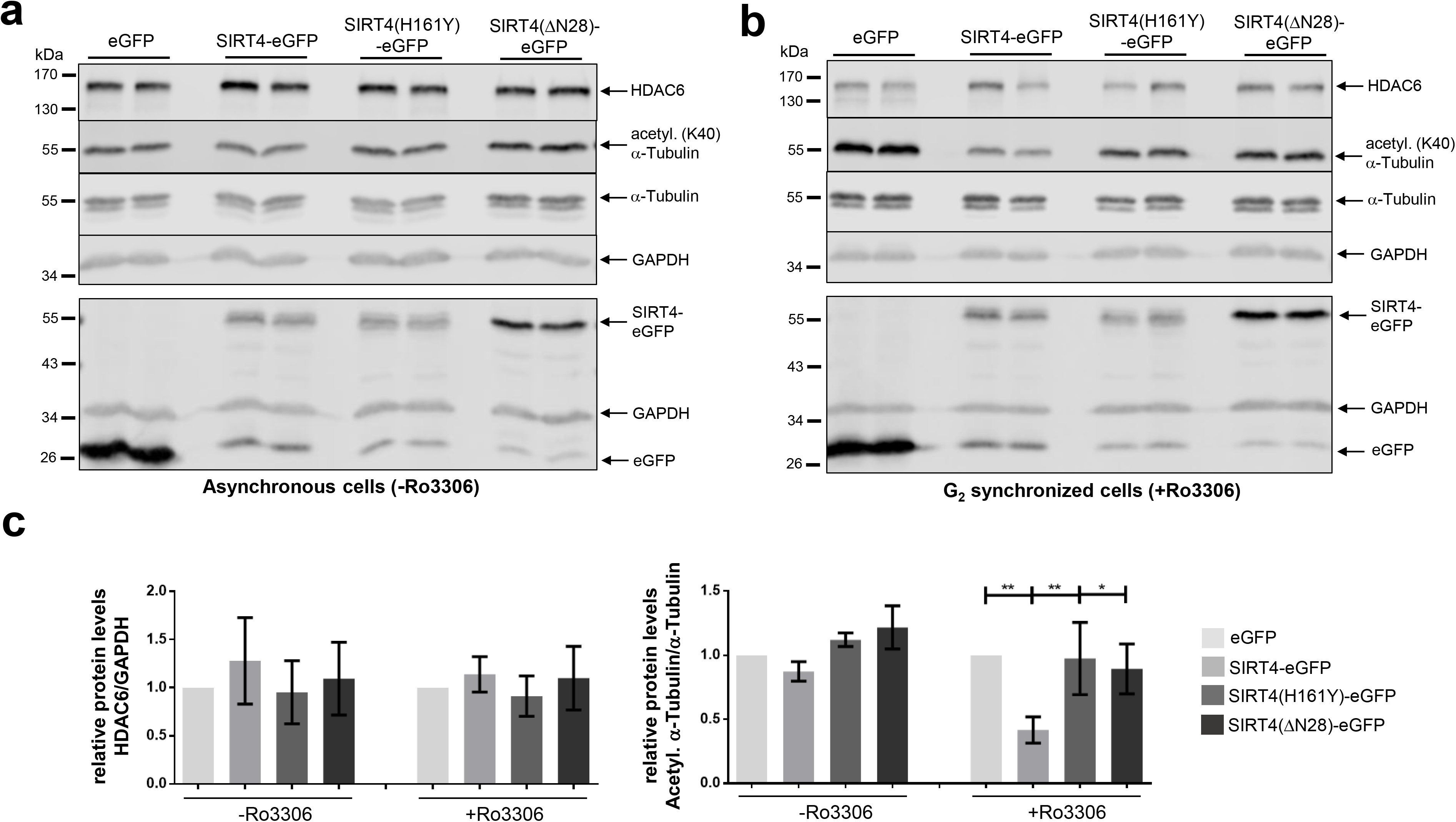
Ectopic expression of SIRT4 impacts negatively on acetylated α-tubulin (K40) during mitosis. HEK293 cells stably expressing eGFP, SIRT4-eGFP, or mutants thereof were either asynchronously grown (**a**) or subjected to RO3306-mediated G_2_ synchronization (**b**) followed by immunoblot analysis of HDAC6, acetylated α-tubulin (K40), and α-tubulin. Expression of eGFP, SIRT4-eGFP, or mutants thereof was analysed in a second immunoblot (lower panels in **a** and **b**). Probing against GAPDH was employed as loading control. Relative HDAC6 amounts and acetylated α-tubulin (K40)/α-tubulin levels were quantified by densitometric analysis (**c**). To evaluate statistical significance one-way ANOVA followed-up by Tukey’s test was performed (three independent experiments; *p<0.05; **p<0.01). All P-values of the analysis of acetylated α-tubulin (K40)/α-tubulin levels (+Ro3306) refer to comparison with SIRT4-eGFP.

## Discussion

This study provides insights into a potential role of extra-mitochondrially localized SIRT4 in mitotic cell division. Our findings show that (i) SIRT4 localizes not only in mitochondria, but also in the cytosol, where it is localized at centrosomes especially in early mitotic phases; (ii) as revealed by mass spectrometric, co-immunoprecipitation, and microtubule pulldown analyses, SIRT4 co-pellets with Taxol-stabilized microtubules and interacts with microtubule components and binding proteins, in particular α-tubulin, with components of the centrosome associated γTURC complex (γ-tubulin, GCP2, GCP3), and with the α-tubulin deacetylase HDAC6; (iii) linked to the SIRT4-HDAC6 interaction, increased SIRT4 expression results in decreased acetyl-α-tubulin (K40) levels, which are typically associated with decreased stability and altered dynamics of mitotic microtubules; (iv) at the cellular level, ectopic expression of SIRT4 or SIRT4(ΔN28) lacking the N-terminal mitochondrial targeting signal prolongs mitotic progression and inhibits cell proliferation. Consistent with the subcellular localization profile of endogenous SIRT4 (Fig. 1, 2, and Fig. S4), it has been recently reported that ectopic SIRT4 expressed even at very low levels shows a dual localization in mitochondria as well as in the cytosol and nucleus. The authors attributed this to a low mitochondrial import kinetics of SIRT4 ^36^. Thus, we propose that SIRT4 may exert its cell cycle inhibitory and tumor suppressor function through both mitochondria, *i.e*. bioenergetics- dependent, and mitochondria-independent, *i.e*. centrosome/mitotic spindle apparatus-linked mechanisms. Interestingly, the dual mitochondrial and centrosomal localization of SIRT4 (shown in this work) and its increased nuclear localization upon mitochondrial stress^36^ is reminiscent of a function of SIRT4 as “moonlighting protein” that per definition localizes at more than one cellular compartment/structure with similar or different functions ^52^. Further examples of mitochondrial and centrosomal localized moonlighting proteins include C21orf33/GATD3A (glutamin amidotransferase like class 1 domain containing 3A), which has been identified within the human protein atlas project ^53^, and the mitochondrial porin VDAC3 (voltage-dependent anion-selective channel protein 3) that localizes at centrosomes and regulates centriole assembly ^54^.

In terms of the regulation of extramitochondrial SIRT4, we observed the highest centrosomal levels in G_2_ and early mitosis (Fig. 1), indicating that centrosomal recruitment of SIRT4 (and possibly its dissociation towards mitotic exit) (suppl. movies 1-4) represents a regulated process. This is supported by the observation that total SIRT4 protein levels did not greatly change during cell cycle progression when cells were released from double thymidine block-mediated G_1_/S synchronization (data not shown), although we cannot exclude that the cytosolic (*i.e.* extramitochondrial) pool of SIRT4 does. The low import kinetics of SIRT4 into mitochondria ^36^ could be further reduced in G_2_/M when mitochondria increasingly undergo cyclin B1-CDK1 driven fission ^55^ to become evenly distributed around the mitotic spindle ^56^ for cell division. Elevated cytosolic SIRT4 levels in G_2_/M might then result in increased recruitment of SIRT4 to the centrosome, a hypothesis that remains to be further tested. A candidate regulator of centrosomal SIRT4 localization is CDK1, given that both proteins interact with each other (Fig. 4) and partially colocalize at centrosomes in G_2_ (Bergmann et al., unpublished). Interestingly, recent findings uncovered an additional intramitochondrial role of CDK1 where it interacts with and phosphorylates SIRT3 to enhance mitochondrial metabolism^57,58^. Given that the CDK1 phosphorylation sites are conserved between SIRT3 and SIRT4 (Bergmann et al., unpublished), it will be important to further analyse the nature and function of a CDK1-SIRT4 axis in mitotic *vs*. non-mitotic cells. Overall, our findings add to the increasing evidence for a centrosomal localization of deacetylases (HDACs and sirtuins) ^59^ and their critical function(s) in centrosome biology, microtubule dynamics, and mitotic regulation. For example, the SIRT1-Plk2 (Polo-like kinase 2) and SIRT1-CCDC84-SAS6 axes control centriole duplication ^60^ and prevent centrosome overduplication ^61^, respectively. Expression of SIRT2, which is involved in the regulation of microtubule dynamics ^8,10,^ is regulated in a cell cycle-dependent manner where SIRT2 localizes to centrosomes and the mitotic spindle ^62^. Phosphorylation of SIRT2 by cyclin A-CDK2 reduces binding of SIRT2 to centrosomes and promotes G_2_/M progression ^63^. In line with this, increased SIRT2 levels due to mitotic stress cause an extension of the mitotic phase ^64^ presumably through the regulatory role of SIRT2 towards the anaphase promoting factor/cyclosome (APC/C) and hence cyclin B1 degradation ^65^.

Our proteome analysis further suggests a role of SIRT4 in the regulation of microtubule dynamics and function which may mechanistically relate to the inhibitory impact of ectopically expressed SIRT4 on cell division and proliferation (Fig. 3). SIRT4 binds microtubules (Fig. 5) and interacts (Fig. 4 and Fig. S12) and co-localizes (Fig. S12) with γ-tubulin, TUBGCP2, and TUBGCP3, which represent core components of the γTURC. The latter is located at the outer region of the pericentriolar material (PCM) ^66,67^ and functions as nucleator of microtubules at their minus poles ^49,50^. It remains to be determined whether SIRT4 directly or indirectly regulates recruitment of the γTURC to centrosomes or its microtubule nucleation activity. However, besides phosphorylation, only few other posttranslational modifications have been so far described for γTURC components, including an acetylation of GCP2 at Lys827 with currently unknown function ^68,69^. A second microtubule linked interaction partner of SIRT4 represents the deacetylase HDAC6 (Fig. 4 and suppl. Movie 6) that was also identified as *bona fide* SIRT4 interactor in the screen by Mathias et al. ^24^. HDAC6 is mainly found in the nucleus and cytosol and also localizes at the centrosome and basal body where it is involved in ciliary disassembly ^70^. HDAC6 targets besides HSP90 and cortactin K40-acetylated α-tubulin, resulting in decreased microtubule stability and altered microtubule dynamics ^9,10^. Interestingly, consistent with a role of SIRT4 in microtubule dynamics, ectopic expression of SIRT4 strongly inhibits the levels of acetylated α-tubulin (K40) in G_2_-synchronized HEK293 cells (Fig. 6). The absent effect of SIRT4(ΔN28)-eGFP expression on K40-acetylated α-tubulin levels (Fig. 6) was unexpected, given that SIRT4ΔN28-eGFP coimmunoprecipitates both with α-tubulin (Fig. S10) and HDAC6 (data not shown). Thus, either SIRT4 requires its N-terminus for its extramitochondrial function towards modulation of K40-acetylated α-tubulin levels, or SIRT4 impacts through additional acetyl-α-tubulin (K40) independent mechanism(s) on euploidy (Fig. S9) and mitotic progression and proliferation (Fig. 3). The latter possibility is supported by the reduced, but still significant inhibitory impact of SIRT4(ΔN28) on mitotic duration and proliferation as compared to the full effect of wild-type SIRT4. It remains to be determined whether SIRT4 regulates acetylated α-tubulin (K40) levels directly as deacetylase or indirectly *via* interaction with the known microtubule deacetylase HDAC6. The latter mechanism has been e.g. described for the tumor suppressor RITA (RBP-J and tubulin-associated protein) that interacts with HDAC6 and thereby modulates levels of K40-acetylated α-tubulin and microtubule dynamics ^71^. Lastly, the NAD^+^-SIRT3 axis has been also implicated in the regulation of microtubule dynamics and chromosomal alignment during mitosis ^32,33^. However, this function of SIRT3 is likely mitochondrial-based, given that SIRT3 is predominantly found in mitochondria (Fig. 2 and Fig. S5) and neither localizes at centrosomes nor at the mitotic spindle (Fig. S5 and data not shown).

Recent overviews of the literature revealed that SIRT4, although first described as a metabolic tumor suppressor ^37^, may display both tumor suppressor and oncogenic/cancer promoting activities, depending on the tumor type and checkpoint activating conditions ^72,73^. Our data on the putative SIRT4-HDAC6-microtubule dynamics axis are rather consistent with an extramitochondrial tumor suppressor function of SIRT4. Thus, stress-induced SIRT4 as in the case of DNA damage and senescence induction may exert its anti-proliferative role through both mitochondrial/metabolism and mitochondria independent functions, the latter associated with its localization and function at the mitotic spindle apparatus.

## Material and Methods

### Cell culture

HEK293, HT1080, Colo-680N, and HeLa cell lines were cultured at 37°C and 5% CO_2_ in DMEM (Dulbecco’s Modified Eagle Medium) containing high glucose (4.5 g/L; Thermo Fisher Scientific) with 10% fetal bovine serum (FBS) and penicillin (100 units/mL)/streptomycin (100 μg/mL).

### Generation of SIRT4 expressing cell lines

HEK293 cell lines stably expressing SIRT4-eGFP from the pcDNA3.1 vector and mutants [enzymatically inactive SIRT4(H161Y)-eGFP and SIRT4(ΔN28)-eGFP lacking the N-terminal mitochondrial targeting signal] have been described ^42^ and cultured in media containing 800 μg/ml Geneticin/G418 (Genaxxon) as permanent selection agent. HEK293 and HeLa cell lines stably expressing SIRT4-eGFP from the retroviral vector puc2CL12IPwo were generated as described elsewhere ^74^ and further enriched by fluorescence activated cell sorting (FACS). Expression of SIRT4-eGFP fusion constructs was validated by immunoblotting and flow cytometry.

### Cell proliferation kinetics

HEK293 cells expressing eGFP, SIRT4-eGFP, or mutants thereof were seeded at 1.5×10^4^ cells / well in triplicates (6-well plates). Total numbers of viable cells were determined after 2 and 4 days using the TC10 cell counter (Bio-Rad).

### Live cell imaging

HEK293 cells expressing eGFP or SIRT4-eGFP (3×10^5^) were seeded on μ-Dish 35 mm plates (ibidi). For live cell imaging, cells were cultured in CO_2_-independent HEPES containing media (Life Technologies) at 37 °C in an isolated incubation chamber essentially as described ^75^. Cells were initially imaged at brightfield and 488 nm and thereafter only at brightfield every 12 min using a Nikon Eclipse TE2000-E microscope under control of the NIS Elements Advanced Research software (Nikon).

### Preparation of total cell lysates for immunoblot analysis

Cleared cell lysates were generated using lysis buffer containing either 0.3% CHAPS (3-[(3-Cholamidopropyl) dimethylammonio]-1-propanesulfonate) or 0.5% NP-40, 50 mM Tris-HCl (pH 7.4), 150 mM NaCl, 1 mM Na_3_VO_4_, 10 mM NaF, 1 mM EDTA, 1 mM EGTA, 2.5 mM Na_4_O_7_P_2_, 1 μM DTT, 1× cOmplete™ protease inhibitor cocktail (Sigma-Aldrich). Lysates were cleared by centrifugation (11.000 × g at 4°C for 20 min). Protein concentration of the supernatants was determined using the Bradford assay (K015.1, Roth). Cell lysates subjected to immunoblot analysis were obtained by lysing cells in lysis buffer containing 0.5% NP-40 (see above). Antibodies used for immunoblot analysis are listed in Table S2.

### Immunoprecipitation of GFP fusion proteins using the anti-GFP nanobody or standard immunoprecipitation protocols

The single-domain-anti-GFP antibody (“nanobody”) method ^76^ was employed to immunoprecipitate SIRT4-eGFP fusion proteins essentially as described^42^. Co-immunoprecipitation of α-tubulin interacting proteins was performed from total cell lysates using α-Tubulin specific antibodies (rabbit anti-α-tubulin, ab52899, Abcam) and Protein A/G Sepharose beads (Santa Cruz Biotechnology). Cell lysates subjected to immunoprecipitation were obtained by lysing cells in lysis buffer containing 0.3% CHAPS (see above).

### Subcellular fractionation analysis

Subcellular fractionation of total cell lysates was performed essentially as described ^77^ with additional centrifugation steps to obtain a cytosolic fraction together with a mitochondrially enriched particulate fraction. Cells were suspended in HEPES buffered solution [20 mM HEPES, pH 7.5; 220 mM mannitol; 70 mM sucrose; 1 mM EDTA; 1× protease inhibitor cocktail (Roche)] and mechanically lysed by repeatedly passing through 20 G syringe needles. The total cell lysate was centrifuged (600 × g, 10 min), and the resulting crude cytoplasmatic fraction without cellular debris was subjected to at least eight further centrifugation steps (600 × g, 1000 × g, 16.000 × g) thereby collecting mitochondria enriched pellets and a pure cytosolic fraction. Mitochondria containing pellets were resuspended in HEPES buffered solution containing 10 mM MgCl_2_ and 250 mM sucrose, centrifuged (12.000 × g, 10 min) twice through a sucrose cushion (HEPES buffered solution containing 0.5 mM MgCl_2_ and 880 mM sucrose). The resulting highly mitochondria enriched pellets were resuspended [40 mM Tris HCl, pH 7.5; 150 mM NaCl; 3% glycerol; 0.5 mM DTT; 1x protease inhibitor cocktail (Roche)] and analysed by SDS-PAGE. Antibodies used for immunoblot analysis of subcellular marker proteins are listed in suppl. Table 2.

### Microtubule pulldown experiments

Pelleting of Taxol-stabilized microtubules from cytosolic fractions was performed essentially as described ^78^. Asynchronously growing HEK293 cells expressing eGFP or SIRT4-eGFP were lysed in PHEM buffer [60 mM PIPES, 25 mM HEPES, 1mM EGTA, 1 mM magnesium acetate, pH 6.8; 1x cOmplete™ protease inhibitor cocktail (Sigma-Aldrich)] using a Dounce homogenizer. Following centrifugation (14.000 × g, 30 min) of the total cell lysate, the supernatant (cytosolic fraction) was supplemented with GTP (1 mM) and Paclitaxel/Taxol (20 μM) (both from Sigma-Aldrich). Samples were incubated at room temperature for 30 min and subjected to centrifugation (14.000g, 15 min) through a sucrose layer (15% sucrose in PHEM buffer) to obtain supernatant and the microtubules containing pellet fraction. The latter was washed one time in Taxol containing PHEM buffer, centrifuged, and sample fractions were analysed by SDS-PAGE.

### RO-3306 mediated G_2_ cell cycle arrest

Cells were treated for 14h with the CDK1 inhibitor RO-3306 (Selleckchem; 10 μM) to achieve synchronization at G_2_. When indicated, cells were released into mitosis by one time washing and addition of fresh media, harvested 45 min later, and analysed as indicated.

### Mass spectrometric analysis of the mitotic SIRT4 interactome

Sample preparation for proteomic analysis, LC-MS analysis, computational mass spectrometric data analysis, and gene ontology/protein network analysis are specified in the Supplementary Materials and Methods section. Primary data obtained from mass spectrometric analysis of SIRT4-eGFP interacting proteins are listed in Table S1.

### Confocal laser scanning microscopy and signal quantification using ImageJ software

Cells were fixed in 4% paraformaldehyde for 20 min and permeabilized with 0.2% Triton X-100 for 20 min followed by a blocking step with 4% BSA/0.05% saponin for 30 min at room temperature. Alternatively, for spinning disk confocal analysis, cells were fixed in 4% paraformaldehyde for 20 min and permeabilized with 0.25% Triton X-100 for 5 min followed by a blocking step with 3% BSA in PBS (phosphate buffered saline) for two hours at room temperature. Cells were stained with primary antibodies in 1% BSA in PBS overnight at 4°C. All primary and secondary antibodies used for confocal imaging analysis are listed in Table S3. In particular, SIRT4 was detected using antibodies from Sigma-Aldrich (SAB1407208; mouse monoclonal; a.a. 1-314 of human SIRT4 as epitope; 1:500) and Santa Cruz Biotechnology (sc-135053; rabbit polyclonal; a.a. 81-314 of human SIRT4 as epitope). DNA was detected by DAPI staining followed by mounting of coverslips with ProLong Gold antifade reagent (Invitrogen, P36934). Analyzes were performed with a LSM510-Meta confocal microscope (Zeiss) equipped with 40/1.3 immersion objectives and excitation wavelengths of 468 nm, 488 nm, 543 nm, and 633 nm. In addition, an UltraVIEW spinning disk confocal microscope (Perkin Elmer, Waltham, MA, USA) with excitation wavelengths of 405 nm, 488 nm, 561 nm, and 633 nm, a 60x/1.4 NA oil objective, and the Volocity 6.3 software (Perkin Elmer) was employed. To increase detection of SIRT4-eGFP fusion proteins, primary antibodies against GFP (Nacalai Tesque, Inc., GF090R; 1:1000) were employed in spinning disk confocal microscopy when indicated. Image processing and quantification of centrosomal SIRT4 and Pericentrin signal intensities were performed based on ImageJ software v1.49k.

### Statistical analysis

Data are presented as mean ± s.d. Multiple comparisons were analyzed by one-way analysis of variance (ANOVA) followed by Tukey’s post hoc test using the GraphPad Prism software.

## Acknowledgements

We thank Nick Stoecklein for providing Colo-680N cells, Hakima Ezzahoini for technical assistance, and Katharina Raba (Core Flow Cytometry Facility, ITZ) for FACS sorting of stable cell lines. This work was funded by the Stiftung für Altersforschung (grant 701.810.783) of the Heinrich Heine-University Düsseldorf (to R.P.P.) and by the research commission (grants 9772643 and 9772690) of the Medical Faculty of the Heinrich Heine-University Düsseldorf (to R.P.P. and M.R.A.).

## Author Contributions

L.B., A.L., S. A-H., and R.P.P. initiated the project. L.B., A.L., and R.P.P designed the study. L.B., A.L., C.B., S. A-H., I.F., P.V., A.K., and R.P.P designed, performed, and analyzed the experiments. N.O., A.S., and K.S. obtained and analyzed the mass spectromectric data. C.W. and H.H. provided expertise and generated retrovirally transduced cell line models. C.M., D.S., and R.U.J. provided expertise and tools for microscopic and cell cycle analysis. J.S., A.S.R., and M.R.A. provided essential reagents and methods for protein complex and mitochondrial analysis and critically corrected the manuscript. L.B. and R.P.P. wrote the manuscript. All authors read, discussed, and approved the final version of the manuscript.

## Additional Information

**Supplementary Information** including supplementary Tables, Figures, and Movies accompanies this paper.

## Competing Interests

The authors declare no competing interests.

