## Supplementary Information for "Subcellular localization and mitotic interactome analyses identify SIRT4 as a centrosomally localized and microtubule associated protein"

### Supplementary Materials and Methods

**Expression constructs and generation of stable cell lines.** Plasmids for eukaryotic expression of TUBGCP2 (pCMV6-GCP2-Flag) and TUBGCP3 (pCMV6-GCP3-Flag) were obtained from Sino Biological Inc. Expression plasmids for C-terminally Flag-tagged Sirtuins were obtained from Origene (pCMV6-SIRT4-Myc-Flag) and Addgene (pcDNA3.1-SIRT3-Flag: #13814; pcDNA3.1-SIRT5-Flag: #13816; [1]). HEK293 cell lines stably expressing flagged SIRT isoforms were generated using standard transfection protocols followed by selection [400 µg/ml Geneticin/G418 (Genaxxon)]. Cell lines were passaged in media containing 400 µg/ml Geneticin/G418 as permanent selection agent.

**Sample preparation for proteomic analysis.** Proteins were extracted from frozen cell pellets as described elsewhere[2]. Briefly, cells were lysed and homogenized in urea buffer with a TissueLyser (Qiagen, Hilden, Germany) and supernatants were collected after centrifugation for 15 min at 14.000 x g and 4°C. Protein concentration was determined using the Pierce 660 nm Protein Assay (Fischer Scientific, Schwerte, Germany) and 10 µg protein per sample were loaded on a SDS-PAGE for in-gel-digestion. The isolated gel pieces were reduced (50 µl, 10 mM DTT), alkylated (50 µl, 50 mM iodoacetamide), and underwent afterwards tryptic digestion (6 µl, 200 ng trypsin in 100 mM ammonium bicarbonate). The peptides were resolved in 15 µl 0.1 % trifluoracetic acid and subjected to liquid chromatography.

**LC-MS analysis.** For the LC-MS analysis a QExactive plus (Thermo Scientific, Bremen, Germany) connected with an Ultimate 3000 Rapid Separation liquid chromatography system (Dionex / Thermo Scientific, Idstein, Germany) equipped with an Acclaim PepMap 100 C18 column (75 µm inner diameter, 25 cm length, 2 mm particle size from Thermo Scientific, Bremen, Germany) was applied. The length of the isocratic LC gradient was 120 minutes. The mass spectrometer was operating in positive mode and coupled with a nano electrospray ionization source. Capillary temperature was set to 250°C and source voltage to 1.4 kV. In the QExactive plus mass spectrometer for the survey scans a mass range from 200 to 2000 m/z at a resolution of 70,000 was used. The automatic gain control was set to 3.000.000 and the maximum fill time was 50 ms. The 10 most intensive peptide ions were isolated and fragmented by high-energy collision dissociation (HCD).

**Computational mass spectrometric data analysis.** Peptide and protein identification and quantification was done by using MaxQuant (version 1.5.3.8, MPI for Biochemistry, Planegg, Germany) applying standard parameters. As human samples were analyzed, searches were conducted using a specific proteome database (UP000005640, downloaded 06/20/16) from UniProt. Methionine oxidation and acetylation at protein N-termini were set as variable modification and carbamidomethylations at cysteines were considered as fixed modification. Peptides and proteins were accepted with a false discovery rate set to 1%. Unique and razor peptides were used for label-free quantification and peptides with variable modifications were included in the quantification. The minimal ratio count was set to two and the matched between runs option was enabled.

The normalized intensities as provided by MaxQuant were analyzed by using Perseus framework (version 1.5.0.15, MPI for Biochemistry, Planegg, Germany). Only proteins containing at least two unique peptides and a minimum of 3 valid values in each group were taken into consideration for protein quantification. Proteins which were identified only by site or marked as contaminant (from the MaxQuant contaminant list) were excluded from the analysis. For the calculation of enriched proteins in the two groups Student's t-tests were applied. The significance analysis was applied on $\log_2$ transformed values using a S0 constant = 0 and a 1 % false discovery rate-based cutoff. The mass spectrometry proteomics data have been deposited to the ProteomeXchange Consortium *via* the PRIDE partner repository[3] with the data set identifier PXD017319. SIRT4-interacting proteins were functionally grouped and subjected to gene ontology and pathway/network analysis using the ClueGO software[4].

### Supplementary Tables

**Table S1.** Differential analysis of SIRT4-interacting proteins in mitotically synchronized HEK293 cells stably expressing SIRT4-eGFP as compared to eGFP expressing control cells.

**Table S2.** List of antibodies used for immunoblot analysis.

**Table S3.** List of antibodies used for confocal imaging analysis.

### Supplementary Figure legends

**Fig. S1.** SIRT4 co-localizes with Pericentrin at the centrosome during interphase. Endogenous SIRT4 was detected in HT1080 fibrosarcoma cells using spinning disk microscopy and a monoclonal antibody against SIRT4 (SAB1407208, Sigma-Aldrich). Antibodies against Pericentrin and $\alpha$-tubulin were employed to visualize centrosomes and microtubules, respectively. DAPI was used to detect DNA. Colocalization of SIRT4 and Pericentrin is indicated by white arrows, whereas colocalizing MTOCs (microtubule organizing centers) are marked by yellow arrows. Bar: 10 µm.

**Fig. S2.** SIRT4 co-localizes with Pericentrin at centrosomes. Endogenous SIRT4 was detected in Colo-680N esophagus squamous carcinoma cells using standard confocal microscopy (cLSM510-Meta, Zeiss) and a monoclonal antibody against SIRT4 (SAB1407208, Sigma-Aldrich). Antibody staining against Pericentrin was employed to visualize centrosomes. DAPI was used to detect DNA. Bar: 5 µm.

**Fig. S3.** SIRT4 co-localizes with the mitotic spindle marker TACC3. (**a**) Endogenous SIRT4 was detected in HEK293 cells using standard confocal microscopy (cLSM510-Meta, Zeiss) and a monoclonal antibody against SIRT4 (SAB1407208, Sigma-Aldrich). Antibody staining against TACC3 was employed to visualize the mitotic and central spindle. DAPI was used to detect DNA. Bar: 10 µm. (**b**) Magnifications (2.6-fold) of cells with SIRT4 localizing at the interphase centrosome (upper panel) as well as at the mitotic and central spindle (middle and lower panels, respectively).

**Fig. S4.** Detection of SIRT4 at the mitotic spindle apparatus and in mitochondria. Endogenous SIRT4 was detected in HEK293 cells using standard confocal microscopy (cLSM510-Meta, Zeiss) and a polyclonal antibody against SIRT4 (sc-135053, Santa Cruz Biotechnology). Antibodies against MTC02 and $\alpha$-tubulin were employed to visualize mitochondria and microtubules, respectively. DAPI was used to detect DNA. Magnifications of SIRT4 co-localizing with either MTC02 (mitochondria) or $\alpha$-tubulin (mitotic spindle) are depicted and indicated by arrows. Bar: 10 µm.

**Fig. S5.** SIRT3 localizes in mitochondria, but is absent from centrosomes. Endogenous SIRT3 was detected in HeLa cells using confocal microscopy (cLSM510-Meta, Zeiss) and a rabbit monoclonal antibody against SIRT3 (#5490, Cell Signaling). MTC02 and γ-tubulin co-stainings were employed to visualize mitochondria and centrosomes, respectively. DAPI was used to detect DNA. Bar: 20 µm (interphase) and 10 µm (mitosis).

**Fig. S6.** Subcellular localization of SIRT4-eGFP (upper panels) and SIRT4(ΔN28)-eGFP (lower panels) in transiently transfected HT1080 fibrosarcoma cells (interphase) as imaged by spinning disk microscopy. Antibodies against Pericentrin were employed to visualize centrosomes. DAPI was used to detect DNA. Bars: 10 µm.

**Fig. S7.** Subcellular fractionation analysis of ectopically expressed SIRT4-eGFP in HEK293 cells. Total cell lysates (TCL; 80 µg) and the respective mitochondrially enriched (Mito-enriched; 50 µg) and cytosolic (Cyto; 80 µg) fractions were subjected to immunoblot analysis. Subcellular marker proteins detected were OPA1 (long and short form; mitochondria), α-tubulin (cytoplasm), and HDAC6 (predominantly cytoplasmatic localized) (**a**) as well as Na/K ATPase as cell membrane marker (**b**). Antibodies against eGFP were employed to visualize eGFP and SIRT4-eGFP.

**Fig. S8.** Subcellular fractionation analysis of ectopically expressed sirtuin proteins. C-terminally Flag-tagged sirtuin isoforms SIRT3, SIRT4, and SIRT5 were ectopically expressed in HEK293 cells. Total cell lysates (TCL; 60 µg) and the respective mitochondrially enriched (Mito-enriched; 60 µg) and cytosolic (Cyto; 60 µg) fractions were subjected to immunoblot analysis. Subcellular marker proteins detected were OPA1 (long and short form; mitochondria) and ATP5A1 (mitochondria), α-tubulin (cytoplasm), and HDAC6 (predominantly cytoplasmatic localized).

**Fig. S9.** HEK293 cells ectopically expressing SIRT4(ΔN28)-eGFP display an increased percentage of polyploidy. (**a**) Quantification of bi- or multinucleated HEK293 cells expressing SIRT4-eGFP or mutants thereof. The total numbers of cells analysed and the percentage of bi- or multinucleated cells are indicated (**b**). Representative pictures of polyploid SIRT4(ΔN28)-eGFP expressing HEK293 cells are depicted. Bar: 20 or 25 µm.

**Fig. S10.** Anti-eGFP nanobody based immunoprecipitation analysis of mitotic SIRT4-eGFP interactors. HEK293 cells stably expressing SIRT4-eGFP, mutants thereof [enzymatically inactive SIRT4(H161Y)-eGFP or SIRT4(ΔN28)-eGFP lacking the N-terminal mitochondrial targeting signal], or eGFP as control were analysed. α-tubulin, γ-tubulin, but not Pericentrin co-immunoprecipitate with SIRT4-eGFP.

**Fig. S11.** Network analysis of the SIRT4-interactom of mitotically synchronized HEK293 cells using the ClueGO software.

**Fig. S12.** SIRT4-eGFP interacts and subcellularly colocalizes with the γTUSC components GCP2 and GCP3. (**a**) Upper panel: Anti-eGFP nanobody based immunoprecipitation analysis of SIRT4-eGFP interaction with ectopically expressed GCP2-Flag in HEK293 cells (upper panel). SIRT4 co-localizes with GCP2 at the mitotic spindle apparatus (lower panel). Endogenous SIRT4 and GCP2 were detected in HT1080 cells using standard confocal microscopy (cLSM510-Meta, Zeiss) and staining with a polyclonal antibody against SIRT4 (sc-135053, Santa Cruz Biotechnology) and a monoclonal antibody against GCP2 (sc. 377117, Santa Cruz Biotechnology). Microtubules were detected by α-Tubulin staining. DAPI was used to visualize DNA. Bar: 5 µm. (**b**) Upper panel: Anti-eGFP nanobody based immunoprecipitation analysis of SIRT4-eGFP interaction with endogenous GCP3 in HEK293 cells (upper panel). SIRT4 co-localizes with GCP3 at the mitotic spindle apparatus. Endogenous SIRT4 and GCP3 were detected in HT1080 cells using standard confocal microscopy (cLSM510-Meta, Zeiss) and staining with a polyclonal antibody against SIRT4 (sc-135053, Santa Cruz Biotechnology) and a monoclonal antibody against GCP3 (sc. 373758, Santa Cruz Biotechnology). Microtubules were detected by α-Tubulin staining. DAPI was used to visualize DNA. Bar: 5 µm.

**Supplementary Movies**

**Suppl. Movie 1:** Subcellular localization of SIRT4-eGFP in transiently transfected HeLa cells (interphase) as imaged by spinning disk microscopy. Antibodies against Pericentrin and $\alpha$-tubulin were employed to visualize centrosomes and microtubules, respectively. DAPI was used to detect DNA. Lower panels: Movies embedded as GIF files into PowerPoint.

**Suppl. Movie 2:** Subcellular localization of SIRT4-eGFP in transiently transfected HeLa cells (metaphase) as imaged by spinning disk microscopy. To increase detection of the SIRT4-eGFP fusion protein a primary antibody against GFP was employed. Antibodies against Pericentrin and $\alpha$-tubulin were employed to visualize centrosomes and microtubules, respectively. DAPI was used to detect DNA. Lower panels: Movies embedded into PowerPoint.

**Suppl. Movie 3:** Subcellular localization of eGFP in transiently transfected HeLa cells (metaphase) as imaged by spinning disk microscopy. To increase detection of the eGFP protein a primary antibody against GFP was employed. Antibodies against Pericentrin and $\alpha$-tubulin were employed to visualize centrosomes and microtubules, respectively. DAPI was used to detect DNA. Lower panels: Movies embedded as GIF files into PowerPoint.

**Suppl. Movie 4:** Subcellular localization of SIRT4-eGFP in in transiently transfected HeLa cells (telophase/cytokinesis) as imaged by spinning disk microscopy. To increase detection of the SIRT4-eGFP fusion protein a primary antibody against GFP was employed. Antibodies against Pericentrin and $\alpha$-tubulin were employed to visualize centrosomes and microtubules, respectively. DAPI was used to detect DNA. Lower panels: Movies embedded as GIF files into PowerPoint.

**Suppl. Movie 5:** Subcellular localization of SIRT4($\Delta$N28)-eGFP in in transiently transfected HeLa cells (interphase) as imaged by spinning disk microscopy. Antibodies against Pericentrin and $\alpha$-tubulin were employed to visualize centrosomes and microtubules, respectively. DAPI was used to detect DNA. Lower panels: Movies embedded as GIF files into PowerPoint.

**Suppl. Movie 6:** SIRT4 and HDCA6 partially colocalize in mitotic cells as imaged by spinning disk microscopy. Antibodies to detect endogenous SIRT4 and HDAC6 were from Sigma-Aldrich (SAB1407208; mouse monoclonal) and Cell Signaling (#7558), respectively. Antibodies against $\alpha$-tubulin were employed to visualize microtubules. DAPI was used to detect DNA. Lower panels: Movies embedded as GIF files into PowerPoint.
