## Supplementary Figures for "Subcellular localization and mitotic interactome analyses identify SIRT4 as a centrosomally localized and microtubule associated protein"

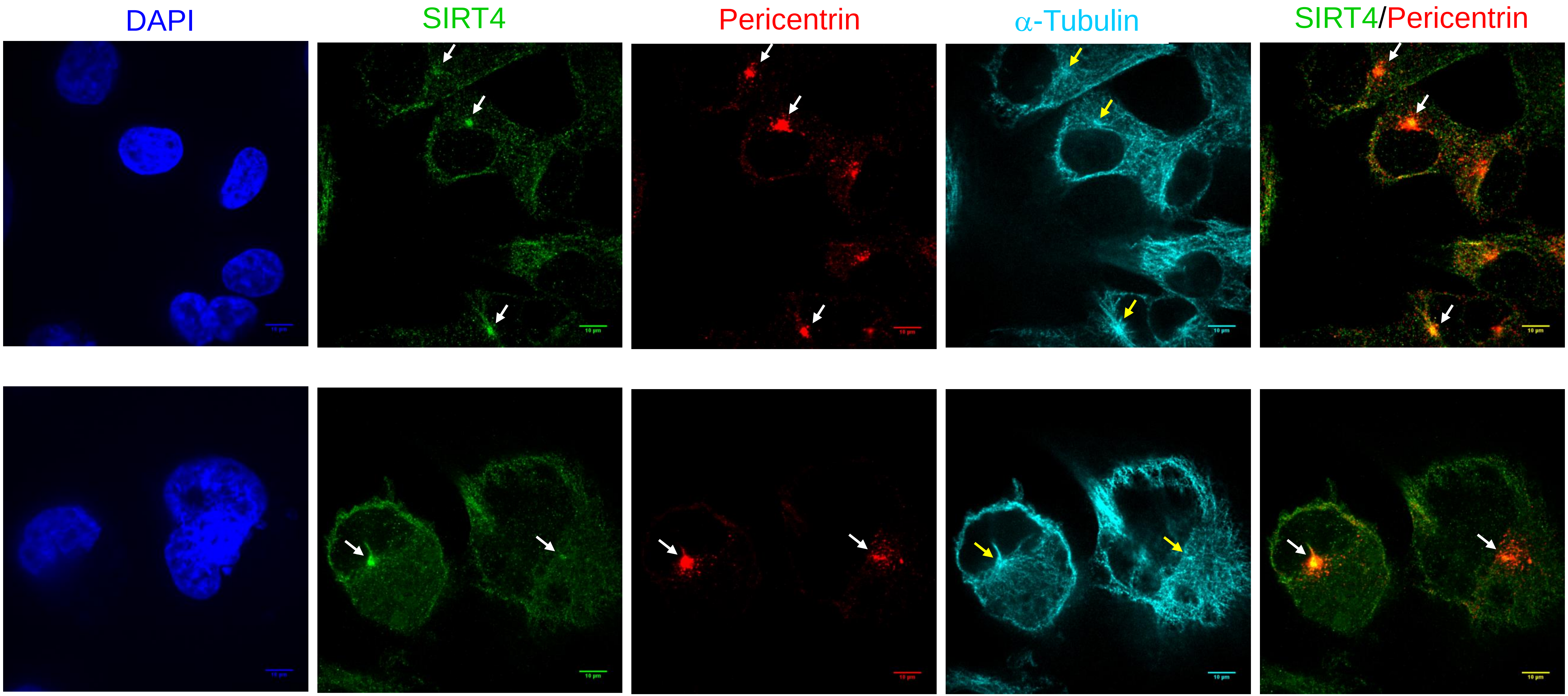


Fig. S1

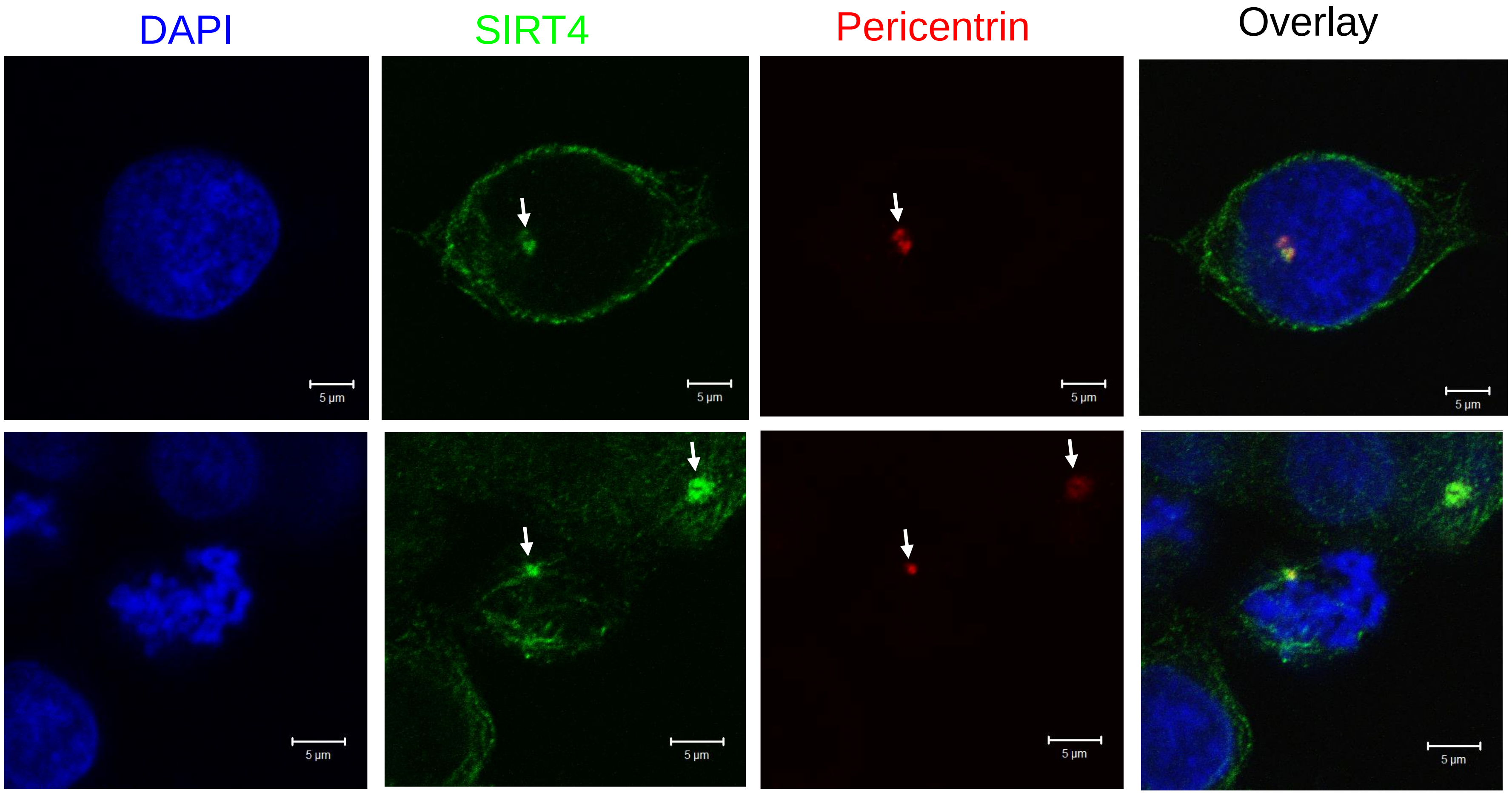


Fig. S2

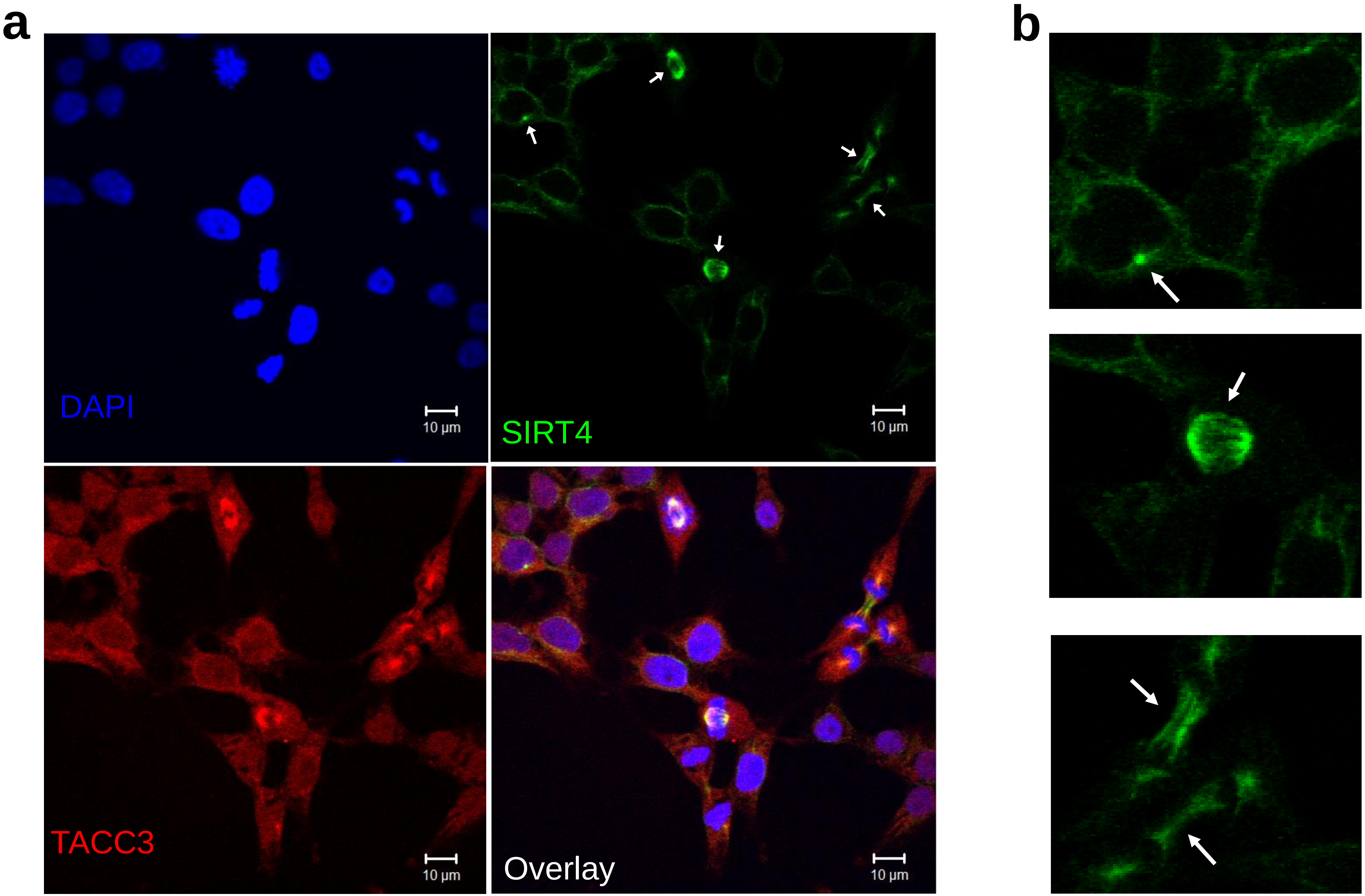


Fig. S3

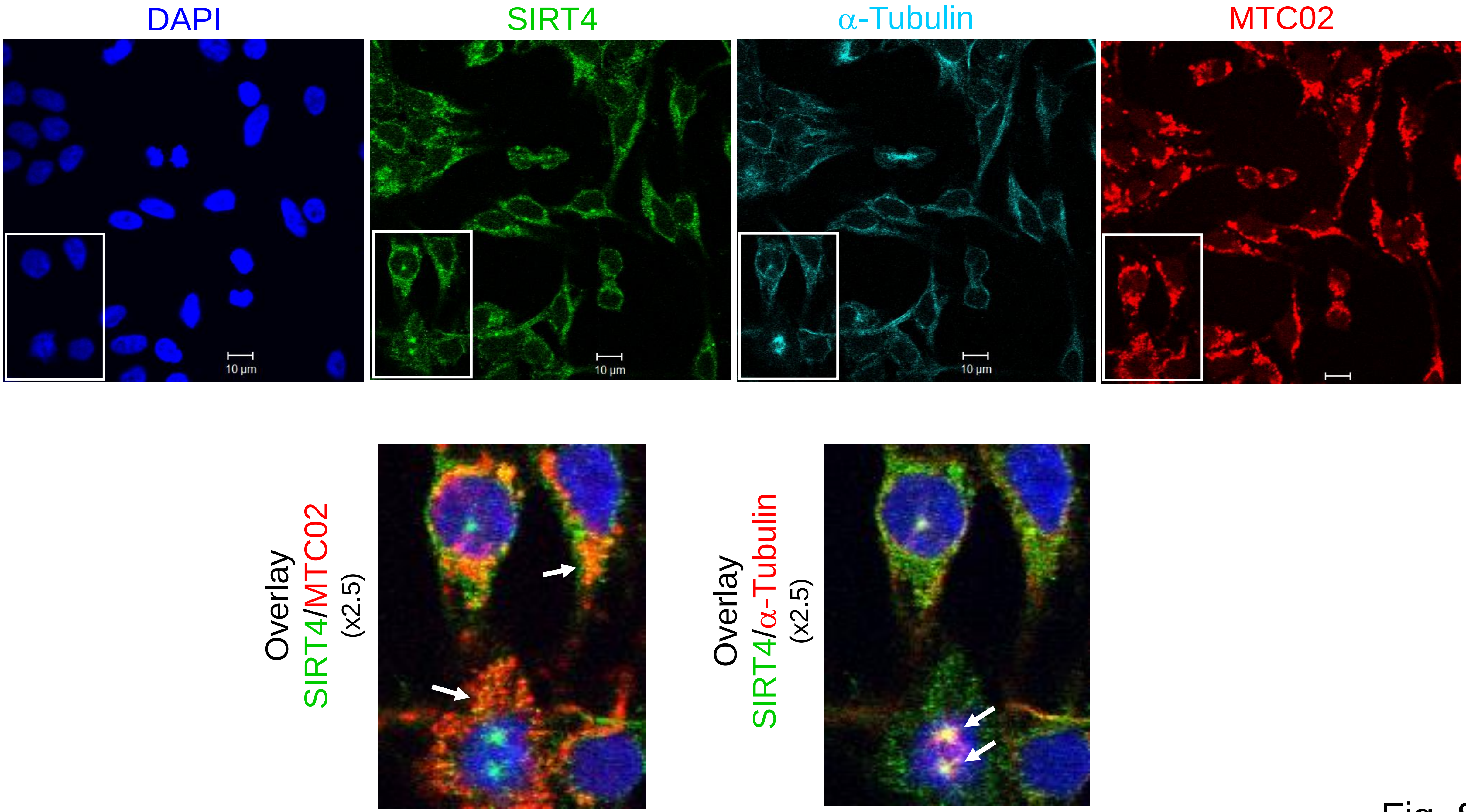
DAPI
SIRT4
α-Tubulin
MTC02
10 µm
10 µm
10 µm
Overlay
SIRT4/MTC02
(x2.5)
Overlay
SIRT4/α-Tubulin
(x2.5)

Fig. S4

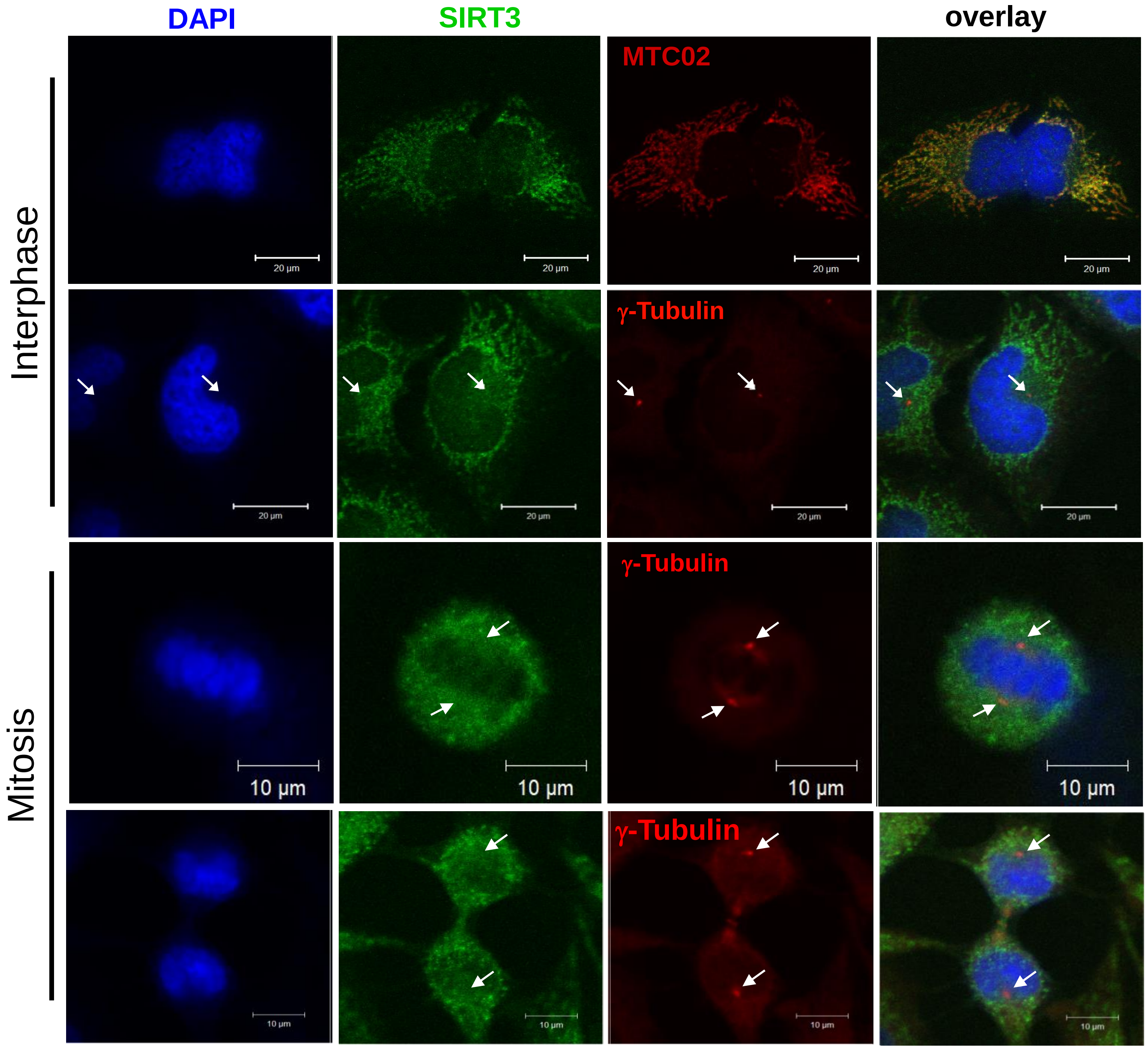
DAPI
SIRT3
overlay
MTC02
γ-Tubulin
γ-Tubulin
γ-Tubulin
Interphase
Mitosis
20 µm
10 µm

Fig. S5

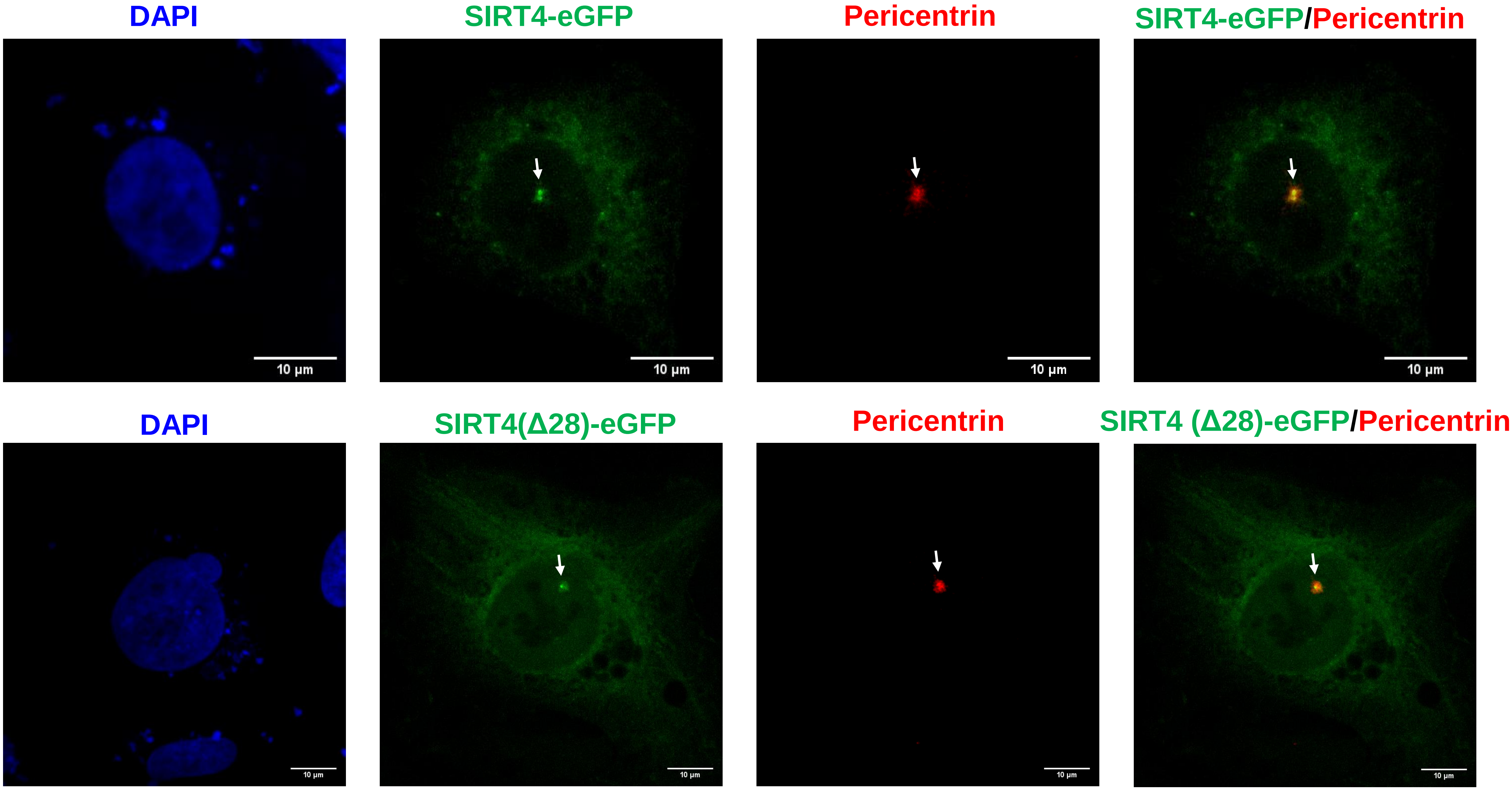
DAPI
SIRT4-eGFP
Pericentrin
SIRT4-eGFP/Pericentrin
10 µm
10 µm
10 µm
10 µm
DAPI
SIRT4(Δ28)-eGFP
Pericentrin
SIRT4 (Δ28)-eGFP/Pericentrin
10 µm
10 µm
10 µm
10 µm

Fig. S6

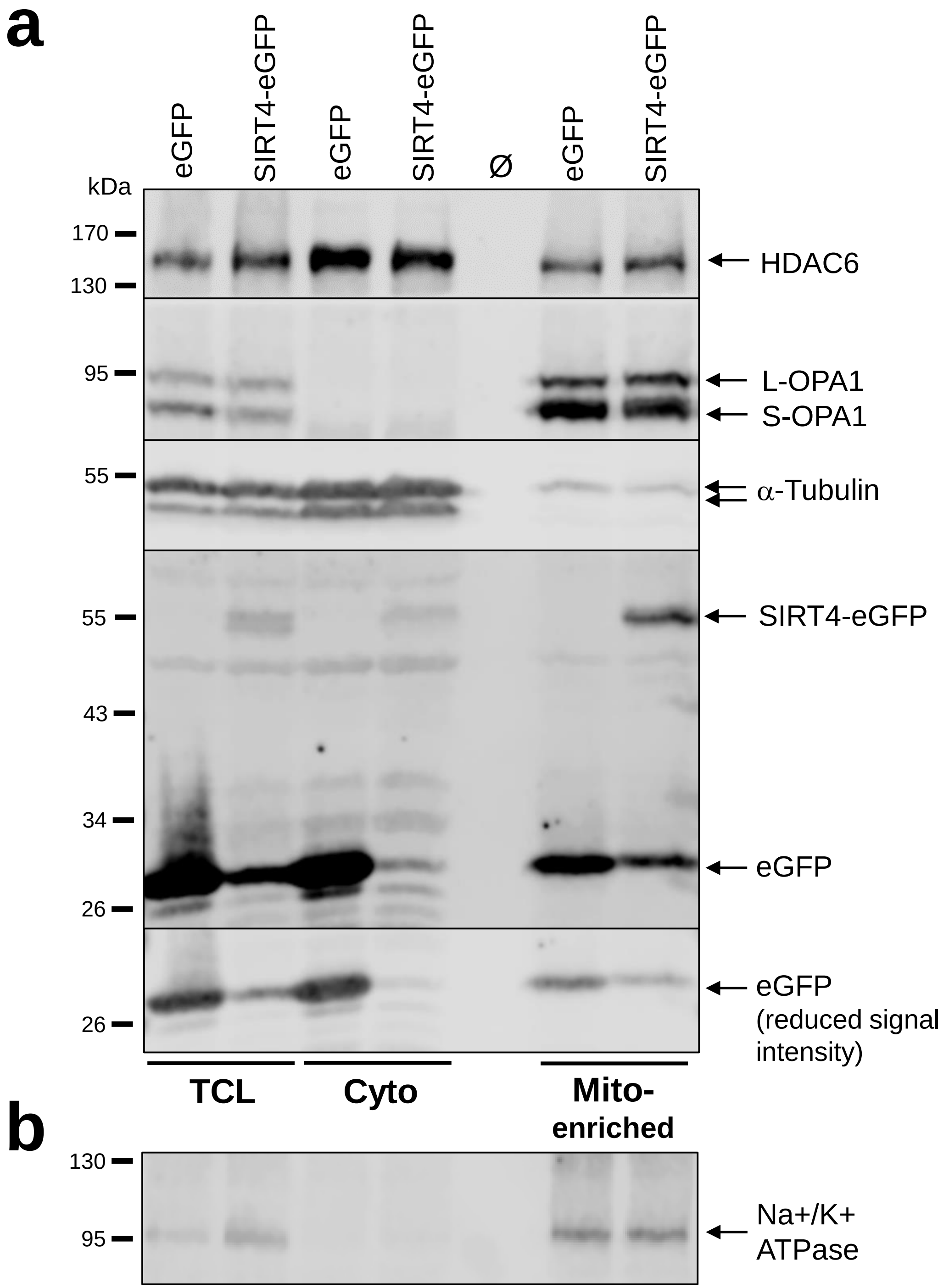
a
eGFP
SIRT4-eGFP
eGFP
SIRT4-eGFP
Ø
eGFP
SIRT4-eGFP
kDa
170
130
95
55
55
43
34
26
26
HDAC6
L-OPA1
S-OPA1
α-Tubulin
SIRT4-eGFP
eGFP
eGFP
(reduced signal intensity)
TCL
Cyto
Mito-enriched
b
130
95
Na+/K+ ATPase

Fig. S7

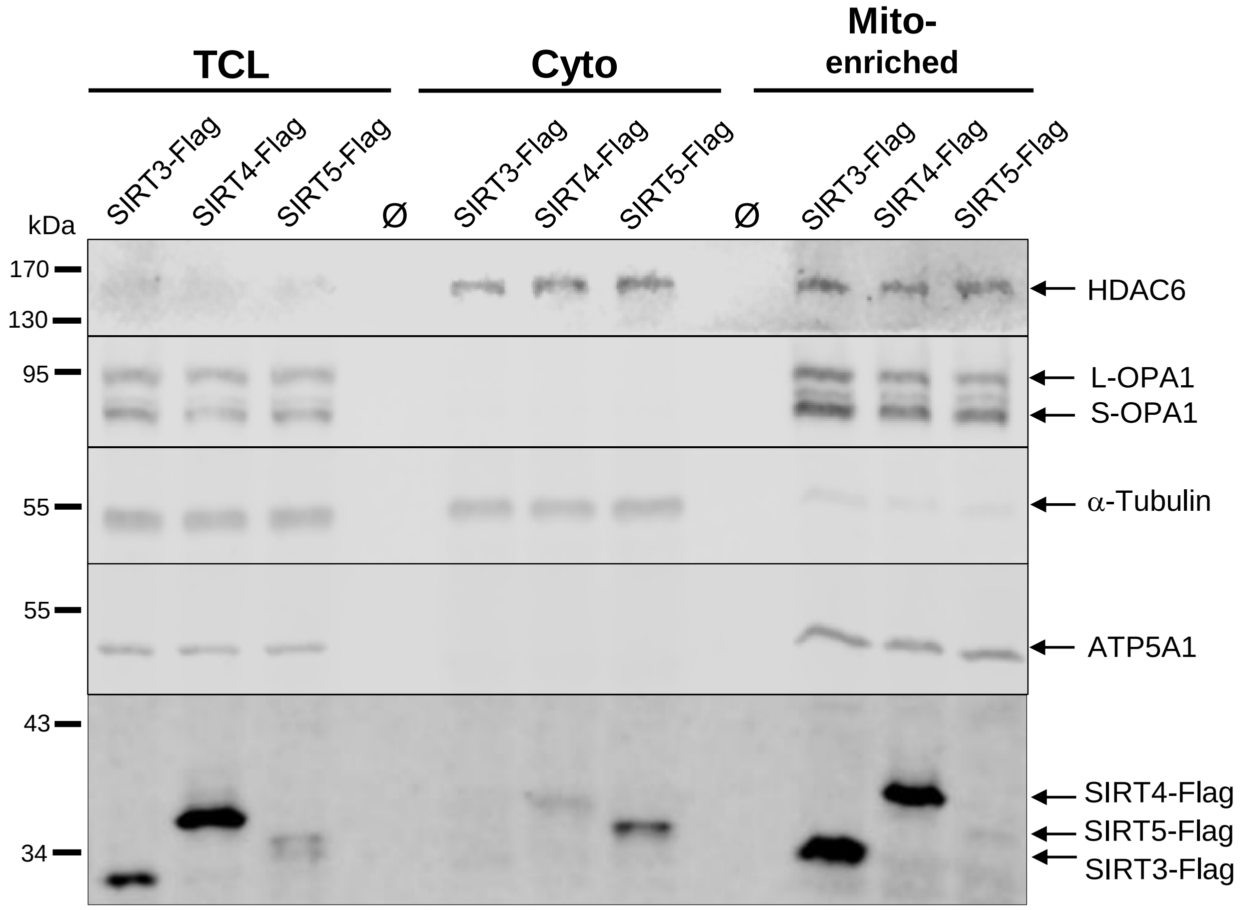

TCL
Cyto
Mito-
enriched
SIRT3-Flag
SIRT4-Flag
SIRT5-Flag
Ø
SIRT3-Flag
SIRT4-Flag
SIRT5-Flag
Ø
SIRT3-Flag
SIRT4-Flag
SIRT5-Flag
kDa
170
130
95
55
55
43
34
HDAC6
L-OPA1
S-OPA1
α-Tubulin
ATP5A1
SIRT4-Flag
SIRT5-Flag
SIRT3-Flag


Fig. S8

**a**

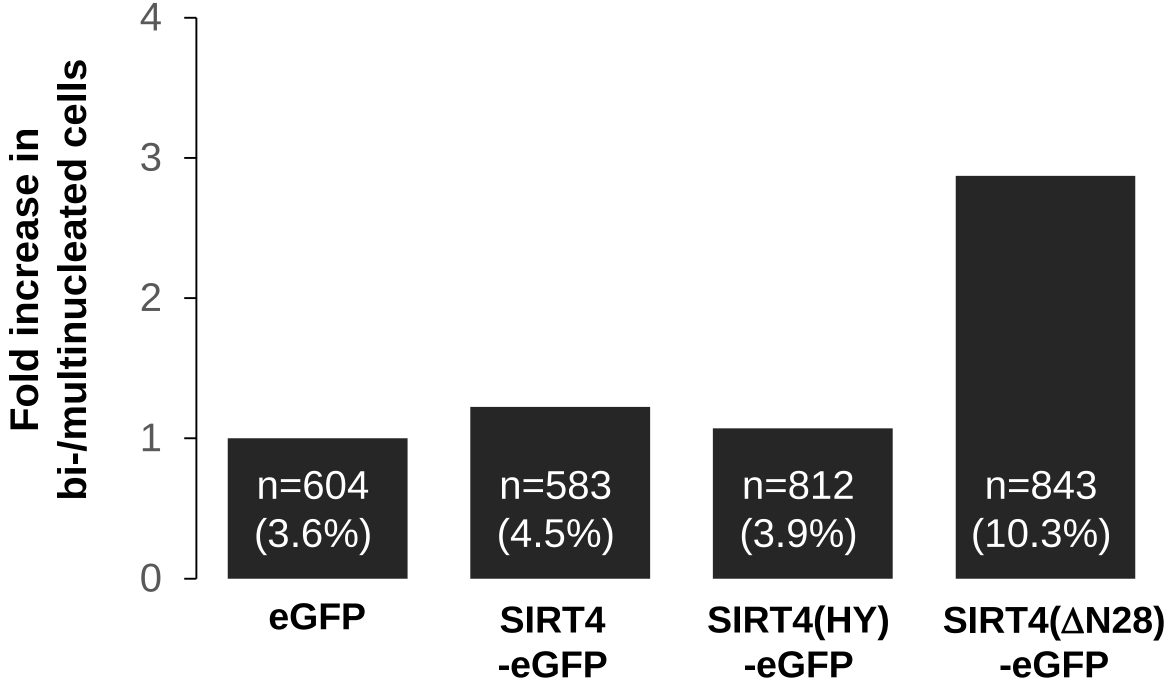

Fold increase in bi-/multinucleated cells
4
3
2
1
0
n=604 (3.6%)
n=583 (4.5%)
n=812 (3.9%)
n=843 (10.3%)
eGFP
SIRT4 -eGFP
SIRT4(HY) -eGFP
SIRT4(ΔN28) -eGFP


**b**

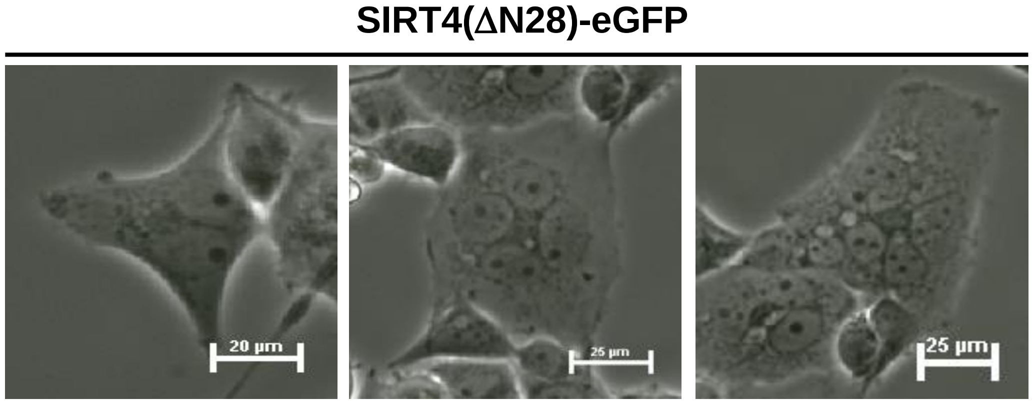

SIRT4(ΔN28)-eGFP
20 µm
25 µm
25 µm


Fig. S9

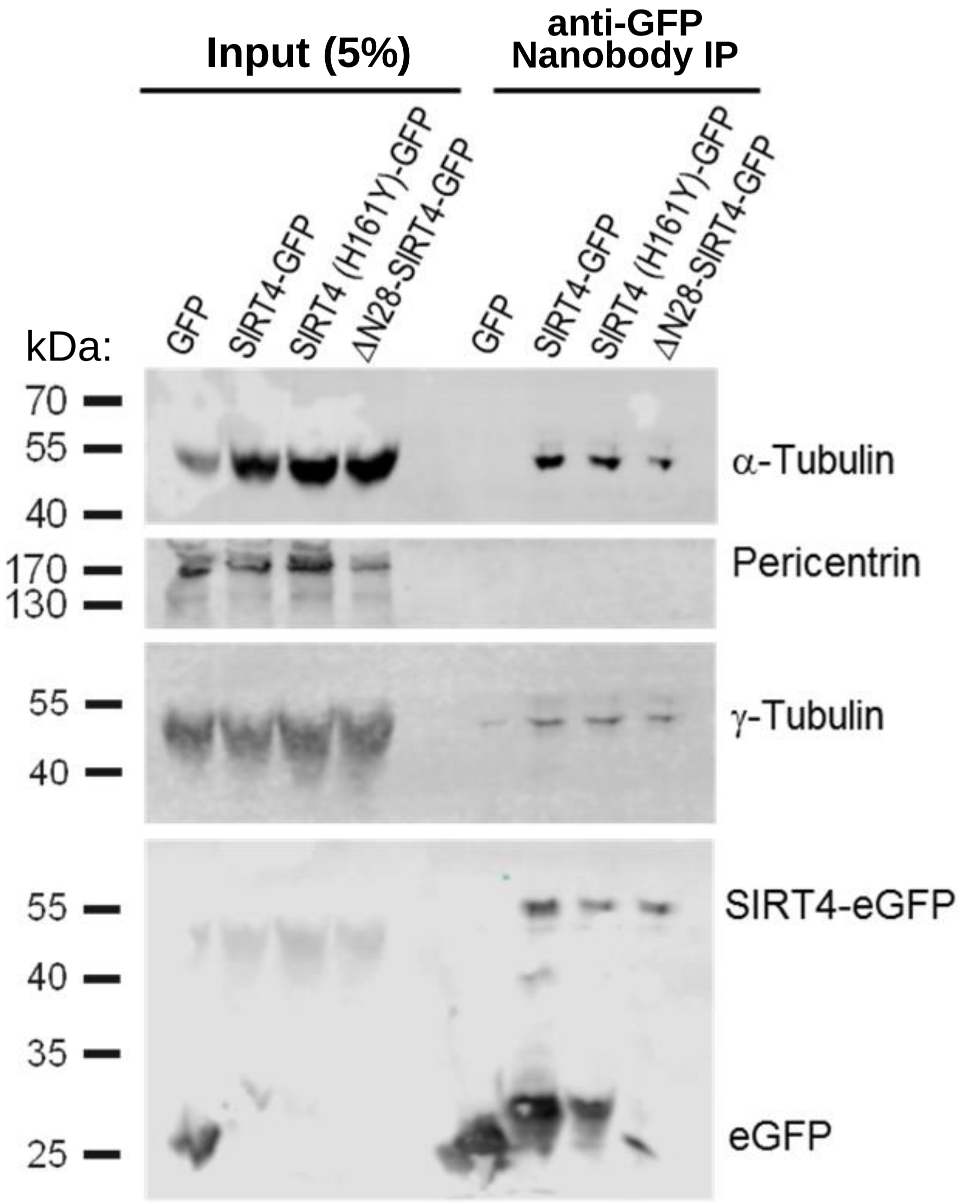

Input (5%)
anti-GFP
Nanobody IP
GFP
SIRT4-GFP
SIRT4 (H161Y)-GFP
ΔN28-SIRT4-GFP
GFP
SIRT4-GFP
SIRT4 (H161Y)-GFP
ΔN28-SIRT4-GFP
kDa:
70
55
40
170
130
55
40
55
40
35
25
α-Tubulin
Pericentrin
γ-Tubulin
SIRT4-eGFP
eGFP


Fig. S10

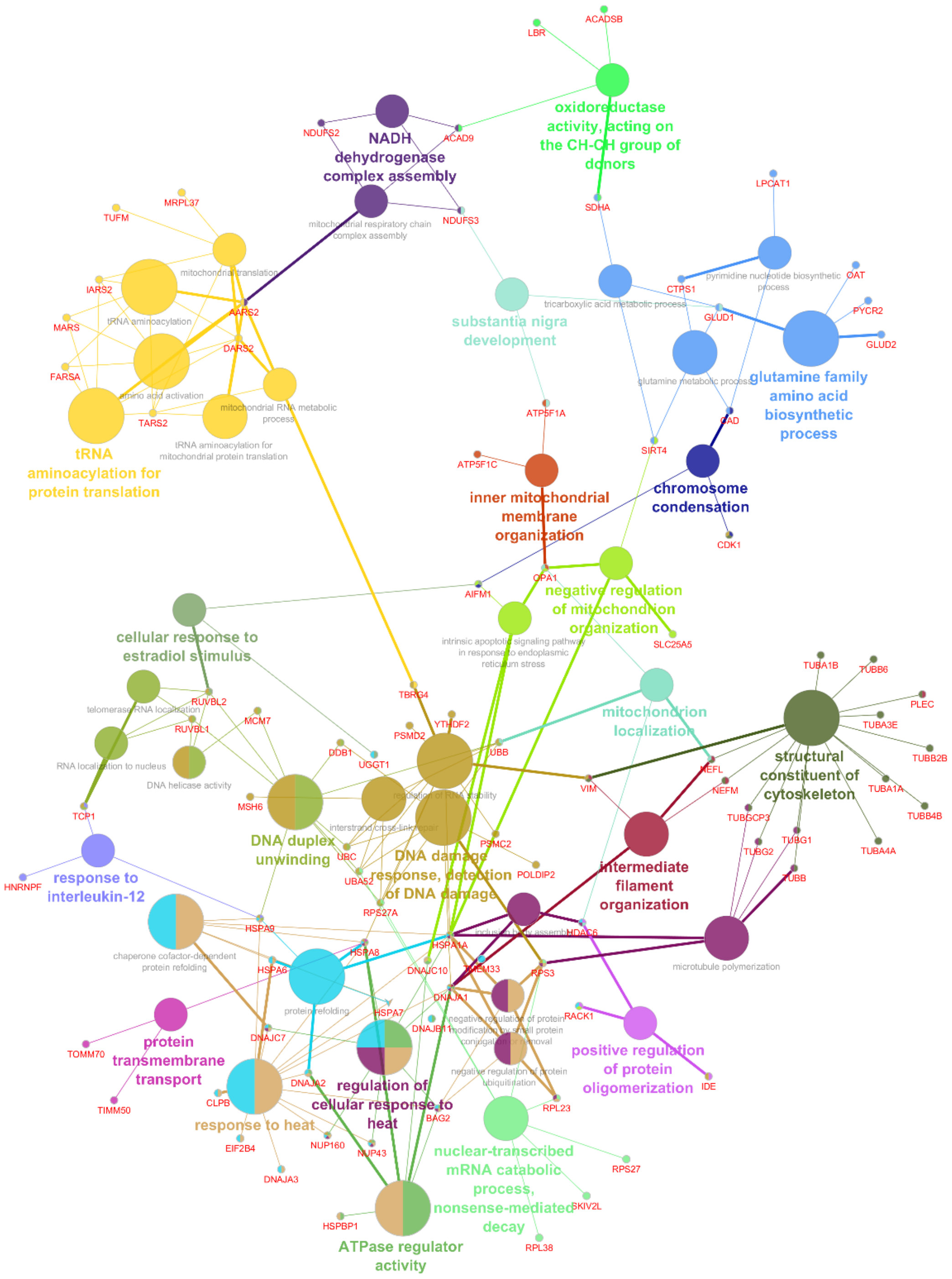
oxidoreductase activity, acting on the CH-CH group of donors
NADH dehydrogenase complex assembly
mitochondrial respiratory chain complex assembly
mitochondrial translation
tRNA aminoacylation
amino acid activation
mitochondrial RNA metabolic process
tRNA aminoacylation for mitochondrial protein translation
tRNA aminoacylation for protein translation
substantia nigra development
tricarboxylic acid metabolic process
pyrimidine nucleotide biosynthetic process
glutamine metabolic process
glutamine family amino acid biosynthetic process
inner mitochondrial membrane organization
chromosome condensation
negative regulation of mitochondrion organization
intrinsic apoptotic signaling pathway in response to endoplasmic reticulum stress
cellular response to estradiol stimulus
telomerase RNA localization
RNA localization to nucleus
DNA helicase activity
mitochondrion localization
structural constituent of cytoskeleton
DNA duplex unwinding
interstrand cross-link repair
DNA damage response, detection of DNA damage
intermediate filament organization
response to interleukin-12
chaperone cofactor-dependent protein refolding
inclusion body assembly
microtubule polymerization
protein refolding
protein transmembrane transport
regulation of cellular response to heat
response to heat
negative regulation of protein modification by small protein conjugation or removal
negative regulation of protein ubiquitination
positive regulation of protein oligomerization
nuclear-transcribed mRNA catabolic process, nonsense-mediated decay
ATPase regulator activity

Fig. S11

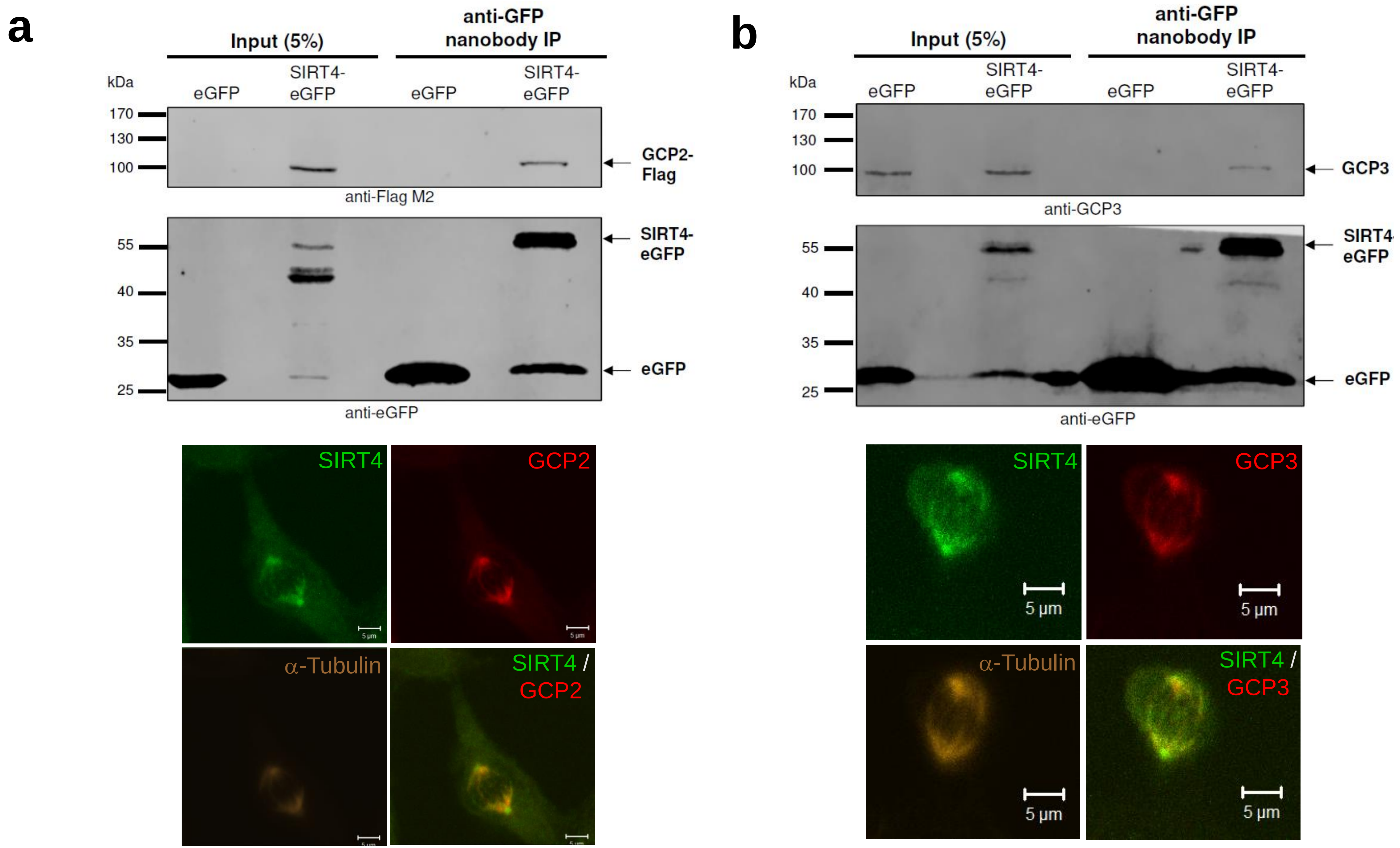
a
Input (5%)
anti-GFP nanobody IP
kDa
eGFP
SIRT4-eGFP
eGFP
SIRT4-eGFP
170
130
100
GCP2-Flag
anti-Flag M2
55
40
35
25
SIRT4-eGFP
eGFP
anti-eGFP
SIRT4
GCP2
α-Tubulin
SIRT4 / GCP2
b
Input (5%)
anti-GFP nanobody IP
kDa
eGFP
SIRT4-eGFP
eGFP
SIRT4-eGFP
170
130
100
GCP3
anti-GCP3
55
40
35
25
SIRT4-eGFP
eGFP
anti-eGFP
SIRT4
GCP3
α-Tubulin
SIRT4 / GCP3
5 µm
5 µm
5 µm
5 µm

Fig. S12
