## Supplementary material for "Subcellular localization and mitotic interactome analyses identify SIRT4 as a centrosomally localized and microtubule associated protein": Table S1

| Significant | Protein IDs | Majority prot | Score | Intensity | MS/MS Count | Unique peptic | t-test p value | t-test Differer | SIRT4-eGFP #1 | SIRT4-eGFP #2 | SIRT4-eGFP #3 | SIRT4-eGFP #4 | eGFP #1 | eGFP #2 | eGFP #3 | eGFP #4 |
|---|---|---|---|---|---|---|---|---|---|---|---|---|---|---|---|---|
| + | sp\|P14735\|IDE_HUMAN;tr\|Q5T5N3\|Q5T5N3_HUMAN | sp\|P14735\|ID | 323,31 | 1802900000 | 83 | 32 | 1,69E-08 | 7,99321985 | 19,48581 | 19,60563 | 19,44071 | 19,65087 | 28,00426 | 27,28982 | 27,70827 | 27,15354 |
| + | sp\|P68363\|TBA1B_HUMAN;sp\|P68366\|TBA4A_HUMAN;tr\|F8VVB9\|F8VVB9_HUMAN;tr\|C9JDS | sp\|P68363\|T | 323,31 | 3,2565E+10 | 200 | 1 | 3,85E-07 | 3,11166763 | 28,28776 | 28,52772 | 28,51608 | 28,53271 | 31,82491 | 31,54324 | 31,67405 | 31,26874 |
| + | sp\|P0DMV9\|HS71B_HUMAN;sp\|P0DMV8\|HS71A_HUMAN;tr\|A0A0G2JIW1\|A0A0G2JIW1_HUM | sp\|P0DMV9\| | 323,31 | 2,5768E+10 | 306 | 25 | 7,88E-07 | 1,62405109 | 29,38527 | 29,33745 | 29,55784 | 29,63843 | 31,19601 | 31,05692 | 31,06561 | 31,09666 |
| + | sp\|Q9BW92\|SYTM_HUMAN;tr\|U3KQG0\|U3KQG0_HUMAN;tr\|F6S7Q7\|F6S7Q7_HUMAN;tr\|U3 | sp\|Q9BW92\| | 24,499 | 74934000 | 19 | 9 | 1,28E-06 | 2,27569532 | 20,70261 | 20,54482 | 20,4247 | 20,40291 | 22,77125 | 22,61993 | 23,06733 | 22,71931 |
| + | tr\|Q5JP53\|Q5JP53_HUMAN;sp\|P07437\|TBB5_HUMAN;tr\|Q5ST81\|Q5ST81_HUMAN | tr\|Q5JP53\|Q | 323,31 | 2,5329E+10 | 152 | 6 | 1,28E-06 | 2,97265863 | 27,77052 | 28,1757 | 28,10993 | 28,23339 | 31,29844 | 31,07304 | 31,06728 | 30,74142 |
| + | sp\|P31689\|DNJA1_HUMAN | sp\|P31689\|D | 242,01 | 418600000 | 29 | 10 | 1,43E-06 | 3,94379663 | 21,55037 | 21,45874 | 21,83082 | 20,9239 | 25,51926 | 25,47636 | 25,41471 | 25,12869 |
| + | sp\|P49411\|EFTU_HUMAN;tr\|H3BNU3\|H3BNU3_HUMAN | sp\|P49411\|E | 323,31 | 2997800000 | 122 | 27 | 1,85E-06 | 2,24556684 | 25,73229 | 25,85262 | 25,82092 | 26,05935 | 28,26526 | 28,08851 | 28,26535 | 27,82833 |
| + | sp\|Q8NB90\|SPAT5_HUMAN;tr\|J3QRR3\|J3QRR3_HUMAN;tr\|J3QRW1\|J3QRW1_HUMAN | sp\|Q8NB90\|S | 25,333 | 48934000 | 18 | 3 | 2,33E-06 | 1,57896423 | 20,05126 | 20,12994 | 20,14805 | 20,23876 | 21,70438 | 21,80963 | 21,87279 | 21,49707 |
| + | tr\|A0A087WXZ3\|A0A087WXZ3_HUMAN;sp\|Q9Y5A9\|YTHD2_HUMAN;tr\|A0A024R7W5\|A0A02 | tr\|A0A087W | 47,221 | 121610000 | 24 | 7 | 3,58E-06 | 1,59357357 | 21,72146 | 21,7348 | 21,76052 | 21,8055 | 23,46936 | 23,23732 | 23,55192 | 23,13798 |
| + | sp\|O60884\|DNJA2_HUMAN;tr\|A0A087WT48\|A0A087WT48_HUMAN;tr\|I3L320\|I3L320_HUMA | sp\|O60884\|D | 150,4 | 269080000 | 32 | 11 | 4,01E-06 | 4,03915548 | 20,80381 | 21,07437 | 20,64157 | 20,19928 | 25,06139 | 24,84024 | 24,74842 | 24,2256 |
| + | sp\|P07196\|NFL_HUMAN;tr\|A0A087X0W2\|A0A087X0W2_HUMAN | sp\|P07196\|N | 323,31 | 4549900000 | 94 | 30 | 4,17E-06 | 1,98017359 | 26,7226 | 26,70859 | 26,81164 | 26,56725 | 28,94999 | 28,54709 | 28,79129 | 28,44241 |
| + | sp\|P27708\|PYR1_HUMAN;tr\|F8VPD4\|F8VPD4_HUMAN;tr\|H7C2E4\|H7C2E4_HUMAN;tr\|H7BZB | sp\|P27708\|P | 323,31 | 2539900000 | 170 | 61 | 4,42E-06 | 2,71483135 | 25,31614 | 25,33006 | 25,06435 | 24,873 | 27,60915 | 27,71914 | 28,22986 | 27,88472 |
| + | sp\|P33993\|MCM7_HUMAN;tr\|C9J8M6\|C9J8M6_HUMAN | sp\|P33993\|M | 271,35 | 437040000 | 64 | 20 | 4,70E-06 | 1,48879766 | 23,98129 | 23,64482 | 23,73738 | 23,64471 | 25,33217 | 25,0857 | 25,24449 | 25,30103 |
| + | sp\|Q14257\|RCN2_HUMAN;tr\|H0YL43\|H0YL43_HUMAN;tr\|A8MXP8\|A8MXP8_HUMAN | sp\|Q14257\|R | 125,94 | 71262000 | 22 | 8 | 6,42E-06 | 2,09923363 | 20,38732 | 20,69634 | 20,36196 | 20,74796 | 22,84169 | 22,49045 | 22,80499 | 22,4534 |
| + | sp\|P30837\|AL1B1_HUMAN;tr\|A0A0U1RQK9\|A0A0U1RQK9_HUMAN;tr\|H0Y2X5\|H0Y2X5_HUM | sp\|P30837\|A | 39,398 | 72075000 | 19 | 7 | 6,50E-06 | 1,66330528 | 20,81495 | 20,7913 | 20,91911 | 21,2237 | 22,45959 | 22,71597 | 22,65679 | 22,56994 |
| + | tr\|A0A087WTA5\|A0A087WTA5_HUMAN;sp\|Q9UI10\|EI2BD_HUMAN;tr\|E7ERK9\|E7ERK9_HUM | tr\|A0A087W | 13,68 | 59907000 | 14 | 8 | 6,76E-06 | 1,78170156 | 20,73983 | 20,46573 | 20,7978 | 20,28974 | 22,27989 | 22,36268 | 22,34852 | 22,42882 |
| + | tr\|E9PNM1\|E9PNM1_HUMAN;sp\|P37268\|FDFT_HUMAN;tr\|E9PS69\|E9PS69_HUMAN;tr\|E9PJG | tr\|E9PNM1\|E | 27,06 | 72029000 | 19 | 8 | 7,32E-06 | 3,00379054 | 19,70889 | NaN | 20,02715 | 19,68522 | 23,08601 | 22,57815 | 22,85719 | 22,72217 |
| + | sp\|Q9BSD7\|NTPCR_HUMAN;tr\|Q5TDF0\|Q5TDF0_HUMAN | sp\|Q9BSD7\|N | 65,852 | 68051000 | 26 | 9 | 7,58E-06 | 2,80820704 | 19,2954 | 19,87048 | 19,74991 | 19,9502 | 22,70094 | 22,80211 | 22,31614 | 22,27963 |
| + | sp\|O95831\|AIFM1_HUMAN;tr\|E9PMA0\|E9PMA0_HUMAN | sp\|O95831\|A | 38,175 | 121380000 | 24 | 8 | 7,90E-06 | 1,34474897 | 22,10008 | 21,88248 | 22,05142 | 22,07535 | 23,48542 | 23,14781 | 23,49997 | 23,35515 |
| + | sp\|P25705\|ATPA_HUMAN;tr\|K7ERX7\|K7ERX7_HUMAN;tr\|K7EK77\|K7EK77_HUMAN;tr\|K7EJP1 | sp\|P25705\|A | 323,31 | 1097300000 | 80 | 17 | 8,03E-06 | 1,33402872 | 25,23004 | 25,25469 | 25,04955 | 25,23213 | 26,63616 | 26,39121 | 26,69564 | 26,37951 |
| + | sp\|Q9H857\|NT5D2_HUMAN;tr\|H7C519\|H7C519_HUMAN | sp\|Q9H857\|N | 18,811 | 67755000 | 15 | 2 | 1,31E-05 | 2,08548069 | 20,37231 | 20,61342 | 20,60494 | 20,2674 | 22,18876 | 22,73074 | 22,78738 | 22,49311 |
| + | sp\|Q3ZCQ8\|TIM50_HUMAN;tr\|M0R0C3\|M0R0C3_HUMAN;tr\|M0R2F8\|M0R2F8_HUMAN;tr\|M | sp\|Q3ZCQ8\|T | 270,17 | 180060000 | 29 | 7 | 1,32E-05 | 2,61073542 | 21,10399 | 21,546 | 21,53621 | 21,6413 | 24,38383 | 24,18064 | 24,08958 | 23,61639 |
| + | sp\|O95816\|BAG2_HUMAN | sp\|O95816\|B | 55,859 | 208970000 | 16 | 5 | 1,42E-05 | 3,75131989 | 20,41648 | 20,74854 | 20,43166 | 20,679 | 24,65992 | 24,78756 | 24,29866 | 23,53481 |
| + | sp\|Q9Y230\|RUVB2_HUMAN;tr\|M0R0Y3\|M0R0Y3_HUMAN;tr\|X6R2L4\|X6R2L4_HUMAN;tr\|M0 | sp\|Q9Y230\|R | 323,31 | 659030000 | 65 | 18 | 1,50E-05 | 0,93503284 | 24,79996 | 24,67336 | 24,80538 | 24,83123 | 25,89349 | 25,59024 | 25,659 | 25,70733 |
| + | tr\|A0A087WTV6\|A0A087WTV6_HUMAN;sp\|Q96C36\|P5CR2_HUMAN;tr\|A0A087WZR9\|A0A08 | tr\|A0A087W | 49,366 | 166030000 | 24 | 9 | 1,84E-05 | 2,11711788 | 21,41085 | 21,78624 | 21,78417 | 22,19436 | 24,05904 | 23,96526 | 23,87615 | 23,74366 |
| + | sp\|P42677\|RS27_HUMAN;tr\|Q5T4L4\|Q5T4L4_HUMAN;tr\|C9J1C5\|C9J1C5_HUMAN | sp\|P42677\|R | 115,25 | 753980000 | 18 | 3 | 2,21E-05 | 2,21135712 | 23,95314 | 24,00813 | 23,62879 | 23,81636 | 26,55249 | 25,87077 | 25,99561 | 25,83297 |
| + | sp\|Q9BZE1\|RM37_HUMAN;tr\|S4R369\|S4R369_HUMAN;tr\|H0Y4J2\|H0Y4J2_HUMAN;tr\|A6NHR | sp\|Q9BZE1\|R | 3,466 | 20280000 | 9 | 3 | 2,51E-05 | 1,5125308 | 19,01822 | 18,8462 | 19,25776 | 19,17473 | 20,66891 | 20,74098 | 20,31385 | 20,62328 |
| + | tr\|A0A0C4DGL3\|A0A0C4DGL3_HUMAN;tr\|H0YNW5\|H0YNW5_HUMAN;tr\|H0YKC5\|H0YKC5_H | tr\|A0A0C4DG | 45,802 | 100110000 | 24 | 6 | 2,69E-05 | -1,37259293 | 23,21444 | 23,35259 | 23,33211 | 23,5836 | 22,04597 | 22,18414 | 21,74468 | 22,01757 |
| + | sp\|Q9Y265\|RUVB1_HUMAN;tr\|E7ETR0\|E7ETR0_HUMAN;tr\|H7C4G5\|H7C4G5_HUMAN;tr\|H7C | sp\|Q9Y265\|R | 62,004 | 439680000 | 55 | 15 | 2,85E-05 | 0,92208052 | 24,01606 | 24,23213 | 24,31417 | 24,22317 | 25,09563 | 24,98412 | 25,18332 | 25,21078 |
| + | sp\|Q9Y6E7\|SIR4_HUMAN | sp\|Q9Y6E7\|S | 323,31 | 1,2696E+10 | 112 | 18 | 2,90E-05 | 8,29285765 | 20,05073 | 23,19656 | 23,05988 | 21,9931 | 30,66306 | 30,11446 | 30,32184 | 30,37233 |
| + | tr\|E9PJM3\|E9PJM3_HUMAN;sp\|Q9UK99\|FBX3_HUMAN;tr\|G3V1E0\|G3V1E0_HUMAN;tr\|Q49A | tr\|E9PJM3\|E | 78,21 | 41665000 | 13 | 5 | 4,60E-05 | 2,4787364 | 19,85462 | 19,59173 | 19,50787 | 18,98565 | 21,52807 | 22,24403 | 22,0099 | 22,07281 |
| + | sp\|P52701\|MSH6_HUMAN;tr\|A0A087WWJ1\|A0A087WWJ1_HUMAN;tr\|A0A087WYT6\|A0A08 | sp\|P52701\|M | 50,005 | 128610000 | 35 | 15 | 4,61E-05 | 1,71747398 | 21,71494 | 22,21247 | 21,61733 | 21,60014 | 23,3648 | 23,39545 | 23,7145 | 23,54002 |
| + | sp\|P51570\|GALK1_HUMAN;tr\|K7ERJ9\|K7ERJ9_HUMAN;tr\|K7EII7\|K7EII7_HUMAN;tr\|K7ERN9\| | sp\|P51570\|G | 38,354 | 55083000 | 15 | 9 | 5,42E-05 | 2,56234312 | 20,35628 | 19,73235 | 19,4705 | 20,18749 | 22,8766 | 22,56895 | 22,16533 | 22,38511 |
| + | sp\|P08670\|VIME_HUMAN;tr\|B0YJC4\|B0YJC4_HUMAN;tr\|B0YJC5\|B0YJC5_HUMAN;tr\|Q5JVS8\| | sp\|P08670\|V | 323,31 | 5736700000 | 174 | 35 | 5,64E-05 | 1,19369173 | 27,56413 | 27,61947 | 27,7965 | 27,64897 | 28,80514 | 28,81922 | 29,14834 | 28,63114 |
| + | sp\|Q15149\|PLEC_HUMAN;tr\|E9PMV1\|E9PMV1_HUMAN;tr\|H0YDN1\|H0YDN1_HUMAN;tr\|E9P | sp\|Q15149\|P | 323,31 | 1014800000 | 193 | 89 | 5,73E-05 | 1,39154196 | 24,98242 | 25,20848 | 25,03649 | 24,74031 | 26,23035 | 26,36139 | 26,67101 | 26,27112 |
| + | sp\|P07355\|ANXA2_HUMAN;tr\|H0YN42\|H0YN42_HUMAN;tr\|H0YMD0\|H0YMD0_HUMAN;tr\|H | sp\|P07355\|A | 308,07 | 414470000 | 51 | 16 | 5,93E-05 | -1,50628281 | 25,18117 | 25,52543 | 25,61591 | 25,57903 | 23,94726 | 24,17988 | 23,65992 | 24,08934 |
| + | sp\|P11142\|HSP7C_HUMAN;tr\|E9PKE3\|E9PKE3_HUMAN;tr\|E9PN89\|E9PN89_HUMAN;tr\|E9PN | sp\|P11142\|H | 323,31 | 6069600000 | 158 | 27 | 6,11E-05 | 0,95021629 | 27,77649 | 27,89058 | 28,01919 | 27,94101 | 29,07217 | 28,86183 | 28,81357 | 28,68057 |
| + | sp\|P17066\|HSP76_HUMAN;sp\|P48741\|HSP77_HUMAN | sp\|P17066\|H | 323,31 | 1759500000 | 7 | 2 | 6,49E-05 | 1,65765524 | 25,44695 | 25,79897 | 25,66117 | 25,84372 | 26,95571 | 27,32543 | 27,62631 | 27,47398 |
| + | sp\|P00367\|DHE3_HUMAN;sp\|P49448\|DHE4_HUMAN | sp\|P00367\|D | 34,797 | 175770000 | 23 | 10 | 7,34E-05 | 0,96682787 | 22,87147 | 22,57646 | 22,92544 | 22,93744 | 23,90371 | 23,84986 | 23,65403 | 23,77052 |
| + | sp\|P30101\|PDIA3_HUMAN | sp\|P30101\|P | 147,39 | 792260000 | 74 | 16 | 7,73E-05 | -1,52623081 | 26,05906 | 26,5747 | 26,50361 | 26,60484 | 24,80907 | 25,05205 | 24,68357 | 25,09261 |
| + | sp\|P17812\|PYRG1_HUMAN;sp\|Q9NRF8\|PYRG2_HUMAN | sp\|P17812\|P | 206,48 | 300720000 | 47 | 14 | 8,09E-05 | 1,16734171 | 23,25624 | 23,37862 | 23,38943 | 23,56813 | 24,83306 | 24,50743 | 24,59747 | 24,32382 |
| + | tr\|E7ESP9\|E7ESP9_HUMAN;sp\|P07197\|NFM_HUMAN;tr\|E7EMV2\|E7EMV2_HUMAN;tr\|A0A08 | tr\|E7ESP9\|E7 | 323,31 | 1,1138E+10 | 254 | 49 | 8,33E-05 | 1,693645 | 28,37501 | 28,35502 | 28,29408 | 27,9042 | 29,8416 | 29,6376 | 30,31786 | 29,90584 |
| + | sp\|P68371\|TBB4B_HUMAN;tr\|A0A075B736\|A0A075B736_HUMAN;tr\|Q5SQY0\|Q5SQY0_HUM | sp\|P68371\|T | 79,011 | 1177700000 | 16 | 2 | 8,84E-05 | 2,59679317 | 23,84355 | 24,19182 | 24,06357 | 24,39663 | 26,8411 | 27,09245 | 26,97662 | 25,97257 |
| + | sp\|Q9NU22\|MDN1_HUMAN;tr\|Q5T795\|Q5T795_HUMAN;tr\|M0QXR3\|M0QXR3_HUMAN | sp\|Q9NU22\| | 271,28 | 322600000 | 73 | 55 | 9,33E-05 | 3,37779951 | 21,512 | 21,99314 | 21,51663 | 20,79297 | 24,42355 | 25,03355 | 25,50724 | 24,36159 |
| + | sp\|Q9NRH3\|TBG2_HUMAN;sp\|P23258\|TBG1_HUMAN;tr\|K7EKE5\|K7EKE5_HUMAN;tr\|K7EIS0\| | sp\|Q9NRH3\| | 41,672 | 49071000 | 8 | 4 | 0,00010381 | 0,78709602 | 21,25569 | NaN | 21,06336 | 21,0921 | 21,86306 | 21,93524 | 22,04006 | 21,85821 |
| + | sp\|Q8NC51\|PAIRB_HUMAN | sp\|Q8NC51\| | 96,365 | 380830000 | 39 | 12 | 0,00010394 | -0,78571033 | 24,8944 | 24,90679 | 25,17866 | 25,19682 | 24,24547 | 24,3332 | 24,24612 | 24,20903 |
| + | sp\|P61962\|DCAF7_HUMAN;tr\|A0A087WWI6\|A0A087WWI6_HUMAN | sp\|P61962\|D | 5,0395 | 20265000 | 7 | 4 | 0,00010861 | 0,94265795 | 19,41332 | 19,65081 | 19,70078 | 19,78328 | 20,51133 | 20,74476 | 20,63343 | 20,42931 |
| + | sp\|P23396\|RS3_HUMAN;tr\|E9PL09\|E9PL09_HUMAN;tr\|E9PPU1\|E9PPU1_HUMAN;tr\|F2Z2S8\| | sp\|P23396\|R | 323,31 | 1651000000 | 71 | 18 | 0,00013305 | 1,04163742 | 25,86665 | 26,05192 | 25,98986 | 26,00996 | 27,3132 | 27,04635 | 26,96157 | 26,76383 |
| + | tr\|A0A087X1B2\|A0A087X1B2_HUMAN;tr\|B9A018\|B9A018_HUMAN;sp\|Q53GS9\|SNUT2_HUM | tr\|A0A087X1 | 7,7371 | 6683600 | 9 | 2 | 0,00014675 | -1,37130499 | 19,2978 | 19,35907 | 19,45069 | 19,40815 | 17,69927 | 18,40711 | 17,81753 | 18,10656 |
| + | sp\|P56192\|SYMC_HUMAN;tr\|F5H2V6\|F5H2V6_HUMAN;tr\|F8VS26\|F8VS26_HUMAN;tr\|F8W0 | sp\|P56192\|S | 139,09 | 292260000 | 42 | 15 | 0,00015626 | 1,0393281 | 23,58818 | 23,52504 | 23,54144 | 23,36038 | 24,75343 | 24,22177 | 24,57606 | 24,62109 |
| + | sp\|P14625\|ENPL_HUMAN;tr\|Q96GW1\|Q96GW1_HUMAN;tr\|H0YIV0\|H0YIV0_HUMAN;tr\|F8W | sp\|P14625\|E | 205,41 | 772570000 | 70 | 20 | 0,00018703 | -0,79020786 | 26,07926 | 25,98895 | 26,10095 | 26,13401 | 25,25613 | 25,23184 | 25,10949 | 25,54489 |
| + | tr\|A0A087WXM6\|A0A087WXM6_HUMAN;tr\|J3QQT2\|J3QQT2_HUMAN;tr\|J3KRX5\|J3KRX5_HU | tr\|A0A087W | 19,026 | 213360000 | 26 | 4 | 0,00022319 | -0,82639456 | 24,29824 | 24,44481 | 24,32079 | 24,09852 | 23,5064 | 23,64812 | 23,42034 | 23,28193 |
| + | sp\|O75822\|EIF3J_HUMAN;tr\|H0YGJ7\|H0YGJ7_HUMAN;tr\|H0YLP3\|H0YLP3_HUMAN | sp\|O75822\|E | 4,8066 | 11310000 | 3 | 3 | 0,00022948 | -0,56224998 | 19,84189 | 19,94492 | 20,06316 | 19,95177 | 19,38062 | 19,33702 | 19,44692 | NaN |
| + | sp\|Q9Y285\|SYFA_HUMAN;tr\|K7ER00\|K7ER00_HUMAN;tr\|K7ER16\|K7ER16_HUMAN;tr\|K7EPH | sp\|Q9Y285\|S | 53,979 | 157870000 | 17 | 6 | 0,00023005 | 1,81800032 | 21,85011 | 21,88703 | 22,4789 | 21,85445 | 24,27441 | 23,81617 | 23,42291 | 23,829 |
| + | sp\|Q9BUF5\|TBB6_HUMAN;tr\|K7ESM5\|K7ESM5_HUMAN;tr\|K7ES63\|K7ES63_HUMAN;tr\|K7EL | sp\|Q9BUF5\|T | 35,551 | 222220000 | 11 | 3 | 0,00023459 | 2,78565788 | 21,65692 | 21,45061 | 21,10636 | 21,52392 | 23,4768 | 24,77274 | 24,81054 | 23,82036 |
| + | sp\|P27824\|CALX_HUMAN;tr\|H0Y9Q7\|H0Y9Q7_HUMAN;tr\|D6RGY2\|D6RGY2_HUMAN;tr\|D6RF | sp\|P27824\|C | 86,802 | 229350000 | 13 | 8 | 0,0002483 | -0,83882284 | 24,11585 | 24,37981 | 24,41493 | 24,09031 | 23,26999 | 23,34204 | 23,57681 | 23,45676 |
| + | tr\|E5KLK1\|E5KLK1_HUMAN;tr\|E5KLJ9\|E5KLJ9_HUMAN;tr\|E5KLJ5\|E5KLJ5_HUMAN;tr\|E5KLJ6\|E | tr\|E5KLK1\|E5 | 37,168 | 116830000 | 21 | 1 | 0,00024838 | 1,8134675 | 21,50368 | 21,53825 | 21,15919 | 21,704 | 23,49094 | 23,42355 | 23,56454 | 22,67997 |
| + | sp\|P78417\|GSTO1_HUMAN;tr\|Q5TA02\|Q5TA02_HUMAN;tr\|Q5TA01\|Q5TA01_HUMAN | sp\|P78417\|G | 12,189 | 83141000 | 17 | 4 | 0,00027883 | -1,30067158 | 22,87085 | 23,16566 | 23,02442 | 23,20784 | 21,79749 | 22,15224 | 21,40145 | 21,7149 |
| + | sp\|P36542\|ATPG_HUMAN | sp\|P36542\|A | 28,01 | 127180000 | 23 | 6 | 0,00028262 | 1,09064007 | 22,25324 | 22,24566 | 22,19346 | 22,23637 | 23,5657 | 23,42944 | 23,39061 | 22,90555 |
| + | sp\|P14314\|GLU2B_HUMAN;tr\|K7ELL7\|K7ELL7_HUMAN;tr\|K7EKX1\|K7EKX1_HUMAN;tr\|K7EJ7 | sp\|P14314\|G | 14,311 | 101410000 | 20 | 7 | 0,0003205 | -0,94900465 | 23,05886 | 23,20647 | 23,4443 | 23,16079 | 22,42637 | 22,38835 | 22,27378 | 21,9859 |
| + | sp\|P05023\|AT1A1_HUMAN;tr\|M0R116\|M0R116_HUMAN;sp\|P13637\|AT1A3_HUMAN;tr\|A0A | sp\|P05023\|A | 13,813 | 39108000 | 12 | 7 | 0,00033114 | -0,97759485 | 21,91987 | 21,78592 | 21,97337 | 21,81464 | 21,03322 | 20,54312 | 20,89378 | 21,11331 |
| + | sp\|Q15477\|SKIV2_HUMAN;tr\|H7C5N0\|H7C5N0_HUMAN;tr\|H7C4L3\|H7C4L3_HUMAN;tr\|F8W | sp\|Q15477\|S | 26,43 | 23625000 | 17 | 8 | 0,00033242 | 2,37813838 | 18,75887 | 18,75392 | NaN | 18,74613 | 21,11832 | 21,50465 | 21,41761 | 20,48385 |
| + | sp\|P07900\|HS90A_HUMAN;tr\|G3V2J8\|G3V2J8_HUMAN;sp\|Q14568\|HS902_HUMAN;sp\|Q58F | sp\|P07900\|H | 323,31 | 1630900000 | 89 | 16 | 0,00033571 | -0,55695677 | 27,04781 | 27,06476 | 27,08487 | 27,03118 | 26,47653 | 26,53224 | 26,3136 | 26,67842 |
| + | sp\|Q9NYU2\|UGGG1_HUMAN;tr\|H7BZG0\|H7BZG0_HUMAN | sp\|Q9NYU2\| | 17,749 | 37467000 | 13 | 7 | 0,00034602 | 1,22390366 | 20,48611 | 20,63476 | 20,14146 | 20,3229 | 21,30279 | 21,51114 | 21,80388 | 21,86302 |
| + | sp\|P26583\|HMGB2_HUMAN;tr\|D6R9A6\|D6R9A6_HUMAN | sp\|P26583\|H | 10,895 | 112640000 | 6 | 2 | 0,00036772 | -1,00065931 | 23,44405 | 23,49644 | NaN | 23,47729 | NaN | 22,33615 | 22,44177 | 22,63788 |
| + | sp\|P08758\|ANXA5_HUMAN;tr\|D6RBE9\|D6RBE9_HUMAN;tr\|D6RBL5\|D6RBL5_HUMAN;tr\|E9P | sp\|P08758\|A | 44,126 | 204970000 | 22 | 8 | 0,00038828 | -0,75522518 | 24,03565 | 24,2786 | 24,24372 | 24,20292 | 23,35582 | 23,54851 | 23,21681 | 23,61885 |
| + | sp\|Q9Y2S7\|PDIP2_HUMAN;tr\|B4DEM9\|B4DEM9_HUMAN | sp\|Q9Y2S7\|P | 10,183 | 62187000 | 11 | 6 | 0,00041839 | 2,16742897 | 19,91003 | 20,32553 | 20,22835 | 20,77394 | 22,87701 | 22,76525 | 21,7565 | 22,50882 |
| + | tr\|J3QS39\|J3QS39_HUMAN;tr\|J3QTR3\|J3QTR3_HUMAN;tr\|F5H6Q2\|F5H6Q2_HUMAN;sp\|P62 | tr\|J3QS39\|J3 | 323,31 | 1500000000 | 39 | 6 | 0,00042066 | 0,66782093 | 26,25308 | 26,18592 | 26,13022 | 26,18215 | 26,76231 | 26,98577 | 26,64361 | 27,03097 |

| | | | | | | | | | | | | | | | | |
|---|---|---|---|---|---|---|---|---|---|---|---|---|---|---|---|---|
| + | sp\|P32969\|RL9_HUMAN;tr\|D6RAN4\|D6RAN4_HUMAN;tr\|H0Y9V9\|H0Y9V9_HUMAN;tr\|H0Y9R | sp\|P32969\|R | 122,8 | 76848000 | 16 | 6 | 0,00044692 | -1,1557889 | 22,98456 | 23,047 | 22,8975 | 22,37241 | 21,66166 | 21,49799 | 21,71892 | 21,79974 |
| + | sp\|P35579\|MYH9_HUMAN;tr\|A0A0C4DFM8\|A0A0C4DFM8_HUMAN;tr\|E7ERA5\|E7ERA5_HUM | sp\|P35579\|M | 86,019 | 191800000 | 52 | 19 | 0,00045389 | -0,79258966 | 24,07863 | 24,07243 | 24,05457 | 23,97727 | 23,00136 | 23,19104 | 23,54191 | 23,27824 |
| + | sp\|Q13200\|PSMD2_HUMAN;tr\|H7C1H2\|H7C1H2_HUMAN;tr\|H7C2Q3\|H7C2Q3_HUMAN;tr\|C9 | sp\|Q13200\|P | 272,49 | 182810000 | 45 | 14 | 0,00049349 | 0,551929 | 23,21846 | 23,05013 | 23,39401 | 23,10396 | 23,67799 | 23,74725 | 23,73376 | 23,81529 |
| + | tr\|A0A087X1S2\|A0A087X1S2_HUMAN;sp\|P67809\|YBOX1_HUMAN;tr\|H0Y449\|H0Y449_HUMA | tr\|A0A087X1S | 270,93 | 210540000 | 39 | 5 | 0,00054162 | -0,59001017 | 24,03607 | 23,93148 | 24,2627 | 23,86872 | 23,38627 | 23,46837 | 23,44139 | 23,44291 |
| + | tr\|B8ZZK4\|B8ZZK4_HUMAN;tr\|H7C2W9\|H7C2W9_HUMAN;tr\|C9JU56\|C9JU56_HUMAN;tr\|B7Z | tr\|B8ZZK4\|B8 | 10,182 | 378870000 | 13 | 4 | 0,00055599 | -1,0531888 | 25,01572 | 25,478 | 25,18044 | 24,89592 | 24,31264 | 24,04335 | 24,14538 | 23,85595 |
| + | sp\|Q92973\|TNPO1_HUMAN;tr\|E7EW37\|E7EW37_HUMAN;tr\|S4R398\|S4R398_HUMAN | sp\|Q92973\|T | 24,541 | 68774000 | 17 | 6 | 0,00055834 | -0,40821171 | 22,35982 | 22,39788 | 22,32833 | 22,41784 | 21,80116 | 22,06161 | 22,02901 | 21,97925 |
| + | tr\|E9PKZ0\|E9PKZ0_HUMAN;sp\|P62917\|RL8_HUMAN;tr\|G3V1A1\|G3V1A1_HUMAN;tr\|E9PKU4 | tr\|E9PKZ0\|E9 | 52,151 | 550350000 | 23 | 5 | 0,00057795 | -0,98334837 | 25,92334 | 25,77957 | 25,67069 | 25,41976 | 24,49375 | 24,65638 | 24,99386 | 24,71597 |
| + | sp\|Q7Z6Z7\|HUWE1_HUMAN;tr\|H0Y659\|H0Y659_HUMAN;tr\|A0A087X1S3\|A0A087X1S3_HUM | sp\|Q7Z6Z7\|H | 11,083 | 20462000 | 8 | 7 | 0,00059228 | 0,78153992 | 20,02296 | 19,91704 | 19,65529 | 19,74787 | 20,37039 | 20,64271 | 20,70269 | 20,75352 |
| + | sp\|P46060\|RAGP1_HUMAN;tr\|B0QYT5\|B0QYT5_HUMAN;tr\|B0QYT4\|B0QYT4_HUMAN;tr\|H0Y | sp\|P46060\|R | 58,542 | 127960000 | 34 | 12 | 0,00059938 | -0,57070732 | 23,23919 | 23,46114 | 23,27469 | 23,31576 | 22,96201 | 22,64309 | 22,67734 | 22,72551 |
| + | sp\|Q96EY1\|DNJA3_HUMAN;tr\|I3L1T6\|I3L1T6_HUMAN | sp\|Q96EY1\|D | 42,708 | 210120000 | 21 | 9 | 0,00060506 | 2,92689514 | 20,92716 | 20,68252 | 21,73198 | 21,34392 | 24,84566 | 24,40886 | 24,0878 | 23,05084 |
| + | sp\|Q9UBN7\|HDAC6_HUMAN;tr\|A6NDI8\|A6NDI8_HUMAN;tr\|E7EPS2\|E7EPS2_HUMAN;tr\|C9JE | sp\|Q9UBN7\|I | 151,94 | 253370000 | 41 | 9 | 0,00060567 | 1,7416563 | 22,47593 | 23,09905 | 22,80473 | 22,54409 | 23,93787 | 24,27597 | 24,75062 | 24,92597 |
| + | sp\|P04181\|OAT_HUMAN | sp\|P04181\|O | 110,17 | 734870000 | 54 | 14 | 0,00061078 | 2,11978769 | 22,99039 | 23,9478 | 24,20508 | 24,45098 | 26,13673 | 26,05397 | 26,00123 | 25,88148 |
| + | sp\|Q16531\|DDB1_HUMAN;tr\|F5GY55\|F5GY55_HUMAN;tr\|F5H2L3\|F5H2L3_HUMAN;tr\|F5GW | sp\|Q16531\|D | 37,577 | 226570000 | 39 | 14 | 0,00061404 | 0,64963007 | 23,56999 | 23,40133 | 23,55415 | 23,52767 | 24,13469 | 23,91534 | 24,2798 | 24,32182 |
| + | sp\|Q9NSE4\|SYIM_HUMAN | sp\|Q9NSE4\|S | 96,834 | 188470000 | 30 | 14 | 0,00067096 | 2,06609297 | 21,73674 | 22,10582 | 21,4108 | 22,7066 | 23,79064 | 23,84019 | 24,49802 | 24,09547 |
| + | sp\|Q9Y490\|TLN1_HUMAN;sp\|Q9Y4G6\|TLN2_HUMAN | sp\|Q9Y490\|T | 9,635 | 46212000 | 10 | 8 | 0,00072246 | -0,43302536 | 21,74082 | 21,75772 | 21,70485 | 21,58568 | 21,18392 | 21,22653 | 21,21573 | 21,43079 |
| + | sp\|P35232\|PHB_HUMAN;tr\|C9JW96\|C9JW96_HUMAN;tr\|C9JZ20\|C9JZ20_HUMAN;tr\|E7ESE2\| | sp\|P35232\|P | 33,803 | 113170000 | 25 | 10 | 0,00072661 | 1,52850962 | 21,5888 | 21,37279 | 21,32729 | 22,16023 | 23,07405 | 23,56546 | 22,8866 | 23,03703 |
| + | sp\|P17987\|TCPA_HUMAN;tr\|E7ERF2\|E7ERF2_HUMAN;tr\|E7EQR6\|E7EQR6_HUMAN;tr\|F5H28 | sp\|P17987\|T | 176,47 | 964890000 | 56 | 20 | 0,00074599 | 0,39363527 | 25,60147 | 25,5846 | 25,80519 | 25,74707 | 26,15509 | 26,05316 | 26,00973 | 26,09489 |
| + | tr\|I3L2C7\|I3L2C7_HUMAN;sp\|P57678\|GEMI4_HUMAN | tr\|I3L2C7\|I3L | 19,787 | 37735000 | 13 | 7 | 0,00075825 | 1,88722563 | 19,99162 | 19,67635 | 19,65017 | 19,58893 | 20,79352 | 21,67616 | 22,0971 | 21,88919 |
| + | sp\|P13995\|MTDC_HUMAN;tr\|B9A062\|B9A062_HUMAN;tr\|B8ZZU9\|B8ZZU9_HUMAN | sp\|P13995\|M | 11,923 | 93732000 | 10 | 4 | 0,00076056 | 0,59327698 | 22,26284 | 22,45194 | 22,0733 | 22,20425 | 22,87049 | 22,97722 | 22,75371 | 22,76401 |
| + | sp\|P12277\|KCRB_HUMAN;tr\|H0YJG0\|H0YJG0_HUMAN;tr\|G3V4N7\|G3V4N7_HUMAN;tr\|G3V4 | sp\|P12277\|K | 175,99 | 704460000 | 44 | 12 | 0,00078612 | -0,98145676 | 25,77228 | 25,89913 | 26,25641 | 26,18741 | 25,04573 | 25,06899 | 24,77575 | 25,29894 |
| + | sp\|Q8NF37\|PCAT1_HUMAN;tr\|A0A0G2JQ62\|A0A0G2JQ62_HUMAN;tr\|A0A0G2JRI7\|A0A0G2JR | sp\|Q8NF37\|P | 13,498 | 15984000 | 4 | 3 | 0,00080639 | 0,54481459 | 19,70818 | 19,6263 | 19,71139 | 19,72384 | 20,08317 | 20,45854 | 20,1278 | 20,27946 |
| + | tr\|Q5JR95\|Q5JR95_HUMAN;sp\|P62241\|RS8_HUMAN | tr\|Q5JR95\|Q | 323,31 | 308230000 | 27 | 7 | 0,00082726 | -0,61090374 | 24,87895 | 24,76875 | 24,61661 | 24,46781 | 24,05084 | 24,05192 | 23,99499 | 24,19077 |
| + | sp\|P24752\|THIL_HUMAN;tr\|H0YEL7\|H0YEL7_HUMAN;tr\|E9PRQ6\|E9PRQ6_HUMAN | sp\|P24752\|T | 26,289 | 133110000 | 17 | 8 | 0,00083332 | -1,04659986 | 23,54097 | 23,61392 | 23,60513 | 23,80548 | 22,60716 | 22,30129 | 22,43489 | 23,03576 |
| + | sp\|P62829\|RL23_HUMAN;tr\|J3KT29\|J3KT29_HUMAN;tr\|C9JD32\|C9JD32_HUMAN;tr\|B9ZVP7\| | sp\|P62829\|R | 120,48 | 1252200000 | 26 | 9 | 0,00086671 | 1,36982346 | 25,02323 | 25,59619 | 25,3839 | 24,94566 | 26,77731 | 26,86597 | 26,6487 | 26,1363 |
| + | sp\|P53618\|COPB_HUMAN;tr\|E9PP73\|E9PP73_HUMAN | sp\|P53618\|C | 15,061 | 93797000 | 15 | 7 | 0,00086886 | 0,82362366 | 22,28351 | 21,86982 | 22,2321 | 22,10965 | 22,77888 | 22,82083 | 23,21202 | 22,97783 |
| + | sp\|Q13838\|DX39B_HUMAN;tr\|A0A0G2JJZ9\|A0A0G2JJZ9_HUMAN;tr\|Q5STU3\|Q5STU3_HUMA | sp\|Q13838\|D | 88,343 | 523240000 | 46 | 6 | 0,00087173 | -0,73430634 | 25,31057 | 25,46005 | 25,48557 | 25,63675 | 24,66503 | 24,88311 | 24,49082 | 24,91675 |
| + | sp\|P18206\|VINC_HUMAN;tr\|Q5JQ13\|Q5JQ13_HUMAN;tr\|A0A096LPE1\|A0A096LPE1_HUMAN | sp\|P18206\|V | 20,001 | 55526000 | 27 | 9 | 0,00090733 | -0,94232559 | 22,40852 | 22,58456 | 22,21824 | 22,35861 | 21,41338 | 21,38337 | 21,1784 | 21,82547 |
| + | sp\|P47914\|RL29_HUMAN | sp\|P47914\|R | 13,433 | 179600000 | 13 | 2 | 0,00096315 | -0,73415279 | 24,0694 | 24,18155 | 23,95748 | 23,72203 | 23,44316 | 23,08918 | 23,23326 | 23,22825 |
| + | sp\|P05198\|IF2A_HUMAN;tr\|H0YJS4\|H0YJS4_HUMAN;tr\|G3V4T5\|G3V4T5_HUMAN | sp\|P05198\|IF | 18,648 | 59429000 | 20 | 7 | 0,001048 | -0,79986954 | 22,08748 | 22,46501 | 22,47551 | 22,2554 | 21,78093 | 21,38685 | 21,35035 | 21,56579 |
| + | sp\|Q9UQ80\|PA2G4_HUMAN;tr\|F8VR77\|F8VR77_HUMAN;tr\|H0YIN7\|H0YIN7_HUMAN;tr\|F8W | sp\|Q9UQ80\|I | 138,58 | 508840000 | 48 | 14 | 0,00106077 | -0,77320242 | 25,36326 | 25,37352 | 25,69067 | 25,58035 | 24,85234 | 24,84364 | 24,41944 | 24,79956 |
| + | sp\|P14550\|AK1A1_HUMAN;tr\|V9GYG2\|V9GYG2_HUMAN;tr\|V9GYP9\|V9GYP9_HUMAN;tr\|Q5T | sp\|P14550\|A | 4,4608 | 46852000 | 10 | 4 | 0,0010795 | -1,20003462 | 22,01808 | 22,17316 | 22,15576 | 22,37289 | 21,08622 | 21,1249 | 20,42234 | 21,2863 |
| + | tr\|K7EPP7\|K7EPP7_HUMAN;tr\|K7ESP1\|K7ESP1_HUMAN;sp\|Q99615\|DNJC7_HUMAN;tr\|K7EIH | tr\|K7EPP7\|K7 | 16,907 | 33388000 | 7 | 5 | 0,00108629 | 0,61263561 | 20,85536 | 20,90353 | 20,83098 | NaN | 21,65932 | 21,5354 | 21,32893 | 21,38005 |
| + | sp\|P09429\|HMGB1_HUMAN;tr\|Q5T7C4\|Q5T7C4_HUMAN;sp\|B2RPK0\|HGB1A_HUMAN;sp\|P23 | sp\|P09429\|H | 57,811 | 410720000 | 25 | 6 | 0,00112742 | -1,25265646 | 25,09317 | 25,5402 | 25,45403 | 25,43654 | 24,61863 | 23,96737 | 23,72067 | 24,20665 |
| + | sp\|P00918\|CAH2_HUMAN;tr\|E5RID5\|E5RID5_HUMAN;tr\|E5RK37\|E5RK37_HUMAN | sp\|P00918\|C | 141,03 | 670490000 | 46 | 9 | 0,00113062 | -1,21690607 | 25,86901 | 26,02035 | 26,04839 | 26,3124 | 24,88428 | 24,97136 | 24,32079 | 25,20609 |
| + | sp\|P45954\|ACDSB_HUMAN | sp\|P45954\|A | 9,5977 | 45390000 | 9 | 6 | 0,00113667 | 1,29606104 | 20,62471 | 20,51912 | 20,24016 | 20,56672 | 21,79634 | 22,32127 | 21,69379 | 21,32356 |
| + | sp\|Q9P258\|RCC2_HUMAN | sp\|Q9P258\|R | 29,334 | 84786000 | 22 | 6 | 0,00115435 | -0,63011026 | 22,47108 | 22,85369 | 22,87341 | 22,87093 | 22,18123 | 22,2127 | 22,00751 | 22,14722 |
| + | sp\|P62826\|RAN_HUMAN;tr\|B5MDF5\|B5MDF5_HUMAN;tr\|J3KQE5\|J3KQE5_HUMAN;tr\|F5H01 | sp\|P62826\|R | 151,53 | 701400000 | 25 | 8 | 0,00119533 | -0,81896305 | 25,78756 | 25,91216 | 26,06003 | 26,0625 | 25,09547 | 25,49485 | 25,04856 | 24,90752 |
| + | tr\|A0A0A6YYJ8\|A0A0A6YYJ8_HUMAN;tr\|A0A0A6YYC3\|A0A0A6YYC3_HUMAN;sp\|Q9Y383\|LC7L | tr\|A0A0A6YY. | 12,778 | 54416000 | 15 | 3 | 0,00119659 | -0,83297491 | 21,9132 | 22,17937 | 22,38287 | 22,38696 | 21,28918 | 21,30196 | 21,28167 | 21,6577 |
| + | sp\|Q9UBS4\|DJB11_HUMAN;tr\|H7C2Y5\|H7C2Y5_HUMAN | sp\|Q9UBS4\|D | 77,51 | 75304000 | 18 | 8 | 0,00121307 | 1,91227913 | 20,51316 | 20,63556 | 20,67736 | 20,74608 | 22,59562 | 22,98722 | 23,02722 | 21,61122 |
| + | sp\|P10398\|ARAF_HUMAN;tr\|Q96II5\|Q96II5_HUMAN;tr\|H7C4S5\|H7C4S5_HUMAN | sp\|P10398\|A | 52,797 | 284800000 | 31 | 9 | 0,00121403 | 3,03146458 | 20,84736 | 22,64229 | 21,6897 | 21,03617 | 23,80892 | 25,46136 | 24,54475 | 24,52636 |
| + | tr\|A0A024QZP7\|A0A024QZP7_HUMAN;sp\|P06493\|CDK1_HUMAN;tr\|E5RIU6\|E5RIU6_HUMAN | tr\|A0A024QZ | 53,959 | 348720000 | 31 | 10 | 0,0012461 | 1,1167531 | 23,39048 | 24,06587 | 23,68067 | 23,7168 | 25,2288 | 24,76263 | 24,59121 | 24,7382 |
| + | sp\|P31040\|SDHA_HUMAN;tr\|A0A087X1I3\|A0A087X1I3_HUMAN;tr\|D6RFM5\|D6RFM5_HUMA | sp\|P31040\|S | 17,419 | 81060000 | 10 | 6 | 0,00130795 | 1,76087284 | 20,64808 | 21,34131 | 21,13458 | 20,82602 | 22,58217 | 23,1888 | 22,05696 | 23,16555 |
| + | sp\|P84077\|ARF1_HUMAN;sp\|P61204\|ARF3_HUMAN;tr\|F5H423\|F5H423_HUMAN;tr\|F5H0C7\| | sp\|P84077\|A | 105,77 | 253460000 | 27 | 5 | 0,00131495 | -1,00833082 | 24,69285 | 24,38667 | 24,56349 | 24,27199 | 23,64262 | 23,7679 | 23,07897 | 23,39218 |
| + | sp\|O15269\|SPTC1_HUMAN | sp\|O15269\|S | 78,421 | 86465000 | 16 | 10 | 0,00132178 | 1,77191353 | 21,20174 | 20,76806 | 21,12666 | 21,15146 | 23,6075 | 22,80043 | 22,77173 | 22,15592 |
| + | sp\|P09960\|LKHA4_HUMAN;tr\|B4DEH5\|B4DEH5_HUMAN | sp\|P09960\|L | 78,287 | 56529000 | 22 | 8 | 0,0013224 | -0,71632719 | 22,04363 | 22,25468 | 22,29286 | 22,26937 | 21,28991 | 21,80613 | 21,521 | 21,37819 |
| + | sp\|Q71U36\|TBA1A_HUMAN;sp\|Q13748\|TBA3C_HUMAN;sp\|Q6PEY2\|TBA3E_HUMAN;tr\|F8VQ | sp\|Q71U36\|T | 17,633 | 187250000 | 12 | 1 | 0,0013516 | 3,5673852 | 20,45192 | 20,00826 | 20,46802 | 20,17719 | 22,79753 | 24,34969 | 22,85217 | 25,37554 |
| + | sp\|P23921\|RIR1_HUMAN;tr\|E9PL69\|E9PL69_HUMAN;tr\|E9PP77\|E9PP77_HUMAN;tr\|H0YCY7\| | sp\|P23921\|R | 23,924 | 80699000 | 27 | 11 | 0,0013625 | -0,79877424 | 22,48866 | 23,05137 | 22,84621 | 22,77181 | 21,75727 | 22,09487 | 21,99814 | 22,11267 |
| + | sp\|Q14739\|LBR_HUMAN;tr\|C9JXK0\|C9JXK0_HUMAN;tr\|C9JES9\|C9JES9_HUMAN | sp\|Q14739\|L | 131,08 | 67814000 | 18 | 4 | 0,00140903 | 0,79206371 | 21,29922 | 21,43731 | 21,76547 | 21,47571 | 22,26405 | 22,10572 | 22,57907 | 22,19713 |
| + | sp\|Q9Y4R8\|TELO2_HUMAN;tr\|H3BRS3\|H3BRS3_HUMAN;tr\|H3BU45\|H3BU45_HUMAN | sp\|Q9Y4R8\|T | 71,605 | 66864000 | 21 | 13 | 0,00141988 | 1,94877386 | 20,70329 | 21,08005 | 19,82208 | 20,5418 | 22,48971 | 22,61865 | 22,96842 | 21,86552 |
| + | sp\|Q9NZL4\|HPBP1_HUMAN;tr\|K7EN20\|K7EN20_HUMAN;tr\|K7EL16\|K7EL16_HUMAN;tr\|K7EN | sp\|Q9NZL4\|H | 13,696 | 32255000 | 11 | 5 | 0,00144032 | 0,58055067 | 20,83391 | 20,77523 | 20,56141 | 20,66622 | 21,47833 | 21,19856 | 21,10162 | 21,38047 |
| + | sp\|Q96AE4\|FUBP1_HUMAN;tr\|E9PEB5\|E9PEB5_HUMAN;tr\|C9JSZ1\|C9JSZ1_HUMAN;sp\|Q96I2 | sp\|Q96AE4\|F | 30,436 | 176010000 | 34 | 9 | 0,00148576 | -0,58518553 | 23,69424 | 23,79005 | 23,77766 | 24,01546 | 23,16378 | 23,16016 | 23,13717 | 23,47556 |
| + | sp\|Q969Z0\|TBRG4_HUMAN;tr\|H7C4R5\|H7C4R5_HUMAN;tr\|C9IZN7\|C9IZN7_HUMAN | sp\|Q969Z0\|T | 6,976 | 15149000 | 7 | 3 | 0,00148813 | 0,2886157 | 19,69569 | 19,61061 | 19,7147 | 19,78931 | 19,95035 | 20,08537 | 19,91615 | 20,01291 |
| + | sp\|Q96HE7\|ERO1A_HUMAN;tr\|G3V2H0\|G3V2H0_HUMAN;tr\|G3V5B3\|G3V5B3_HUMAN;tr\|Q5 | sp\|Q96HE7\|E | 29,697 | 39694000 | 18 | 5 | 0,00149457 | -0,79173279 | 21,69387 | 22,03224 | 21,90183 | 21,85152 | 21,10738 | 21,22594 | 20,71549 | 21,26373 |
| + | sp\|Q5JTZ9\|SYAM_HUMAN | sp\|Q5JTZ9\|S | 37,946 | 50867000 | 24 | 8 | 0,00152133 | 1,00837517 | 20,94185 | 21,36906 | 20,78564 | 21,10495 | 22,24025 | 21,66583 | 22,10137 | 22,22756 |
| + | sp\|Q8N1F7\|NUP93_HUMAN;tr\|H3BVG0\|H3BVG0_HUMAN;tr\|H3BNN5\|H3BNN5_HUMAN;tr\|H | sp\|Q8N1F7\|N | 14,611 | 36334000 | 16 | 8 | 0,00154619 | -0,6501565 | 21,55075 | 21,61796 | 21,80148 | 21,64518 | 20,90551 | 21,08037 | 20,77 | 21,25885 |
| + | sp\|P50990\|TCPQ_HUMAN;tr\|H7C4C8\|H7C4C8_HUMAN;tr\|H7C2U0\|H7C2U0_HUMAN | sp\|P50990\|T | 294,5 | 868830000 | 85 | 23 | 0,00175494 | -0,49495173 | 25,92334 | 26,00759 | 26,17491 | 26,23392 | 25,50585 | 25,64878 | 25,48081 | 25,72453 |
| + | tr\|A0A087X054\|A0A087X054_HUMAN;sp\|Q9Y4L1\|HYOU1_HUMAN;tr\|E9PJ21\|E9PJ21_HUMA | tr\|A0A087X0 | 11,5 | 34830000 | 11 | 6 | 0,00176657 | -1,40338929 | 21,52057 | 22,01318 | 22,20786 | 22,07114 | NaN | 20,33604 | 20,40997 | 20,90338 |
| + | tr\|B7Z7P8\|B7Z7P8_HUMAN;sp\|P62495\|ERF1_HUMAN;tr\|I3L492\|I3L492_HUMAN;tr\|D6RCB3\| | tr\|B7Z7P8\|B7 | 16,12 | 57649000 | 18 | 7 | 0,00177915 | -0,60273838 | 22,08047 | 22,112 | 22,28481 | 22,32237 | 21,41183 | 21,86469 | 21,55544 | 21,55675 |
| + | sp\|Q15084\|PDIA6_HUMAN | sp\|Q15084\|P | 219,42 | 296170000 | 34 | 9 | 0,0017825 | -0,84444523 | 24,62556 | 24,80336 | 24,80312 | 24,97998 | 23,78596 | 24,12629 | 23,66036 | 24,26163 |
| + | sp\|Q13724\|MOGS_HUMAN;tr\|C9J8D4\|C9J8D4_HUMAN;tr\|C9JDQ1\|C9JDQ1_HUMAN | sp\|Q13724\|M | 20,964 | 51140000 | 20 | 9 | 0,00178258 | 1,1351018 | 20,88442 | 20,88181 | 20,99514 | 20,78421 | 22,38393 | 22,11032 | 22,171 | 21,42075 |
| + | tr\|Q5HYB6\|Q5HYB6_HUMAN;tr\|A0A087WWU8\|A0A087WWU8_HUMAN;tr\|D6R904\|D6R904_ | tr\|Q5HYB6\|Q | 102,23 | 328610000 | 44 | 7 | 0,00179774 | -0,79473543 | 24,5335 | 24,8412 | 24,72817 | 24,76759 | 24,05631 | 24,23586 | 23,68014 | 23,71921 |
| + | sp\|P04075\|ALDOA_HUMAN;tr\|J3KPS3\|J3KPS3_HUMAN;tr\|H3BQN4\|H3BQN4_HUMAN;tr\|H3B | sp\|P04075\|A | 323,31 | 2723800000 | 92 | 16 | 0,00182535 | -0,788692 | 27,67963 | 27,73635 | 27,87203 | 27,99863 | 27,05051 | 27,25124 | 26,66166 | 27,16844 |
| + | tr\|E9PBK6\|E9PBK6_HUMAN;tr\|E9PDU5\|E9PDU5_HUMAN;sp\|Q9NNW5\|WDR6_HUMAN;tr\|A0 | tr\|E9PBK6\|E9 | 2,1418 | 9605000 | 4 | 2 | 0,00184011 | 0,84693702 | 18,96747 | NaN | 18,70215 | 18,6989 | 19,77043 | 19,84387 | 19,42232 | 19,50916 |
| + | sp\|O43670\|ZN207_HUMAN;tr\|J3QRS9\|J3QRS9_HUMAN;tr\|X6R4W8\|X6R4W8_HUMAN;tr\|J3K | sp\|O43670\|Z | 10,189 | 37035000 | 11 | 3 | 0,00186501 | -0,68453598 | 21,56211 | 21,27725 | 21,86533 | 21,59578 | 20,98294 | 20,9556 | 20,85064 | 20,77314 |
| + | sp\|P42704\|LPPRC_HUMAN;tr\|B8ZZ38\|B8ZZ38_HUMAN;tr\|A0A0C4DG06\|A0A0C4DG06_HUMA | sp\|P42704\|L | 200,57 | 435190000 | 71 | 28 | 0,00190324 | -0,57439518 | 25,03381 | 25,18643 | 25,19888 | 25,25429 | 24,62199 | 24,47494 | 24,41886 | 24,86003 |
| + | sp\|P27797\|CALR_HUMAN;tr\|K7EJB9\|K7EJB9_HUMAN | sp\|P27797\|C | 61,769 | 178700000 | 26 | 7 | 0,00194021 | -0,60956621 | 23,85833 | 23,93814 | 24,06242 | 24,10597 | 23,24367 | 23,61122 | 23,18126 | 23,49045 |
| + | tr\|E7EPM6\|E7EPM6_HUMAN;sp\|P33121\|ACSL1_HUMAN;tr\|B7Z3Z9\|B7Z3Z9_HUMAN;tr\|H0Y9 | tr\|E7EPM6\|E | 17,224 | 122920000 | 17 | 11 | 0,00197248 | 2,85295359 | 20,53488 | 20,42039 | 20,66926 | NaN | 23,03063 | 23,64284 | 24,39146 | 22,51292 |
| + | sp\|P30040\|ERP29_HUMAN;tr\|F8VY02\|F8VY02_HUMAN;tr\|F8W1G0\|F8W1G0_HUMAN | sp\|P30040\|E | 7,0946 | 38650000 | 12 | 4 | 0,00203031 | -1,10064936 | 21,43201 | 21,85871 | 21,97386 | 21,90228 | 20,89112 | 21,01842 | 20,23538 | 20,61935 |
| + | sp\|P09012\|SNRPA_HUMAN;tr\|M0R268\|M0R268_HUMAN;tr\|M0R0G9\|M0R0G9_HUMAN;tr\|N | sp\|P09012\|S | 6,5371 | 85966000 | 12 | 2 | 0,00214446 | -0,57025957 | 22,77167 | 22,55494 | 22,95807 | 22,8938 | 22,12174 | 22,22726 | 22,13655 | 22,41189 |

| | | | | | | | | | | | | | | | | |
|---|---|---|---|---|---|---|---|---|---|---|---|---|---|---|---|---|
| + | sp\|P26440\|IVD_HUMAN;tr\|A0A0A0MT83\|A0A0A0MT83_HUMAN;tr\|H0YKV0\|H0YKV0_HUMAI | sp\|P26440\|IV | 4,6962 | 19020000 | 8 | 2 | 0,00216746 | -1,15484476 | 20,78205 | 20,92904 | 20,88696 | 21,23433 | 20,15326 | 19,77703 | 19,47945 | NaN |
| + | sp\|Q6PI48\|SYDM_HUMAN | sp\|Q6PI48\|SY | 7,5667 | 24740000 | 8 | 5 | 0,00220421 | 1,04465151 | 19,7408 | 19,86497 | 19,51549 | 19,98663 | 20,60467 | 21,04946 | 21,19135 | 20,44101 |
| + | sp\|P52597\|HNRPF_HUMAN | sp\|P52597\|H | 187,36 | 115920000 | 19 | 4 | 0,00221584 | 0,53738356 | 22,58456 | 22,36284 | 22,63808 | 22,47013 | 23,06651 | 23,2029 | 23,12726 | 22,80847 |
| + | sp\|P62750\|RL23A_HUMAN;tr\|H7BY10\|H7BY10_HUMAN;tr\|K7EJV9\|K7EJV9_HUMAN;tr\|K7ERT | sp\|P62750\|R | 206,98 | 419160000 | 34 | 8 | 0,00229548 | -0,81980753 | 25,14048 | 25,3946 | 25,19242 | 24,90981 | 24,37504 | 24,68105 | 24,20628 | 24,09571 |
| + | sp\|P14868\|SYDC_HUMAN;tr\|C9J7S3\|C9J7S3_HUMAN;tr\|C9JLC1\|C9JLC1_HUMAN;tr\|C9JQM9\| | sp\|P14868\|S' | 31,406 | 139240000 | 30 | 11 | 0,00232226 | -0,26332855 | 23,25136 | 23,30438 | 23,43275 | 23,23997 | 23,01698 | 23,12367 | 22,99926 | 23,03525 |
| + | sp\|P09661\|RU2A_HUMAN;tr\|H0YKK0\|H0YKK0_HUMAN;tr\|H0YMA0\|H0YMA0_HUMAN;tr\|H0Y | sp\|P09661\|R | 59,101 | 61274000 | 15 | 6 | 0,00237895 | -0,30197191 | 22,30897 | 22,10789 | 22,2232 | 22,35531 | 21,97914 | 21,98992 | 21,9411 | 21,87733 |
| + | sp\|P35998\|PRS7_HUMAN;tr\|C9JLS9\|C9JLS9_HUMAN | sp\|P35998\|P | 72,267 | 194570000 | 33 | 15 | 0,00244237 | 0,51536226 | 23,04845 | 23,14436 | 23,37703 | 23,39008 | 23,7214 | 23,92687 | 23,6796 | 23,69349 |
| + | sp\|P61981\|1433G_HUMAN | sp\|P61981\|1 | 88,21 | 65321000 | 14 | 3 | 0,00253232 | -0,41030025 | 22,21386 | 22,29911 | 22,36327 | 22,34335 | 21,93844 | 21,87721 | 21,6996 | 22,06313 |
| + | sp\|P62424\|RL7A_HUMAN;tr\|Q5T8U3\|Q5T8U3_HUMAN;tr\|Q5T8U2\|Q5T8U2_HUMAN | sp\|P62424\|R | 150,81 | 550420000 | 42 | 11 | 0,0026583 | -0,68934727 | 25,66405 | 25,69988 | 25,73684 | 25,23524 | 25,1031 | 24,91766 | 24,77465 | 24,78322 |
| + | sp\|P04083\|ANXA1_HUMAN;tr\|Q5T3N1\|Q5T3N1_HUMAN;tr\|Q5T3N0\|Q5T3N0_HUMAN | sp\|P04083\|A | 46,482 | 27491000 | 13 | 5 | 0,00267362 | -0,80038357 | 21,1327 | 21,59094 | 21,48419 | 21,15777 | 20,38005 | 20,65368 | 20,32158 | 20,80876 |
| + | sp\|Q9BTT0\|AN32E_HUMAN;tr\|E9PPH5\|E9PPH5_HUMAN;tr\|Q5TB19\|Q5TB19_HUMAN;tr\|E9P | sp\|Q9BTT0\|A | 21,987 | 49568000 | 17 | 5 | 0,00271371 | -1,64076185 | 21,72655 | 22,66562 | 22,55166 | 22,56377 | 21,04434 | 21,27042 | 20,45443 | 20,17536 |
| + | sp\|Q6PGP7\|TTC37_HUMAN;tr\|D6RCE2\|D6RCE2_HUMAN | sp\|Q6PGP7\|T | 16,588 | 33843000 | 11 | 8 | 0,00273207 | 2,46581364 | 19,261 | 18,5908 | 18,8842 | 18,94341 | 21,42347 | 20,84053 | 22,73355 | 20,5451 |
| + | sp\|P42766\|RL35_HUMAN;tr\|F2Z388\|F2Z388_HUMAN | sp\|P42766\|R | 61,536 | 292940000 | 17 | 3 | 0,00285157 | -1,29214859 | 25,01014 | 25,0127 | 25,23787 | 24,09418 | 23,73892 | 23,50591 | 23,58967 | 23,35178 |
| + | sp\|P13667\|PDIA4_HUMAN | sp\|P13667\|P | 35,324 | 157100000 | 34 | 12 | 0,00289328 | -0,77079439 | 23,34109 | 23,77946 | 23,64119 | 23,66427 | 22,64742 | 23,10559 | 22,58472 | 23,00512 |
| + | sp\|P06733\|ENOA_HUMAN;tr\|K7EM90\|K7EM90_HUMAN;sp\|P13929\|ENOB_HUMAN;sp\|P091( | sp\|P06733\|E | 323,31 | 3310600000 | 92 | 19 | 0,00296564 | -0,52375031 | 27,98354 | 28,02454 | 28,06286 | 28,21002 | 27,34257 | 27,59413 | 27,45591 | 27,79334 |
| + | sp\|Q9BRK5\|CAB45_HUMAN;tr\|G3V1E2\|G3V1E2_HUMAN;tr\|H0Y3T6\|H0Y3T6_HUMAN | sp\|Q9BRK5\|C | 6,6076 | 38840000 | 14 | 4 | 0,00298467 | 1,11890507 | 20,34505 | 20,55056 | 20,4491 | 20,39686 | 22,00247 | 21,82912 | 21,40468 | 20,98092 |
| + | sp\|Q9H845\|ACAD9_HUMAN;tr\|H0Y8Z9\|H0Y8Z9_HUMAN;tr\|D6RCD8\|D6RCD8_HUMAN | sp\|Q9H845\|A | 4,5969 | 14291000 | 8 | 4 | 0,00310701 | 1,54251862 | 18,81863 | 18,43713 | 18,62881 | 18,91751 | 19,66293 | 20,60123 | 20,91612 | 19,79187 |
| + | sp\|P40926\|MDHM_HUMAN;tr\|G3XAL0\|G3XAL0_HUMAN | sp\|P40926\|M | 90,861 | 706180000 | 35 | 10 | 0,00311689 | -0,91548252 | 25,73575 | 25,9327 | 26,13128 | 26,29626 | 25,0912 | 25,19809 | 24,71508 | 25,42969 |
| + | sp\|Q8IXB1\|DJC10_HUMAN;tr\|E7EP04\|E7EP04_HUMAN;tr\|A0A087WXH7\|A0A087WXH7_HUM | sp\|Q8IXB1\|D | 121,25 | 129590000 | 23 | 11 | 0,00313407 | 2,47935692 | NaN | 21,34743 | 20,78883 | 20,92586 | 24,2712 | 24,01265 | 22,92582 | 22,79058 |
| + | sp\|O75306\|NDUS2_HUMAN | sp\|O75306\|N | 13,935 | 49756000 | 17 | 7 | 0,00315491 | 1,8867979 | 19,97603 | 20,15078 | 20,03255 | 20,23023 | 21,75103 | 22,41343 | 22,77925 | 20,99307 |
| + | tr\|K7EJT5\|K7EJT5_HUMAN;tr\|K7EP65\|K7EP65_HUMAN;tr\|K7EKS7\|K7EKS7_HUMAN;tr\|K7ELC4 | tr\|K7EJT5\|K7 | 11,737 | 142410000 | 11 | 2 | 0,00321051 | -0,76941967 | 23,71513 | 23,54581 | 23,80597 | 23,31493 | 22,898 | 23,09058 | 22,80843 | 22,50715 |
| + | sp\|P14618\|KPYM_HUMAN;tr\|H3BTN5\|H3BTN5_HUMAN;tr\|B4DNK4\|B4DNK4_HUMAN;tr\|H3B | sp\|P14618\|K | 323,31 | 1377900000 | 89 | 20 | 0,00321538 | -0,85748863 | 26,63893 | 27,03044 | 27,03842 | 27,07948 | 26,05654 | 26,21653 | 25,69254 | 26,39172 |
| + | sp\|P27348\|1433T_HUMAN;tr\|E9PG15\|E9PG15_HUMAN | sp\|P27348\|1 | 214,97 | 177110000 | 24 | 8 | 0,00338744 | -0,65762329 | 23,6903 | 23,9188 | 24,01299 | 23,87493 | 23,30215 | 23,45073 | 22,87185 | 23,24179 |
| + | tr\|J3QR09\|J3QR09_HUMAN;tr\|J3KTE4\|J3KTE4_HUMAN;sp\|P84098\|RL19_HUMAN;tr\|J3QL15\|. | tr\|J3QR09\|J3 | 40,287 | 291980000 | 9 | 3 | 0,00350743 | -0,61463118 | 24,66871 | 24,76527 | 25,02069 | 24,41893 | 24,03255 | 24,10925 | 24,22817 | 24,0451 |
| + | sp\|P05141\|ADT2_HUMAN | sp\|P05141\|A | 23,111 | 176950000 | 16 | 1 | 0,00361502 | 0,9159193 | 22,92136 | 22,65773 | 22,62582 | 22,8697 | 23,63243 | 24,15792 | 23,56139 | 23,29656 |
| + | sp\|P36957\|ODO2_HUMAN;tr\|Q86SW4\|Q86SW4_HUMAN | sp\|P36957\|O | 11,1 | 56567000 | 12 | 6 | 0,00366467 | -0,43711853 | 22,16346 | 21,95387 | 22,23634 | 22,07887 | 21,79027 | 21,58284 | 21,79935 | 21,51162 |
| + | tr\|H7C3I1\|H7C3I1_HUMAN;tr\|F6VDH7\|F6VDH7_HUMAN;tr\|Q3KNR6\|Q3KNR6_HUMAN;sp\|Q8 | tr\|H7C3I1\|H7 | 5,4402 | 63691000 | 16 | 3 | 0,0038984 | -0,64564466 | 22,16907 | 22,49346 | 22,4839 | 22,5395 | 21,64821 | 21,73769 | 21,61014 | 22,10732 |
| + | tr\|F8W1A4\|F8W1A4_HUMAN;sp\|P54819\|KAD2_HUMAN;tr\|F8VZG5\|F8VZG5_HUMAN;tr\|F8V | tr\|F8W1A4\|F | 77,785 | 228900000 | 41 | 10 | 0,00417223 | -0,90143299 | 24,14196 | 24,19144 | 24,30382 | 24,57213 | 23,41442 | 23,73593 | 22,90971 | 23,54357 |
| + | tr\|A0A0C4DFV9\|A0A0C4DFV9_HUMAN;sp\|Q01105\|SET_HUMAN;tr\|A0A087X027\|A0A087X027 | tr\|A0A0C4DF | 283,38 | 325600000 | 25 | 6 | 0,00423565 | -1,36027813 | 24,29334 | 25,26191 | 25,61172 | 24,98037 | 23,70988 | 23,57057 | 23,43097 | 23,99481 |
| + | sp\|Q6P1M0\|S27A4_HUMAN | sp\|Q6P1M0\|! | 3,0704 | 6965400 | 5 | 3 | 0,00433192 | 0,38803212 | 18,7091 | 18,49635 | 18,57913 | NaN | 18,90416 | 19,0631 | 19,07517 | 18,88914 |
| + | sp\|P60842\|IF4A1_HUMAN;tr\|J3KT12\|J3KT12_HUMAN;tr\|J3KTB5\|J3KTB5_HUMAN;tr\|J3QL43\|. | sp\|P60842\|IF | 252,46 | 1187500000 | 89 | 9 | 0,00437112 | -0,53152418 | 26,33514 | 26,54597 | 26,62171 | 26,54146 | 25,9608 | 26,10536 | 25,69726 | 26,15476 |
| + | sp\|P13639\|EF2_HUMAN;tr\|K7EJ74\|K7EJ74_HUMAN;tr\|K7EP67\|K7EP67_HUMAN | sp\|P13639\|E | 323,31 | 3653900000 | 150 | 38 | 0,00438507 | -0,41952324 | 28,11123 | 28,08806 | 28,15957 | 28,14836 | 27,68732 | 27,71424 | 27,48639 | 27,94117 |
| + | sp\|P49321\|NASP_HUMAN;tr\|Q5T624\|Q5T624_HUMAN;tr\|H0YDS9\|H0YDS9_HUMAN;tr\|E9PR | sp\|P49321\|N | 71,932 | 289350000 | 29 | 12 | 0,00454214 | -0,67090178 | 24,68859 | 24,55713 | 24,60371 | 24,55292 | 23,62344 | 23,95828 | 23,81039 | 24,32663 |
| + | sp\|P63104\|1433Z_HUMAN;tr\|B0AZS6\|B0AZS6_HUMAN;tr\|E7EX29\|E7EX29_HUMAN;tr\|B7Z2E | sp\|P63104\|1 | 323,31 | 398910000 | 46 | 8 | 0,00461586 | -0,59601402 | 25,01945 | 25,09933 | 25,03246 | 25,07867 | 24,58675 | 24,77826 | 24,20105 | 24,2798 |
| + | tr\|F5GXX5\|F5GXX5_HUMAN;sp\|P61803\|DAD1_HUMAN;tr\|F5H895\|F5H895_HUMAN;tr\|A0A0E | tr\|F5GXX5\|F | 16,966 | 66729000 | 7 | 4 | 0,00471883 | -1,00984653 | 22,0036 | NaN | 22,54903 | 22,21285 | 21,52401 | 21,39765 | 20,9113 | 21,1483 |
| + | sp\|Q9BVA1\|TBB2B_HUMAN;tr\|G3V5W4\|G3V5W4_HUMAN;tr\|G3V2R8\|G3V2R8_HUMAN;tr\|G | sp\|Q9BVA1\|T | 25,907 | 407880000 | 9 | 2 | 0,00471946 | 1,96319723 | 22,63057 | 22,86524 | 22,70725 | 22,93912 | 24,84211 | 23,66362 | 25,82936 | 24,65987 |
| + | sp\|O75694\|NU155_HUMAN;tr\|E9PF10\|E9PF10_HUMAN | sp\|O75694\|N | 20,56 | 83978000 | 26 | 12 | 0,00478979 | -0,74694872 | 23,06062 | 22,76203 | 22,42452 | 23,11862 | 22,19003 | 21,93049 | 22,19743 | 22,06006 |
| + | tr\|M0R3D6\|M0R3D6_HUMAN;tr\|M0R1A7\|M0R1A7_HUMAN;tr\|M0R117\|M0R117_HUMAN;sp | tr\|M0R3D6\|N | 57,728 | 287110000 | 28 | 4 | 0,00483517 | -0,61495972 | 24,56483 | 24,63893 | 24,61526 | 24,22869 | 23,79233 | 23,9178 | 24,17904 | 23,69869 |
| + | tr\|H3BMH2\|H3BMH2_HUMAN;tr\|H3BSC1\|H3BSC1_HUMAN;sp\|P62491\|RB11A_HUMAN;sp\|Q | tr\|H3BMH2\|I | 6,41 | 117870000 | 17 | 5 | 0,00493035 | -0,62263441 | 23,1275 | 23,41274 | 23,20131 | 23,36693 | 22,73246 | 22,82621 | 22,27782 | 22,78145 |
| + | sp\|P41227\|NAA10_HUMAN;tr\|F8W808\|F8W808_HUMAN;tr\|A8MWP7\|A8MWP7_HUMAN;tr\|Q5T | sp\|P41227\|N | 16,91 | 28980000 | 8 | 3 | 0,00493732 | -0,4614954 | 21,10942 | 21,06815 | 21,25032 | 21,26802 | 20,7381 | 20,93272 | 20,47338 | 20,70573 |
| + | sp\|P41091\|IF2G_HUMAN;tr\|F8W810\|F8W810_HUMAN;sp\|Q2VIR3\|IF2GL_HUMAN;tr\|H7BZU1 | sp\|P41091\|IF | 37,002 | 85161000 | 23 | 8 | 0,00497699 | -0,47553778 | 22,74337 | 22,53825 | 22,96395 | 22,62874 | 22,31396 | 22,35114 | 22,07874 | 22,22832 |
| + | sp\|P16152\|CBR1_HUMAN;tr\|E9PQ63\|E9PQ63_HUMAN;tr\|A8MTM1\|A8MTM1_HUMAN;sp\|O7 | sp\|P16152\|C | 104,5 | 80448000 | 24 | 7 | 0,00507766 | -0,79719257 | 22,55223 | 22,89624 | 22,7977 | 22,92823 | 21,90647 | 22,38601 | 21,60141 | 22,09175 |
| + | sp\|P04040\|CATA_HUMAN | sp\|P04040\|C | 6,7078 | 23459000 | 12 | 5 | 0,00508515 | -0,71529865 | 20,85575 | 21,24464 | 21,00313 | 20,94592 | 20,47912 | 20,57487 | 19,93905 | 20,1952 |
| + | tr\|G3V198\|G3V198_HUMAN;sp\|Q12769\|NU160_HUMAN;tr\|E9PR16\|E9PR16_HUMAN;tr\|E9P | tr\|G3V198\|G | 52,738 | 47681000 | 18 | 8 | 0,00519233 | 1,36075211 | 20,50659 | 20,59195 | 20,48787 | 20,38617 | 21,08401 | 22,03275 | 22,59149 | 21,70734 |
| + | tr\|Q32Q12\|Q32Q12_HUMAN;sp\|P22392\|NDKB_HUMAN;tr\|J3KPD9\|J3KPD9_HUMAN;sp\|O60 | tr\|Q32Q12\|C | 117,44 | 760700000 | 48 | 2 | 0,00525774 | -1,00419712 | 26,03103 | 25,87371 | 25,97612 | 26,46234 | 25,34736 | 25,48726 | 24,72338 | 24,7684 |
| + | tr\|I3L397\|I3L397_HUMAN;sp\|P63241\|IF5A1_HUMAN;tr\|I3L504\|I3L504_HUMAN;sp\|Q6IS14\|IF | tr\|I3L397\|I3L | 164,62 | 458020000 | 31 | 6 | 0,00525882 | -0,93362427 | 25,09865 | 25,49677 | 25,31731 | 25,49945 | 24,53831 | 24,92506 | 24,09031 | 24,124 |
| + | sp\|P38646\|GRP75_HUMAN;tr\|D6RJI2\|D6RJI2_HUMAN;tr\|D6RA73\|D6RA73_HUMAN;tr\|H0Y8S | sp\|P38646\|G | 323,31 | 1118300000 | 98 | 23 | 0,00529435 | 0,22913885 | 26,06723 | 26,00329 | 25,9514 | 26,17467 | 26,3195 | 26,21334 | 26,30895 | 26,27135 |
| + | sp\|P25205\|MCM3_HUMAN;tr\|J3KQ69\|J3KQ69_HUMAN;tr\|Q7Z6P5\|Q7Z6P5_HUMAN | sp\|P25205\|M | 84,668 | 190210000 | 53 | 21 | 0,00531815 | 0,30282688 | 23,62098 | 23,36145 | 23,41712 | 23,47469 | 23,6609 | 23,74396 | 23,85994 | 23,82075 |
| + | tr\|M0QXU7\|M0QXU7_HUMAN;sp\|O43615\|TIM44_HUMAN;tr\|M0R301\|M0R301_HUMAN | tr\|M0QXU7\|I | 12,842 | 61540000 | 9 | 5 | 0,00534169 | -0,61178923 | 22,72386 | 22,12698 | 22,31083 | 22,36022 | 21,75071 | 21,92506 | 21,58522 | 21,81374 |
| + | tr\|A0A087WWJ2\|A0A087WWJ2_HUMAN;tr\|C9J5D1\|C9J5D1_HUMAN;tr\|E7EQ69\|E7EQ69_HUI | tr\|A0A087W\ | 2,4156 | 9659500 | 5 | 2 | 0,00534184 | -0,42299954 | 19,58327 | 19,6855 | 19,74394 | 19,83712 | 19,41844 | 19,29762 | NaN | 19,15232 |
| + | tr\|C9JRH2\|C9JRH2_HUMAN;tr\|C9JMJ4\|C9JMJ4_HUMAN;tr\|C9J3R0\|C9J3R0_HUMAN;tr\|C9JQZ | tr\|C9JRH2\|C9 | 11,338 | 41493000 | 7 | 3 | 0,00575809 | -0,47922182 | 21,31595 | 21,66631 | 21,74763 | 21,68551 | 21,26413 | 21,13496 | 20,97176 | 21,12767 |
| + | sp\|Q02878\|RL6_HUMAN;tr\|F8VZ45\|F8VZ45_HUMAN;tr\|U3KQR5\|U3KQR5_HUMAN;tr\|F8VR6 | sp\|Q02878\|R | 157,98 | 822330000 | 41 | 13 | 0,00581462 | -0,7302084 | 26,34669 | 26,27915 | 26,37003 | 25,76025 | 25,45861 | 25,25764 | 25,72669 | 25,39234 |
| + | sp\|Q01518\|CAP1_HUMAN;tr\|Q5T0R7\|Q5T0R7_HUMAN;tr\|Q5T0R6\|Q5T0R6_HUMAN;tr\|Q5T0 | sp\|Q01518\|C | 21,847 | 154660000 | 38 | 11 | 0,00584187 | -0,5572753 | 23,30466 | 23,68848 | 23,87014 | 23,6633 | 22,90674 | 23,10093 | 23,08594 | 23,20386 |
| + | sp\|P23528\|COF1_HUMAN;tr\|G3V1A4\|G3V1A4_HUMAN;tr\|E9PP50\|E9PP50_HUMAN;tr\|E9PK2 | sp\|P23528\|C | 287,43 | 486440000 | 36 | 9 | 0,00590807 | -1,09135199 | 25,41616 | 25,37951 | 25,21108 | 25,78995 | 24,12297 | 25,04097 | 24,02732 | 24,24002 |
| + | sp\|P46777\|RL5_HUMAN;tr\|Q5T7N0\|Q5T7N0_HUMAN | sp\|P46777\|R | 183,25 | 240400000 | 28 | 8 | 0,00594497 | -0,71329689 | 24,52773 | 24,0242 | 24,34245 | 24,37776 | 23,36279 | 23,52886 | 23,53611 | 23,99118 |
| + | sp\|P15531\|NDKA_HUMAN;tr\|E7ERL0\|E7ERL0_HUMAN;tr\|E5RHP0\|E5RHP0_HUMAN;tr\|C9K02 | sp\|P15531\|N | 31,581 | 70338000 | 14 | 1 | 0,00604461 | -1,16162062 | 22,36581 | 22,64315 | 23,06802 | 22,65347 | 21,7205 | 22,01869 | 21,45573 | 20,88904 |
| + | sp\|P63173\|RL38_HUMAN;tr\|J3KT73\|J3KT73_HUMAN;tr\|J3QL01\|J3QL01_HUMAN;tr\|J3KSP2\|J: | sp\|P63173\|R | 11,243 | 406450000 | 19 | 5 | 0,00609778 | 1,13730001 | 23,50191 | 23,94976 | 23,89264 | 23,88558 | 25,4788 | 25,23301 | 24,72536 | 24,34191 |
| + | sp\|Q9NR30\|DDX21_HUMAN | sp\|Q9NR30\|[ | 108,78 | 604670000 | 82 | 24 | 0,00617578 | -0,56758451 | 25,65952 | 25,60493 | 25,75307 | 25,49317 | 24,87633 | 24,87465 | 25,41119 | 25,07818 |
| + | tr\|E7EVA0\|E7EVA0_HUMAN;sp\|P27816\|MAP4_HUMAN;tr\|H7C4C5\|H7C4C5_HUMAN;tr\|H0Y2 | tr\|E7EVA0\|E | 52,996 | 93443000 | 29 | 9 | 0,00636403 | -0,47696733 | 22,9446 | 22,90139 | 23,06112 | 22,67161 | 22,19668 | 22,41186 | 22,46852 | 22,5938 |
| + | sp\|P06744\|G6PI_HUMAN;tr\|A0A0A0MTS2\|A0A0A0MTS2_HUMAN;tr\|K7EQ48\|K7EQ48_HUMA | sp\|P06744\|G | 83,289 | 327060000 | 27 | 9 | 0,00637343 | -0,93017721 | 24,44474 | 24,79576 | 25,24576 | 24,97259 | 23,85785 | 23,99774 | 23,57358 | 24,30897 |
| + | sp\|P52209\|6PGD_HUMAN;tr\|K7EMN2\|K7EMN2_HUMAN;tr\|K7EM49\|K7EM49_HUMAN;tr\|K7I | sp\|P52209\|6 | 16,679 | 70379000 | 11 | 6 | 0,00651081 | -0,80250645 | 22,48753 | 22,34308 | 22,86931 | 22,70774 | 21,90511 | 22,03668 | 21,33172 | 21,92412 |
| + | sp\|Q8NFH3\|NUP43_HUMAN | sp\|Q8NFH3\|I | 2,9215 | 15149000 | 6 | 2 | 0,00688356 | 0,48616473 | NaN | 19,78539 | 19,80945 | 19,92988 | 20,3626 | 20,45222 | NaN | 20,1684 |
| + | sp\|P63244\|RACK1_HUMAN;tr\|H0YAF8\|H0YAF8_HUMAN;tr\|D6R9Z1\|D6R9Z1_HUMAN;tr\|H0YA | sp\|P63244\|R | 114,79 | 298750000 | 22 | 4 | 0,00716314 | 0,55865669 | 23,57207 | 24,04935 | 23,97762 | 23,86089 | 24,58463 | 24,58049 | 24,28905 | 24,24038 |
| + | tr\|D3YTB1\|D3YTB1_HUMAN;sp\|P62910\|RL32_HUMAN;tr\|F8W727\|F8W727_HUMAN | tr\|D3YTB1\|D | 18,418 | 204150000 | 15 | 3 | 0,00733527 | -0,81990623 | 24,52337 | 24,34069 | 24,28221 | 23,68368 | 23,63201 | 23,45601 | 23,22897 | 23,23333 |
| + | sp\|P23246\|SFPQ_HUMAN;tr\|H0Y9K7\|H0Y9K7_HUMAN;tr\|H0Y9U2\|H0Y9U2_HUMAN | sp\|P23246\|S | 294,95 | 938820000 | 86 | 18 | 0,00743994 | -0,49545765 | 26,2637 | 26,058 | 26,23456 | 26,15143 | 25,63857 | 25,52079 | 25,54483 | 26,02168 |
| + | sp\|Q04837\|SSBP_HUMAN;tr\|C9K0U8\|C9K0U8_HUMAN;tr\|A0A0G2JLD8\|A0A0G2JLD8_HUMAN | sp\|Q04837\|S | 5,1814 | 171810000 | 9 | 4 | 0,00746978 | -0,69866276 | 23,90701 | 23,86183 | 23,8187 | 23,82182 | 23,59889 | 23,24674 | 22,79223 | 22,97685 |
| + | tr\|H0YN26\|H0YN26_HUMAN;sp\|P39687\|AN32A_HUMAN;tr\|H7BZ09\|H7BZ09_HUMAN;sp\|O9 | tr\|H0YN26\|H | 27,805 | 108770000 | 15 | 4 | 0,00787889 | -1,06729269 | 22,72821 | 22,96001 | 23,74704 | 23,58646 | 22,19442 | 22,28726 | 21,85125 | 22,41962 |
| + | tr\|C9JXB8\|C9JXB8_HUMAN;tr\|C9JNW5\|C9JNW5_HUMAN;sp\|P83731\|RL24_HUMAN | tr\|C9JXB8\|C9 | 30,86 | 147330000 | 11 | 4 | 0,00838501 | -0,91138458 | 23,69051 | 23,92561 | 23,82406 | 23,35595 | 23,15675 | 22,5255 | 22,68069 | NaN |
| + | sp\|P28066\|PSA5_HUMAN | sp\|P28066\|P | 51,687 | 95658000 | 12 | 6 | 0,00848357 | -0,30734094 | 22,69485 | 22,86435 | 22,76893 | 22,8121 | 22,48013 | 22,59991 | 22,3531 | NaN |

| | | | | | | | | | | | | | | | | |
|---|---|---|---|---|---|---|---|---|---|---|---|---|---|---|---|---|
| + | sp\|Q15427\|SF3B4_HUMAN;tr\|Q5SZ64\|Q5SZ64_HUMAN | sp\|Q15427\|S | 17,664 | 17603000 | 7 | 4 | 0,00886468 | -0,30121565 | 20,40343 | 20,15165 | 20,29109 | 20,12956 | 19,82299 | 20,043 | 19,92829 | 19,97659 |
| + | sp\|Q92522\|H1X_HUMAN | sp\|Q92522\|H | 25,677 | 78590000 | 16 | 4 | 0,00887523 | -0,80284452 | 22,50753 | 23,18345 | 23,0112 | 22,38603 | 22,04103 | 21,88401 | 22,16984 | 21,78197 |
| + | sp\|Q9H078\|CLPB_HUMAN;tr\|H0YGM0\|H0YGM0_HUMAN;tr\|F5H392\|F5H392_HUMAN | sp\|Q9H078\|C | 16,662 | 27626000 | 9 | 2 | 0,00903796 | 0,83583482 | NaN | 20,5682 | 19,90029 | 20,27401 | 21,20126 | 21,03234 | 21,2815 | 20,81824 |
| + | sp\|Q9P2J5\|SYLC_HUMAN;tr\|A0A087WXY1\|A0A087WXY1_HUMAN | sp\|Q9P2J5\|S | 9,6947 | 52475000 | 21 | 6 | 0,00905716 | -0,52890587 | 22,15354 | 22,30129 | 21,93038 | 22,08232 | 21,30952 | 21,48513 | 21,80487 | 21,75238 |
| + | sp\|Q00688\|FKBP3_HUMAN;tr\|G3V5F2\|G3V5F2_HUMAN | sp\|Q00688\|F | 4,2708 | 43805000 | 8 | 3 | 0,00912528 | -0,814116 | 21,42578 | 21,9272 | 22,0353 | 22,38422 | 21,18659 | 21,2502 | 20,88122 | 21,19802 |
| + | sp\|Q96CW5\|GCP3_HUMAN;tr\|A0A087WWB5\|A0A087WWB5_HUMAN;tr\|A0A087WU06\|A0A | sp\|Q96CW5\| | 7,528 | 13511000 | 6 | 5 | 0,00918123 | 0,45266469 | NaN | 19,4974 | 19,31115 | 19,40195 | 19,82842 | 19,73455 | 20,00553 | NaN |
| + | tr\|F8W914\|F8W914_HUMAN;sp\|Q9NQC3\|RTN4_HUMAN;tr\|H7C106\|H7C106_HUMAN;tr\|A0A | tr\|F8W914\|F | 19,944 | 91502000 | 19 | 5 | 0,00934968 | -0,46598196 | 22,72939 | 22,97072 | 22,90303 | 22,8348 | 22,36172 | 22,45043 | 22,6525 | 22,10936 |
| + | sp\|P31939\|PUR9_HUMAN;tr\|H7C1S2\|H7C1S2_HUMAN;tr\|F8WEF0\|F8WEF0_HUMAN;tr\|C9JLK | sp\|P31939\|P | 112,65 | 162830000 | 37 | 16 | 0,00954089 | -0,70534801 | 23,59275 | 23,6872 | 23,82464 | 23,96499 | 22,8678 | 23,1861 | 22,71614 | 23,47816 |
| + | tr\|E7EU96\|E7EU96_HUMAN;sp\|Q8NEV1\|CSK23_HUMAN;sp\|P68400\|CSK21_HUMAN;tr\|Q5U5 | tr\|E7EU96\|E7 | 5,4784 | 20225000 | 7 | 4 | 0,00964983 | -0,6264658 | 20,30263 | 20,74329 | 20,96169 | 20,696 | 19,78879 | 20,24656 | 20,11553 | 20,04687 |
| + | tr\|E7EQR4\|E7EQR4_HUMAN;sp\|P15311\|EZRI_HUMAN | tr\|E7EQR4\|E7 | 111,24 | 272880000 | 35 | 7 | 0,00967817 | -0,4265728 | 24,29313 | 24,39034 | 24,45098 | 24,57837 | 23,86041 | 24,13399 | 23,81078 | 24,20135 |
| + | sp\|P57088\|TMM33_HUMAN;tr\|H0Y8N0\|H0Y8N0_HUMAN;tr\|D6RAA6\|D6RAA6_HUMAN | sp\|P57088\|T | 5,296 | 62096000 | 11 | 3 | 0,00974205 | 0,7141223 | 21,5443 | 21,66049 | 21,668 | 21,24284 | 22,618 | 22,1134 | 22,37894 | 21,86178 |
| + | sp\|O75616\|ERAL1_HUMAN;tr\|J3QT61\|J3QT61_HUMAN;tr\|J3QSB2\|J3QSB2_HUMAN;tr\|J3QRV | sp\|O75616\|E | 4,9013 | 25529000 | 5 | 3 | 0,01008015 | 0,51378012 | 20,01672 | 20,47289 | 20,37369 | 20,33625 | 20,78748 | 21,09352 | 20,72709 | 20,64658 |
| + | sp\|Q9H9Y6\|RPA2_HUMAN;tr\|C9JJG2\|C9JJG2_HUMAN;tr\|C9JS83\|C9JS83_HUMAN | sp\|Q9H9Y6\|F | 6,4863 | 20118000 | 10 | 6 | 0,01017488 | 0,94793272 | 19,78112 | 19,59244 | 19,18401 | 19,64091 | 19,93027 | 20,707 | 20,96247 | 20,39048 |
| + | sp\|Q92769\|HDAC2_HUMAN;tr\|E5RFI6\|E5RFI6_HUMAN;tr\|E5RJ04\|E5RJ04_HUMAN;tr\|E5RGV4 | sp\|Q92769\|H | 12,714 | 95073000 | 10 | 1 | 0,01028619 | 0,38362932 | 22,05947 | 22,11181 | 22,19487 | 21,997 | 22,30669 | 22,44038 | 22,74772 | 22,40289 |
| + | sp\|P39023\|RL3_HUMAN;tr\|G5E9G0\|G5E9G0_HUMAN;tr\|B5MCW2\|B5MCW2_HUMAN;tr\|H7C | sp\|P39023\|R | 305,86 | 560160000 | 34 | 8 | 0,01032995 | -0,53735399 | 25,57516 | 25,39617 | 25,56666 | 25,1653 | 24,8775 | 24,58852 | 25,10113 | 24,98672 |
| + | sp\|P27695\|APEX1_HUMAN;tr\|G3V3M6\|G3V3M6_HUMAN;tr\|G3V5M0\|G3V5M0_HUMAN;tr\|G | sp\|P27695\|A | 35,541 | 56693000 | 19 | 6 | 0,01039999 | -0,81561899 | 21,97572 | 22,09607 | 22,29751 | 22,23617 | 21,2577 | 21,38548 | 20,84007 | 21,85973 |
| + | sp\|P62258\|1433E_HUMAN;tr\|K7EM20\|K7EM20_HUMAN;tr\|K7EIT4\|K7EIT4_HUMAN;tr\|B4DJF | sp\|P62258\|1 | 302,98 | 1625500000 | 59 | 16 | 0,01060341 | -0,54405785 | 26,75595 | 26,89412 | 27,17988 | 26,94347 | 26,22172 | 26,74535 | 26,25818 | 26,37194 |
| + | sp\|P54577\|SYYC_HUMAN;tr\|A0A0C4DGZ5\|A0A0C4DGZ5_HUMAN | sp\|P54577\|S | 46,158 | 270150000 | 30 | 15 | 0,01068888 | -0,57543516 | 24,32389 | 24,38311 | 24,58852 | 24,54657 | 23,96807 | 23,83932 | 23,52013 | 24,21282 |
| + | sp\|Q8TEQ6\|GEMI5_HUMAN | sp\|Q8TEQ6\|G | 8,7732 | 25150000 | 10 | 7 | 0,01124197 | 0,63502884 | 20,20192 | 20,25523 | 20,37994 | 19,8994 | 20,64869 | 20,73686 | 21,24464 | 20,64641 |
| + | tr\|H3BQF1\|H3BQF1_HUMAN;sp\|P07741\|APT_HUMAN;tr\|H3BQB1\|H3BQB1_HUMAN;tr\|H3BS | tr\|H3BQF1\|H | 15,393 | 172950000 | 30 | 6 | 0,01125575 | -0,7317729 | 23,71513 | 23,8531 | 23,72151 | 24,11347 | 23,41738 | 23,40795 | 22,67618 | 22,9746 |
| + | sp\|P28074\|PSB5_HUMAN;tr\|H0YJM8\|H0YJM8_HUMAN | sp\|P28074\|P | 57,37 | 161110000 | 30 | 8 | 0,01141585 | -0,47265816 | 23,55309 | 23,57011 | 23,82833 | 23,6936 | 23,4438 | 23,12044 | 22,90663 | 23,28362 |
| + | sp\|Q9UQE7\|SMC3_HUMAN | sp\|Q9UQE7\|S | 5,0321 | 21308000 | 7 | 5 | 0,01141772 | -0,35085154 | 20,56607 | 20,50795 | 20,32433 | 20,35423 | 20,01468 | 20,02296 | 20,22423 | NaN |
| + | sp\|P60228\|EIF3E_HUMAN;tr\|H0YBR5\|H0YBR5_HUMAN;tr\|E5RGA2\|E5RGA2_HUMAN;tr\|E5RIT | sp\|P60228\|E | 10,642 | 79615000 | 14 | 6 | 0,01197811 | -0,32090902 | 22,55457 | 22,52378 | 22,71846 | 22,59439 | 22,29499 | 22,11994 | 22,20281 | 22,48981 |
| + | tr\|A0A075B730\|A0A075B730_HUMAN;tr\|A0A087X1U6\|A0A087X1U6_HUMAN;sp\|P58107\|EPI | tr\|A0A075B7 | 23,268 | 49679000 | 18 | 7 | 0,01210063 | -1,1282258 | 21,77318 | 22,7017 | 22,34199 | 22,00768 | 20,61423 | 21,34716 | 21,63125 | 20,719 |
| + | sp\|Q9NR31\|SAR1A_HUMAN;tr\|H0Y5E8\|H0Y5E8_HUMAN | sp\|Q9NR31\|S | 33,657 | 313790000 | 18 | 3 | 0,01214241 | -1,40760279 | 24,64488 | 24,82818 | 25,44503 | 24,95762 | 24,42073 | 23,46513 | 23,68024 | 22,67919 |
| + | sp\|O14744\|ANM5_HUMAN;tr\|C9JSX3\|C9JSX3_HUMAN;tr\|H0YJX6\|H0YJX6_HUMAN;tr\|G3V58 | sp\|O14744\|A | 94,102 | 108610000 | 14 | 8 | 0,01222411 | 0,3037281 | 22,59864 | 22,57448 | 22,54026 | 22,5242 | 23,03382 | 22,9287 | 22,63629 | 22,85369 |
| + | sp\|O95202\|LETM1_HUMAN | sp\|O95202\|L | 4,6557 | 25495000 | 7 | 4 | 0,01226566 | -0,87552261 | 20,60919 | 21,4314 | 20,94235 | 21,65373 | 20,15882 | 20,27776 | 20,2024 | 20,49561 |
| + | tr\|C9JG87\|C9JG87_HUMAN;sp\|Q9NYK5\|RM39_HUMAN | tr\|C9JG87\|C9 | 3,5846 | 9901600 | 5 | 2 | 0,01226851 | -0,34299564 | 19,5514 | 19,61924 | 19,65517 | 19,46653 | 19,21817 | 19,25622 | 19,00939 | 19,43659 |
| + | sp\|P04080\|CYTB_HUMAN | sp\|P04080\|C | 8,4856 | 52927000 | 11 | 2 | 0,01247187 | -1,51815192 | 22,52652 | 22,43639 | 22,5004 | 22,25908 | NaN | 21,79249 | 20,19484 | 20,75001 |
| + | sp\|Q15031\|SYLM_HUMAN;tr\|C9JYR8\|C9JYR8_HUMAN;tr\|E9PHM2\|E9PHM2_HUMAN | sp\|Q15031\|S | 2,6913 | 16462000 | 5 | 2 | 0,01264495 | 0,80585384 | 19,60771 | 19,53096 | 19,40775 | 19,62038 | 20,89689 | 20,52888 | 19,95291 | 20,01154 |
| + | sp\|O43707\|ACTN4_HUMAN;tr\|F5GXS2\|F5GXS2_HUMAN;tr\|H7C144\|H7C144_HUMAN;tr\|G3V | sp\|O43707\|A | 81,1 | 167120000 | 38 | 11 | 0,01274842 | -0,4687252 | 23,82036 | 23,58715 | 23,78776 | 23,75124 | 23,24787 | 23,24136 | 22,99099 | 23,59138 |
| + | sp\|P26038\|MOES_HUMAN;tr\|V9GZ54\|V9GZ54_HUMAN | sp\|P26038\|M | 22,192 | 96948000 | 11 | 8 | 0,01288976 | -0,49819851 | 22,64823 | 22,93971 | 22,94244 | 22,7666 | 22,26705 | 22,2854 | 22,08261 | 22,66912 |
| + | tr\|A0A024R4M0\|A0A024R4M0_HUMAN;sp\|P46781\|RS9_HUMAN;tr\|B5MCT8\|B5MCT8_HUMA | tr\|A0A024R4 | 26,537 | 816250000 | 34 | 11 | 0,01295552 | -0,61784029 | 26,14367 | 26,34349 | 26,09523 | 25,72578 | 25,76476 | 25,37156 | 25,51111 | 25,18937 |
| + | sp\|P62269\|RS18_HUMAN;tr\|A0A0G2JQH2\|A0A0G2JQH2_HUMAN;tr\|Q5GGW2\|Q5GGW2_HU | sp\|P62269\|R | 291,58 | 1624400000 | 41 | 10 | 0,01302904 | 0,57323694 | 26,17465 | 26,64828 | 26,3186 | 26,05759 | 27,1671 | 26,69471 | 26,7753 | 26,85496 |
| + | sp\|Q9UBT2\|SAE2_HUMAN;tr\|U3KQ93\|U3KQ93_HUMAN;tr\|K7ESK7\|K7ESK7_HUMAN;tr\|K7EPI | sp\|Q9UBT2\|S | 39,042 | 31281000 | 16 | 8 | 0,0130669 | -0,71759987 | 21,50548 | 21,50208 | 21,49194 | 21,24621 | 20,31617 | 21,01903 | 20,82128 | NaN |
| + | tr\|A0A087WT27\|A0A087WT27_HUMAN;tr\|J3KN95\|J3KN95_HUMAN;sp\|O95394\|AGM1_HUM | tr\|A0A087WT | 37,437 | 20869000 | 13 | 4 | 0,01308845 | 1,63961124 | 19,44113 | 19,18255 | 19,38 | 17,96871 | 20,18192 | 21,0229 | 21,32092 | 20,00512 |
| + | tr\|A0A0D9SF54\|A0A0D9SF54_HUMAN;sp\|Q13813\|SPTN1_HUMAN;tr\|A0A0D9SGF6\|A0A0D9S | tr\|A0A0D9SF | 30,093 | 75435000 | 29 | 16 | 0,01327359 | -0,60460758 | 22,92732 | 22,69183 | 22,60815 | 22,25733 | 21,91951 | 21,78085 | 22,10447 | 22,26138 |
| + | tr\|C9J4M6\|C9J4M6_HUMAN;tr\|C9J2Y9\|C9J2Y9_HUMAN;sp\|P30876\|RPB2_HUMAN | tr\|C9J4M6\|C | 15,555 | 37161000 | 15 | 10 | 0,01328791 | 1,78423198 | 19,76812 | NaN | 19,69459 | 19,84144 | 20,83969 | 21,52176 | 22,68027 | 21,16742 |
| + | sp\|Q15366\|PCBP2_HUMAN;tr\|H3BRU6\|H3BRU6_HUMAN;tr\|F8VZX2\|F8VZX2_HUMAN;tr\|F8W | sp\|Q15366\|P | 37,708 | 142510000 | 22 | 6 | 0,01333092 | -0,48533535 | 23,2339 | 23,33892 | 23,60093 | 23,62344 | 22,82712 | 23,08391 | 22,76048 | 23,18433 |
| + | tr\|F8VZJ2\|F8VZJ2_HUMAN;tr\|F8W0W4\|F8W0W4_HUMAN;tr\|H0YHX9\|H0YHX9_HUMAN;sp\|Q | tr\|F8VZJ2\|F8 | 9,0649 | 98724000 | 14 | 4 | 0,013415 | -0,98908854 | 22,62631 | 23,05909 | 23,83663 | 23,04589 | 22,34262 | 22,16555 | 21,77096 | 22,33243 |
| + | sp\|P40429\|RL13A_HUMAN;tr\|M0QYS1\|M0QYS1_HUMAN;sp\|Q6NVV1\|R13P3_HUMAN;tr\|Q8J | sp\|P40429\|R | 7,4493 | 229640000 | 18 | 4 | 0,01351677 | -0,68699265 | 24,23922 | 24,66628 | 24,31181 | 24,03364 | 24,00333 | 23,47853 | 23,70398 | 23,31714 |
| + | tr\|C9J9K3\|C9J9K3_HUMAN;sp\|P08865\|RSSA_HUMAN;tr\|A0A0C4DG17\|A0A0C4DG17_HUMAN | tr\|C9J9K3\|C9 | 323,31 | 465250000 | 45 | 13 | 0,01380034 | -0,61745739 | 25,12372 | 25,1106 | 25,23517 | 25,43759 | 24,64807 | 24,57358 | 24,21082 | 25,00479 |
| + | sp\|P61247\|RS3A_HUMAN;tr\|D6RG13\|D6RG13_HUMAN;tr\|D6RAT0\|D6RAT0_HUMAN;tr\|H0Y9 | sp\|P61247\|R | 300,37 | 500570000 | 43 | 8 | 0,01415593 | -0,31191349 | 25,30998 | 25,18658 | 25,34547 | 25,03523 | 24,89855 | 24,9795 | 25,00517 | 24,74638 |
| + | sp\|P22102\|PUR2_HUMAN;tr\|C9JBJ1\|C9JBJ1_HUMAN;tr\|C9JTV6\|C9JTV6_HUMAN;tr\|F8WD69\| | sp\|P22102\|P | 111,58 | 375700000 | 71 | 22 | 0,01419263 | -0,26649475 | 24,7305 | 24,85291 | 24,83543 | 24,77671 | 24,42349 | 24,4222 | 24,55374 | 24,73014 |
| + | tr\|J3KPX7\|J3KPX7_HUMAN;sp\|Q99623\|PHB2_HUMAN;tr\|F5GY37\|F5GY37_HUMAN;tr\|F5H3X | tr\|J3KPX7\|J3 | 21,622 | 68735000 | 16 | 10 | 0,01419859 | 1,23408556 | 21,43268 | 20,61243 | 20,85003 | 21,46493 | 21,68693 | 23,0925 | 22,34981 | 22,16717 |
| + | sp\|P00558\|PGK1_HUMAN;sp\|P07205\|PGK2_HUMAN | sp\|P00558\|P | 299,99 | 841260000 | 59 | 14 | 0,0152625 | -0,66768885 | 25,775 | 26,29556 | 26,13046 | 26,34518 | 25,4073 | 25,68563 | 25,06571 | 25,7168 |
| + | sp\|Q8IYD1\|ERF3B_HUMAN | sp\|Q8IYD1\|E | 14,07 | 44400000 | 14 | 1 | 0,01529154 | -0,28772068 | 21,60589 | 21,5904 | 21,81409 | 21,55689 | 21,29401 | 21,35896 | 21,23649 | 21,52693 |
| + | sp\|P62314\|SMD1_HUMAN;tr\|J3QLR7\|J3QLR7_HUMAN;tr\|J3QLI9\|J3QLI9_HUMAN | sp\|P62314\|S | 4,9921 | 51791000 | 13 | 2 | 0,01542155 | -0,52858877 | 22,21836 | 22,01638 | 22,10053 | 21,86242 | 21,17164 | 21,50281 | 21,55974 | 21,84915 |
| + | sp\|P51665\|PSMD7_HUMAN;tr\|H3BNT7\|H3BNT7_HUMAN;tr\|H3BTM8\|H3BTM8_HUMAN | sp\|P51665\|P | 8,111 | 34312000 | 6 | 6 | 0,01563391 | -0,47063828 | 21,44743 | 21,4894 | 21,24039 | 21,31976 | 20,66718 | 20,9132 | 21,13044 | NaN |
| + | sp\|Q07021\|C1QBP_HUMAN;tr\|I3L3Q7\|I3L3Q7_HUMAN;tr\|I3L3B0\|I3L3B0_HUMAN | sp\|Q07021\|C | 298,9 | 201850000 | 24 | 6 | 0,015653 | -0,48895836 | 24,0023 | 23,71858 | 24,33722 | 23,95056 | 23,54981 | 23,70092 | 23,41571 | 23,3864 |
| + | sp\|P62316\|SMD2_HUMAN;tr\|K7ERG4\|K7ERG4_HUMAN;tr\|K7EJB5\|K7EJB5_HUMAN | sp\|P62316\|S | 11,502 | 39174000 | 10 | 3 | 0,01600694 | -0,33204015 | 21,84311 | 21,76498 | 21,76085 | NaN | 21,38237 | 21,61432 | 21,37613 | NaN |
| + | sp\|P61978\|HNRPK_HUMAN;tr\|Q5T6W2\|Q5T6W2_HUMAN;tr\|S4R457\|S4R457_HUMAN;tr\|S4 | sp\|P61978\|H | 294,95 | 1043200000 | 90 | 17 | 0,01605701 | -0,27533436 | 26,08347 | 25,98941 | 26,26166 | 26,33657 | 25,82355 | 25,92341 | 25,91351 | 25,90931 |
| + | sp\|Q99714\|HCD2_HUMAN;tr\|Q5H928\|Q5H928_HUMAN | sp\|Q99714\|H | 55,523 | 119800000 | 29 | 7 | 0,01618167 | -0,43927574 | 23,15741 | 23,21576 | 23,06149 | 23,22187 | 22,69341 | 23,03066 | 22,41093 | 22,76444 |
| + | sp\|Q04917\|1433F_HUMAN;tr\|A2IDB2\|A2IDB2_HUMAN | sp\|Q04917\|1 | 25,399 | 40595000 | 11 | 5 | 0,0169469 | -0,68645239 | 21,24853 | 22,01981 | 21,80243 | 21,85776 | 21,24336 | 21,20341 | 21,04547 | 20,69047 |
| + | sp\|P41250\|SYG_HUMAN;tr\|H7C443\|H7C443_HUMAN | sp\|P41250\|S | 38,81 | 132740000 | 27 | 9 | 0,01698833 | -0,32668781 | 23,08691 | 23,19625 | 23,25451 | 23,32882 | 22,92767 | 23,00345 | 22,63739 | 22,99124 |
| | tr\|D6RHZ5\|D6RHZ5_HUMAN;tr\|H7BXG7\|H7BXG7_HUMAN;sp\|O94979\|SC31A_HUMAN;tr\|D6 | tr\|D6RHZ5\|D | 3,4856 | 19865000 | 6 | 3 | 0,01731335 | -0,2381986 | NaN | 20,70117 | 20,61522 | 20,69319 | NaN | 20,3625 | 20,39373 | 20,53877 |
| + | sp\|P05455\|LA_HUMAN;tr\|E7ERC4\|E7ERC4_HUMAN;tr\|E9PGX9\|E9PGX9_HUMAN;tr\|E9PFL9\|E | sp\|P05455\|L | 64,152 | 129160000 | 37 | 8 | 0,01750869 | -0,55094337 | 23,24013 | 23,0558 | 23,51713 | 23,47023 | 22,68368 | 23,09325 | 22,83505 | 22,46755 |
| + | sp\|P10155\|RO60_HUMAN;tr\|H0Y9N5\|H0Y9N5_HUMAN;tr\|G5E9R9\|G5E9R9_HUMAN | sp\|P10155\|R | 9,4226 | 64285000 | 9 | 4 | 0,01756114 | 1,07726177 | 20,9775 | 21,04793 | 20,96261 | 21,15283 | 21,8081 | 22,83798 | NaN | 21,69136 |
| + | sp\|O75937\|DNJC8_HUMAN;tr\|S4R3J5\|S4R3J5_HUMAN | sp\|O75937\|D | 9,4041 | 63035000 | 11 | 4 | 0,01769836 | -0,4986736 | 22,37247 | 22,13734 | 22,30025 | 22,46643 | NaN | 21,78676 | 21,59587 | 22,07871 |
| + | sp\|Q01082\|SPTB2_HUMAN;tr\|A0A087WUZ3\|A0A087WUZ3_HUMAN;tr\|F8W6C1\|F8W6C1_HU | sp\|Q01082\|S | 10,086 | 50217000 | 18 | 8 | 0,01811352 | -0,64821768 | 22,33759 | 22,22573 | 22,1928 | 21,74288 | 21,34316 | 21,15047 | 21,5475 | 21,86499 |
| + | tr\|M0QXF7\|M0QXF7_HUMAN;sp\|Q969H8\|MYDGF_HUMAN;tr\|M0QYN0\|M0QYN0_HUMAN | tr\|M0QXF7\|M | 2,9626 | 35875000 | 4 | 2 | 0,01836792 | -0,70679776 | 21,37814 | 21,59108 | 21,80463 | 22,17231 | 21,1283 | 20,94564 | NaN | 21,01529 |
| + | sp\|Q9H3N1\|TMX1_HUMAN;tr\|G3V448\|G3V448_HUMAN | sp\|Q9H3N1\|T | 33,73 | 68474000 | 9 | 6 | 0,01843914 | -0,54205608 | 22,79009 | 22,59823 | 22,13624 | 22,58534 | 21,78009 | 21,95731 | 21,95677 | 22,24751 |
| + | sp\|P60900\|PSA6_HUMAN;tr\|G3V5Z7\|G3V5Z7_HUMAN;tr\|G3V295\|G3V295_HUMAN;tr\|G3V3 | sp\|P60900\|P | 53,591 | 133050000 | 23 | 6 | 0,0185487 | -0,42355108 | 23,41532 | 23,28503 | 23,39807 | 23,46139 | 23,17821 | 23,06199 | 22,59837 | 23,02704 |
| + | sp\|Q53GQ0\|DHB12_HUMAN;tr\|E9PI21\|E9PI21_HUMAN | sp\|Q53GQ0\|I | 25,339 | 156220000 | 19 | 7 | 0,01860571 | 0,90383768 | 22,25525 | 23,01803 | 22,66171 | 22,29387 | 23,70314 | 23,07287 | 23,95465 | 23,11356 |
| + | tr\|F5GYN4\|F5GYN4_HUMAN;sp\|Q96FW1\|OTUB1_HUMAN;tr\|J3KR44\|J3KR44_HUMAN;tr\|F5H | tr\|F5GYN4\|F | 18,629 | 53110000 | 20 | 5 | 0,01891751 | -0,60546398 | 21,76393 | 22,28133 | 22,13 | 22,0902 | 21,42281 | 21,76709 | 21,0442 | 21,6095 |
| + | sp\|P48047\|ATPO_HUMAN;tr\|H7C0C1\|H7C0C1_HUMAN;tr\|H7C086\|H7C086_HUMAN;tr\|H7C0 | sp\|P48047\|A | 8,1415 | 46092000 | 14 | 4 | 0,01900868 | -0,33362913 | 21,778 | 21,48949 | 21,77896 | 21,51567 | 21,41132 | 21,43446 | 21,2165 | 21,16533 |
| + | sp\|P30050\|RL12_HUMAN | sp\|P30050\|R | 40,751 | 384750000 | 35 | 6 | 0,01903145 | -0,30392933 | 24,74929 | 24,78042 | 25,00967 | 24,58228 | 24,43065 | 24,57045 | 24,50045 | 24,40438 |
| + | sp\|P00505\|AATM_HUMAN | sp\|P00505\|A | 56,656 | 162360000 | 18 | 7 | 0,0190778 | -0,68390703 | 23,17668 | 23,88138 | 23,83287 | 23,832 | 23,02989 | 23,09981 | 22,61755 | 23,24006 |
| + | sp\|P61086\|UBE2K_HUMAN;tr\|D6RDM7\|D6RDM7_HUMAN | sp\|P61086\|U | 11,718 | 50281000 | 14 | 4 | 0,01931677 | -0,56931543 | 21,67681 | 21,85018 | 22,28311 | 22,24867 | 21,35499 | 21,71733 | 21,45558 | 21,25361 |

| | | | | | | | | | | | | | | | | |
|---|---|---|---|---|---|---|---|---|---|---|---|---|---|---|---|---|
| + | sp\|Q07666\|KHDR1_HUMAN | sp\|Q07666\|K | 59,74 | 113520000 | 16 | 4 | 0,01932657 | -0,46714401 | 23,29586 | 22,72823 | 23,26428 | 22,97094 | 22,47778 | 22,53205 | 22,62158 | 22,75931 |
| + | sp\|Q53H96\|P5CR3_HUMAN;tr\|A0A0A0MQS1\|A0A0A0MQS1_HUMAN;tr\|F8WEI0\|F8WEI0_HUI | sp\|Q53H96\|F | 3,6983 | 7054600 | 2 | 2 | 0,01943331 | 0,79138438 | 18,51542 | NaN | 18,37105 | 18,43718 | NaN | 19,61829 | 18,91853 | 19,16098 |
| + | sp\|Q96CS3\|FAF2_HUMAN | sp\|Q96CS3\|F | 37,742 | 23905000 | 8 | 5 | 0,01951841 | 0,62145185 | 20,19292 | 19,88408 | 20,10303 | 20,00977 | 20,794 | 21,00717 | 20,73198 | 20,14246 |
| + | tr\|H3BTA2\|H3BTA2_HUMAN;sp\|P60510\|PP4C_HUMAN;tr\|H3BV22\|H3BV22_HUMAN;tr\|I3L4X | tr\|H3BTA2\|H | 9,2841 | 18504000 | 6 | 2 | 0,01952442 | 0,5677096 | NaN | 19,95205 | 19,96846 | 20,25396 | 20,68723 | 20,78548 | 20,40489 | NaN |
| + | sp\|P40925\|MDHC_HUMAN;tr\|B9A041\|B9A041_HUMAN;tr\|B8ZZ51\|B8ZZ51_HUMAN;tr\|C9JF7 | sp\|P40925\|M | 24,299 | 122880000 | 18 | 5 | 0,01964679 | -1,06502453 | 23,29446 | 23,51953 | 23,60162 | 23,7166 | NaN | 21,97232 | 22,2764 | 23,15536 |
| + | sp\|Q14166\|TTL12_HUMAN | sp\|Q14166\|T | 5,9564 | 37099000 | 8 | 6 | 0,01987841 | -0,47847271 | 21,33276 | 21,82427 | 21,44673 | 21,45171 | 21,1266 | 21,18659 | 20,71515 | 21,11324 |
| + | sp\|P61313\|RL15_HUMAN;tr\|E7EQV9\|E7EQV9_HUMAN;tr\|E7EX53\|E7EX53_HUMAN;tr\|E7ENU | sp\|P61313\|R | 148,12 | 437700000 | 24 | 7 | 0,02011304 | -0,70598507 | 25,39002 | 25,53157 | 25,17695 | 24,74396 | 24,22133 | 24,43714 | 24,91118 | 24,44891 |
| + | sp\|P19105\|ML12A_HUMAN;sp\|O14950\|ML12B_HUMAN;tr\|J3QRS3\|J3QRS3_HUMAN;tr\|J3KTJ | sp\|P19105\|M | 8,0677 | 21446000 | 7 | 4 | 0,02022174 | -0,50033855 | 20,73115 | 20,43553 | 20,90866 | 20,908 | 20,55684 | 20,107 | 20,27196 | 20,0462 |
| + | tr\|E9PIA8\|E9PIA8_HUMAN;sp\|P50897\|PPT1_HUMAN;tr\|E9PSE5\|E9PSE5_HUMAN;tr\|Q5T0S4\| | tr\|E9PIA8\|E9 | 3,6415 | 20623000 | 3 | 2 | 0,02030718 | 0,99368858 | 19,59992 | 19,61367 | 19,42614 | 19,79454 | 21,51176 | 20,29961 | 20,43675 | 20,16091 |
| + | sp\|P31946\|1433B_HUMAN;tr\|Q4VY20\|Q4VY20_HUMAN;tr\|A0A0J9YWZ2\|A0A0J9YWZ2_HUMA | sp\|P31946\|1 | 139,49 | 102330000 | 22 | 4 | 0,02079415 | -0,59059286 | 22,8786 | 22,75778 | 23,45526 | 23,01178 | 22,44208 | 22,75022 | 22,23541 | 22,31335 |
| + | sp\|Q8NI60\|ADCK3_HUMAN | sp\|Q8NI60\|A | 2,9443 | 7581900 | 4 | 3 | 0,02089976 | 0,89383094 | 18,54898 | NaN | 18,76526 | 17,89936 | 19,18015 | 19,69554 | 19,19556 | 19,1222 |
| + | sp\|Q13428\|TCOF_HUMAN;tr\|J3KQ96\|J3KQ96_HUMAN;tr\|E7ETY2\|E7ETY2_HUMAN;tr\|H0YA9 | sp\|Q13428\|T | 9,8033 | 44300000 | 12 | 6 | 0,02093694 | -0,56597328 | 21,62015 | 22,10569 | 21,94314 | 21,70299 | 20,91452 | 21,17475 | 21,51648 | 21,50232 |
| | sp\|P40938\|RFC3_HUMAN;tr\|A0A087X270\|A0A087X270_HUMAN | sp\|P40938\|R | 8,036 | 14228000 | 9 | 4 | 0,02183168 | 0,21730566 | 19,79877 | 19,6686 | 19,64324 | 19,67071 | 20,01876 | 19,8608 | 20,00758 | 19,7634 |
| + | sp\|P60174\|TPIS_HUMAN;tr\|U3KPZ0\|U3KPZ0_HUMAN;tr\|U3KQF3\|U3KQF3_HUMAN;tr\|U3KPS | sp\|P60174\|T | 238,77 | 576120000 | 79 | 13 | 0,0220559 | -0,62715721 | 25,44436 | 25,47417 | 25,55798 | 25,7304 | 24,95403 | 25,09297 | 24,3763 | 25,27498 |
| + | sp\|P09211\|GSTP1_HUMAN;tr\|A8MX94\|A8MX94_HUMAN;tr\|A0A087X2E9\|A0A087X2E9_HUM | sp\|P09211\|G | 106,79 | 146430000 | 11 | 6 | 0,02207613 | -0,56889343 | 23,29754 | 23,28616 | 23,99757 | 23,60093 | 22,88861 | 23,2123 | 22,8642 | 22,94152 |
| + | sp\|Q9NTK5\|OLA1_HUMAN;tr\|J3KQ32\|J3KQ32_HUMAN;tr\|C9JTK6\|C9JTK6_HUMAN | sp\|Q9NTK5\|( | 4,3863 | 41770000 | 5 | 4 | 0,02239891 | -1,29992104 | NaN | 21,48267 | 22,5417 | 21,6737 | 20,70033 | 20,79614 | 20,30184 | NaN |
| + | sp\|O00299\|CLIC1_HUMAN | sp\|O00299\|C | 52,508 | 117500000 | 29 | 9 | 0,02277999 | -0,56982279 | 22,95216 | 23,12089 | 23,10262 | 23,39428 | 22,63068 | 22,74948 | 22,09642 | 22,81408 |
| + | tr\|E9PLL6\|E9PLL6_HUMAN;sp\|P46776\|RL27A_HUMAN;tr\|E9PJD9\|E9PJD9_HUMAN;tr\|E9PLX7 | tr\|E9PLL6\|E9 | 20,372 | 265070000 | 13 | 3 | 0,02278772 | -0,52694337 | 24,42892 | 24,81939 | 24,55315 | 24,2281 | 24,07284 | 24,05556 | 23,81294 | NaN |
| + | tr\|F8W7C6\|F8W7C6_HUMAN;tr\|A0A087WV22\|A0A087WV22_HUMAN;tr\|X1WI28\|X1WI28_HU | tr\|F8W7C6\|F | 19,698 | 264680000 | 22 | 5 | 0,0229246 | -0,67729092 | 24,76243 | 24,37425 | 24,89004 | 23,97229 | 23,71156 | 23,65862 | 24,01026 | 23,9094 |
| + | sp\|P43490\|NAMPT_HUMAN;tr\|A0A0C4DFS8\|A0A0C4DFS8_HUMAN | sp\|P43490\|N | 6,1468 | 32838000 | 12 | 5 | 0,02325624 | -0,44271469 | 21,154 | 21,36922 | 21,55791 | 21,76276 | 21,1432 | 20,91145 | 20,89349 | 21,1249 |
| + | sp\|Q15008\|PSMD6_HUMAN;tr\|H7C531\|H7C531_HUMAN;tr\|C9J7B7\|C9J7B7_HUMAN;tr\|C9J0 | sp\|Q15008\|P | 21,132 | 44770000 | 17 | 7 | 0,0234856 | -0,34590928 | 21,72713 | 22,0126 | 21,94831 | 21,71791 | 21,37968 | 21,62574 | 21,51133 | NaN |
| + | sp\|P36578\|RL4_HUMAN;tr\|H3BM89\|H3BM89_HUMAN;tr\|H3BTP7\|H3BTP7_HUMAN;tr\|H3BU | sp\|P36578\|R | 28,989 | 400950000 | 25 | 6 | 0,02357385 | -0,46175718 | 24,91365 | 24,95978 | 25,21608 | 24,80055 | 24,30327 | 24,38726 | 24,87122 | 24,4813 |
| + | sp\|Q04637\|IF4G1_HUMAN;tr\|E7EX73\|E7EX73_HUMAN;tr\|E9PGM1\|E9PGM1_HUMAN;tr\|E7E | sp\|Q04637\|I | 123,35 | 208480000 | 30 | 16 | 0,02366756 | -0,49980307 | 24,23535 | 24,12123 | 24,17843 | 23,73603 | 23,31147 | 23,49826 | 23,56744 | 23,89467 |
| + | tr\|H0Y2W2\|H0Y2W2_HUMAN;sp\|Q9NVI7\|ATD3A_HUMAN;tr\|Q5SV16\|Q5SV16_HUMAN | tr\|H0Y2W2\|H | 29,313 | 93794000 | 15 | 1 | 0,02445898 | 1,08218336 | 21,46643 | 21,67642 | 21,08284 | 22,74917 | 22,92573 | 22,94324 | 22,70824 | 22,72638 |
| + | tr\|A0A0A0MSQ0\|A0A0A0MSQ0_HUMAN;sp\|P13797\|PLST_HUMAN;sp\|P13796\|PLSL_HUMAN; | tr\|A0A0A0MS | 37,702 | 191170000 | 27 | 9 | 0,02471147 | -0,57372379 | 23,43314 | 24,14686 | 23,97334 | 24,02496 | 23,22202 | 23,56848 | 23,06819 | 23,4247 |
| + | sp\|P32119\|PRDX2_HUMAN;tr\|A6NIW5\|A6NIW5_HUMAN | sp\|P32119\|P | 9,433 | 163500000 | 14 | 3 | 0,02483824 | -0,41559601 | 23,49863 | 23,84995 | 23,62979 | 23,70535 | 23,48493 | 23,42803 | 22,998 | 23,11038 |
| + | tr\|J3KT10\|J3KT10_HUMAN;sp\|Q9BW27\|NUP85_HUMAN;tr\|J3KSH3\|J3KSH3_HUMAN;tr\|J3KRC | tr\|J3KT10\|J3 | 7,0456 | 29948000 | 7 | 6 | 0,02500511 | 1,49535418 | 19,90046 | 19,96353 | 20,01386 | 19,0044 | 20,16066 | 21,43781 | 20,98273 | 22,28246 |
| + | sp\|P52907\|CAZA1_HUMAN;tr\|C9JUG7\|C9JUG7_HUMAN;tr\|F8W9N7\|F8W9N7_HUMAN;tr\|A0 | sp\|P52907\|C | 13,75 | 56916000 | 20 | 6 | 0,02537738 | -0,28595638 | 21,824 | 21,88342 | 22,16628 | 22,059 | 21,61063 | 21,81386 | 21,5925 | 21,77189 |
| + | sp\|P30086\|PEBP1_HUMAN | sp\|P30086\|P | 7,9942 | 140690000 | 16 | 5 | 0,02554978 | -0,58832455 | 23,41326 | 23,34096 | 23,2037 | 23,96296 | 23,09875 | 22,98531 | 22,58898 | 22,89454 |
| + | tr\|K7EQA1\|K7EQA1_HUMAN;sp\|O14737\|PDCD5_HUMAN;tr\|K7ESJ4\|K7ESJ4_HUMAN | tr\|K7EQA1\|K | 5,0121 | 21463000 | 8 | 3 | 0,02622635 | -0,39694357 | 20,6759 | 20,56179 | 20,90911 | 20,80262 | 20,2815 | 20,645 | 20,10534 | 20,32981 |
| + | sp\|P53680\|AP2S1_HUMAN;tr\|M0R0N4\|M0R0N4_HUMAN;tr\|M0QYZ2\|M0QYZ2_HUMAN;tr\|M | sp\|P53680\|A | 2,5375 | 41933000 | 3 | 3 | 0,02626943 | 0,77085686 | 21,23175 | 20,89608 | 20,62194 | 20,6943 | 21,74624 | 22,21798 | 21,33041 | 21,23286 |
| | tr\|B1AHB1\|B1AHB1_HUMAN;sp\|P33992\|MCM5_HUMAN;tr\|B1AHB2\|B1AHB2_HUMAN;tr\|B1/ | tr\|B1AHB1\|B | 22,119 | 53991000 | 15 | 5 | 0,02741495 | -0,25472212 | 22,13824 | 22,06494 | 21,7634 | 22,07051 | 21,76652 | 21,67857 | 21,76624 | 21,80688 |
| + | sp\|P55060\|XPO2_HUMAN | sp\|P55060\|X | 116,49 | 340470000 | 67 | 26 | 0,0275836 | -0,45089245 | 24,44563 | 24,82881 | 24,6634 | 24,78985 | 23,97448 | 24,28849 | 24,09273 | 24,56842 |
| + | sp\|Q13263\|TIF1B_HUMAN;tr\|M0R0K9\|M0R0K9_HUMAN;tr\|M0R3C0\|M0R3C0_HUMAN;tr\|M( | sp\|Q13263\|T | 151,02 | 301080000 | 32 | 16 | 0,02784337 | 0,64439583 | 23,62288 | 23,56267 | 24,23352 | 23,96137 | 24,30452 | 24,2839 | 24,40912 | 24,96049 |
| + | sp\|Q14247\|SRC8_HUMAN;tr\|H0YEV2\|H0YEV2_HUMAN;tr\|H0YCD9\|H0YCD9_HUMAN | sp\|Q14247\|S | 7,7053 | 21512000 | 7 | 4 | 0,02806582 | -0,33043734 | NaN | 20,72983 | 20,7159 | 20,75638 | 20,57598 | 20,39582 | 20,239 | NaN |
| + | sp\|P62753\|RS6_HUMAN;tr\|A2A3R7\|A2A3R7_HUMAN;tr\|A2A3R5\|A2A3R5_HUMAN | sp\|P62753\|R | 14,549 | 179590000 | 14 | 3 | 0,02883162 | -1,09243345 | 23,81029 | 23,94101 | 24,64702 | 23,61964 | 22,60198 | 23,83634 | 22,53424 | 22,67566 |
| + | tr\|K7ENG2\|K7ENG2_HUMAN;sp\|P26368\|U2AF2_HUMAN | tr\|K7ENG2\|K | 21,179 | 43261000 | 15 | 4 | 0,02884216 | -0,44436312 | 21,48704 | 21,61225 | 21,84869 | 21,72034 | 20,83105 | 21,42726 | 21,37066 | 21,2619 |
| + | sp\|P09972\|ALDOC_HUMAN;tr\|K7EKH5\|K7EKH5_HUMAN;tr\|C9J8F3\|C9J8F3_HUMAN;tr\|J3KSV | sp\|P09972\|A | 18,473 | 42890000 | 9 | 2 | 0,02885066 | -0,75108608 | NaN | 21,8385 | 22,18807 | 21,45141 | 20,94871 | 21,24592 | 20,73115 | 21,37385 |
| + | tr\|A0A087WUT6\|A0A087WUT6_HUMAN;sp\|O60841\|IF2P_HUMAN | tr\|A0A087WU | 5,1968 | 26140000 | 7 | 4 | 0,02909065 | -0,60641289 | 20,91765 | 21,06415 | 21,38047 | 20,7997 | 19,93962 | 20,73066 | 20,50911 | 20,55693 |
| + | sp\|Q13162\|PRDX4_HUMAN;tr\|H7C3T4\|H7C3T4_HUMAN;tr\|A6NG45\|A6NG45_HUMAN;tr\|A6I | sp\|Q13162\|P | 29,15 | 260170000 | 28 | 6 | 0,02963216 | -0,40049601 | 24,36306 | 24,18026 | 24,23856 | 24,33108 | 23,96173 | 23,93715 | 23,49228 | 24,11981 |
| + | tr\|A0A087WTP3\|A0A087WTP3_HUMAN;sp\|Q92945\|FUBP2_HUMAN;tr\|M0R0I5\|M0R0I5_HUN | tr\|A0A087WT | 83,796 | 351590000 | 47 | 15 | 0,03082711 | -0,43512201 | 24,72208 | 24,72401 | 24,79074 | 24,97075 | 24,39598 | 24,30862 | 24,03313 | 24,72936 |
| + | sp\|Q16658\|FSCN1_HUMAN;tr\|C9JFC0\|C9JFC0_HUMAN;tr\|A0A0A0MSB2\|A0A0A0MSB2_HUM | sp\|Q16658\|F | 81,022 | 419780000 | 56 | 14 | 0,03142751 | -0,40405607 | 24,80386 | 24,92823 | 24,97976 | 25,07928 | 24,45086 | 24,74319 | 24,20978 | 24,77107 |
| + | sp\|Q15365\|PCBP1_HUMAN;tr\|F8VTZ0\|F8VTZ0_HUMAN;tr\|H3BSP4\|H3BSP4_HUMAN;tr\|C9K0 | sp\|Q15365\|P | 241,45 | 404520000 | 42 | 6 | 0,03234676 | -0,49562502 | 24,9608 | 25,00058 | 24,82512 | 25,27352 | 24,64422 | 24,46675 | 24,12502 | 24,84153 |
| + | tr\|B1AHF3\|B1AHF3_HUMAN;sp\|P00387\|NB5R3_HUMAN | tr\|B1AHF3\|B | 8,5449 | 43656000 | 6 | 2 | 0,03247584 | -0,46938086 | 21,45634 | 21,6868 | 22,09513 | 21,65351 | 21,27333 | 21,4913 | 21,26459 | 20,98503 |
| | sp\|O43242\|PSMD3_HUMAN | sp\|O43242\|P | 35,956 | 62642000 | 18 | 10 | 0,03268939 | -0,167665 | 22,00518 | 22,1379 | 22,23468 | 22,04434 | 21,93506 | 21,91506 | 22,02736 | 21,87395 |
| | sp\|P63208\|SKP1_HUMAN;tr\|E7ERH2\|E7ERH2_HUMAN;tr\|F8W8N3\|F8W8N3_HUMAN;tr\|E5RJ | sp\|P63208\|S | 31,048 | 47289000 | 10 | 3 | 0,03396073 | 0,28589106 | 21,55178 | 21,2921 | 21,24383 | 21,47581 | 21,62796 | 21,72521 | 21,85289 | 21,50101 |
| + | sp\|P14923\|PLAK_HUMAN;tr\|C9JTX4\|C9JTX4_HUMAN;tr\|C9J826\|C9J826_HUMAN;tr\|C9JK18\|( | sp\|P14923\|P | 14,16 | 52100000 | 14 | 7 | 0,03404141 | -1,14036179 | 22,59092 | 23,1437 | 21,48812 | 22,09462 | 21,62819 | 21,02513 | 21,46174 | 20,64086 |
| + | sp\|P49915\|GUAA_HUMAN | sp\|P49915\|G | 30,691 | 108200000 | 28 | 10 | 0,03411056 | -0,34590101 | 22,63761 | 22,98888 | 23,12589 | 23,04737 | 22,56458 | 22,71678 | 22,43334 | 22,70145 |
| + | sp\|P10515\|ODP2_HUMAN;tr\|H0YDD4\|H0YDD4_HUMAN;tr\|E9PEJ4\|E9PEJ4_HUMAN | sp\|P10515\|O | 11,235 | 29519000 | 15 | 4 | 0,03485199 | -0,9895277 | 21,29132 | 21,20174 | 21,59455 | 21,52492 | 20,56895 | 20,86469 | 20,84437 | 19,37641 |
| + | sp\|P31948\|STIP1_HUMAN;tr\|F5H783\|F5H783_HUMAN;tr\|F5GXD8\|F5GXD8_HUMAN;tr\|H0YG | sp\|P31948\|S | 44,63 | 326340000 | 33 | 12 | 0,03519777 | -0,44911575 | 24,21978 | 24,67126 | 24,75129 | 24,92746 | 24,15352 | 24,3927 | 24,0721 | 24,15499 |
| + | sp\|P12956\|XRCC6_HUMAN;tr\|B1AHC9\|B1AHC9_HUMAN | sp\|P12956\|X | 153,96 | 425010000 | 61 | 18 | 0,03540524 | -0,41616392 | 24,80247 | 24,95527 | 25,0542 | 25,16645 | 24,30299 | 24,82813 | 24,78397 | 24,39865 |
| + | sp\|P25786\|PSA1_HUMAN;tr\|F5GX11\|F5GX11_HUMAN;tr\|B4DEV8\|B4DEV8_HUMAN | sp\|P25786\|P | 22,768 | 83272000 | 14 | 6 | 0,0354676 | -0,37268257 | 22,25488 | 22,64665 | 22,80136 | 22,28122 | 22,11709 | 22,09278 | 22,08528 | 22,19823 |
| | tr\|D6RD66\|D6RD66_HUMAN;sp\|O75083\|WDR1_HUMAN | tr\|D6RD66\|D | 4,5092 | 14281000 | 2 | 3 | 0,03602224 | -0,16207027 | 19,86097 | 19,90718 | 19,89996 | 20,11108 | 19,79357 | 19,73757 | 19,7632 | 19,83657 |
| + | sp\|Q15293\|RCN1_HUMAN;tr\|E9PP27\|E9PP27_HUMAN | sp\|Q15293\|R | 47,289 | 69569000 | 16 | 6 | 0,03606879 | 0,69627762 | 21,37369 | 22,23357 | 21,55211 | 21,52903 | 22,55642 | 22,36842 | 21,87938 | 22,66928 |
| | sp\|Q9HAV4\|XPO5_HUMAN;tr\|H0Y9I3\|H0Y9I3_HUMAN | sp\|Q9HAV4\|) | 4,0811 | 11734000 | 8 | 4 | 0,03632218 | -0,28377295 | 19,7312 | 19,78545 | 19,80708 | 19,81731 | 19,27935 | 19,60708 | 19,38315 | 19,73638 |
| + | sp\|O75369\|FLNB_HUMAN;tr\|E7EN95\|E7EN95_HUMAN | sp\|O75369\|F | 20,611 | 53953000 | 27 | 13 | 0,03633214 | -0,62109041 | 22,20353 | 22,32149 | 22,31005 | 21,90481 | 21,04946 | 21,44081 | 21,73032 | 22,03493 |
| + | sp\|P56545\|CTBP2_HUMAN;tr\|Q5SQP8\|Q5SQP8_HUMAN;tr\|E9PGB1\|E9PGB1_HUMAN;tr\|E7E | sp\|P56545\|C | 11,175 | 11951000 | 8 | 3 | 0,03645986 | -0,47541746 | NaN | 19,80573 | 20,0713 | 19,96084 | NaN | 19,61072 | 19,59623 | 19,20467 |
| | tr\|G5E9R3\|G5E9R3_HUMAN;tr\|M0R0A1\|M0R0A1_HUMAN;tr\|C9J4Z3\|C9J4Z3_HUMAN;tr\|Q6F | tr\|G5E9R3\|G | 2,4585 | 59791000 | 7 | 3 | 0,03664276 | -0,29493141 | NaN | 22,33276 | 22,2265 | 21,96716 | 21,97631 | 21,85878 | 21,91397 | 21,7731 |
| + | sp\|Q9Y266\|NUDC_HUMAN;tr\|A0A0A0MSS4\|A0A0A0MSS4_HUMAN;tr\|A0A0A0MSU9\|A0A0A0 | sp\|Q9Y266\|N | 5,0729 | 99511000 | 8 | 5 | 0,0367068 | -0,47319508 | 22,64454 | 22,77746 | 22,93959 | 23,00012 | 22,15035 | 22,82052 | 22,15694 | 22,34112 |
| | sp\|Q13283\|G3BP1_HUMAN;tr\|E5RIZ6\|E5RIZ6_HUMAN;tr\|E5RH00\|E5RH00_HUMAN;tr\|D6R9 | sp\|Q13283\|G | 69,872 | 139130000 | 28 | 6 | 0,03672985 | -0,26642036 | 23,23488 | 23,11962 | 23,41995 | 23,41687 | 22,88565 | 22,94969 | 23,13215 | 23,15815 |
| + | sp\|P39748\|FEN1_HUMAN;tr\|I3L3E9\|I3L3E9_HUMAN;tr\|F5H1Y3\|F5H1Y3_HUMAN | sp\|P39748\|F | 21,474 | 38008000 | 14 | 4 | 0,03757439 | -0,50291681 | 21,43553 | 22,00388 | 21,63781 | 21,63112 | 21,59751 | 21,14065 | 21,02486 | 20,93366 |
| + | sp\|O75821\|EIF3G_HUMAN;tr\|K7EP16\|K7EP16_HUMAN;tr\|K7ER90\|K7ER90_HUMAN;tr\|K7EL2 | sp\|O75821\|E | 18,907 | 91998000 | 18 | 5 | 0,03766112 | -0,55716324 | 22,33655 | 22,7274 | 22,96932 | 22,82018 | 21,8338 | 22,49962 | 22,35348 | 21,9379 |
| + | sp\|P82979\|SARNP_HUMAN;tr\|F8VZQ9\|F8VZQ9_HUMAN;tr\|H0YHG0\|H0YHG0_HUMAN | sp\|P82979\|S | 7,2802 | 63375000 | 15 | 4 | 0,03804926 | -0,32326365 | 22,42029 | 22,01229 | 22,22535 | 22,37692 | 21,90746 | 21,80306 | 21,86476 | 22,16653 |
| + | sp\|O76031\|CLPX_HUMAN | sp\|O76031\|C | 16,04 | 49262000 | 6 | 7 | 0,03817021 | 0,60101827 | 20,92412 | 21,37729 | 21,31705 | 21,3811 | NaN | 22,01971 | 22,08994 | 21,44309 |
| + | tr\|E7EQB2\|E7EQB2_HUMAN;tr\|E7ER44\|E7ER44_HUMAN;sp\|P02788\|TRFL_HUMAN;tr\|C9JCF5 | tr\|E7EQB2\|E | 5,1511 | 27395000 | 11 | 4 | 0,03884417 | -0,85915327 | 22,14808 | 20,95333 | 20,97155 | 21,53227 | 20,91481 | 20,14358 | 20,45944 | 20,65079 |
| + | sp\|P11177\|ODPB_HUMAN;tr\|C9J634\|C9J634_HUMAN;tr\|F8WF02\|F8WF02_HUMAN | sp\|P11177\|O | 19,204 | 42708000 | 7 | 5 | 0,03892641 | 0,42365885 | 21,11794 | 20,67444 | 21,18646 | 21,37883 | 21,55154 | 21,47338 | 21,65779 | 21,36959 |
| + | sp\|P23588\|IF4B_HUMAN;tr\|E7EX17\|E7EX17_HUMAN;tr\|F8W0K0\|F8W0K0_HUMAN;tr\|F8VRU | sp\|P23588\|I | 13,417 | 59830000 | 10 | 3 | 0,04020749 | -0,43703461 | 22,29732 | 22,37132 | 22,2596 | 22,2676 | NaN | 21,98458 | 21,4958 | 22,1054 |
| + | tr\|A0A140T936\|A0A140T936_HUMAN;sp\|P26640\|SYVC_HUMAN;tr\|A0A140T954\|A0A140T95 | tr\|A0A140T9 | 38,311 | 63452000 | 24 | 11 | 0,04080661 | -0,46546936 | 22,25716 | 22,56225 | 22,00624 | 22,46827 | 22,23681 | 21,71255 | 21,66887 | 21,81382 |
| + | sp\|Q96PK6\|RBM14_HUMAN;tr\|A0A0A6YYI9\|A0A0A6YYI9_HUMAN;tr\|F8WDX3\|F8WDX3_HUN | sp\|Q96PK6\|R | 18,241 | 82957000 | 18 | 9 | 0,04081104 | 0,3901763 | 22,23515 | 22,13019 | 22,24348 | 22,17871 | 22,67373 | 22,14995 | 22,72752 | 22,79703 |

<table>
<tr><td>+</td><td>sp|P62081|RS7_HUMAN;tr|B5MCP9|B5MCP9_HUMAN</td><td>sp|P62081|R</td><td>24,424</td><td>251770000</td><td>21</td><td>5</td><td>0,04290173</td><td>-0,56008005</td><td>24,40646</td><td>24,30452</td><td>24,55713</td><td>24,08716</td><td>23,38851</td><td>23,95376</td><td>24,24096</td><td>23,53172</td></tr>
<tr><td>+</td><td>tr|A0A0C4DGI3|A0A0C4DGI3_HUMAN;tr|B4DJV2|B4DJV2_HUMAN;sp|O75390|CISY_HUMAN;</td><td>tr|A0A0C4DG</td><td>72,532</td><td>444870000</td><td>41</td><td>9</td><td>0,04298975</td><td>-0,46163034</td><td>25,05846</td><td>25,09442</td><td>25,10322</td><td>25,30414</td><td>24,81142</td><td>24,67487</td><td>24,20903</td><td>25,01839</td></tr>
<tr><td>+</td><td>sp|P26641|EF1G_HUMAN</td><td>sp|P26641|E</td><td>119,63</td><td>572530000</td><td>51</td><td>10</td><td>0,04329094</td><td>-0,36885309</td><td>25,34194</td><td>25,43435</td><td>25,57888</td><td>25,50812</td><td>25,19551</td><td>25,04143</td><td>24,75471</td><td>25,39624</td></tr>
<tr><td></td><td>sp|P20618|PSB1_HUMAN</td><td>sp|P20618|P</td><td>14,238</td><td>137110000</td><td>13</td><td>7</td><td>0,0438934</td><td>-0,31348419</td><td>23,23498</td><td>23,17811</td><td>23,22582</td><td>23,43797</td><td>23,17936</td><td>23,0412</td><td>22,6644</td><td>22,93799</td></tr>
<tr><td>+</td><td>tr|A0A087WZH7|A0A087WZH7_HUMAN;sp|P29966|MARCS_HUMAN</td><td>tr|A0A087W</td><td>216,74</td><td>173920000</td><td>34</td><td>6</td><td>0,0439693</td><td>-0,68944693</td><td>23,6147</td><td>23,70398</td><td>23,85823</td><td>23,52097</td><td>22,80392</td><td>23,46799</td><td>23,33879</td><td>22,3294</td></tr>
<tr><td>+</td><td>sp|Q9BQ39|DDX50_HUMAN;tr|A0A087WVC1|A0A087WVC1_HUMAN</td><td>sp|Q9BQ39|</td><td>5,8621</td><td>37400000</td><td>15</td><td>5</td><td>0,04404501</td><td>-0,34939623</td><td>21,77688</td><td>21,38906</td><td>21,4314</td><td>21,34262</td><td>21,10591</td><td>20,88398</td><td>21,22305</td><td>21,32943</td></tr>
<tr><td>+</td><td>tr|H0Y8E6|H0Y8E6_HUMAN;sp|P49736|MCM2_HUMAN</td><td>tr|H0Y8E6|H</td><td>32,421</td><td>104890000</td><td>38</td><td>12</td><td>0,04448233</td><td>-0,47129011</td><td>23,26728</td><td>23,16665</td><td>23,01243</td><td>22,79622</td><td>22,41589</td><td>22,49833</td><td>22,39312</td><td>23,05008</td></tr>
<tr><td>+</td><td>sp|P07954|FUMH_HUMAN</td><td>sp|P07954|F</td><td>17,141</td><td>37825000</td><td>12</td><td>5</td><td>0,0446785</td><td>-0,4368</td><td>21,26562</td><td>21,58045</td><td>21,70683</td><td>21,69859</td><td>21,01679</td><td>21,14345</td><td>20,84537</td><td>21,49868</td></tr>
<tr><td></td><td>sp|P02786|TFR1_HUMAN;tr|G3V0E5|G3V0E5_HUMAN;tr|H7C3V5|H7C3V5_HUMAN;tr|F8WB</td><td>sp|P02786|T</td><td>10,259</td><td>44524000</td><td>10</td><td>7</td><td>0,04512877</td><td>-0,24558163</td><td>21,60639</td><td>21,75422</td><td>21,59291</td><td>21,60883</td><td>21,18973</td><td>21,3009</td><td>21,56137</td><td>21,52803</td></tr>
<tr><td></td><td>sp|Q13155|AIMP2_HUMAN;tr|A8MU58|A8MU58_HUMAN;tr|F8W950|F8W950_HUMAN</td><td>sp|Q13155|A</td><td>6,8678</td><td>14442000</td><td>5</td><td>3</td><td>0,0477264</td><td>-0,25801086</td><td>19,94235</td><td>19,67883</td><td>20,05973</td><td>20,05245</td><td>19,54772</td><td>19,80747</td><td>19,69518</td><td>19,65094</td></tr>
<tr><td>+</td><td>tr|M0QWZ7|M0QWZ7_HUMAN;sp|Q9NP81|SYSM_HUMAN;tr|B4DJM9|B4DJM9_HUMAN;tr|</td><td>tr|M0QWZ7|</td><td>4,235</td><td>25979000</td><td>8</td><td>4</td><td>0,04783429</td><td>0,57996368</td><td>20,31717</td><td>20,35165</td><td>20,32301</td><td>20,39194</td><td>21,48513</td><td>21,08128</td><td>20,74296</td><td>20,39425</td></tr>
<tr><td>+</td><td>sp|Q02790|FKBP4_HUMAN;tr|H0YFG2|H0YFG2_HUMAN;tr|F5H1U3|F5H1U3_HUMAN</td><td>sp|Q02790|F</td><td>120,83</td><td>261940000</td><td>35</td><td>14</td><td>0,0481356</td><td>-0,45043707</td><td>24,08205</td><td>24,36486</td><td>24,53996</td><td>24,61689</td><td>23,9486</td><td>24,08529</td><td>23,56663</td><td>24,2015</td></tr>
<tr><td>+</td><td>sp|P46821|MAP1B_HUMAN;tr|D6RA32|D6RA32_HUMAN;tr|D6RCL2|D6RCL2_HUMAN;tr|D6R</td><td>sp|P46821|M</td><td>5,6041</td><td>10276000</td><td>5</td><td>4</td><td>0,04814222</td><td>0,63013744</td><td>18,93759</td><td>18,88369</td><td>18,71102</td><td>18,88313</td><td>18,96101</td><td>19,70737</td><td>20,06999</td><td>19,19761</td></tr>
<tr><td>+</td><td>tr|G3V203|G3V203_HUMAN;sp|Q07020|RL18_HUMAN;tr|J3QQ67|J3QQ67_HUMAN;tr|H0YH</td><td>tr|G3V203|G</td><td>183,54</td><td>341810000</td><td>21</td><td>6</td><td>0,04844183</td><td>-0,66135979</td><td>25,17919</td><td>25,29302</td><td>25,0599</td><td>24,4279</td><td>24,85191</td><td>24,03733</td><td>24,32697</td><td>24,09836</td></tr>
<tr><td>+</td><td>sp|P23526|SAHH_HUMAN</td><td>sp|P23526|S</td><td>55,992</td><td>389680000</td><td>28</td><td>7</td><td>0,04898085</td><td>-0,357131</td><td>24,55941</td><td>24,83788</td><td>24,88721</td><td>24,82211</td><td>24,26549</td><td>24,76981</td><td>24,21756</td><td>24,42522</td></tr>
<tr><td>+</td><td>sp|P78347|GTF2I_HUMAN</td><td>sp|P78347|G</td><td>79,061</td><td>38222000</td><td>18</td><td>6</td><td>0,04955229</td><td>-0,67649889</td><td>22,22506</td><td>20,90272</td><td>21,56263</td><td>21,48488</td><td>20,90095</td><td>20,80349</td><td>20,76474</td><td>21,0001</td></tr>
<tr><td></td><td>sp|P06576|ATPB_HUMAN;tr|H0YH81|H0YH81_HUMAN;tr|F8W079|F8W079_HUMAN;tr|F8W</td><td>sp|P06576|A</td><td>323,31</td><td>600530000</td><td>74</td><td>15</td><td>0,04986613</td><td>0,17950487</td><td>25,09362</td><td>25,04293</td><td>25,14663</td><td>25,32306</td><td>25,31437</td><td>25,43259</td><td>25,34201</td><td>25,23528</td></tr>
<tr><td>+</td><td>sp|P43686|PRS6B_HUMAN</td><td>sp|P43686|P</td><td>26,722</td><td>35922000</td><td>6</td><td>4</td><td>0,04993586</td><td>0,5918808</td><td>20,64878</td><td>20,46981</td><td>20,87252</td><td>20,85392</td><td>20,80112</td><td>21,83248</td><td>21,11566</td><td>21,46329</td></tr>
<tr><td>+</td><td>sp|P61081|UBC12_HUMAN;tr|M0QX69|M0QX69_HUMAN;tr|M0QYI6|M0QYI6_HUMAN</td><td>sp|P61081|U</td><td>16,362</td><td>45684000</td><td>17</td><td>5</td><td>0,05052219</td><td>-0,67232291</td><td>NaN</td><td>22,00038</td><td>21,8723</td><td>22,09139</td><td>21,51581</td><td>21,41225</td><td>20,68123</td><td>21,65351</td></tr>
<tr><td>+</td><td>sp|Q9Y3F4|STRAP_HUMAN;tr|H0YH33|H0YH33_HUMAN</td><td>sp|Q9Y3F4|S</td><td>18,552</td><td>99513000</td><td>16</td><td>7</td><td>0,05080591</td><td>-0,65475035</td><td>22,61648</td><td>22,98163</td><td>23,10181</td><td>23,09692</td><td>22,59181</td><td>22,68389</td><td>22,29367</td><td>21,60847</td></tr>
<tr><td>+</td><td>tr|A0A087WVM4|A0A087WVM4_HUMAN;sp|Q6UB35|C1TM_HUMAN;tr|B7ZM99|B7ZM99_H</td><td>tr|A0A087W</td><td>26,457</td><td>223840000</td><td>35</td><td>14</td><td>0,05141246</td><td>0,42850447</td><td>23,38903</td><td>23,3232</td><td>23,61088</td><td>23,53552</td><td>23,56813</td><td>23,71272</td><td>23,97832</td><td>24,31347</td></tr>
<tr><td>+</td><td>sp|Q99536|VAT1_HUMAN;tr|K7ERT7|K7ERT7_HUMAN;tr|K7EJM4|K7EJM4_HUMAN;tr|K7ER8</td><td>sp|Q99536|V</td><td>30,726</td><td>21659000</td><td>9</td><td>5</td><td>0,05162074</td><td>-0,60836935</td><td>20,43685</td><td>20,56653</td><td>21,24144</td><td>20,77435</td><td>20,09352</td><td>20,62827</td><td>20,09854</td><td>19,76536</td></tr>
<tr><td>+</td><td>sp|P23284|PPIB_HUMAN</td><td>sp|P23284|P</td><td>17,803</td><td>65828000</td><td>8</td><td>4</td><td>0,05185497</td><td>-0,94159889</td><td>22,19736</td><td>21,71749</td><td>23,13189</td><td>22,98612</td><td>21,46707</td><td>21,58261</td><td>21,12692</td><td>22,08987</td></tr>
<tr><td>+</td><td>sp|Q96QK1|VPS35_HUMAN</td><td>sp|Q96QK1|</td><td>15,746</td><td>32901000</td><td>14</td><td>7</td><td>0,0525898</td><td>-0,395576</td><td>21,45046</td><td>21,31396</td><td>21,18144</td><td>21,34044</td><td>20,88733</td><td>20,73595</td><td>20,70649</td><td>21,37422</td></tr>
<tr><td>+</td><td>tr|A0A0A0MSI0|A0A0A0MSI0_HUMAN;sp|Q06830|PRDX1_HUMAN;tr|A0A0A0MRQ5|A0A0A0</td><td>tr|A0A0A0M</td><td>25,947</td><td>2886700000</td><td>36</td><td>7</td><td>0,05292747</td><td>-0,40839481</td><td>27,48516</td><td>27,82705</td><td>27,66099</td><td>27,91028</td><td>27,52103</td><td>27,56935</td><td>26,97509</td><td>27,18442</td></tr>
<tr><td>+</td><td>sp|Q96P70|IPO9_HUMAN</td><td>sp|Q96P70|I</td><td>80,507</td><td>21735000</td><td>10</td><td>4</td><td>0,05367894</td><td>0,68706703</td><td>20,3469</td><td>19,8142</td><td>19,92647</td><td>20,53934</td><td>20,72758</td><td>20,56076</td><td>20,563</td><td>21,52382</td></tr>
<tr><td></td><td>sp|P38919|IF4A3_HUMAN;tr|I3L3H2|I3L3H2_HUMAN</td><td>sp|P38919|I</td><td>15,213</td><td>45714000</td><td>11</td><td>5</td><td>0,05374339</td><td>0,18948603</td><td>21,37719</td><td>21,51735</td><td>21,37183</td><td>21,32092</td><td>21,4402</td><td>21,73909</td><td>21,51523</td><td>21,65071</td></tr>
<tr><td>+</td><td>sp|P18077|RL35A_HUMAN;tr|F8WBS5|F8WBS5_HUMAN;tr|F8WB72|F8WB72_HUMAN;tr|C9</td><td>sp|P18077|R</td><td>76,291</td><td>385120000</td><td>11</td><td>5</td><td>0,05482654</td><td>-0,71045828</td><td>24,95593</td><td>25,59446</td><td>25,11577</td><td>24,48757</td><td>24,49454</td><td>23,76638</td><td>24,63378</td><td>24,41719</td></tr>
<tr><td>+</td><td>sp|Q12788|TBL3_HUMAN;tr|J3KNP2|J3KNP2_HUMAN;tr|A0A087WYP7|A0A087WYP7_HUMA</td><td>sp|Q12788|T</td><td>5,5018</td><td>9777300</td><td>7</td><td>4</td><td>0,05510277</td><td>0,47468042</td><td>18,9061</td><td>18,89454</td><td>18,91713</td><td>19,06834</td><td>18,9502</td><td>19,50996</td><td>19,8919</td><td>19,33276</td></tr>
<tr><td>+</td><td>tr|M0R026|M0R026_HUMAN;sp|A1L0T0|ILVBL_HUMAN;tr|M0R1B5|M0R1B5_HUMAN;tr|E9F</td><td>tr|M0R026|M</td><td>60,854</td><td>31529000</td><td>10</td><td>5</td><td>0,0554821</td><td>1,49037981</td><td>NaN</td><td>19,45932</td><td>19,73319</td><td>19,66706</td><td>19,78556</td><td>21,65923</td><td>22,08317</td><td>20,91298</td></tr>
<tr><td>+</td><td>sp|P61088|UBE2N_HUMAN;tr|F8VZ29|F8VZ29_HUMAN;tr|F8VSD4|F8VSD4_HUMAN;tr|F8VV</td><td>sp|P61088|U</td><td>12,864</td><td>63229000</td><td>7</td><td>3</td><td>0,05595312</td><td>-0,99633837</td><td>21,6197</td><td>22,09394</td><td>22,88276</td><td>23,24766</td><td>21,50033</td><td>21,94095</td><td>21,46623</td><td>20,9512</td></tr>
<tr><td>+</td><td>sp|Q9BZX2|UCK2_HUMAN;sp|Q9HA47|UCK1_HUMAN</td><td>sp|Q9BZX2|U</td><td>7,5105</td><td>20202000</td><td>9</td><td>5</td><td>0,0577204</td><td>0,38935137</td><td>19,88178</td><td>20,23795</td><td>20,13521</td><td>20,29569</td><td>20,3932</td><td>20,73727</td><td>20,20048</td><td>20,77708</td></tr>
<tr><td>+</td><td>sp|Q9BQA1|MEP50_HUMAN;tr|H0Y711|H0Y711_HUMAN</td><td>sp|Q9BQA1|</td><td>12,859</td><td>31014000</td><td>10</td><td>4</td><td>0,05844179</td><td>0,60659599</td><td>20,32003</td><td>20,37284</td><td>20,77113</td><td>20,45964</td><td>21,39482</td><td>21,5935</td><td>20,74468</td><td>20,61702</td></tr>
<tr><td></td><td>sp|Q13347|EIF3I_HUMAN;tr|Q5TFK1|Q5TFK1_HUMAN</td><td>sp|Q13347|E</td><td>25,429</td><td>101130000</td><td>22</td><td>8</td><td>0,05853696</td><td>-0,35648537</td><td>22,70898</td><td>22,8596</td><td>23,23664</td><td>22,78734</td><td>22,57309</td><td>22,73107</td><td>22,26508</td><td>22,59737</td></tr>
<tr><td></td><td>sp|Q15645|PCH2_HUMAN;tr|H0YAL2|H0YAL2_HUMAN</td><td>sp|Q15645|P</td><td>5,546</td><td>37683000</td><td>7</td><td>4</td><td>0,05930415</td><td>0,25971699</td><td>20,82912</td><td>21,04527</td><td>21,07294</td><td>21,13984</td><td>21,41736</td><td>21,40857</td><td>21,03436</td><td>21,26574</td></tr>
<tr><td>+</td><td>sp|P61604|CH10_HUMAN;tr|B8ZZL8|B8ZZL8_HUMAN;tr|S4R3N1|S4R3N1_HUMAN;tr|B8ZZ54</td><td>sp|P61604|C</td><td>55,388</td><td>237180000</td><td>14</td><td>6</td><td>0,0600002</td><td>0,70047696</td><td>NaN</td><td>23,53445</td><td>23,06886</td><td>23,80193</td><td>24,56477</td><td>24,3678</td><td>23,68603</td><td>24,05697</td></tr>
<tr><td></td><td>sp|Q92499|DDX1_HUMAN;tr|F1T0B3|F1T0B3_HUMAN;tr|A0A087X2G1|A0A087X2G1_HUMA</td><td>sp|Q92499|D</td><td>82,242</td><td>72139000</td><td>33</td><td>10</td><td>0,06011566</td><td>-0,34326839</td><td>22,34101</td><td>22,28382</td><td>22,42322</td><td>22,50758</td><td>21,67719</td><td>22,27125</td><td>22,2569</td><td>21,97722</td></tr>
<tr><td>+</td><td>tr|E7EPB3|E7EPB3_HUMAN;sp|P50914|RL14_HUMAN</td><td>tr|E7EPB3|E</td><td>8,6939</td><td>246370000</td><td>13</td><td>3</td><td>0,06231429</td><td>-0,60150766</td><td>24,85334</td><td>24,06497</td><td>24,61127</td><td>23,81411</td><td>23,65185</td><td>23,5438</td><td>24,04618</td><td>23,69583</td></tr>
<tr><td>+</td><td>tr|F8W1K5|F8W1K5_HUMAN;tr|F8VXJ7|F8VXJ7_HUMAN;sp|Q9Y2B0|CNPY2_HUMAN;tr|F8W</td><td>tr|F8W1K5|F</td><td>22,818</td><td>18762000</td><td>9</td><td>4</td><td>0,06244805</td><td>-0,45456648</td><td>20,19713</td><td>20,79725</td><td>20,89852</td><td>20,64386</td><td>NaN</td><td>20,25327</td><td>20,05748</td><td>20,22811</td></tr>
<tr><td></td><td>sp|P50991|TCPD_HUMAN</td><td>sp|P50991|T</td><td>165,38</td><td>812450000</td><td>66</td><td>20</td><td>0,06310362</td><td>-0,28531075</td><td>25,57764</td><td>25,89384</td><td>25,86689</td><td>26,1541</td><td>25,56506</td><td>25,66947</td><td>25,47822</td><td>25,63848</td></tr>
<tr><td>+</td><td>sp|P61353|RL27_HUMAN;tr|K7ELC7|K7ELC7_HUMAN;tr|K7EQQ9|K7EQQ9_HUMAN;tr|K7ERY</td><td>sp|P61353|R</td><td>24,056</td><td>190470000</td><td>12</td><td>4</td><td>0,0634605</td><td>-0,50155544</td><td>23,74479</td><td>24,03322</td><td>23,82182</td><td>23,39545</td><td>23,56605</td><td>23,40639</td><td>23,26556</td><td>22,75105</td></tr>
<tr><td>+</td><td>sp|P30048|PRDX3_HUMAN</td><td>sp|P30048|P</td><td>14,312</td><td>135750000</td><td>28</td><td>6</td><td>0,06412876</td><td>-0,51013374</td><td>23,18828</td><td>23,26857</td><td>23,67228</td><td>23,89966</td><td>23,0845</td><td>23,20086</td><td>22,55368</td><td>23,14922</td></tr>
<tr><td>+</td><td>sp|P62191|PRS4_HUMAN;tr|G3V4X1|G3V4X1_HUMAN</td><td>sp|P62191|P</td><td>105,45</td><td>88205000</td><td>10</td><td>7</td><td>0,06413009</td><td>0,39917946</td><td>22,20494</td><td>22,50756</td><td>22,30349</td><td>22,36175</td><td>22,25419</td><td>22,85933</td><td>22,89877</td><td>22,96215</td></tr>
<tr><td></td><td>tr|J3QT28|J3QT28_HUMAN;sp|O43684|BUB3_HUMAN;tr|J3QSX4|J3QSX4_HUMAN</td><td>tr|J3QT28|J</td><td>10,966</td><td>57097000</td><td>13</td><td>4</td><td>0,06426207</td><td>-0,27774572</td><td>21,89682</td><td>22,04507</td><td>22,12682</td><td>22,08648</td><td>21,90415</td><td>21,73723</td><td>21,45408</td><td>21,94874</td></tr>
<tr><td>+</td><td>tr|A0A087WZZ5|A0A087WZZ5_HUMAN;sp|Q13435|SF3B2_HUMAN;tr|E9PJ04|E9PJ04_HUMA</td><td>tr|A0A087W</td><td>14,899</td><td>88058000</td><td>16</td><td>6</td><td>0,06436892</td><td>-0,42831182</td><td>22,8226</td><td>22,69515</td><td>22,81423</td><td>22,42278</td><td>21,98423</td><td>22,01764</td><td>22,35251</td><td>22,68714</td></tr>
<tr><td>+</td><td>sp|Q9Y696|CLIC4_HUMAN</td><td>sp|Q9Y696|C</td><td>12,052</td><td>23291000</td><td>6</td><td>3</td><td>0,06455772</td><td>-0,46645641</td><td>20,57339</td><td>20,97218</td><td>20,57967</td><td>21,02709</td><td>20,76182</td><td>20,12149</td><td>20,01713</td><td>20,38606</td></tr>
<tr><td></td><td>sp|P62266|RS23_HUMAN;tr|D6RD47|D6RD47_HUMAN;tr|D6RDJ2|D6RDJ2_HUMAN;tr|D6RIX</td><td>sp|P62266|R</td><td>32,37</td><td>323820000</td><td>24</td><td>7</td><td>0,06463604</td><td>-0,39170218</td><td>24,95323</td><td>24,66855</td><td>24,23411</td><td>24,28263</td><td>24,21971</td><td>24,14383</td><td>24,1536</td><td>24,05457</td></tr>
<tr><td>+</td><td>tr|M0R0P1|M0R0P1_HUMAN;tr|M0R299|M0R299_HUMAN;tr|M0QXL5|M0QXL5_HUMAN;tr|</td><td>tr|M0R0P1|M</td><td>6,8239</td><td>33751000</td><td>4</td><td>3</td><td>0,06505102</td><td>0,75184584</td><td>20,5966</td><td>20,22176</td><td>20,49775</td><td>20,37624</td><td>20,73207</td><td>21,04627</td><td>22,12414</td><td>20,79725</td></tr>
<tr><td></td><td>sp|P52292|IMA1_HUMAN;tr|J3QLL0|J3QLL0_HUMAN;tr|J3KS65|J3KS65_HUMAN</td><td>sp|P52292|I</td><td>70,427</td><td>167110000</td><td>30</td><td>8</td><td>0,06567912</td><td>0,3229413</td><td>23,40315</td><td>23,05755</td><td>23,34394</td><td>23,14886</td><td>23,52013</td><td>23,44063</td><td>23,37822</td><td>23,90628</td></tr>
<tr><td>+</td><td>sp|Q9H0U4|RAB1B_HUMAN;tr|E9PLD0|E9PLD0_HUMAN;tr|A0A087WTI1|A0A087WTI1_HUM</td><td>sp|Q9H0U4|</td><td>183,78</td><td>163580000</td><td>24</td><td>2</td><td>0,06667763</td><td>-0,64152861</td><td>23,27256</td><td>24,14709</td><td>23,29516</td><td>23,42021</td><td>22,86671</td><td>23,43505</td><td>22,78213</td><td>22,48503</td></tr>
<tr><td></td><td>sp|Q9NZI8|IF2B1_HUMAN</td><td>sp|Q9NZI8|I</td><td>88,672</td><td>311700000</td><td>40</td><td>12</td><td>0,06683914</td><td>0,34544182</td><td>24,01699</td><td>23,71796</td><td>24,19648</td><td>23,90408</td><td>24,06102</td><td>24,16875</td><td>24,58469</td><td>24,40282</td></tr>
<tr><td></td><td>sp|P15880|RS2_HUMAN;tr|E9PQD7|E9PQD7_HUMAN;tr|H0YEN5|H0YEN5_HUMAN;tr|E9PM</td><td>sp|P15880|R</td><td>80,599</td><td>791100000</td><td>37</td><td>8</td><td>0,06703049</td><td>-0,29196024</td><td>25,83048</td><td>26,06919</td><td>25,92802</td><td>25,52046</td><td>25,67906</td><td>25,55183</td><td>25,56098</td><td>25,38844</td></tr>
<tr><td></td><td>tr|B1AK87|B1AK87_HUMAN;tr|B1AK88|B1AK88_HUMAN;tr|B1AK85|B1AK85_HUMAN;sp|P47</td><td>tr|B1AK87|B</td><td>4,157</td><td>23261000</td><td>9</td><td>3</td><td>0,06706537</td><td>-0,37556458</td><td>20,78093</td><td>20,89178</td><td>20,62658</td><td>20,81495</td><td>20,77467</td><td>20,53687</td><td>20,05378</td><td>20,24667</td></tr>
<tr><td></td><td>sp|P29401|TKT_HUMAN;tr|A0A0B4J1R6|A0A0B4J1R6_HUMAN;tr|E9PFF2|E9PFF2_HUMAN;tr</td><td>sp|P29401|T</td><td>51,114</td><td>582600000</td><td>37</td><td>15</td><td>0,06731055</td><td>-0,32649946</td><td>25,24104</td><td>25,43317</td><td>25,27402</td><td>25,62642</td><td>24,94855</td><td>25,18835</td><td>24,80582</td><td>25,32591</td></tr>
<tr><td></td><td>sp|Q5VTE0|EF1A3_HUMAN;sp|P68104|EF1A1_HUMAN;tr|A0A087WV01|A0A087WV01_HUM</td><td>sp|Q5VTE0|E</td><td>323,31</td><td>8615500000</td><td>84</td><td>11</td><td>0,06859263</td><td>0,15701294</td><td>28,83744</td><td>29,05938</td><td>28,9493</td><td>29,1159</td><td>29,12196</td><td>29,25169</td><td>29,10262</td><td>29,11379</td></tr>
<tr><td></td><td>sp|P30041|PRDX6_HUMAN</td><td>sp|P30041|P</td><td>242,47</td><td>302420000</td><td>47</td><td>10</td><td>0,06908716</td><td>-0,39642572</td><td>24,2617</td><td>24,45419</td><td>24,501</td><td>24,64312</td><td>24,34001</td><td>24,34422</td><td>23,71712</td><td>23,87296</td></tr>
<tr><td></td><td>tr|E5RHW4|E5RHW4_HUMAN;sp|O94905|ERLN2_HUMAN;tr|E5RJ09|E5RJ09_HUMAN;tr|B0Q</td><td>tr|E5RHW4|E</td><td>8,1707</td><td>46480000</td><td>9</td><td>5</td><td>0,06931958</td><td>0,21535349</td><td>21,47308</td><td>21,30374</td><td>21,49492</td><td>21,51152</td><td>21,91284</td><td>21,54274</td><td>21,60558</td><td>21,58353</td></tr>
<tr><td></td><td>tr|C9J8M3|C9J8M3_HUMAN;tr|C9JZI1|C9JZI1_HUMAN;sp|P35249|RFC4_HUMAN;tr|C9JXZ7|</td><td>tr|C9J8M3|C</td><td>1,7462</td><td>21725000</td><td>6</td><td>2</td><td>0,06967797</td><td>0,2237668</td><td>20,26465</td><td>20,38501</td><td>20,37093</td><td>20,17779</td><td>20,78636</td><td>20,46364</td><td>20,39163</td><td>20,45181</td></tr>
<tr><td>+</td><td>sp|Q9BQ67|GRWD1_HUMAN;tr|M0QX71|M0QX71_HUMAN</td><td>sp|Q9BQ67|</td><td>7,3949</td><td>19100000</td><td>8</td><td>4</td><td>0,06983388</td><td>0,46941884</td><td>20,30508</td><td>NaN</td><td>20,43573</td><td>20,07848</td><td>20,46184</td><td>20,75001</td><td>21,0157</td><td>NaN</td></tr>
<tr><td></td><td>sp|Q86VP6|CAND1_HUMAN;tr|A0A0C4DGH5|A0A0C4DGH5_HUMAN;tr|H0YH27|H0YH27_HU</td><td>sp|Q86VP6|</td><td>26,168</td><td>192440000</td><td>38</td><td>16</td><td>0,06999916</td><td>-0,40903759</td><td>23,99826</td><td>23,74263</td><td>23,75236</td><td>24,01052</td><td>23,21105</td><td>23,32375</td><td>23,36413</td><td>23,96869</td></tr>
<tr><td>+</td><td>sp|O00410|IPO5_HUMAN;tr|H0Y8C6|H0Y8C6_HUMAN;tr|E7ETV3|E7ETV3_HUMAN;tr|E7EQT</td><td>sp|O00410|I</td><td>114,79</td><td>117920000</td><td>33</td><td>11</td><td>0,07102946</td><td>-0,48742247</td><td>23,15643</td><td>23,12127</td><td>23,53457</td><td>23,30076</td><td>22,579</td><td>22,56041</td><td>22,62939</td><td>23,39454</td></tr>
<tr><td></td><td>sp|P46109|CRKL_HUMAN</td><td>sp|P46109|C</td><td>2,9743</td><td>7408000</td><td>4</td><td>2</td><td>0,0711893</td><td>-0,2193354</td><td>NaN</td><td>19,0162</td><td>19,10034</td><td>19,29134</td><td>18,77684</td><td>18,94567</td><td>18,89205</td><td>19,05195</td></tr>
<tr><td>+</td><td>tr|A0A087WX29|A0A087WX29_HUMAN;tr|B1AKP7|B1AKP7_HUMAN;tr|G3V162|G3V162_HU</td><td>tr|A0A087W</td><td>23,477</td><td>52700000</td><td>5</td><td>3</td><td>0,07158843</td><td>0,4991641</td><td>20,79408</td><td>21,40432</td><td>21,35515</td><td>21,83996</td><td>21,61764</td><td>21,96088</td><td>21,93614</td><td>21,87549</td></tr>
<tr><td></td><td>sp|Q8WUM4|PDC6I_HUMAN;tr|F8WDK9|F8WDK9_HUMAN</td><td>sp|Q8WUM4</td><td>40,711</td><td>56211000</td><td>22</td><td>9</td><td>0,07221416</td><td>-0,28700018</td><td>22,0624</td><td>22,12921</td><td>21,95269</td><td>22,09291</td><td>21,73698</td><td>21,83642</td><td>21,45367</td><td>22,06214</td></tr>
<tr><td>+</td><td>sp|P35637|FUS_HUMAN;tr|H3BPE7|H3BPE7_HUMAN;tr|A0A075B7D9|A0A075B7D9_HUMAN</td><td>sp|P35637|F</td><td>69,795</td><td>86463000</td><td>12</td><td>4</td><td>0,07286724</td><td>-0,85631466</td><td>22,69905</td><td>22,7091</td><td>23,13863</td><td>22,89774</td><td>22,4465</td><td>22,54875</td><td>22,12884</td><td>20,89519</td></tr>
<tr><td>+</td><td>sp|P04259|K2C6B_HUMAN</td><td>sp|P04259|K</td><td>163,61</td><td>610840000</td><td>60</td><td>1</td><td>0,0734617</td><td>-1,41838837</td><td>25,6126</td><td>27,11734</td><td>24,89606</td><td>26,09033</td><td>25,36429</td><td>23,6413</td><td>25,24514</td><td>23,79203</td></tr>
<tr><td></td><td>tr|A0A0B4J2E5|A0A0B4J2E5_HUMAN;sp|Q15269|PWP2_HUMAN</td><td>tr|A0A0B4J2E</td><td>5,0086</td><td>9690500</td><td>6</td><td>2</td><td>0,07409676</td><td>0,34461896</td><td>19,01362</td><td>NaN</td><td>19,05502</td><td>19,10352</td><td>19,11319</td><td>19,43011</td><td>19,73047</td><td>19,33425</td></tr>
<tr><td>+</td><td>sp|Q15691|MARE1_HUMAN</td><td>sp|Q15691|N</td><td>15,913</td><td>84595000</td><td>21</td><td>6</td><td>0,07434192</td><td>-0,66049719</td><td>22,0746</td><td>23,02933</td><td>23,08876</td><td>22,92038</td><td>22,31703</td><td>22,09529</td><td>21,58545</td><td>22,47331</td></tr>
<tr><td></td><td>sp|P07814|SYEP_HUMAN;tr|V9GYZ6|V9GYZ6_HUMAN;tr|V9GZ76|V9GZ76_HUMAN</td><td>sp|P07814|S</td><td>134,53</td><td>265510000</td><td>39</td><td>19</td><td>0,07437472</td><td>0,36668205</td><td>24,02293</td><td>23,80499</td><td>24,00624</td><td>23,62187</td><td>24,03984</td><td>23,97954</td><td>24,30682</td><td>24,59656</td></tr>
</table>

| | | | | | | | | | | | | | | | | |
|---|---|---|---|---|---|---|---|---|---|---|---|---|---|---|---|---|
| | sp\|P16403\|H12_HUMAN;sp\|P10412\|H14_HUMAN;sp\|P16402\|H13_HUMAN;sp\|P22492\|H1T_ | sp\|P16403\|H | 71,322 | 1495000000 | 40 | 8 | 0,07475707 | -0,22583675 | 26,6476 | 26,7082 | 26,7571 | 26,7057 | 26,19342 | 26,50253 | 26,67936 | 26,53993 |
| + | sp\|Q15843\|NEDD8_HUMAN;tr\|F8VSA6\|F8VSA6_HUMAN;tr\|E9PS38\|E9PS38_HUMAN;tr\|S4R3( | sp\|Q15843\|N | 1,8077 | 9127100 | 2 | 2 | 0,07509098 | 0,46106593 | NaN | 19,07041 | 19,13786 | 18,67131 | 19,27733 | 19,79308 | 19,14803 | 19,46527 |
| | tr\|B5MCX3\|B5MCX3_HUMAN;sp\|Q15019\|SEPT2_HUMAN;tr\|C9J2Q4\|C9J2Q4_HUMAN;tr\|C9IY | tr\|B5MCX3\|B | 46,892 | 44933000 | 17 | 7 | 0,07518187 | -0,31184149 | 21,62885 | 21,58063 | 21,67189 | 21,7402 | 21,55272 | 21,31689 | 20,95419 | 21,55042 |
| | tr\|G5EA06\|G5EA06_HUMAN;sp\|Q92552\|RT27_HUMAN;tr\|D6RJC7\|D6RJC7_HUMAN;tr\|D6RH2 | tr\|G5EA06\|G | 2,1188 | 18637000 | 4 | 2 | 0,07530489 | 0,25480127 | 20,09995 | 20,34006 | 20,29412 | 20,17865 | 20,26075 | 20,57015 | 20,61809 | NaN |
| + | sp\|Q71DI3\|H32_HUMAN | sp\|Q71DI3\|H | 250,06 | 2575100000 | 19 | 1 | 0,07615468 | 1,57246733 | 25,45164 | 24,70007 | 26,65499 | 26,22601 | 25,70279 | 27,65799 | 28,54657 | 27,41522 |
| | sp\|Q99832\|TCPH_HUMAN;tr\|F8WAM2\|F8WAM2_HUMAN | sp\|Q99832\|T | 104,33 | 723810000 | 69 | 15 | 0,07626191 | -0,14729309 | 25,50963 | 25,61608 | 25,55195 | 25,74065 | 25,51881 | 25,48873 | 25,31759 | 25,504 |
| | sp\|P12004\|PCNA_HUMAN | sp\|P12004\|P | 7,1592 | 68543000 | 9 | 4 | 0,08043469 | -0,32506752 | 22,09291 | 22,30627 | 22,55705 | 22,53174 | 22,36303 | 21,97722 | 21,8519 | 21,99555 |
| | sp\|P28838\|AMPL_HUMAN;tr\|H0Y983\|H0Y983_HUMAN;tr\|H0Y9Q1\|H0Y9Q1_HUMAN | sp\|P28838\|A | 20,556 | 31896000 | 9 | 6 | 0,08328562 | -0,3528738 | 21,14781 | 21,20234 | NaN | 21,50349 | 20,98266 | 20,9507 | 20,62266 | 21,17066 |
| + | sp\|P40937\|RFC5_HUMAN;tr\|E9PEP3\|E9PEP3_HUMAN;tr\|F5H5S0\|F5H5S0_HUMAN | sp\|P40937\|R | 4,1916 | 53601000 | 5 | 4 | 0,08349595 | 0,53347635 | 21,68752 | 21,52172 | 21,69417 | 21,61679 | 22,05854 | 22,4066 | 21,5048 | 22,68417 |
| | sp\|P38117\|ETFB_HUMAN;tr\|M0QY67\|M0QY67_HUMAN | sp\|P38117\|E | 13,915 | 116010000 | 22 | 7 | 0,08381332 | -0,45845413 | 23,12398 | 23,20044 | 23,04348 | 23,41274 | 23,0234 | 22,63429 | 22,19559 | 23,09354 |
| | sp\|P22314\|UBA1_HUMAN;tr\|Q5JRR6\|Q5JRR6_HUMAN;tr\|Q5JRR9\|Q5JRR9_HUMAN;tr\|Q5JRS( | sp\|P22314\|U | 323,31 | 554990000 | 89 | 22 | 0,08460817 | -0,44711256 | 25,46105 | 25,40224 | 25,40688 | 25,48551 | 24,95082 | 24,86802 | 24,56058 | 25,58781 |
| | sp\|P63261\|ACTG_HUMAN;sp\|P60709\|ACTB_HUMAN;sp\|P63267\|ACTH_HUMAN;sp\|P68133\|A | sp\|P63261\|A | 323,31 | 1,1263E+10 | 85 | 12 | 0,08616405 | -0,39780283 | 29,47628 | 29,72224 | 29,73841 | 30,04726 | 29,41689 | 29,33153 | 28,94737 | 29,69719 |
| | sp\|P51148\|RAB5C_HUMAN;tr\|F8VVK3\|F8VVK3_HUMAN;tr\|K7ERI8\|K7ERI8_HUMAN;tr\|K7ERQ | sp\|P51148\|R | 100,44 | 93073000 | 28 | 2 | 0,08727119 | -0,45473719 | 22,29061 | 23,15372 | 22,65755 | 22,79275 | 22,56519 | 22,42335 | 22,06121 | 22,02594 |
| | sp\|O75534\|CSDE1_HUMAN;tr\|E9PLT0\|E9PLT0_HUMAN;tr\|E9PLD4\|E9PLD4_HUMAN;tr\|E9PNC | sp\|O75534\|C | 17,046 | 84755000 | 17 | 10 | 0,08737773 | -0,17092323 | 22,32501 | 22,66941 | 22,54738 | 22,59735 | 22,39796 | 22,25278 | 22,37472 | 22,43 |
| | sp\|Q7KZF4\|SND1_HUMAN;tr\|H7C597\|H7C597_HUMAN | sp\|Q7KZF4\|S | 32,138 | 156400000 | 50 | 13 | 0,08758826 | -0,32325125 | 23,61189 | 23,4325 | 23,66275 | 23,64724 | 23,26799 | 23,05336 | 23,05303 | 23,68699 |
| | sp\|O43776\|SYNC_HUMAN;tr\|K7EQ35\|K7EQ35_HUMAN;tr\|K7EJ19\|K7EJ19_HUMAN;tr\|K7EIU7 | sp\|O43776\|S | 40,6 | 81152000 | 27 | 10 | 0,08810745 | 0,37095499 | 21,63914 | 21,98096 | 22,22075 | 22,16947 | 22,12921 | 22,56134 | 22,61899 | 22,18459 |
| | sp\|O14818\|PSA7_HUMAN;tr\|H0Y586\|H0Y586_HUMAN;sp\|Q8TAA3\|PSA7L_HUMAN;tr\|F5GY3( | sp\|O14818\|P | 40,174 | 136350000 | 23 | 8 | 0,08915091 | -0,30678129 | 23,29081 | 23,01419 | 23,54085 | 23,27014 | 23,1817 | 23,08508 | 22,69347 | 22,92861 |
| | sp\|P49588\|SYAC_HUMAN;tr\|H3BPK7\|H3BPK7_HUMAN | sp\|P49588\|S | 14,057 | 82194000 | 13 | 6 | 0,08939716 | -0,45694542 | 22,41232 | 22,75609 | 22,78921 | 22,63657 | 22,55852 | 21,86522 | 21,79563 | 22,54706 |
| | sp\|Q9Y333\|LSM2_HUMAN | sp\|Q9Y333\|L | 18,665 | 6663100 | 7 | 2 | 0,08948242 | -0,20594041 | 18,83306 | 18,9851 | 19,08755 | 18,95589 | 18,74931 | 18,92135 | 18,60773 | NaN |
| | sp\|Q96T76\|MMS19_HUMAN;tr\|H0Y746\|H0Y746_HUMAN;tr\|H7C1W5\|H7C1W5_HUMAN;tr\|C | sp\|Q96T76\|N | 10,816 | 26867000 | 7 | 7 | 0,09024501 | 0,22738123 | 20,63343 | 20,60648 | 20,64403 | 20,38911 | 20,61468 | 20,91772 | 20,65255 | 20,99762 |
| | sp\|Q15459\|SF3A1_HUMAN;tr\|F8WC79\|F8WC79_HUMAN;tr\|F8WB66\|F8WB66_HUMAN | sp\|Q15459\|S | 16,783 | 68708000 | 15 | 7 | 0,09090692 | -0,33076096 | 22,34589 | 22,22926 | 22,52499 | 22,28116 | 21,78089 | 21,7383 | 22,18737 | 22,3517 |
| | sp\|O00303\|EIF3F_HUMAN;tr\|H0YDT6\|H0YDT6_HUMAN;tr\|E9PQV8\|E9PQV8_HUMAN;tr\|A0A( | sp\|O00303\|E | 51,276 | 53206000 | 11 | 5 | 0,0920955 | 0,27354097 | 21,2307 | 21,86283 | 21,6759 | 21,71469 | 21,91944 | 21,87916 | 21,87605 | 21,90364 |
| + | sp\|Q15181\|IPYR_HUMAN;tr\|Q5SQT6\|Q5SQT6_HUMAN | sp\|Q15181\|I | 12,569 | 156450000 | 16 | 6 | 0,09238405 | -0,55057859 | 23,2535 | 23,50942 | 23,74366 | 24,09362 | 22,93184 | 23,59069 | 22,61767 | 23,25768 |
| | sp\|Q14103\|HNRPD_HUMAN;tr\|H0YA96\|H0YA96_HUMAN;tr\|H0Y8G5\|H0Y8G5_HUMAN;tr\|D6 | sp\|Q14103\|H | 58,601 | 250710000 | 17 | 7 | 0,09325007 | -0,42933846 | 23,86844 | 23,71931 | 24,1141 | 24,23965 | 23,27469 | 23,51905 | 24,07553 | 23,35488 |
| + | sp\|Q13404\|UB2V1_HUMAN;tr\|I3L0A0\|I3L0A0_HUMAN;tr\|A0A0A0MSL3\|A0A0A0MSL3_HUMA | sp\|Q13404\|U | 3,9341 | 65575000 | 10 | 3 | 0,09381718 | -0,60307471 | 22,05815 | NaN | 22,36185 | 22,4032 | 21,90437 | 22,15761 | 21,07666 | 21,54666 |
| + | sp\|P28340\|DPOD1_HUMAN;tr\|M0R2B7\|M0R2B7_HUMAN;tr\|M0QZR8\|M0QZR8_HUMAN;tr\|N | sp\|P28340\|D | 6,0462 | 15385000 | 10 | 5 | 0,09444932 | 0,56234964 | NaN | 19,95674 | 19,05022 | 19,94392 | 19,94707 | 20,33963 | 20,31628 | 20,2476 |
| | tr\|E9PKP7\|E9PKP7_HUMAN;sp\|P17480\|UBF1_HUMAN;tr\|E9PLT2\|E9PLT2_HUMAN | tr\|E9PKP7\|E | 2,4579 | 17171000 | 3 | 2 | 0,09532972 | -0,12893804 | NaN | 20,18302 | 20,39352 | 20,2383 | 20,11553 | 20,09275 | 20,22482 | 20,13759 |
| + | tr\|D6R974\|D6R974_HUMAN;sp\|P09936\|UCHL1_HUMAN;tr\|D6R956\|D6R956_HUMAN;tr\|D6RI | tr\|D6R974\|D | 39,807 | 134180000 | 13 | 4 | 0,09560425 | -0,5838294 | 23,0581 | 23,21079 | 23,74755 | 23,48296 | 22,69341 | 23,4025 | 22,17725 | 22,89093 |
| | sp\|P40227\|TCPZ_HUMAN;sp\|Q92526\|TCPW_HUMAN;tr\|J3KRI6\|J3KRI6_HUMAN | sp\|P40227\|T | 320,1 | 1048700000 | 53 | 19 | 0,09645953 | 0,20556927 | 25,93818 | 25,87352 | 25,83328 | 26,01148 | 25,94829 | 26,35788 | 25,97631 | 26,19625 |
| | tr\|Q3BDU5\|Q3BDU5_HUMAN;sp\|P02545\|LMNA_HUMAN;tr\|Q5TCI8\|Q5TCI8_HUMAN;tr\|H0Y/ | tr\|Q3BDU5\|C | 22,935 | 99286000 | 24 | 11 | 0,09711616 | 0,28241062 | 22,48242 | 22,12436 | 22,30446 | 22,63112 | 22,4877 | 22,6093 | 22,92723 | 22,64777 |
| + | sp\|Q709C8\|VP13C_HUMAN | sp\|Q709C8\|V | 24,222 | 59499000 | 24 | 15 | 0,09869647 | -1,61485052 | 21,28771 | 22,67805 | 21,48045 | 23,84163 | 21,56973 | 20,53403 | 21,5833 | 19,14138 |
| | sp\|P07237\|PDIA1_HUMAN;tr\|H7BZ94\|H7BZ94_HUMAN;tr\|H0Y3Z3\|H0Y3Z3_HUMAN;tr\|I3L39 | sp\|P07237\|P | 123,6 | 265930000 | 38 | 14 | 0,09920975 | 0,17733622 | 23,87961 | 23,88995 | 24,12258 | 24,04419 | 24,05532 | 24,30382 | 24,03238 | 24,25415 |
| + | sp\|Q01469\|FABP5_HUMAN;tr\|I6L8B7\|I6L8B7_HUMAN | sp\|Q01469\|F | 47,63 | 106070000 | 15 | 3 | 0,09927779 | -0,6759367 | 23,26728 | 23,27668 | 23,02102 | NaN | 22,91989 | 22,90896 | 21,74263 | 22,47808 |
| | tr\|E7ES33\|E7ES33_HUMAN;tr\|E7EPK1\|E7EPK1_HUMAN;sp\|Q16181\|SEPT7_HUMAN;tr\|G3V1C | tr\|E7ES33\|E | 7,2208 | 50028000 | 12 | 5 | 0,09982743 | -0,31792879 | 21,53488 | 21,82695 | 21,72974 | 21,67401 | 21,419 | 21,27378 | 21,03812 | 21,76296 |
| | sp\|P45974\|UBP5_HUMAN;tr\|F5H571\|F5H571_HUMAN | sp\|P45974\|U | 28,481 | 19869000 | 13 | 6 | 0,10069731 | -0,4816103 | 20,45342 | 20,76733 | 20,62979 | 20,56086 | 19,70315 | 20,18507 | 19,82537 | 20,77137 |
| | sp\|P46779\|RL28_HUMAN;tr\|H0YKD8\|H0YKD8_HUMAN;tr\|H0YLP6\|H0YLP6_HUMAN;tr\|H0YMI | sp\|P46779\|R | 18,748 | 194450000 | 9 | 4 | 0,10229955 | -0,32105637 | 24,1043 | 23,97176 | 23,74929 | 23,56523 | 23,55063 | 23,41455 | 23,84029 | 23,3009 |
| | sp\|P11172\|UMPS_HUMAN;tr\|E9PFD2\|E9PFD2_HUMAN | sp\|P11172\|U | 3,2636 | 23664000 | 10 | 3 | 0,10272557 | 0,4859004 | 19,7552 | 20,71339 | 20,37029 | 20,29849 | 20,7432 | 21,21721 | 20,5093 | 20,61126 |
| | sp\|Q9Y295\|DRG1_HUMAN | sp\|Q9Y295\|D | 4,4206 | 14367000 | 5 | 2 | 0,10326112 | -0,36081839 | 20,23841 | 20,1061 | 20,10738 | 20,23351 | 19,99397 | 19,48695 | 19,52081 | 20,24039 |
| | tr\|H7C1U0\|H7C1U0_HUMAN;sp\|P13798\|ACPH_HUMAN;tr\|C9JIF9\|C9JIF9_HUMAN;tr\|C9JLK2\| | tr\|H7C1U0\|H | 5,777 | 9411700 | 4 | 2 | 0,1037114 | -0,17235422 | 19,44581 | 19,40707 | 19,37063 | 19,55165 | 19,23804 | 19,19648 | 19,14385 | 19,50736 |
| | sp\|P12273\|PIP_HUMAN | sp\|P12273\|P | 14,554 | 31158000 | 8 | 4 | 0,10430597 | -0,41471195 | 21,64961 | 20,97939 | 20,82943 | 21,63923 | 20,87065 | 20,80554 | 20,90566 | 20,85696 |
| | sp\|P62701\|RS4X_HUMAN;tr\|C9JEH7\|C9JEH7_HUMAN;sp\|P22090\|RS4Y1_HUMAN;sp\|Q8TD47 | sp\|P62701\|R | 56,941 | 994590000 | 35 | 8 | 0,10509357 | -0,37617731 | 26,10159 | 26,35301 | 26,38096 | 25,87516 | 25,55514 | 25,9205 | 26,19263 | 25,53774 |
| | tr\|B4DXZ6\|B4DXZ6_HUMAN;sp\|P51114\|FXR1_HUMAN;tr\|E7EU85\|E7EU85_HUMAN;tr\|E9PFF! | tr\|B4DXZ6\|B | 75,704 | 40630000 | 17 | 3 | 0,10655255 | 0,34382725 | 21,29995 | 20,86537 | 21,43939 | 20,75197 | 21,27355 | 21,45287 | 21,38527 | 21,62029 |
| | tr\|J3QSB4\|J3QSB4_HUMAN;sp\|P26373\|RL13_HUMAN | tr\|J3QSB4\|J3 | 323,31 | 270640000 | 7 | 3 | 0,10655361 | -0,51061201 | 24,80454 | 24,3536 | 24,72728 | 23,92624 | 23,82474 | 23,62232 | 23,86872 | 24,45344 |
| | sp\|P30520\|PURA2_HUMAN | sp\|P30520\|P | 28,394 | 47076000 | 17 | 6 | 0,10712594 | 0,44738102 | 21,53853 | 21,28884 | 21,44935 | 20,86688 | 21,82236 | 21,5285 | 21,37449 | 22,20777 |
| | sp\|Q9UMS4\|PRP19_HUMAN;tr\|F5GY56\|F5GY56_HUMAN;tr\|H0YGF3\|H0YGF3_HUMAN;tr\|F5H | sp\|Q9UMS4\| | 16,806 | 233790000 | 20 | 7 | 0,10935396 | 0,27235365 | 23,37663 | 23,61897 | 23,96543 | 23,60851 | 23,80754 | 23,94735 | 24,1255 | 23,77856 |
| | tr\|E9PCY7\|E9PCY7_HUMAN;sp\|P31943\|HNRH1_HUMAN;tr\|G8JLB6\|G8JLB6_HUMAN;tr\|D6RIU | tr\|E9PCY7\|E | 276,89 | 550200000 | 48 | 4 | 0,11098454 | 0,2419076 | 24,70525 | 24,80966 | 24,97631 | 25,2787 | 25,08865 | 25,19197 | 25,22494 | 25,23199 |
| | sp\|P10809\|CH60_HUMAN;tr\|E7EXB4\|E7EXB4_HUMAN;tr\|E7ESH4\|E7ESH4_HUMAN;tr\|C9JL25 | sp\|P10809\|C | 323,31 | 5182700000 | 125 | 32 | 0,11323955 | -0,24165916 | 28,17131 | 28,45787 | 28,50154 | 28,77859 | 28,14375 | 28,29321 | 28,1969 | 28,30882 |
| | sp\|Q8N163\|CCAR2_HUMAN;tr\|G3V119\|G3V119_HUMAN;tr\|H0YB24\|H0YB24_HUMAN;tr\|H0Y | sp\|Q8N163\|C | 17,731 | 51214000 | 22 | 8 | 0,11535343 | 0,39793968 | 21,516 | 21,49634 | 21,68787 | 21,0764 | 21,67077 | 21,45478 | 22,01356 | 22,22926 |
| | tr\|C9JBI3\|C9JBI3_HUMAN;sp\|P78330\|SERB_HUMAN | tr\|C9JBI3\|C9J | 4,506 | 14356000 | 4 | 3 | 0,11608391 | -0,18942515 | 20,10457 | NaN | 20,06815 | 19,96113 | 20,02351 | 19,78094 | 19,76113 | NaN |
| | sp\|Q08J23\|NSUN2_HUMAN;tr\|A0A140T9Y7\|A0A140T9Y7_HUMAN | sp\|Q08J23\|N | 11,019 | 49579000 | 15 | 5 | 0,11610128 | 0,2670579 | 21,25119 | 21,63378 | 21,40401 | 21,58192 | 21,44779 | 21,9333 | 21,64342 | 21,91463 |
| | sp\|P11586\|C1TC_HUMAN;tr\|F5H2F4\|F5H2F4_HUMAN;tr\|V9GYY3\|V9GYY3_HUMAN | sp\|P11586\|C | 93,415 | 417540000 | 76 | 23 | 0,11618624 | 0,18716192 | 24,54251 | 24,78362 | 24,56721 | 24,80563 | 24,92034 | 24,79313 | 24,69588 | 25,03825 |
| | sp\|Q15185\|TEBP_HUMAN;tr\|A0A087WYT3\|A0A087WYT3_HUMAN | sp\|Q15185\|T | 18,443 | 48910000 | 13 | 3 | 0,11656907 | -0,46360159 | 21,66375 | 21,97456 | 22,06046 | 22,28881 | 21,38158 | 22,1169 | 21,08245 | 21,55225 |
| | sp\|P04843\|RPN1_HUMAN;tr\|B7Z4L4\|B7Z4L4_HUMAN;tr\|F8WF32\|F8WF32_HUMAN | sp\|P04843\|R | 96,578 | 243140000 | 47 | 12 | 0,11977585 | -0,18811512 | 23,96288 | 24,00573 | 24,19746 | 24,08966 | 23,89901 | 23,86656 | 24,08837 | 23,64933 |
| | tr\|C9J8B8\|C9J8B8_HUMAN;sp\|Q969S8\|HDA10_HUMAN | tr\|C9J8B8\|C9 | 1,6331 | 4228600 | 6 | 1 | 0,12041853 | -0,54951874 | NaN | 18,88804 | 18,65179 | 18,3169 | 18,33447 | 17,42955 | 18,3699 | 18,14365 |
| | sp\|P13010\|XRCC5_HUMAN;tr\|C9JZ81\|C9JZ81_HUMAN;tr\|H7C0H9\|H7C0H9_HUMAN | sp\|P13010\|X | 168,84 | 295950000 | 50 | 17 | 0,12253405 | -0,2440424 | 24,50936 | 24,20053 | 24,64581 | 24,36853 | 23,90159 | 24,24235 | 24,32574 | 24,27838 |
| | sp\|O15067\|PUR4_HUMAN;tr\|H0YGH1\|H0YGH1_HUMAN;tr\|J3KTQ5\|J3KTQ5_HUMAN;tr\|J3KTS | sp\|O15067\|P | 38,998 | 88998000 | 22 | 9 | 0,12256413 | 0,32659864 | 22,44096 | 22,36933 | 22,2247 | 22,22635 | 22,64625 | 22,26396 | 22,55569 | 23,10183 |
| + | sp\|P01876\|IGHA1_HUMAN | sp\|P01876\|I | 4,0982 | 27526000 | 6 | 2 | 0,12271062 | -1,20516332 | 22,65513 | NaN | 20,60991 | 21,12452 | 20,28704 | 20,13646 | 20,35057 | NaN |
| | sp\|P47897\|SYQ_HUMAN;tr\|A0A0U1RQT0\|A0A0U1RQT0_HUMAN;tr\|A0A0U1RQM8\|A0A0U1R( | sp\|P47897\|S | 36,361 | 138240000 | 26 | 14 | 0,12595124 | -0,14303923 | 23,37518 | 23,19768 | 23,4467 | 23,19301 | 23,11009 | 23,06154 | 23,28673 | 23,18204 |
| | tr\|A0A087WYU1\|A0A087WYU1_HUMAN;sp\|Q9Y5X1\|SNX9_HUMAN | tr\|A0A087W | 8,4674 | 10741000 | 6 | 3 | 0,12632349 | 0,16749048 | 19,62412 | 19,35954 | 19,4616 | 19,52338 | 19,75589 | 19,50959 | 19,71347 | NaN |
| | sp\|Q8N766\|EMC1_HUMAN;tr\|Q5TG59\|Q5TG59_HUMAN | sp\|Q8N766\|E | 17,418 | 39526000 | 15 | 11 | 0,12762022 | 0,32606554 | 20,90492 | 20,91933 | 21,00093 | 20,82306 | 20,96077 | 21,0466 | 21,76332 | 21,1818 |
| | sp\|P49257\|LMAN1_HUMAN | sp\|P49257\|L | 7,6047 | 57603000 | 6 | 3 | 0,12812458 | 0,42639494 | 21,55581 | 21,60598 | 21,74833 | 21,48493 | 21,67263 | 22,30003 | 22,54399 | 21,58399 |
| | sp\|P16989\|YBOX3_HUMAN;tr\|A0A0D9SEI8\|A0A0D9SEI8_HUMAN | sp\|P16989\|Y | 52,366 | 31941000 | 8 | 3 | 0,13050955 | -0,29836559 | 20,97435 | 20,83368 | 21,3849 | 21,22317 | 20,71482 | 20,63741 | 21,15351 | 20,71691 |
| | sp\|P46977\|STT3A_HUMAN;tr\|A0A0C4DH80\|A0A0C4DH80_HUMAN;tr\|E9PN73\|E9PN73_HUM/ | sp\|P46977\|S | 3,4546 | 29146000 | 4 | 3 | 0,13122143 | -0,56363153 | 20,9693 | 21,23806 | 21,98621 | 21,06848 | NaN | 20,68765 | 21,0959 | 20,47209 |
| | sp\|P55884\|EIF3B_HUMAN;tr\|C9JQN7\|C9JQN7_HUMAN;tr\|C9JZG1\|C9JZG1_HUMAN | sp\|P55884\|E | 29,746 | 147490000 | 21 | 10 | 0,131658 | -0,24601269 | 23,48333 | 23,3948 | 23,51977 | 23,20271 | 23,0443 | 22,98536 | 23,07097 | 23,51593 |
| | sp\|O60701\|UGDH_HUMAN;tr\|D6RHF4\|D6RHF4_HUMAN;tr\|E9PBD2\|E9PBD2_HUMAN;tr\|E7EF | sp\|O60701\|U | 3,8425 | 17435000 | 10 | 3 | 0,13325023 | -0,51021385 | 20,41473 | 20,56123 | 20,27435 | 20,76871 | 20,35402 | 19,79992 | 19,31134 | 20,51287 |
| | sp\|Q9P0L0\|VAPA_HUMAN;tr\|J3QKM9\|J3QKM9_HUMAN | sp\|Q9P0L0\|V | 14,683 | 60335000 | 12 | 3 | 0,1335238 | 0,4712615 | 21,20467 | 21,95787 | 22,0483 | 21,91605 | 22,09152 | 22,10776 | 22,55967 | NaN |
| | tr\|G3V1B3\|G3V1B3_HUMAN;tr\|M0R181\|M0R181_HUMAN;sp\|P46778\|RL21_HUMAN | tr\|G3V1B3\|G | 3,956 | 193080000 | 6 | 3 | 0,13466814 | -0,40020561 | 23,98833 | 23,79729 | 24,37465 | 23,48726 | NaN | 23,50543 | 23,61054 | 23,41905 |
| | sp\|P04406\|G3P_HUMAN;tr\|E7EUT5\|E7EUT5_HUMAN;sp\|O14556\|G3PT_HUMAN | sp\|P04406\|G | 323,31 | 1583200000 | 43 | 9 | 0,13521725 | -0,37837029 | 26,65676 | 26,64636 | 27,0203 | 27,15364 | 26,4447 | 26,65349 | 26,01844 | 26,84697 |
| | tr\|A0A0A0MRA5\|A0A0A0MRA5_HUMAN;tr\|B7Z4B8\|B7Z4B8_HUMAN;sp\|Q9BUJ2\|HNRL1_HUI | tr\|A0A0A0MI | 11,932 | 38540000 | 12 | 5 | 0,13540792 | -0,21211433 | 21,44035 | 21,3048 | 21,64917 | 21,38427 | 21,38021 | 21,00867 | 21,12622 | 21,41504 |

| | | | | | | | | | | | | | | | |
|---|---|---|---|---|---|---|---|---|---|---|---|---|---|---|---|
| sp\|Q15046\|SYK_HUMAN;tr\|J3KRL2\|J3KRL2_HUMAN;tr\|H3BPV7\|H3BPV7_HUMAN | sp\|Q15046\|S | 7,4094 | 22847000 | 12 | 3 | 0,13570844 | -0,29636908 | 20,58838 | 20,40312 | 20,86771 | 20,74911 | 20,1545 | 20,36954 | 20,74517 | 20,15363 |
| sp\|Q13561\|DCTN2_HUMAN;tr\|H0YI98\|H0YI98_HUMAN;tr\|F8W1I6\|F8W1I6_HUMAN;tr\|F8VW | sp\|Q13561\|D | 7,2322 | 35801000 | 13 | 6 | 0,13758418 | -0,15215111 | 21,15728 | 21,20371 | 21,34066 | 21,26264 | 20,87673 | 21,2191 | 21,05729 | 21,20258 |
| sp\|P51858\|HDGF_HUMAN | sp\|P51858\|H | 15,619 | 80460000 | 14 | 4 | 0,13960727 | -0,42066813 | 22,56086 | 22,54328 | 22,30123 | 22,75654 | 21,90936 | 22,40026 | 21,58325 | 22,58637 |
| sp\|P18124\|RL7_HUMAN;tr\|A8MUD9\|A8MUD9_HUMAN;tr\|C9JIJ5\|C9JIJ5_HUMAN;tr\|C9JZ88\|( | sp\|P18124\|R | 52,349 | 584430000 | 42 | 10 | 0,1411243 | -0,32907152 | 25,17043 | 25,50054 | 25,60908 | 25,14767 | 24,91862 | 25,13073 | 25,39663 | 24,66546 |
| sp\|P10599\|THIO_HUMAN | sp\|P10599\|T | 38,109 | 233240000 | 13 | 5 | 0,14169246 | 0,48632813 | 23,98616 | 23,92071 | 23,20939 | 23,47209 | 24,43619 | 24,2467 | 23,48444 | 24,36633 |
| sp\|P05109\|S10A8_HUMAN | sp\|P05109\|S | 7,3461 | 66786000 | 9 | 3 | 0,14510511 | -0,54869223 | 23,11824 | 22,65176 | 21,77278 | 22,62536 | 22,23386 | 22,26436 | 21,53269 | 21,94246 |
| tr\|J3QLR8\|J3QLR8_HUMAN;sp\|Q9Y3D9\|RT23_HUMAN | tr\|J3QLR8\|J3 | 5,7921 | 26361000 | 7 | 4 | 0,14645904 | -0,15183115 | 20,82376 | 20,81026 | 21,12408 | 20,96684 | 20,76247 | 20,81855 | NaN | 20,75719 |
| sp\|P81605\|DCD_HUMAN | sp\|P81605\|D | 6,5784 | 148150000 | 11 | 3 | 0,14690426 | -0,39113569 | 23,75929 | 23,46139 | 23,38034 | 23,67206 | 23,75348 | 22,7633 | 23,25609 | 22,93566 |
| sp\|P63151\|2ABA_HUMAN;tr\|E5RFR9\|E5RFR9_HUMAN;tr\|E5RIY1\|E5RIY1_HUMAN;sp\|Q00005 | sp\|P63151\|2 | 11,782 | 59601000 | 20 | 6 | 0,14746177 | 0,18046856 | 21,53588 | 21,76927 | 21,81769 | 21,86876 | 21,98211 | 21,89667 | 21,72713 | 22,10757 |
| sp\|P33176\|KINH_HUMAN;tr\|A0A0G2JMZ6\|A0A0G2JMZ6_HUMAN;tr\|J3KNA1\|J3KNA1_HUMAI | sp\|P33176\|K | 26,828 | 67643000 | 38 | 12 | 0,14750611 | -0,31894732 | 22,40026 | 22,35305 | 22,36076 | 22,27674 | 22,03842 | 21,53564 | 22,07858 | 22,46236 |
| tr\|E9PLK3\|E9PLK3_HUMAN;sp\|P55786\|PSA_HUMAN;sp\|A6NEC2\|PSAL_HUMAN;tr\|H0YCQ5\|H | tr\|E9PLK3\|E9 | 18,983 | 121930000 | 25 | 11 | 0,14955002 | -0,18057251 | 22,82409 | 23,08413 | 22,87793 | 23,12605 | 22,86221 | 22,69211 | 22,64414 | 22,99144 |
| sp\|Q13243\|SRSF5_HUMAN;tr\|G3V5K8\|G3V5K8_HUMAN;tr\|B4DJK0\|B4DJK0_HUMAN;tr\|B4DU | sp\|Q13243\|S | 12,33 | 23300000 | 15 | 3 | 0,15406353 | -0,40751839 | 19,9073 | 20,51528 | 20,54952 | 20,13884 | 19,51032 | 20,06697 | 20,32663 | 19,57694 |
| sp\|P34932\|HSP74_HUMAN;tr\|A0A087WYC1\|A0A087WYC1_HUMAN;tr\|A0A087WTS8\|A0A087 | sp\|P34932\|H | 115,98 | 224170000 | 40 | 14 | 0,15669938 | 0,12815237 | 23,829 | 23,91369 | 23,71471 | 23,75746 | 23,80361 | 23,83306 | 24,03036 | 24,06044 |
| tr\|F8W1R7\|F8W1R7_HUMAN;sp\|P60660\|MYL6_HUMAN;tr\|J3KND3\|J3KND3_HUMAN;tr\|G8JL | tr\|F8W1R7\|F | 6,9853 | 55008000 | 10 | 4 | 0,15719874 | -0,19595098 | 21,96782 | 21,88807 | 22,01086 | 22,1549 | 22,08832 | 21,85068 | 21,5855 | 21,71335 |
| sp\|Q9UBE0\|SAE1_HUMAN;tr\|M0QZS6\|M0QZS6_HUMAN;tr\|M0QX65\|M0QX65_HUMAN;tr\|B3 | sp\|Q9UBE0\|S | 71,714 | 31240000 | 13 | 7 | 0,15811278 | -0,49092102 | 21,02378 | 21,19701 | 21,47788 | 21,87429 | 20,9132 | 21,20508 | 20,21703 | 21,27395 |
| sp\|Q9UBB6\|NCDN_HUMAN;tr\|C9J5H8\|C9J5H8_HUMAN | sp\|Q9UBB6\|N | 3,9546 | 8359200 | 8 | 2 | 0,15903865 | 0,3370444 | 18,9922 | NaN | 18,94524 | 19,27542 | NaN | 19,10319 | 19,44845 | 19,67235 |
| sp\|P36404\|ARL2_HUMAN | sp\|P36404\|A | 3,0058 | 14781000 | 5 | 2 | 0,15912224 | 0,8961633 | NaN | 19,35281 | 19,26091 | 19,30672 | 21,32405 | 20,57404 | 19,50999 | 19,40383 |
| sp\|O15355\|PPM1G_HUMAN | sp\|O15355\|P | 13,976 | 37478000 | 13 | 5 | 0,15918311 | -0,39740038 | 21,6189 | 21,25523 | 21,813 | 21,53022 | 21,09539 | 21,13515 | 20,66709 | 21,73012 |
| sp\|P34897\|GLYM_HUMAN;tr\|H0YIZ0\|H0YIZ0_HUMAN;tr\|G3V5L0\|G3V5L0_HUMAN;tr\|G3V2Y | sp\|P34897\|G | 54,759 | 334280000 | 38 | 13 | 0,15927192 | -0,27146053 | 24,59206 | 24,3573 | 24,69678 | 24,82026 | 24,30932 | 24,30375 | 24,05142 | 24,71607 |
| sp\|Q14204\|DYHC1_HUMAN;tr\|H0YJ21\|H0YJ21_HUMAN | sp\|Q14204\|D | 159,39 | 464100000 | 110 | 58 | 0,16078751 | 0,24344444 | 24,68528 | 24,86858 | 24,68233 | 24,54828 | 24,57323 | 25,01665 | 25,23257 | 24,9358 |
| sp\|P26196\|DDX6_HUMAN;tr\|Q8IV96\|Q8IV96_HUMAN | sp\|P26196\|D | 11,828 | 43629000 | 13 | 6 | 0,16197725 | -0,22703981 | 21,27855 | 21,60937 | 21,8273 | 21,7195 | 21,52363 | 21,44486 | 21,15845 | 21,39963 |
| sp\|P24390\|ERD21_HUMAN;tr\|M0R1Y2\|M0R1Y2_HUMAN | sp\|P24390\|E | 43,566 | 67132000 | 20 | 2 | 0,16295209 | 0,58507633 | 22,0124 | 20,96212 | 22,2287 | 21,27923 | 22,50582 | 22,35081 | 22,39616 | 21,56997 |
| sp\|Q9NVP1\|DDX18_HUMAN;tr\|H7C452\|H7C452_HUMAN | sp\|Q9NVP1\|D | 5,2104 | 10479000 | 6 | 3 | 0,16438328 | 0,66341591 | 19,13016 | 18,78625 | 19,19302 | 18,06844 | 18,67731 | 20,05179 | 19,95617 | 19,14627 |
| sp\|Q13409\|DC1I2_HUMAN;tr\|E9PGG1\|E9PGG1_HUMAN;tr\|E7EUM4\|E7EUM4_HUMAN;tr\|E7I | sp\|Q13409\|D | 27,671 | 70679000 | 9 | 3 | 0,16439875 | -0,56391144 | 22,224 | 22,22205 | 22,45249 | 23,20653 | 22,26316 | 21,87867 | 21,24197 | 22,46563 |
| sp\|P00492\|HPRT_HUMAN | sp\|P00492\|H | 14,916 | 90418000 | 22 | 6 | 0,1679295 | -0,18386412 | 22,24284 | 22,46037 | 22,66573 | 22,55985 | 22,19376 | 22,51944 | 22,21877 | 22,26135 |
| sp\|Q92616\|GCN1_HUMAN | sp\|Q92616\|G | 200,78 | 257500000 | 81 | 36 | 0,16868516 | 0,36971331 | 23,40341 | 23,97544 | 23,51509 | 23,83547 | 23,51328 | 24,22949 | 24,4168 | 24,04868 |
| sp\|P33991\|MCM4_HUMAN;tr\|E5RG31\|E5RG31_HUMAN;tr\|E5RFJ8\|E5RFJ8_HUMAN | sp\|P33991\|M | 22,808 | 133630000 | 28 | 13 | 0,16966084 | 0,16920424 | 23,01141 | 22,90533 | 22,9985 | 23,15417 | 22,99016 | 23,05752 | 23,3243 | 23,37425 |
| tr\|K7EJE8\|K7EJE8_HUMAN;tr\|K7EKE6\|K7EKE6_HUMAN;sp\|P36776\|LONM_HUMAN;tr\|K7ER27 | tr\|K7EJE8\|K7 | 11,265 | 35084000 | 22 | 6 | 0,17093158 | -0,34970856 | 21,28433 | 21,46339 | 21,33222 | 21,60616 | 21,01025 | 20,70868 | 20,88517 | 21,68316 |
| sp\|O15160\|RPAC1_HUMAN;tr\|E7EQB9\|E7EQB9_HUMAN;tr\|D6RDJ3\|D6RDJ3_HUMAN;tr\|H0Y | sp\|O15160\|R | 7,9664 | 15572000 | 8 | 3 | 0,17159407 | 0,25740671 | NaN | 19,93229 | 20,01808 | 19,7957 | 20,20693 | 19,88011 | 20,1047 | 20,49931 |
| sp\|P24534\|EF1B_HUMAN;tr\|F2Z2G2\|F2Z2G2_HUMAN;tr\|C9JZW3\|C9JZW3_HUMAN;tr\|F8WF6 | sp\|P24534\|E | 121,45 | 156350000 | 8 | 3 | 0,17190672 | -0,2573781 | 23,13016 | 23,3868 | 23,53172 | 23,72067 | 23,0353 | 23,3507 | 22,95955 | 23,39428 |
| sp\|Q00059\|TFAM_HUMAN;tr\|H7BYN3\|H7BYN3_HUMAN | sp\|Q00059\|T | 3,4353 | 72939000 | 7 | 3 | 0,17592566 | -0,33678722 | 21,87804 | 22,47586 | 22,51058 | 22,69271 | 22,02296 | 22,04126 | 22,38987 | 21,75593 |
| tr\|A0A0B4J1W3\|A0A0B4J1W3_HUMAN;sp\|Q9BXJ9\|NAA15_HUMAN;sp\|Q6N069\|NAA16_HUM | tr\|A0A0B4J1W | 12,039 | 23147000 | 8 | 5 | 0,17775511 | -0,21805843 | 20,57228 | 20,53136 | 20,86718 | 20,58683 | NaN | 20,43288 | 20,19785 | 20,63334 |
| sp\|P78527\|PRKDC_HUMAN | sp\|P78527\|P | 273,63 | 572230000 | 145 | 55 | 0,1787566 | 0,20104694 | 24,98011 | 25,15661 | 24,8739 | 25,04181 | 24,8624 | 25,29204 | 25,35867 | 25,3435 |
| sp\|P61106\|RAB14_HUMAN;tr\|X6RFL8\|X6RFL8_HUMAN | sp\|P61106\|R | 93,583 | 249500000 | 27 | 8 | 0,17893438 | -0,53139353 | 23,406 | 24,52898 | 24,31693 | 24,27987 | 23,61728 | 24,28221 | 23,34516 | 23,16156 |
| sp\|O43395\|PRPF3_HUMAN | sp\|O43395\|P | 2,7448 | 8852900 | 3 | 2 | 0,18304553 | -0,22639132 | 19,48911 | 19,27912 | 19,32133 | 19,33756 | 19,02948 | 18,79592 | 19,22651 | 19,46965 |
| sp\|Q14008\|CKAP5_HUMAN;tr\|H0YDX5\|H0YDX5_HUMAN | sp\|Q14008\|C | 55,078 | 31429000 | 19 | 9 | 0,18405575 | 0,19549036 | 21,04733 | 20,89267 | 20,90139 | 20,73355 | 20,74985 | 21,21537 | 21,18101 | 21,21068 |
| sp\|P26599\|PTBP1_HUMAN;tr\|A0A0U1RRM4\|A0A0U1RRM4_HUMAN;tr\|A6NLN1\|A6NLN1_HU | sp\|P26599\|P | 148,16 | 424820000 | 51 | 13 | 0,18493656 | -0,29860687 | 25,01979 | 24,6371 | 24,91534 | 24,92089 | 24,07022 | 24,59309 | 24,71046 | 24,92493 |
| tr\|F8WAU4\|F8WAU4_HUMAN;tr\|C9IZ01\|C9IZ01_HUMAN;sp\|Q96RP9\|EFGM_HUMAN | tr\|F8WAU4\|F | 13,996 | 41660000 | 20 | 7 | 0,18861781 | 0,16956425 | 21,41875 | 21,34646 | 21,34061 | 21,53825 | 21,62123 | 21,43858 | 21,4026 | 21,85992 |
| sp\|Q8NBJ5\|GT251_HUMAN;tr\|M0QYH0\|M0QYH0_HUMAN;tr\|M0QX72\|M0QX72_HUMAN | sp\|Q8NBJ5\|G | 8,4581 | 32379000 | 11 | 5 | 0,19254319 | -0,27780724 | 20,8155 | 21,20848 | 21,32899 | 21,66384 | 21,09642 | 21,01311 | 21,02823 | 20,76782 |
| tr\|A2A2D0\|A2A2D0_HUMAN;sp\|P16949\|STMN1_HUMAN;tr\|E5RGX5\|E5RGX5_HUMAN;sp\|Q9 | tr\|A2A2D0\|A | 5,1669 | 53851000 | 3 | 2 | 0,19692365 | -0,40443802 | 22,12918 | 21,89138 | 21,92027 | 22,09861 | 21,81355 | 22,19214 | 21,51167 | 20,90434 |
| sp\|Q99613\|EIF3C_HUMAN;sp\|B5ME19\|EIFCL_HUMAN;tr\|H3BPE3\|H3BPE3_HUMAN;tr\|H3BPE | sp\|Q99613\|E | 17,126 | 145800000 | 18 | 8 | 0,19909856 | -0,31782198 | 23,68453 | 23,36132 | 23,30647 | 22,8303 | 23,06911 | 22,64092 | 23,27071 | 22,93059 |
| sp\|P26639\|SYTC_HUMAN;tr\|D6RBR8\|D6RBR8_HUMAN;tr\|D6RCA5\|D6RCA5_HUMAN;tr\|D6R9 | sp\|P26639\|S | 28,354 | 131410000 | 29 | 13 | 0,20135263 | -0,20811462 | 23,3362 | 23,22142 | 23,22005 | 23,29011 | 23,12907 | 23,03539 | 22,6917 | 23,37915 |
| sp\|P39019\|RS19_HUMAN;tr\|A0A075B6E2\|A0A075B6E2_HUMAN;tr\|M0R2L9\|M0R2L9_HUMAI | sp\|P39019\|R | 24,3 | 243710000 | 19 | 6 | 0,2031218 | -0,34204435 | 24,6284 | 23,67066 | 24,09869 | 24,30855 | 24,10406 | 23,91743 | 23,48087 | 23,83576 |
| tr\|E9PD53\|E9PD53_HUMAN;sp\|Q9NTJ3\|SMC4_HUMAN;tr\|C9JR83\|C9JR83_HUMAN | tr\|E9PD53\|E | 33,535 | 11944000 | 12 | 4 | 0,20403778 | 0,26890198 | 19,42173 | 19,7376 | NaN | 19,39427 | 19,77077 | 19,42577 | 19,88527 | 20,06526 |
| tr\|E9PMD8\|E9PMD8_HUMAN;tr\|E9PKD5\|E9PKD5_HUMAN;tr\|E9PM69\|E9PM69_HUMAN;tr\|R | tr\|E9PMD8\|E | 9,1517 | 40658000 | 12 | 5 | 0,20709165 | 0,45402431 | 21,19063 | 20,52965 | 20,84023 | 20,99106 | 21,21555 | 21,34706 | 22,10287 | 20,70219 |
| sp\|P53396\|ACLY_HUMAN;tr\|K7ESG8\|K7ESG8_HUMAN;tr\|K7EIE7\|K7EIE7_HUMAN | sp\|P53396\|A | 87,441 | 230470000 | 47 | 12 | 0,212533 | -0,22579288 | 24,22817 | 24,06587 | 24,06266 | 23,9581 | 23,49692 | 24,02183 | 23,71796 | 24,17493 |
| tr\|H0Y4R1\|H0Y4R1_HUMAN;sp\|P12268\|IMDH2_HUMAN;tr\|E7ETK5\|E7ETK5_HUMAN | tr\|H0Y4R1\|H | 93,24 | 202260000 | 38 | 10 | 0,21384732 | -0,14635611 | 23,85043 | 23,73913 | 23,99412 | 23,74324 | 23,65895 | 23,67508 | 23,49351 | 23,91397 |
| sp\|P55036\|PSMD4_HUMAN;tr\|Q5VWC4\|Q5VWC4_HUMAN;tr\|A6PVX3\|A6PVX3_HUMAN;tr\|H | sp\|P55036\|P | 9,9217 | 37102000 | 4 | 4 | 0,21409296 | -0,10973438 | 21,39305 | NaN | 21,51123 | 21,39021 | 21,19394 | 21,38042 | 21,45453 | 21,25816 |
| sp\|O00116\|ADAS_HUMAN;tr\|B8ZZ81\|B8ZZ81_HUMAN | sp\|O00116\|A | 9,4745 | 15289000 | 9 | 6 | 0,21738728 | -0,16874313 | NaN | 20,21703 | 20,32531 | 20,24621 | NaN | 19,87263 | 20,20383 | 20,20586 |
| tr\|H0YMZ1\|H0YMZ1_HUMAN;tr\|H0YL69\|H0YL69_HUMAN;sp\|P25789\|PSA4_HUMAN;tr\|H0YM | tr\|H0YMZ1\|H | 8,5554 | 120950000 | 12 | 5 | 0,2180686 | -0,31953764 | 23,25123 | 22,80532 | 23,52241 | 22,98251 | 22,72096 | 22,90773 | 22,41841 | 23,23621 |
| sp\|Q9NZ01\|TECR_HUMAN;tr\|M0R3C3\|M0R3C3_HUMAN;tr\|M0QXM3\|M0QXM3_HUMAN | sp\|Q9NZ01\|T | 13,032 | 43106000 | 9 | 5 | 0,21934331 | -0,22462368 | 21,50983 | 21,34435 | 21,77668 | 21,38142 | 21,11997 | 21,35014 | 21,61598 | 21,02769 |
| sp\|P49189\|AL9A1_HUMAN | sp\|P49189\|A | 8,3825 | 40894000 | 10 | 4 | 0,21981056 | -0,24303579 | 21,32147 | 21,40047 | 21,75516 | 21,56425 | 21,36212 | 21,62992 | 20,93352 | 21,14364 |
| sp\|Q58FF8\|H90B2_HUMAN | sp\|Q58FF8\|H | 2,5584 | 93121000 | 3 | 1 | 0,22147285 | -0,52075434 | 22,79741 | 22,72833 | 22,90718 | 22,82834 | 21,19093 | 22,90472 | 22,44083 | 22,64176 |
| tr\|H0YLA2\|H0YLA2_HUMAN;sp\|P37108\|SRP14_HUMAN | tr\|H0YLA2\|H | 9,1154 | 47662000 | 7 | 4 | 0,22215115 | -0,45265532 | 22,04263 | 21,43359 | 21,40966 | 22,60139 | 21,48944 | 21,79685 | 20,95992 | 21,43043 |
| sp\|P13804\|ETFA_HUMAN;tr\|H0YK49\|H0YK49_HUMAN;tr\|H0YLU7\|H0YLU7_HUMAN;tr\|H0YL1 | sp\|P13804\|E | 70,583 | 65227000 | 14 | 5 | 0,22491495 | -0,26713657 | 22,23359 | 22,33976 | 22,11613 | 22,58417 | 21,89126 | 22,3783 | 21,6497 | 22,28585 |
| sp\|Q9Y6M1\|IF2B2_HUMAN;tr\|F8W930\|F8W930_HUMAN | sp\|Q9Y6M1\|I | 24,342 | 25767000 | 12 | 6 | 0,22592891 | 0,23802058 | 20,66501 | NaN | 20,5696 | 20,53877 | 20,49473 | 20,84529 | 21,19244 | 20,78413 |
| tr\|B7Z9I1\|B7Z9I1_HUMAN;sp\|P11310\|ACADM_HUMAN;tr\|Q5T4U5\|Q5T4U5_HUMAN;tr\|H0YD | tr\|B7Z9I1\|B7 | 19,107 | 37091000 | 20 | 5 | 0,2274598 | 0,13929605 | 21,24284 | 21,08466 | 21,34846 | 21,01182 | 21,40499 | 21,19743 | 21,18441 | 21,45814 |
| sp\|Q14152\|EIF3A_HUMAN | sp\|Q14152\|E | 46,584 | 59342000 | 19 | 9 | 0,22895053 | -0,13958025 | 22,27373 | 21,9289 | 22,26196 | 22,10322 | 21,94321 | 21,86291 | 22,03463 | 22,16873 |
| tr\|F8W0J6\|F8W0J6_HUMAN;tr\|F5H4R6\|F5H4R6_HUMAN;tr\|H0YIV4\|H0YIV4_HUMAN;sp\|P552 | tr\|F8W0J6\|F8 | 94,01 | 128030000 | 22 | 7 | 0,22919886 | -0,09939766 | 22,95846 | 23,13002 | 23,15225 | 23,15019 | 23,06456 | 23,03825 | 22,82611 | 23,06441 |
| sp\|P25398\|RS12_HUMAN | sp\|P25398\|R | 106,42 | 270470000 | 23 | 4 | 0,2295568 | 0,28188467 | 23,8146 | 23,91169 | 23,65163 | 24,2717 | 24,63445 | 24,15677 | 23,83451 | 24,15143 |
| sp\|P01040\|CYTA_HUMAN;tr\|C9J0E4\|C9J0E4_HUMAN | sp\|P01040\|C | 4,8723 | 11773000 | 5 | 3 | 0,23050966 | -0,29877758 | 19,77001 | 19,97029 | 20,02648 | 19,51651 | 19,99466 | 19,09962 | 19,35767 | 19,63623 |
| sp\|Q9HD45\|TM9S3_HUMAN;tr\|Q5TB53\|Q5TB53_HUMAN | sp\|Q9HD45\|T | 4,4946 | 29787000 | 8 | 3 | 0,23119854 | -0,64063597 | 20,85855 | 20,80947 | 21,5048 | 21,29961 | 19,22055 | 20,68517 | 21,36057 | 20,64359 |
| sp\|P27694\|RFA1_HUMAN;tr\|I3L4R8\|I3L4R8_HUMAN;tr\|I3L524\|I3L524_HUMAN;tr\|I3L2M5\|I3 | sp\|P27694\|R | 32,133 | 73953000 | 24 | 10 | 0,23856193 | -0,29987049 | 22,42772 | 22,157 | 22,28836 | 22,35641 | 21,45608 | 21,94935 | 22,53258 | 22,092 |
| sp\|Q15233\|NONO_HUMAN;tr\|H7C367\|H7C367_HUMAN;tr\|C9IZL7\|C9IZL7_HUMAN;tr\|C9JYS8 | sp\|Q15233\|N | 173,2 | 444450000 | 27 | 12 | 0,23932759 | -0,23298073 | 25,02445 | 24,7599 | 24,99391 | 25,24329 | 24,51683 | 24,69216 | 24,67928 | 25,20135 |
| sp\|P51572\|BAP31_HUMAN;tr\|C9JSP1\|C9JSP1_HUMAN;tr\|C9JQ75\|C9JQ75_HUMAN;tr\|C9J0M | sp\|P51572\|B | 8,2766 | 237880000 | 5 | 5 | 0,23975269 | 0,22769403 | 23,82347 | 23,61133 | 23,64361 | 23,71314 | 23,81509 | 23,9478 | 24,37119 | 23,56825 |
| sp\|Q9BTV4\|TMM43_HUMAN | sp\|Q9BTV4\|T | 2,2229 | 13354000 | 5 | 2 | 0,24059599 | 0,43595521 | 19,85792 | 19,89318 | 19,88526 | 18,90177 | 19,97323 | 19,81209 | 20,42614 | NaN |
| sp\|Q9NP97\|DLRB1_HUMAN;tr\|B1AKR6\|B1AKR6_HUMAN;sp\|Q8TF09\|DLRB2_HUMAN;tr\|H3B( | sp\|Q9NP97\|D | 3,4589 | 7631000 | 8 | 2 | 0,2455295 | -0,26841418 | 18,96781 | NaN | 19,28834 | 19,12813 | 18,9221 | 19,12563 | 18,53131 | NaN |
| sp\|O14980\|XPO1_HUMAN;tr\|C9JF49\|C9JF49_HUMAN;tr\|C9IYM2\|C9IYM2_HUMAN;tr\|F8WF7 | sp\|O14980\|X | 98,799 | 136010000 | 18 | 10 | 0,24598663 | -0,07196045 | 22,99108 | 23,09779 | 23,02335 | 23,10266 | 22,90218 | 22,9581 | 22,94339 | 23,12337 |
| sp\|Q15637\|SF01_HUMAN;tr\|H7C561\|H7C561_HUMAN | sp\|Q15637\|S | 31,401 | 43695000 | 14 | 3 | 0,24768116 | -0,39262772 | 21,67349 | 21,50606 | 21,55754 | 21,79404 | 21,41916 | 21,82268 | 21,31639 | 20,40239 |

<table>
<tr><th></th><th></th><th></th><th></th><th></th><th></th><th></th><th></th><th></th><th></th><th></th><th></th><th></th><th></th><th></th><th></th></tr>
<tr><td>sp|P61254|RL26_HUMAN;tr|J3KTJ8|J3KTJ8_HUMAN;tr|J3QRI7|J3QRI7_HUMAN;tr|J3QQQ9|J3</td><td>sp|P61254|R</td><td>42,292</td><td>246150000</td><td>19</td><td>2</td><td>0,2484922</td><td>-0,5977397</td><td>24,41306</td><td>24,16469</td><td>24,1916</td><td>23,78097</td><td>23,87708</td><td>23,58875</td><td>24,40224</td><td>22,29129</td></tr>
<tr><td>tr|A0A0J9YXZ5|A0A0J9YXZ5_HUMAN;sp|P46940|IQGA1_HUMAN;tr|H0YKA5|H0YKA5_HUMAN</td><td>tr|A0A0J9YXZ</td><td>8,4215</td><td>22001000</td><td>11</td><td>5</td><td>0,25022245</td><td>-0,22628307</td><td>20,37708</td><td>20,62355</td><td>20,64095</td><td>20,70573</td><td>20,0021</td><td>20,16962</td><td>20,61171</td><td>20,65875</td></tr>
<tr><td>sp|Q92598|HS105_HUMAN;tr|A0A0A0MSM0|A0A0A0MSM0_HUMAN;tr|R4GN69|R4GN69_H</td><td>sp|Q92598|H</td><td>45,013</td><td>140530000</td><td>36</td><td>14</td><td>0,25454013</td><td>0,11903143</td><td>22,93418</td><td>23,25379</td><td>23,15957</td><td>23,12214</td><td>23,09957</td><td>23,15201</td><td>23,30966</td><td>23,38456</td></tr>
<tr><td>sp|P00491|PNPH_HUMAN;tr|G3V5M2|G3V5M2_HUMAN</td><td>sp|P00491|P</td><td>9,3734</td><td>73366000</td><td>10</td><td>4</td><td>0,25494684</td><td>-0,4996562</td><td>21,90176</td><td>22,02807</td><td>22,43916</td><td>22,52169</td><td>21,8196</td><td>22,72178</td><td>21,11407</td><td>21,2366</td></tr>
<tr><td>sp|P00338|LDHA_HUMAN;tr|F5GXY2|F5GXY2_HUMAN;tr|F5GXH2|F5GXH2_HUMAN;tr|F5GY</td><td>sp|P00338|L</td><td>114,06</td><td>1303200000</td><td>33</td><td>11</td><td>0,25524608</td><td>-0,22719145</td><td>26,69352</td><td>26,50139</td><td>26,12566</td><td>26,77316</td><td>26,16123</td><td>26,43743</td><td>26,06577</td><td>26,52054</td></tr>
<tr><td>sp|O15144|ARPC2_HUMAN;tr|H7C3F9|H7C3F9_HUMAN</td><td>sp|O15144|A</td><td>22,109</td><td>39130000</td><td>15</td><td>6</td><td>0,25694553</td><td>-0,22455215</td><td>21,31976</td><td>21,31324</td><td>21,59231</td><td>21,55112</td><td>20,81933</td><td>21,5545</td><td>21,10201</td><td>21,40239</td></tr>
<tr><td>sp|P13861|KAP2_HUMAN;tr|C9J830|C9J830_HUMAN;tr|H7C330|H7C330_HUMAN;tr|H7C1L0</td><td>sp|P13861|K</td><td>11,994</td><td>25870000</td><td>11</td><td>4</td><td>0,25877805</td><td>-0,14814329</td><td>20,69693</td><td>20,70759</td><td>21,05225</td><td>20,72692</td><td>20,78987</td><td>20,75148</td><td>20,62507</td><td>20,4247</td></tr>
<tr><td>sp|O15371|EIF3D_HUMAN;tr|B0QYA4|B0QYA4_HUMAN;tr|B0QYA5|B0QYA5_HUMAN</td><td>sp|O15371|E</td><td>5,7172</td><td>30278000</td><td>6</td><td>2</td><td>0,2621654</td><td>-0,17551359</td><td>NaN</td><td>21,08031</td><td>21,42183</td><td>21,36464</td><td>20,95943</td><td>21,24603</td><td>21,13477</td><td>NaN</td></tr>
<tr><td>tr|B1ANR0|B1ANR0_HUMAN;sp|Q13310|PABP4_HUMAN;tr|H0YCC8|H0YCC8_HUMAN;tr|B1A</td><td>tr|B1ANR0|B</td><td>43,615</td><td>236550000</td><td>40</td><td>1</td><td>0,26316812</td><td>0,14686394</td><td>23,7683</td><td>23,75746</td><td>24,08156</td><td>23,62165</td><td>23,84422</td><td>23,84134</td><td>24,00813</td><td>24,12274</td></tr>
<tr><td>sp|O43390|HNRPR_HUMAN;tr|B4DT28|B4DT28_HUMAN</td><td>sp|O43390|H</td><td>12,664</td><td>46494000</td><td>19</td><td>6</td><td>0,26413665</td><td>-0,34801197</td><td>21,29967</td><td>21,66999</td><td>21,87433</td><td>21,45453</td><td>20,71431</td><td>21,5029</td><td>21,79317</td><td>20,89608</td></tr>
<tr><td>sp|Q99848|EBP2_HUMAN;tr|H7C2Q8|H7C2Q8_HUMAN</td><td>sp|Q99848|E</td><td>4,3562</td><td>15765000</td><td>7</td><td>3</td><td>0,26474014</td><td>-0,32326508</td><td>NaN</td><td>20,04166</td><td>20,3868</td><td>20,46473</td><td>NaN</td><td>19,72807</td><td>20,39843</td><td>19,7969</td></tr>
<tr><td>sp|P61026|RAB10_HUMAN;tr|H0YLJ8|H0YLJ8_HUMAN;tr|H0YL94|H0YL94_HUMAN;sp|P5919</td><td>sp|P61026|R</td><td>26,887</td><td>68818000</td><td>15</td><td>4</td><td>0,26835525</td><td>-0,14071608</td><td>22,04966</td><td>22,4852</td><td>22,27946</td><td>22,44998</td><td>22,17916</td><td>22,29734</td><td>22,20753</td><td>22,0174</td></tr>
<tr><td>sp|P52272|HNRPM_HUMAN;tr|A0A087X0X3|A0A087X0X3_HUMAN;tr|M0R019|M0R019_HU</td><td>sp|P52272|H</td><td>198,67</td><td>635150000</td><td>37</td><td>14</td><td>0,27644635</td><td>0,21553898</td><td>25,00573</td><td>25,22295</td><td>25,3679</td><td>24,85885</td><td>25,20154</td><td>25,32158</td><td>25,72151</td><td>25,07296</td></tr>
<tr><td>sp|Q14697|GANAB_HUMAN;tr|F5H6X6|F5H6X6_HUMAN;tr|E9PKU7|E9PKU7_HUMAN;tr|E9P</td><td>sp|Q14697|G</td><td>323,31</td><td>494650000</td><td>59</td><td>18</td><td>0,27742044</td><td>-0,1255908</td><td>24,97338</td><td>25,1189</td><td>25,1949</td><td>24,95753</td><td>24,8551</td><td>24,79248</td><td>24,90408</td><td>25,19069</td></tr>
<tr><td>sp|P55084|ECHB_HUMAN;tr|F5GZQ3|F5GZQ3_HUMAN;tr|B5MD38|B5MD38_HUMAN;tr|C9J</td><td>sp|P55084|E</td><td>7,903</td><td>38747000</td><td>11</td><td>5</td><td>0,2808212</td><td>-0,30711953</td><td>22,20323</td><td>21,48296</td><td>21,36672</td><td>21,51648</td><td>21,39147</td><td>21,54882</td><td>21,0654</td><td>NaN</td></tr>
<tr><td>sp|P61956|SUMO2_HUMAN;sp|Q6EEV6|SUMO4_HUMAN;sp|P55854|SUMO3_HUMAN;tr|A8</td><td>sp|P61956|S</td><td>10,697</td><td>105470000</td><td>9</td><td>2</td><td>0,28121828</td><td>-0,32252105</td><td>NaN</td><td>22,72293</td><td>23,00154</td><td>23,16082</td><td>22,72299</td><td>23,15023</td><td>22,15718</td><td>22,52657</td></tr>
<tr><td>sp|Q9HB71|CYBP_HUMAN;tr|B2ZWH1|B2ZWH1_HUMAN</td><td>sp|Q9HB71|</td><td>37,96</td><td>55912000</td><td>9</td><td>4</td><td>0,28379408</td><td>0,14752913</td><td>21,98109</td><td>21,67784</td><td>21,5595</td><td>21,82089</td><td>22,065</td><td>21,95978</td><td>21,66206</td><td>21,9426</td></tr>
<tr><td>sp|P62249|RS16_HUMAN;tr|M0R210|M0R210_HUMAN;tr|A0A087WZ27|A0A087WZ27_HUM</td><td>sp|P62249|R</td><td>158,91</td><td>1631000000</td><td>50</td><td>8</td><td>0,2862231</td><td>0,27242517</td><td>26,26209</td><td>26,83219</td><td>26,73518</td><td>26,11333</td><td>27,07286</td><td>26,93955</td><td>26,61989</td><td>26,4002</td></tr>
<tr><td>sp|P18669|PGAM1_HUMAN;sp|P15259|PGAM2_HUMAN;sp|Q8N0Y7|PGAM4_HUMAN</td><td>sp|P18669|P</td><td>26,118</td><td>94667000</td><td>11</td><td>4</td><td>0,29036196</td><td>-0,23536444</td><td>22,77509</td><td>22,65823</td><td>22,81617</td><td>23,02862</td><td>22,43145</td><td>22,88069</td><td>22,12414</td><td>22,90036</td></tr>
<tr><td>tr|A0A0A0MSJ0|A0A0A0MSJ0_HUMAN;sp|Q86XP3|DDX42_HUMAN;tr|J3KRE3|J3KRE3_HUMA</td><td>tr|A0A0A0M</td><td>4,1091</td><td>12491000</td><td>3</td><td>3</td><td>0,29064308</td><td>-0,11228466</td><td>19,74319</td><td>19,78705</td><td>19,76633</td><td>19,87938</td><td>19,67984</td><td>19,47508</td><td>19,64883</td><td>19,92305</td></tr>
<tr><td>tr|F8WDD7|F8WDD7_HUMAN;sp|P59998|ARPC4_HUMAN;tr|F8WCF6|F8WCF6_HUMAN;tr|A</td><td>tr|F8WDD7|F</td><td>3,2186</td><td>29835000</td><td>7</td><td>3</td><td>0,29376031</td><td>0,73793443</td><td>21,34408</td><td>19,88341</td><td>19,75285</td><td>NaN</td><td>21,66787</td><td>21,31341</td><td>21,36164</td><td>19,91595</td></tr>
<tr><td>tr|J3KN66|J3KN66_HUMAN;tr|A0A0A0MSK5|A0A0A0MSK5_HUMAN;sp|Q5JTV8|TOIP1_HUM</td><td>tr|J3KN66|J3</td><td>3,6801</td><td>17080000</td><td>5</td><td>4</td><td>0,29452134</td><td>0,25689888</td><td>NaN</td><td>20,23327</td><td>19,79927</td><td>20,33123</td><td>20,60738</td><td>20,1342</td><td>NaN</td><td>20,39289</td></tr>
<tr><td>sp|Q8WWM7|ATX2L_HUMAN;tr|H3BUF6|H3BUF6_HUMAN;tr|H3BSK9|H3BSK9_HUMAN</td><td>sp|Q8WWM7</td><td>5,0985</td><td>31274000</td><td>9</td><td>4</td><td>0,29630579</td><td>0,32029867</td><td>21,15221</td><td>20,72267</td><td>20,14967</td><td>20,83854</td><td>21,24045</td><td>20,4804</td><td>21,17426</td><td>21,24916</td></tr>
<tr><td>sp|P20700|LMNB1_HUMAN;tr|E9PBF6|E9PBF6_HUMAN;tr|A0A0D9SFE5|A0A0D9SFE5_HUMA</td><td>sp|P20700|L</td><td>80,849</td><td>445540000</td><td>54</td><td>19</td><td>0,29846155</td><td>0,45529795</td><td>24,32971</td><td>23,90242</td><td>24,12755</td><td>24,57906</td><td>23,92877</td><td>25,71487</td><td>24,6142</td><td>24,50209</td></tr>
<tr><td>sp|Q15029|U5S1_HUMAN</td><td>sp|Q15029|U</td><td>19,558</td><td>63837000</td><td>23</td><td>8</td><td>0,30163424</td><td>0,12923574</td><td>21,97495</td><td>21,8711</td><td>21,9441</td><td>21,87845</td><td>21,77961</td><td>21,94621</td><td>22,21045</td><td>22,24928</td></tr>
<tr><td>sp|P21796|VDAC1_HUMAN;tr|C9JI87|C9JI87_HUMAN</td><td>sp|P21796|V</td><td>6,2642</td><td>22092000</td><td>6</td><td>3</td><td>0,30304258</td><td>0,43690157</td><td>19,55127</td><td>19,64484</td><td>19,66622</td><td>20,02134</td><td>19,92093</td><td>21,21957</td><td>20,02296</td><td>19,46782</td></tr>
<tr><td>sp|Q9Y678|COPG1_HUMAN;tr|H0Y8X7|H0Y8X7_HUMAN</td><td>sp|Q9Y678|C</td><td>9,5376</td><td>20421000</td><td>11</td><td>4</td><td>0,30337307</td><td>-0,25499868</td><td>20,43145</td><td>20,3721</td><td>20,47773</td><td>20,69753</td><td>20,14892</td><td>20,8233</td><td>20,19976</td><td>19,78683</td></tr>
<tr><td>tr|A0A0C4DGS1|A0A0C4DGS1_HUMAN;sp|P39656|OST48_HUMAN;tr|U3KQ84|U3KQ84_HUM</td><td>tr|A0A0C4DG</td><td>18,763</td><td>94076000</td><td>29</td><td>8</td><td>0,30342991</td><td>0,15580225</td><td>22,48508</td><td>22,2856</td><td>22,63285</td><td>22,61385</td><td>22,39482</td><td>22,60865</td><td>22,69598</td><td>22,94113</td></tr>
<tr><td>sp|Q15005|SPCS2_HUMAN;tr|A0A087WUC6|A0A087WUC6_HUMAN;tr|E9PI68|E9PI68_HUM</td><td>sp|Q15005|S</td><td>3,7288</td><td>17229000</td><td>4</td><td>2</td><td>0,30597664</td><td>0,38098796</td><td>NaN</td><td>20,10815</td><td>19,92929</td><td>20,07509</td><td>21,19292</td><td>19,88445</td><td>20,41142</td><td>20,18519</td></tr>
<tr><td>tr|A0A0U1RQH7|A0A0U1RQH7_HUMAN;tr|H0Y4X3|H0Y4X3_HUMAN;tr|G3XAC6|G3XAC6_HU</td><td>tr|A0A0U1RC</td><td>32,333</td><td>40157000</td><td>12</td><td>5</td><td>0,31028886</td><td>-0,45032263</td><td>21,61526</td><td>21,4079</td><td>20,56347</td><td>22,4986</td><td>21,13621</td><td>21,02608</td><td>21,26264</td><td>20,85901</td></tr>
<tr><td>sp|P38606|VATA_HUMAN;tr|C9JA17|C9JA17_HUMAN;tr|C9JVW8|C9JVW8_HUMAN</td><td>sp|P38606|V</td><td>11,257</td><td>38875000</td><td>14</td><td>7</td><td>0,31521889</td><td>0,13559771</td><td>21,61275</td><td>21,20652</td><td>21,08219</td><td>21,3274</td><td>21,47353</td><td>21,30011</td><td>21,46996</td><td>21,52765</td></tr>
<tr><td>sp|P21281|VATB2_HUMAN;tr|H0YC04|H0YC04_HUMAN;tr|C9JL73|C9JL73_HUMAN;sp|P1531</td><td>sp|P21281|V</td><td>3,693</td><td>7152300</td><td>7</td><td>2</td><td>0,31972039</td><td>-0,16228819</td><td>18,97958</td><td>18,90745</td><td>19,02443</td><td>19,05973</td><td>18,89457</td><td>18,83556</td><td>18,44298</td><td>19,14892</td></tr>
<tr><td>sp|P60953|CDC42_HUMAN;tr|Q5JYX0|Q5JYX0_HUMAN</td><td>sp|P60953|C</td><td>85,291</td><td>28187000</td><td>10</td><td>3</td><td>0,3198627</td><td>0,64153147</td><td>19,94192</td><td>19,60206</td><td>20,07744</td><td>19,93531</td><td>20,05854</td><td>22,24519</td><td>19,64146</td><td>20,17767</td></tr>
<tr><td>tr|K7EJC1|K7EJC1_HUMAN;tr|K7EJR3|K7EJR3_HUMAN;tr|R4GMR5|R4GMR5_HUMAN;sp|P48</td><td>tr|K7EJC1|K7</td><td>3,8714</td><td>49682000</td><td>8</td><td>3</td><td>0,32208442</td><td>-0,1616621</td><td>21,85962</td><td>21,89504</td><td>21,82889</td><td>21,55225</td><td>21,80365</td><td>21,80664</td><td>21,2635</td><td>21,61535</td></tr>
<tr><td>sp|P62805|H4_HUMAN</td><td>sp|P62805|H</td><td>322,23</td><td>3232700000</td><td>41</td><td>9</td><td>0,32843309</td><td>0,92654943</td><td>25,93159</td><td>25,05001</td><td>27,38105</td><td>27,92603</td><td>25,91593</td><td>27,99814</td><td>28,56518</td><td>27,51563</td></tr>
<tr><td>sp|O95373|IPO7_HUMAN</td><td>sp|O95373|I</td><td>31,034</td><td>72663000</td><td>22</td><td>7</td><td>0,33486215</td><td>-0,27458477</td><td>22,24826</td><td>22,2252</td><td>22,35329</td><td>22,41612</td><td>21,27748</td><td>22,17234</td><td>22,2723</td><td>22,42242</td></tr>
<tr><td>sp|P51398|RT29_HUMAN;tr|V9GYL9|V9GYL9_HUMAN;tr|V9GYF7|V9GYF7_HUMAN;tr|V9GZ6</td><td>sp|P51398|R</td><td>9,0526</td><td>15504000</td><td>9</td><td>4</td><td>0,33667055</td><td>0,17797184</td><td>19,65979</td><td>19,79532</td><td>19,8624</td><td>19,85641</td><td>20,18628</td><td>19,67735</td><td>20,31628</td><td>19,7059</td></tr>
<tr><td>sp|Q9UN86|G3BP2_HUMAN</td><td>sp|Q9UN86|</td><td>9,1773</td><td>54517000</td><td>10</td><td>2</td><td>0,33715839</td><td>0,2196784</td><td>21,75022</td><td>21,51031</td><td>21,84157</td><td>21,27219</td><td>21,45643</td><td>21,86219</td><td>22,24916</td><td>21,68521</td></tr>
<tr><td>sp|P09874|PARP1_HUMAN;tr|Q5VX84|Q5VX84_HUMAN;tr|Q5VX85|Q5VX85_HUMAN</td><td>sp|P09874|P</td><td>323,31</td><td>588440000</td><td>79</td><td>25</td><td>0,3383473</td><td>-0,14956188</td><td>25,32663</td><td>25,32244</td><td>25,32543</td><td>25,28602</td><td>24,97176</td><td>24,91315</td><td>25,23685</td><td>25,54053</td></tr>
<tr><td>sp|Q86V81|THOC4_HUMAN;tr|E9PB61|E9PB61_HUMAN</td><td>sp|Q86V81|T</td><td>111,95</td><td>83532000</td><td>15</td><td>4</td><td>0,34206675</td><td>0,23822069</td><td>22,21617</td><td>21,46234</td><td>22,36586</td><td>22,1717</td><td>22,42355</td><td>22,5087</td><td>22,00014</td><td>22,23658</td></tr>
<tr><td>sp|Q92688|AN32B_HUMAN</td><td>sp|Q92688|A</td><td>5,7779</td><td>70821000</td><td>7</td><td>3</td><td>0,34214004</td><td>-0,31714964</td><td>22,19547</td><td>22,32808</td><td>22,42406</td><td>22,93776</td><td>21,90205</td><td>22,71123</td><td>NaN</td><td>21,84931</td></tr>
<tr><td>tr|A0A0B4J1Z1|A0A0B4J1Z1_HUMAN;tr|C9JAB2|C9JAB2_HUMAN;sp|Q16629|SRSF7_HUMAN</td><td>tr|A0A0B4J1Z</td><td>40,601</td><td>67688000</td><td>11</td><td>3</td><td>0,34316575</td><td>0,45636384</td><td>22,01509</td><td>20,92492</td><td>22,42627</td><td>22,00206</td><td>NaN</td><td>22,18882</td><td>22,78531</td><td>21,92122</td></tr>
<tr><td>sp|Q16777|H2A2C_HUMAN;sp|Q6FI13|H2A2A_HUMAN</td><td>sp|Q16777|H</td><td>323,31</td><td>2491900000</td><td>27</td><td>1</td><td>0,34458425</td><td>0,84839821</td><td>25,74491</td><td>25,18079</td><td>27,06332</td><td>27,22803</td><td>25,33214</td><td>27,51758</td><td>28,47127</td><td>27,28965</td></tr>
<tr><td>sp|P50454|SERPH_HUMAN;tr|E9PJH8|E9PJH8_HUMAN;tr|E9PIG2|E9PIG2_HUMAN;tr|E9PRS3</td><td>sp|P50454|S</td><td>6,1155</td><td>17399000</td><td>3</td><td>3</td><td>0,35691633</td><td>0,11546993</td><td>20,12351</td><td>20,12931</td><td>19,86582</td><td>20,32893</td><td>20,39184</td><td>20,14445</td><td>20,09893</td><td>20,27424</td></tr>
<tr><td>sp|Q5SRE5|NU188_HUMAN</td><td>sp|Q5SRE5|N</td><td>1,9449</td><td>6438000</td><td>3</td><td>3</td><td>0,35807135</td><td>-0,13048172</td><td>18,87781</td><td>18,75012</td><td>18,92187</td><td>18,78302</td><td>18,5713</td><td>18,48129</td><td>18,70819</td><td>19,05011</td></tr>
<tr><td>sp|Q01813|PFKAP_HUMAN;tr|V9GY25|V9GY25_HUMAN;tr|B1APP6|B1APP6_HUMAN;tr|B1A</td><td>sp|Q01813|P</td><td>4,994</td><td>58581000</td><td>8</td><td>2</td><td>0,36096174</td><td>-0,12445545</td><td>21,6657</td><td>22,05043</td><td>22,05295</td><td>22,09388</td><td>22,06194</td><td>21,81917</td><td>21,74456</td><td>21,73946</td></tr>
<tr><td>sp|Q96AG4|LRC59_HUMAN;tr|I3L223|I3L223_HUMAN</td><td>sp|Q96AG4|L</td><td>155,83</td><td>218760000</td><td>26</td><td>7</td><td>0,36227134</td><td>0,15451956</td><td>23,41828</td><td>23,78347</td><td>23,9393</td><td>23,51461</td><td>23,99162</td><td>23,94842</td><td>23,78766</td><td>23,54604</td></tr>
<tr><td>tr|A0A0A0MTN3|A0A0A0MTN3_HUMAN;sp|P21266|GSTM3_HUMAN;tr|E9PLF1|E9PLF1_HUM</td><td>tr|A0A0A0MT</td><td>5,9298</td><td>19197000</td><td>6</td><td>2</td><td>0,36325052</td><td>0,33429019</td><td>20,6856</td><td>20,51557</td><td>20,43777</td><td>20,48227</td><td>20,27889</td><td>21,60923</td><td>20,70565</td><td>NaN</td></tr>
<tr><td>tr|E9PID8|E9PID8_HUMAN;sp|P33240|CSTF2_HUMAN;tr|E7EWR4|E7EWR4_HUMAN;sp|Q9H</td><td>tr|E9PID8|E9</td><td>4,1863</td><td>17073000</td><td>5</td><td>2</td><td>0,3644673</td><td>-0,0767142</td><td>20,25026</td><td>20,26957</td><td>20,40748</td><td>20,22847</td><td>20,28681</td><td>20,06815</td><td>20,28173</td><td>NaN</td></tr>
<tr><td>sp|O95347|SMC2_HUMAN</td><td>sp|O95347|S</td><td>7,2129</td><td>28674000</td><td>8</td><td>6</td><td>0,36754493</td><td>-0,10429239</td><td>20,9885</td><td>21,16183</td><td>20,97771</td><td>20,7128</td><td>20,71011</td><td>20,96874</td><td>20,87079</td><td>20,87403</td></tr>
<tr><td>sp|P08238|HS90B_HUMAN;sp|Q58FF7|H90B3_HUMAN</td><td>sp|P08238|H</td><td>323,31</td><td>4256900000</td><td>152</td><td>13</td><td>0,36845341</td><td>-0,12559986</td><td>28,15822</td><td>28,12169</td><td>28,31592</td><td>28,24082</td><td>28,03318</td><td>28,05533</td><td>27,83032</td><td>28,41542</td></tr>
<tr><td>sp|P49755|TMEDA_HUMAN;tr|G3V2K7|G3V2K7_HUMAN</td><td>sp|P49755|T</td><td>20,992</td><td>69604000</td><td>12</td><td>5</td><td>0,3695902</td><td>-0,10311031</td><td>22,06145</td><td>22,16002</td><td>22,38184</td><td>22,02181</td><td>22,05165</td><td>22,14127</td><td>22,1618</td><td>21,85795</td></tr>
<tr><td>sp|Q96CX2|KCD12_HUMAN;sp|Q68DU8|KCD16_HUMAN</td><td>sp|Q96CX2|K</td><td>37,436</td><td>39727000</td><td>15</td><td>6</td><td>0,37353564</td><td>-0,07158899</td><td>21,31689</td><td>21,44673</td><td>21,27804</td><td>21,40681</td><td>21,27071</td><td>21,27923</td><td>21,15233</td><td>21,45984</td></tr>
<tr><td>sp|P31153|METK2_HUMAN;sp|Q00266|METK1_HUMAN</td><td>sp|P31153|M</td><td>21,773</td><td>107640000</td><td>19</td><td>6</td><td>0,37363819</td><td>-0,2018981</td><td>22,84573</td><td>23,0012</td><td>22,89992</td><td>23,13127</td><td>22,88327</td><td>22,96104</td><td>22,17509</td><td>23,05114</td></tr>
<tr><td>sp|P62937|PPIA_HUMAN;tr|F8WE65|F8WE65_HUMAN;tr|C9J5S7|C9J5S7_HUMAN;tr|E5RIZ5|</td><td>sp|P62937|P</td><td>323,31</td><td>644280000</td><td>33</td><td>7</td><td>0,37387399</td><td>-0,42782068</td><td>25,33412</td><td>25,29747</td><td>25,41783</td><td>25,83178</td><td>24,72765</td><td>25,10221</td><td>24,15429</td><td>26,18577</td></tr>
<tr><td>tr|A0A024RCR6|A0A024RCR6_HUMAN;tr|A0A0G2JK23|A0A0G2JK23_HUMAN;sp|P46379|BAG</td><td>tr|A0A024RC</td><td>26,116</td><td>32358000</td><td>19</td><td>5</td><td>0,37425469</td><td>-0,3959136</td><td>20,62952</td><td>21,4224</td><td>20,63901</td><td>22,24771</td><td>21,12584</td><td>20,46204</td><td>20,75849</td><td>21,00861</td></tr>
<tr><td>sp|Q9Y3U8|RL36_HUMAN;tr|J3QSB5|J3QSB5_HUMAN;tr|J3KTD3|J3KTD3_HUMAN</td><td>sp|Q9Y3U8|F</td><td>12,636</td><td>88961000</td><td>12</td><td>4</td><td>0,38435316</td><td>-0,41551399</td><td>21,48015</td><td>23,48591</td><td>23,0806</td><td>22,69013</td><td>22,54571</td><td>22,177</td><td>22,14622</td><td>22,2058</td></tr>
<tr><td>sp|P61221|ABCE1_HUMAN;tr|D6R9I9|D6R9I9_HUMAN;tr|D6RGF4|D6RGF4_HUMAN;tr|H0Y9</td><td>sp|P61221|A</td><td>37,584</td><td>143890000</td><td>28</td><td>12</td><td>0,38481038</td><td>-0,19151974</td><td>23,3181</td><td>23,48971</td><td>23,31948</td><td>23,37981</td><td>23,02753</td><td>23,13955</td><td>22,82118</td><td>23,75277</td></tr>
<tr><td>sp|P30084|ECHM_HUMAN</td><td>sp|P30084|E</td><td>33,718</td><td>57076000</td><td>15</td><td>4</td><td>0,38504904</td><td>-0,16725779</td><td>21,70134</td><td>21,87811</td><td>21,85011</td><td>22,29393</td><td>21,78161</td><td>21,89182</td><td>21,4065</td><td>21,97453</td></tr>
<tr><td>sp|Q00796|DHSO_HUMAN;tr|H0YLA4|H0YLA4_HUMAN;tr|H0YKB3|H0YKB3_HUMAN</td><td>sp|Q00796|D</td><td>4,1062</td><td>27191000</td><td>10</td><td>3</td><td>0,38632788</td><td>-0,30461693</td><td>20,63112</td><td>20,95851</td><td>21,43695</td><td>21,53758</td><td>20,7102</td><td>21,56983</td><td>20,53479</td><td>20,53089</td></tr>
<tr><td>sp|Q9BRA2|TXD17_HUMAN;tr|I3L0K2|I3L0K2_HUMAN;tr|I3L3M7|I3L3M7_HUMAN;tr|I3L2R6</td><td>sp|Q9BRA2|T</td><td>5,1843</td><td>17722000</td><td>3</td><td>3</td><td>0,38770963</td><td>-0,19406541</td><td>20,27912</td><td>20,31606</td><td>20,3739</td><td>NaN</td><td>20,06026</td><td>20,57034</td><td>19,7336</td><td>20,15165</td></tr>
<tr><td>tr|H0YL72|H0YL72_HUMAN;sp|P50213|IDH3A_HUMAN;tr|H0YMU3|H0YMU3_HUMAN;tr|H0</td><td>tr|H0YL72|H</td><td>7,4602</td><td>87143000</td><td>7</td><td>6</td><td>0,38947229</td><td>0,41639694</td><td>21,67211</td><td>21,95369</td><td>22,54402</td><td>22,5612</td><td>23,15971</td><td>22,87418</td><td>NaN</td><td>21,76357</td></tr>
<tr><td>sp|P40939|ECHA_HUMAN;tr|H0YFD6|H0YFD6_HUMAN</td><td>sp|P40939|E</td><td>15,19</td><td>54436000</td><td>21</td><td>7</td><td>0,40387191</td><td>-0,15970707</td><td>21,80298</td><td>21,54213</td><td>22,15106</td><td>21,80077</td><td>21,54279</td><td>21,79606</td><td>21,9426</td><td>21,37666</td></tr>
<tr><td>tr|K7EQQ8|K7EQQ8_HUMAN;sp|Q9NXH9|TRM1_HUMAN;tr|K7EQU7|K7EQU7_HUMAN;tr|K7</td><td>tr|K7EQQ8|K</td><td>3,0056</td><td>6840800</td><td>4</td><td>3</td><td>0,40990098</td><td>0,16286659</td><td>18,56253</td><td>NaN</td><td>18,75411</td><td>18,86876</td><td>18,82208</td><td>19,18403</td><td>18,66789</td><td>NaN</td></tr>
<tr><td>tr|F2Z2V0|F2Z2V0_HUMAN;sp|Q99829|CPNE1_HUMAN;tr|B0QZ18|B0QZ18_HUMAN;tr|A6P</td><td>tr|F2Z2V0|F</td><td>46,997</td><td>31837000</td><td>15</td><td>5</td><td>0,41032643</td><td>0,12018204</td><td>21,10169</td><td>20,85049</td><td>20,90529</td><td>21,29569</td><td>20,98029</td><td>21,39367</td><td>21,06112</td><td>21,1988</td></tr>
<tr><td>sp|P08243|ASNS_HUMAN;tr|F8WEJ5|F8WEJ5_HUMAN;tr|C9J057|C9J057_HUMAN</td><td>sp|P08243|A</td><td>9,1305</td><td>46887000</td><td>9</td><td>5</td><td>0,41059825</td><td>-0,23399878</td><td>21,40484</td><td>21,62649</td><td>21,51917</td><td>22,37132</td><td>21,5657</td><td>21,50257</td><td>21,42111</td><td>NaN</td></tr>
<tr><td>tr|A0A096LNY6|A0A096LNY6_HUMAN;sp|P30566|PUR8_HUMAN;tr|A0A0A6YY92|A0A0A6YY9</td><td>tr|A0A096LN</td><td>3,2346</td><td>18991000</td><td>6</td><td>2</td><td>0,41113468</td><td>-0,08750407</td><td>20,59113</td><td>20,53592</td><td>20,65875</td><td>20,49843</td><td>20,54821</td><td>20,27685</td><td>NaN</td><td>20,6256</td></tr>
<tr><td>sp|P35606|COPB2_HUMAN;tr|D6R997|D6R997_HUMAN;tr|D6RBZ7|D6RBZ7_HUMAN;tr|D6R</td><td>sp|P35606|C</td><td>21,204</td><td>33037000</td><td>12</td><td>8</td><td>0,4113377</td><td>-0,1376214</td><td>21,21175</td><td>21,17286</td><td>21,18519</td><td>20,96522</td><td>21,08661</td><td>20,56449</td><td>21,15468</td><td>21,17877</td></tr>
<tr><td>sp|Q15021|CND1_HUMAN;tr|E7EN77|E7EN77_HUMAN</td><td>sp|Q15021|C</td><td>5,3087</td><td>8809600</td><td>4</td><td>3</td><td>0,4118117</td><td>-0,24851863</td><td>NaN</td><td>19,56657</td><td>19,65571</td><td>18,84252</td><td>NaN</td><td>18,94241</td><td>19,14298</td><td>19,23386</td></tr>
</table>

| | | | | | | | | | | | | | | | |
|---|---|---|---|---|---|---|---|---|---|---|---|---|---|---|---|
| sp\|P49006\|MRP_HUMAN | sp\|P49006\|M | 27,905 | 29785000 | 7 | 2 | 0,41585255 | -0,23492304 | 21,10604 | 20,97715 | NaN | 21,0426 | 20,84069 | 21,23333 | 20,347 | NaN |
| sp\|Q8N257\|H2B3B_HUMAN;sp\|Q16778\|H2B2E_HUMAN;sp\|P33778\|H2B1B_HUMAN;sp\|P235 | sp\|Q8N257\|H | 321,21 | 1781800000 | 23 | 2 | 0,41951206 | 0,8823185 | 24,97058 | 24,073 | 26,73609 | 26,99118 | 24,63323 | 26,8992 | 28,20191 | 26,56578 |
| sp\|P48444\|COPD_HUMAN;tr\|B0YIW6\|B0YIW6_HUMAN;tr\|Q6P1Q5\|Q6P1Q5_HUMAN;tr\|E9PK | sp\|P48444\|C | 18,195 | 41728000 | 12 | 4 | 0,42436217 | -0,06409168 | 21,27469 | 21,53711 | 21,45623 | 21,40322 | 21,40265 | 21,30536 | 21,46742 | 21,23946 |
| sp\|O75533\|SF3B1_HUMAN;tr\|H7C341\|H7C341_HUMAN | sp\|O75533\|S | 71,404 | 99284000 | 22 | 9 | 0,43031619 | -0,25784922 | 22,88373 | 22,80554 | 22,73411 | 22,61113 | 21,66891 | 22,46107 | 22,89946 | 22,97369 |
| sp\|P42166\|LAP2A_HUMAN | sp\|P42166\|L | 44,134 | 178700000 | 48 | 10 | 0,43482211 | -0,2349844 | 23,29866 | 23,29642 | 23,75929 | 23,76415 | 23,34394 | 23,19831 | 23,91743 | 22,7189 |
| sp\|Q9BSJ8\|ESYT1_HUMAN;tr\|F8VZB1\|F8VZB1_HUMAN | sp\|Q9BSJ8\|E | 14,69 | 34666000 | 22 | 5 | 0,43817812 | -0,07815981 | 21,35692 | 21,17804 | 21,15851 | 21,16484 | 21,05484 | 20,94999 | 21,24034 | 21,3005 |
| sp\|Q9UHB9\|SRP68_HUMAN | sp\|Q9UHB9\| | 23,709 | 34451000 | 12 | 6 | 0,4385765 | -0,07146263 | 21,22117 | 21,14892 | 21,29182 | 21,33615 | 21,18235 | 21,38785 | 21,04019 | 21,10181 |
| sp\|P55072\|TERA_HUMAN;tr\|C9JUP7\|C9JUP7_HUMAN;tr\|C9IZA5\|C9IZA5_HUMAN | sp\|P55072\|T | 41,414 | 258990000 | 52 | 15 | 0,44006306 | -0,06981325 | 23,99981 | 23,93517 | 24,2522 | 24,08229 | 23,87998 | 24,08869 | 23,95296 | 24,06858 |
| sp\|O95436\|NPT2B_HUMAN | sp\|O95436\|N | 1,626 | 81232000 | 5 | 2 | 0,44460138 | -0,25023985 | 23,00703 | 22,91614 | 21,75307 | 22,59798 | 22,5078 | 22,48311 | 22,0459 | 22,23646 |
| sp\|P63000\|RAC1_HUMAN;sp\|P60763\|RAC3_HUMAN;sp\|P15153\|RAC2_HUMAN;tr\|J3KSC4\|J3 | sp\|P63000\|R | 2,5272 | 50325000 | 7 | 3 | 0,44501891 | 0,52294525 | 20,14669 | 20,8064 | 21,73715 | 21,10495 | 20,9686 | 22,65176 | 20,79487 | NaN |
| tr\|J9JID7\|J9JID7_HUMAN;sp\|Q03252\|LMNB2_HUMAN | tr\|J9JID7\|J9J | 17,799 | 74198000 | 15 | 8 | 0,45109064 | 0,29917192 | 21,82877 | 21,74053 | 22,07633 | 21,90367 | 21,46832 | 23,10537 | 22,41235 | 21,75996 |
| sp\|P04844\|RPN2_HUMAN;tr\|Q5JYR7\|Q5JYR7_HUMAN;tr\|F2Z3K5\|F2Z3K5_HUMAN;tr\|Q5JYR4 | sp\|P04844\|R | 190,49 | 132650000 | 36 | 12 | 0,45475998 | -0,10420609 | 23,04006 | 22,92922 | 23,32045 | 23,14331 | 22,96534 | 22,84378 | 23,29614 | 22,91095 |
| sp\|O76094\|SRP72_HUMAN;tr\|R4GNC1\|R4GNC1_HUMAN;tr\|D6RDY6\|D6RDY6_HUMAN | sp\|O76094\|S | 11,37 | 25793000 | 9 | 5 | 0,45793915 | -0,20364507 | 21,28167 | 20,48572 | 21,13659 | NaN | NaN | 20,68243 | 20,80719 | 20,80341 |
| sp\|P11387\|TOP1_HUMAN;tr\|E5RJ95\|E5RJ95_HUMAN;tr\|E5RFS0\|E5RFS0_HUMAN;tr\|E5RIC7\|I | sp\|P11387\|T | 13,023 | 43731000 | 7 | 4 | 0,46299284 | 0,2031827 | 21,47526 | 21,06881 | 20,96437 | 21,68282 | 21,12231 | 21,32075 | 22,03644 | 21,52449 |
| sp\|P07195\|LDHB_HUMAN;tr\|A8MW50\|A8MW50_HUMAN;tr\|F5H793\|F5H793_HUMAN;tr\|C9. | sp\|P07195\|L | 323,31 | 1487400000 | 50 | 9 | 0,46365622 | -0,13263178 | 27,00185 | 26,53614 | 26,53906 | 26,79266 | 26,54822 | 26,71751 | 26,24525 | 26,8282 |
| sp\|O43175\|SERA_HUMAN;tr\|Q5SZU1\|Q5SZU1_HUMAN | sp\|O43175\|S | 30,501 | 299510000 | 39 | 12 | 0,4710846 | -0,12537861 | 24,22619 | 24,39126 | 24,3123 | 24,52103 | 24,20464 | 24,21823 | 23,89634 | 24,63006 |
| sp\|P35250\|RFC2_HUMAN;tr\|H7C5P4\|H7C5P4_HUMAN;tr\|A0A087WVY3\|A0A087WVY3_HUM | sp\|P35250\|R | 4,7165 | 20837000 | 8 | 4 | 0,47307405 | 0,0928731 | 20,23585 | 20,46533 | 20,45393 | 20,57191 | 20,68286 | 20,26511 | 20,47645 | 20,67409 |
| sp\|Q14974\|IMB1_HUMAN;tr\|J3KTM9\|J3KTM9_HUMAN;tr\|J3QR48\|J3QR48_HUMAN;tr\|J3QKC | sp\|Q14974\|I | 323,31 | 414920000 | 54 | 16 | 0,47679676 | 0,12569046 | 24,79615 | 24,61341 | 24,64548 | 24,67357 | 24,74181 | 24,55894 | 24,6536 | 25,27704 |
| sp\|O43852\|CALU_HUMAN;tr\|H0Y875\|H0Y875_HUMAN | sp\|O43852\|C | 24,381 | 102760000 | 21 | 7 | 0,48899149 | 0,13771915 | 22,23017 | 22,66262 | 22,8477 | 22,52578 | 22,7599 | 22,87693 | 22,87044 | 22,30989 |
| sp\|P31942\|HNRH3_HUMAN | sp\|P31942\|H | 23,963 | 60412000 | 16 | 4 | 0,49053858 | 0,23476648 | 21,18804 | 21,17883 | 22,09497 | 21,87083 | 21,52979 | 22,10124 | 22,26728 | 21,37342 |
| sp\|P29692\|EF1D_HUMAN;tr\|E9PK01\|E9PK01_HUMAN;tr\|E9PRY8\|E9PRY8_HUMAN;tr\|E9PMV | sp\|P29692\|E | 71,865 | 161890000 | 22 | 5 | 0,49741701 | -0,11526489 | 23,48431 | 23,15647 | 23,62255 | 23,52456 | 23,3839 | 23,32334 | 23,0103 | 23,6093 |
| sp\|Q9Y617\|SERC_HUMAN | sp\|Q9Y617\|S | 13,727 | 119570000 | 20 | 8 | 0,49923489 | -0,09596348 | 22,74189 | 23,01711 | 22,94156 | 23,16873 | 22,63163 | 23,07233 | 22,78903 | 22,99245 |
| sp\|Q12931\|TRAP1_HUMAN;tr\|I3L0K7\|I3L0K7_HUMAN;tr\|I3L239\|I3L239_HUMAN;tr\|I3L253\|I3 | sp\|Q12931\|T | 323,31 | 363390000 | 53 | 14 | 0,50407184 | -0,11430693 | 24,5948 | 24,76233 | 24,5625 | 24,89458 | 24,67131 | 24,39833 | 24,33559 | 24,95176 |
| sp\|P48643\|TCPE_HUMAN;tr\|E7ENZ3\|E7ENZ3_HUMAN;tr\|B7ZAR1\|B7ZAR1_HUMAN;tr\|E9PCA | sp\|P48643\|T | 323,31 | 535120000 | 64 | 18 | 0,50731832 | -0,07798386 | 24,79516 | 25,16434 | 25,22733 | 24,9389 | 24,88106 | 25,07985 | 24,88297 | 24,96992 |
| tr\|E9PMI6\|E9PMI6_HUMAN;tr\|E9PJF4\|E9PJF4_HUMAN;tr\|J3KN38\|J3KN38_HUMAN;sp\|P5410 | tr\|E9PMI6\|E | 7,8106 | 14464000 | 3 | 2 | 0,50795326 | -0,08696906 | 20,06592 | 19,98968 | 20,03778 | 20,17487 | 20,03282 | 19,72457 | NaN | 20,18289 |
| sp\|Q8N1G4\|LRC47_HUMAN | sp\|Q8N1G4\|L | 11,285 | 16448000 | 10 | 3 | 0,5163547 | -0,19526958 | NaN | 20,19508 | 20,13132 | 20,46314 | 20,39467 | 19,77054 | 19,59858 | 20,50785 |
| tr\|A0A0D9SFB3\|A0A0D9SFB3_HUMAN;tr\|A0A0D9SG12\|A0A0D9SG12_HUMAN;sp\|O00571\|DD | tr\|A0A0D9SF | 50,184 | 171150000 | 45 | 12 | 0,52058551 | -0,11161137 | 23,29908 | 23,37782 | 23,76739 | 23,37796 | 22,97179 | 23,5093 | 23,43695 | 23,45776 |
| sp\|P68871\|HBB_HUMAN;tr\|F8W6P5\|F8W6P5_HUMAN;sp\|P02042\|HBD_HUMAN;tr\|E9PFT6\|E | sp\|P68871\|H | 233,61 | 305450000 | 35 | 6 | 0,52385666 | -0,53465128 | 24,73738 | 24,35964 | 23,56221 | 25,00894 | 25,79434 | 23,59218 | 23,86231 | 22,28074 |
| sp\|Q15155\|NOMO1_HUMAN;sp\|P69849\|NOMO3_HUMAN;tr\|J3KN36\|J3KN36_HUMAN;tr\|A0 | sp\|Q15155\|N | 9,3513 | 17500000 | 7 | 4 | 0,52461204 | -0,06999874 | 20,36378 | 20,36965 | 20,22294 | 20,06342 | 20,10546 | 20,01781 | 20,28048 | 20,33604 |
| sp\|P05388\|RLA0_HUMAN;tr\|F8VWS0\|F8VWS0_HUMAN;tr\|F8VU65\|F8VU65_HUMAN;tr\|F8VP | sp\|P05388\|R | 286,75 | 771760000 | 48 | 11 | 0,52534426 | -0,13632536 | 25,74319 | 25,64576 | 25,87044 | 25,3652 | 25,26137 | 25,76162 | 25,86512 | 25,19118 |
| sp\|P53597\|SUCA_HUMAN | sp\|P53597\|S | 3,7078 | 22886000 | 5 | 2 | 0,5293824 | 0,10190248 | 20,38068 | 20,39142 | 20,40873 | 20,66484 | 20,59268 | 20,93869 | 20,34397 | 20,37793 |
| sp\|Q13895\|BYST_HUMAN;tr\|H7BY94\|H7BY94_HUMAN | sp\|Q13895\|B | 2,4791 | 7576200 | 3 | 2 | 0,53031353 | 0,09520245 | 18,88762 | 18,85538 | 18,88908 | 18,877 | 18,87658 | 18,65522 | 19,33632 | 19,02177 |
| sp\|P25788\|PSA3_HUMAN;tr\|G3V3W4\|G3V3W4_HUMAN;tr\|G3V5N4\|G3V5N4_HUMAN;tr\|G3 | sp\|P25788\|P | 58,925 | 81110000 | 19 | 6 | 0,53063341 | -0,23614931 | 22,50942 | 22,54463 | 22,97699 | 21,828 | 22,93135 | 22,33576 | 21,78676 | 21,86056 |
| sp\|Q14566\|MCM6_HUMAN | sp\|Q14566\|M | 46,097 | 117230000 | 28 | 12 | 0,53363054 | -0,14305353 | 23,20766 | 22,72507 | 23,14782 | 22,99519 | 22,68132 | 22,58072 | 22,82128 | 23,42021 |
| sp\|P06748\|NPM_HUMAN;tr\|E5RGW4\|E5RGW4_HUMAN;tr\|E5RI98\|E5RI98_HUMAN | sp\|P06748\|N | 323,31 | 1460600000 | 23 | 4 | 0,53485278 | -0,191154 | 26,06614 | 26,58492 | 26,99432 | 26,54963 | 25,96193 | 26,6983 | 26,7758 | 25,99436 |
| sp\|Q7L2H7\|EIF3M_HUMAN;tr\|J3KNJ2\|J3KNJ2_HUMAN;tr\|H0YCQ8\|H0YCQ8_HUMAN;tr\|E9PRI | sp\|Q7L2H7\|E | 128,5 | 16726000 | 11 | 4 | 0,54386547 | -0,18598509 | 19,77624 | NaN | 20,70658 | 20,28816 | 20,11273 | 20,09623 | 19,67623 | 20,39885 |
| tr\|E9PKG1\|E9PKG1_HUMAN;sp\|Q99873\|ANM1_HUMAN;tr\|H7C2I1\|H7C2I1_HUMAN;tr\|A0A0 | tr\|E9PKG1\|E | 13,77 | 41681000 | 9 | 4 | 0,54479822 | 0,29400508 | 21,37978 | NaN | 20,88904 | 21,48286 | 20,92158 | 22,30541 | 21,40671 | NaN |
| sp\|P19338\|NUCL_HUMAN;tr\|H7BY16\|H7BY16_HUMAN;tr\|C9JYW2\|C9JYW2_HUMAN;tr\|C9JLB | sp\|P19338\|N | 263,48 | 2701500000 | 82 | 24 | 0,55281107 | 0,11093235 | 27,48631 | 27,34942 | 27,61224 | 27,09356 | 27,15692 | 27,39034 | 27,72592 | 27,71208 |
| tr\|G8JLD5\|G8JLD5_HUMAN;sp\|O00429\|DNM1L_HUMAN;tr\|F8VZ52\|F8VZ52_HUMAN | tr\|G8JLD5\|G | 2,6087 | 8784800 | 5 | 3 | 0,55399981 | 0,06312799 | 19,16624 | 19,19371 | 19,17553 | 18,98618 | 19,04484 | 19,28134 | 19,04793 | 19,40006 |
| sp\|O60264\|SMCA5_HUMAN;sp\|P28370\|SMCA1_HUMAN;tr\|A0A0A0MRP6\|A0A0A0MRP6_HU | sp\|O60264\|S | 10,333 | 31764000 | 10 | 5 | 0,56334172 | 0,22339296 | 20,31661 | 20,34093 | 20,97673 | 20,7615 | 20,26178 | 20,72517 | 21,76211 | 20,54028 |
| tr\|H9KV75\|H9KV75_HUMAN;sp\|P12814\|ACTN1_HUMAN;tr\|H0YJ11\|H0YJ11_HUMAN;tr\|H0YJ | tr\|H9KV75\|H | 2,7192 | 10236000 | 3 | 2 | 0,57153878 | -0,11474705 | 19,75344 | 18,9725 | 19,69542 | 19,61597 | 19,38821 | 19,38924 | 19,2463 | 19,55461 |
| sp\|P62333\|PRS10_HUMAN;tr\|A0A087X2I1\|A0A087X2I1_HUMAN;tr\|H0YJC0\|H0YJC0_HUMAN; | sp\|P62333\|P | 19,546 | 90567000 | 24 | 8 | 0,57154257 | 0,06986332 | 22,239 | 22,52931 | 22,38118 | 22,58095 | 22,72038 | 22,55258 | 22,42437 | 22,31257 |
| sp\|P62913\|RL11_HUMAN;tr\|Q5VVC8\|Q5VVC8_HUMAN;tr\|Q5VVC9\|Q5VVC9_HUMAN | sp\|P62913\|R | 96,374 | 202950000 | 20 | 6 | 0,57330551 | 0,32159567 | 23,38139 | 23,40393 | 23,46239 | 23,31907 | 23,02284 | 24,92619 | 24,29039 | 22,61374 |
| tr\|A0A087WUL2\|A0A087WUL2_HUMAN;sp\|P49720\|PSB3_HUMAN;tr\|A0A087WXQ8\|A0A087\ | tr\|A0A087WU | 2,7611 | 31223000 | 6 | 2 | 0,5770458 | 0,17760754 | 21,16993 | 20,07483 | 21,34234 | 21,0654 | 21,07457 | 21,25753 | 20,8233 | 21,20753 |
| sp\|P35580\|MYH10_HUMAN | sp\|P35580\|M | 24,004 | 28565000 | 11 | 7 | 0,57914482 | -0,19866753 | 21,33604 | 21,12742 | 21,23298 | 20,6486 | 20,3393 | 20,3931 | 21,48822 | 21,32975 |
| sp\|Q00610\|CLH1_HUMAN;tr\|A0A087WVQ6\|A0A087WVQ6_HUMAN;tr\|J3KS13\|J3KS13_HUMA | sp\|Q00610\|C | 217,92 | 353700000 | 86 | 28 | 0,5825193 | -0,10971975 | 24,7113 | 24,60665 | 24,68362 | 24,15004 | 24,12747 | 24,26899 | 24,64669 | 24,66958 |
| sp\|Q9Y3I0\|RTCB_HUMAN | sp\|Q9Y3I0\|R | 18,11 | 54825000 | 11 | 5 | 0,58395948 | -0,23568678 | 21,50586 | 21,64878 | 21,63768 | 22,49629 | 20,5861 | 22,07874 | 21,86189 | 21,81913 |
| sp\|Q15717\|ELAV1_HUMAN;tr\|M0QZR9\|M0QZR9_HUMAN;tr\|M0R055\|M0R055_HUMAN | sp\|Q15717\|E | 10,866 | 65001000 | 18 | 7 | 0,58610058 | 0,22893 | 21,61216 | 21,33685 | 22,04217 | 21,93991 | 21,10661 | 22,16162 | 22,83954 | 21,73905 |
| tr\|A0A0U1RQF0\|A0A0U1RQF0_HUMAN;sp\|P49327\|FAS_HUMAN;tr\|A0A0U1RRG3\|A0A0U1RR | tr\|A0A0U1RC | 323,31 | 858820000 | 164 | 55 | 0,59260778 | -0,10887957 | 25,91593 | 26,06437 | 25,75659 | 25,59477 | 25,27753 | 25,68715 | 25,91472 | 26,01674 |
| sp\|P15924\|DESP_HUMAN | sp\|P15924\|D | 84,401 | 212490000 | 66 | 29 | 0,59430773 | -0,14674759 | 24,03507 | 24,23936 | 23,09713 | 23,81763 | 23,67982 | 23,52671 | 23,8624 | 23,53326 |
| sp\|P62263\|RS14_HUMAN;tr\|H0YB22\|H0YB22_HUMAN;tr\|E5RH77\|E5RH77_HUMAN | sp\|P62263\|R | 80,38 | 263380000 | 12 | 5 | 0,5993745 | -0,50638962 | NaN | 23,71931 | 25,11831 | 23,40276 | 24,06077 | 23,64372 | 21,72451 | 24,86594 |
| sp\|O60506\|HNRPQ_HUMAN;tr\|B7Z645\|B7Z645_HUMAN;tr\|F6UXX1\|F6UXX1_HUMAN | sp\|O60506\|H | 94,154 | 383690000 | 50 | 11 | 0,60183853 | -0,11222744 | 24,74566 | 24,43352 | 24,78561 | 24,38482 | 24,02732 | 24,67314 | 24,82221 | 24,37802 |
| sp\|P78371\|TCPB_HUMAN;tr\|F5GWF6\|F5GWF6_HUMAN;tr\|F8VQ14\|F8VQ14_HUMAN | sp\|P78371\|T | 255,23 | 634870000 | 77 | 21 | 0,6030227 | -0,07910967 | 25,17615 | 25,41351 | 25,4204 | 25,38752 | 24,99278 | 25,3669 | 25,13038 | 25,59109 |
| sp\|O75643\|U520_HUMAN;tr\|B4E0P5\|B4E0P5_HUMAN | sp\|O75643\|U | 142,26 | 151800000 | 48 | 22 | 0,61001557 | 0,13451958 | 23,27185 | 22,85177 | 22,95463 | 23,04382 | 22,49211 | 23,26069 | 23,56732 | 23,34001 |
| sp\|P48147\|PPCE_HUMAN | sp\|P48147\|P | 4,6291 | 10131000 | 10 | 3 | 0,61255191 | 0,06940889 | 19,12116 | 19,35518 | 19,36765 | 19,54202 | 19,43335 | 19,29167 | 19,25509 | 19,68353 |
| sp\|O94925\|GLSK_HUMAN;tr\|H7BZD1\|H7BZD1_HUMAN;tr\|C9JIJ6\|C9JIJ6_HUMAN | sp\|O94925\|G | 3,2434 | 31983000 | 3 | 2 | 0,61486616 | -0,15028667 | 20,92738 | 21,15875 | 21,08719 | 21,28963 | 21,16454 | 21,48822 | 21,00546 | 20,20359 |
| sp\|Q13049\|TRI32_HUMAN;tr\|Q5JVY0\|Q5JVY0_HUMAN | sp\|Q13049\|T | 34,444 | 125270000 | 25 | 13 | 0,61517467 | 0,41684151 | 21,0901 | 22,98141 | 21,73802 | 24,16131 | 23,98224 | 22,16383 | 22,97418 | 22,51795 |
| sp\|Q9P035\|HACD3_HUMAN;tr\|H3BPZ1\|H3BPZ1_HUMAN;tr\|H3BS72\|H3BS72_HUMAN;tr\|H3B | sp\|Q9P035\|H | 44,752 | 37970000 | 13 | 4 | 0,62031208 | -0,05809339 | 21,38052 | 21,65403 | 21,58449 | 21,45543 | 21,33631 | 21,39032 | 21,65495 | NaN |
| sp\|Q92900\|RENT1_HUMAN | sp\|Q92900\|R | 17,222 | 55507000 | 23 | 9 | 0,62665175 | 0,09114408 | 21,94174 | 21,96526 | 21,56132 | 21,79951 | 21,93146 | 21,53089 | 21,89667 | 22,27338 |
| sp\|P62906\|RL10A_HUMAN | sp\|P62906\|R | 34,061 | 592450000 | 26 | 8 | 0,6307285 | 0,15449905 | 24,3835 | 24,90834 | 25,48299 | 25,44525 | 24,73753 | 25,27508 | 25,41542 | 25,41006 |
| sp\|P08133\|ANXA6_HUMAN;tr\|E5RK69\|E5RK69_HUMAN;tr\|E7EMC6\|E7EMC6_HUMAN;tr\|H0Y | sp\|P08133\|A | 8,5505 | 35884000 | 10 | 6 | 0,63696423 | -0,09697533 | 21,12534 | 21,30145 | 20,88017 | 21,69477 | 20,99252 | 21,13239 | 21,0692 | 21,41972 |
| sp\|P62277\|RS13_HUMAN;tr\|J3KMX5\|J3KMX5_HUMAN;tr\|E9PS50\|E9PS50_HUMAN | sp\|P62277\|R | 72,03 | 185700000 | 20 | 6 | 0,63750698 | 0,16981745 | 23,81176 | 22,67816 | 23,8944 | 23,1897 | 23,96622 | 23,49692 | 23,07574 | 23,7144 |
| sp\|Q9BY77\|PDIP3_HUMAN;tr\|F6VRR5\|F6VRR5_HUMAN;tr\|Q8WUT1\|Q8WUT1_HUMAN;tr\|F8 | sp\|Q9BY77\|P | 4,4483 | 12174000 | 5 | 3 | 0,63802592 | -0,09670719 | NaN | 19,67908 | 19,75739 | 19,69147 | 19,88927 | 19,21516 | 19,47965 | 19,86635 |
| tr\|E9PBS1\|E9PBS1_HUMAN;sp\|P22234\|PUR6_HUMAN;tr\|D6RF62\|D6RF62_HUMAN | tr\|E9PBS1\|E | 66,062 | 271020000 | 31 | 10 | 0,6414089 | -0,14387178 | 24,21801 | 23,3652 | 24,38936 | 24,55625 | 23,79838 | 24,10957 | 23,75908 | 24,2863 |
| sp\|P49368\|TCPG_HUMAN;tr\|B4DUR8\|B4DUR8_HUMAN;tr\|E9PRC8\|E9PRC8_HUMAN;tr\|Q5SZ | sp\|P49368\|T | 62,936 | 582630000 | 60 | 15 | 0,64324817 | 0,06640625 | 25,04085 | 24,98272 | 25,40682 | 25,23301 | 25,26417 | 25,09684 | 25,07631 | 25,4917 |
| sp\|P07737\|PROF1_HUMAN;tr\|K7EJ44\|K7EJ44_HUMAN;CON__P02584;tr\|I3L3D5\|I3L3D5_HUM | sp\|P07737\|P | 64,41 | 301980000 | 26 | 6 | 0,64377029 | -0,1805048 | 24,94033 | 23,80882 | 23,88157 | 24,15522 | 24,11601 | 24,05001 | 23,30619 | 24,59172 |
| tr\|H7BXY3\|H7BXY3_HUMAN;sp\|Q7L2E3\|DHX30_HUMAN | tr\|H7BXY3\|H | 28,381 | 55371000 | 19 | 12 | 0,64430982 | 0,14969301 | 21,40447 | 21,46503 | 22,0192 | 21,39383 | 21,146 | 21,94167 | 22,35877 | 21,43487 |
| sp\|Q9UBB4\|ATX10_HUMAN;tr\|B1AHE3\|B1AHE3_HUMAN;tr\|B1AHE4\|B1AHE4_HUMAN | sp\|Q9UBB4\| | 5,8452 | 11836000 | 9 | 3 | 0,64563632 | 0,14484549 | 19,26061 | 19,38275 | 19,60357 | 19,72106 | 19,54836 | 19,48235 | 19,0936 | 20,42306 |
| tr\|A0A024RA52\|A0A024RA52_HUMAN;sp\|P25787\|PSA2_HUMAN;tr\|H3BT36\|H3BT36_HUMAN | tr\|A0A024RA | 21,048 | 34895000 | 14 | 4 | 0,64604543 | -0,11150122 | 21,25194 | 20,92904 | 21,31567 | 20,91539 | 21,06789 | 20,88881 | 20,51007 | 21,49926 |

| | | | | | | | | | | | | | | | |
|---|---|---|---|---|---|---|---|---|---|---|---|---|---|---|---|
| sp\|O00264\|PGRC1_HUMAN;tr\|U3KQM0\|U3KQM0_HUMAN | sp\|O00264\|P | 9,2433 | 135690000 | 13 | 5 | 0,64612085 | -0,10084009 | 22,8547 | 22,71389 | 23,49277 | 23,20262 | 23,12704 | 23,09553 | 23,00263 | 22,63542 |
| sp\|Q14978\|NOLC1_HUMAN;tr\|A0A0A0MRM9\|A0A0A0MRM9_HUMAN;tr\|S4R349\|S4R349_HU | sp\|Q14978\|N | 24,632 | 54395000 | 8 | 6 | 0,64745269 | 0,1388135 | 21,8665 | 21,56384 | 22,21848 | 21,4708 | 21,38401 | 22,14762 | 22,44017 | 21,70307 |
| tr\|A0A024R4E5\|A0A024R4E5_HUMAN;sp\|Q00341\|VIGLN_HUMAN;tr\|H0Y394\|H0Y394_HUMA | tr\|A0A024R4 | 7,6908 | 39687000 | 10 | 7 | 0,65257105 | -0,0569253 | 21,50096 | 21,34657 | 21,41617 | 21,22058 | 21,14824 | 21,23637 | 21,25159 | 21,62037 |
| sp\|Q9H444\|CHM4B_HUMAN | sp\|Q9H444\|C | 127,67 | 36889000 | 8 | 4 | 0,65735292 | 0,46270227 | 20,01332 | 20,31352 | 20,72376 | 20,23806 | 23,71534 | 19,95077 | 19,55822 | 19,91513 |
| tr\|Q5T3Q7\|Q5T3Q7_HUMAN;sp\|Q9H583\|HEAT1_HUMAN | tr\|Q5T3Q7\|Q | 11,601 | 13914000 | 7 | 6 | 0,65922615 | -0,07660087 | 19,84592 | 19,91139 | 19,85271 | 19,91069 | 19,44164 | 19,86427 | 20,10482 | NaN |
| tr\|J3KTF8\|J3KTF8_HUMAN;sp\|P52565\|GDIR1_HUMAN;tr\|J3QQX2\|J3QQX2_HUMAN;tr\|J3KRE2 | tr\|J3KTF8\|J3K | 24,262 | 62291000 | 12 | 4 | 0,65950559 | 0,05018759 | 21,79229 | 21,91747 | 21,9668 | 22,11737 | 22,16303 | 22,07682 | 21,76765 | 21,98718 |
| tr\|A0A087WV29\|A0A087WV29_HUMAN;sp\|Q9H0A0\|NAT10_HUMAN | tr\|A0A087WV | 8,2658 | 12076000 | 6 | 4 | 0,66155752 | 0,12489112 | NaN | 19,5441 | 19,60634 | 19,25306 | 19,23678 | 19,48613 | 20,21288 | 19,4351 |
| sp\|P51659\|DHB4_HUMAN;tr\|E7ER27\|E7ER27_HUMAN;tr\|E7EWE5\|E7EWE5_HUMAN;tr\|G5E9 | sp\|P51659\|D | 71,401 | 160410000 | 33 | 11 | 0,66186585 | -0,09632158 | 23,25782 | 23,40834 | 23,46463 | 23,70356 | 23,34706 | 23,23301 | 22,9893 | 23,8797 |
| sp\|P37802\|TAGL2_HUMAN;tr\|X6RJP6\|X6RJP6_HUMAN;tr\|C9J5W6\|C9J5W6_HUMAN;sp\|Q9UI | sp\|P37802\|T | 175,57 | 320260000 | 35 | 7 | 0,66200617 | -0,10676908 | 24,17333 | 24,46631 | 24,45174 | 24,64642 | 24,46039 | 24,54699 | 23,70134 | 24,60201 |
| sp\|Q9Y262\|EIF3L_HUMAN;tr\|B0QY89\|B0QY89_HUMAN;tr\|B0QY90\|B0QY90_HUMAN;tr\|C9JH | sp\|Q9Y262\|E | 27,943 | 173900000 | 42 | 13 | 0,66340845 | -0,06530428 | 23,63733 | 23,47222 | 23,55415 | 23,25184 | 23,17048 | 23,40743 | 23,34854 | 23,72786 |
| sp\|Q08945\|SSRP1_HUMAN;tr\|E9PMD4\|E9PMD4_HUMAN;tr\|A0A0U1RRK2\|A0A0U1RRK2_HUN | sp\|Q08945\|S | 94,317 | 103260000 | 18 | 10 | 0,66406505 | 0,19456053 | 22,29496 | 22,16999 | 22,58155 | 22,80853 | 22,0255 | 22,30922 | 23,83026 | 22,46829 |
| tr\|E9PDE8\|E9PDE8_HUMAN;sp\|O95757\|HS74L_HUMAN;tr\|D6RJ96\|D6RJ96_HUMAN | tr\|E9PDE8\|E9 | 14,458 | 33283000 | 16 | 8 | 0,66568268 | 0,1009841 | 21,17487 | 21,15072 | 20,89726 | 21,07764 | 21,17432 | 20,97386 | 20,78604 | 21,7702 |
| sp\|P62851\|RS25_HUMAN | sp\|P62851\|R | 24,245 | 454900000 | 14 | 4 | 0,66828124 | -0,09971523 | 24,53291 | 25,04623 | 24,91857 | 24,7348 | 24,88008 | 25,14344 | 24,53843 | 24,2717 |
| tr\|F8VYE8\|F8VYE8_HUMAN;sp\|P36873\|PP1G_HUMAN;tr\|F8W0W8\|F8W0W8_HUMAN;tr\|F8V | tr\|F8VYE8\|F8 | 70,494 | 71259000 | 31 | 1 | 0,67178726 | 0,07748318 | 21,97876 | 21,9827 | 22,3747 | 22,09313 | 22,17536 | 22,49512 | 22,27736 | 21,79138 |
| tr\|I3L1L3\|I3L1L3_HUMAN;sp\|Q9BQG0\|MBB1A_HUMAN;tr\|I3L2H8\|I3L2H8_HUMAN | tr\|I3L1L3\|I3L | 23,601 | 52376000 | 14 | 13 | 0,67494626 | -0,13042688 | 21,81014 | 21,81577 | 21,92673 | 21,57565 | 21,10021 | 21,46827 | 22,45408 | 21,58403 |
| sp\|Q07955\|SRSF1_HUMAN;tr\|J3KTL2\|J3KTL2_HUMAN;tr\|J3QQV5\|J3QQV5_HUMAN;tr\|J3KSW | sp\|Q07955\|S | 44,686 | 118180000 | 22 | 9 | 0,67582443 | -0,0796895 | 22,80802 | 22,61113 | 23,00122 | 23,22162 | 22,55112 | 22,68607 | 23,02744 | 23,05859 |
| sp\|P51149\|RAB7A_HUMAN;tr\|C9J8S3\|C9J8S3_HUMAN;tr\|C9J4V0\|C9J4V0_HUMAN;tr\|C9J4S4 | sp\|P51149\|R | 52,424 | 317310000 | 30 | 9 | 0,67634331 | 0,11392355 | 23,625 | 24,3532 | 24,21712 | 24,40633 | 24,41654 | 24,72265 | 23,91971 | 23,99843 |
| sp\|P22626\|ROA2_HUMAN;tr\|A0A087WUI2\|A0A087WUI2_HUMAN | sp\|P22626\|R | 316,15 | 660500000 | 58 | 12 | 0,68207698 | 0,14356995 | 25,15815 | 24,64339 | 25,51947 | 25,34973 | 24,71639 | 25,53769 | 25,96005 | 25,03091 |
| tr\|J3KR24\|J3KR24_HUMAN;sp\|P41252\|SYIC_HUMAN;tr\|A0A0A0MSX9\|A0A0A0MSX9_HUMAN | tr\|J3KR24\|J3 | 21,02 | 143770000 | 31 | 12 | 0,68784522 | -0,05679417 | 23,44771 | 23,44329 | 23,16054 | 23,05565 | 23,09719 | 23,03428 | 23,39663 | 23,35192 |
| sp\|Q8IZL8\|PELP1_HUMAN;tr\|C9JFV4\|C9JFV4_HUMAN;tr\|I3L445\|I3L445_HUMAN;tr\|E7EV54\|E | sp\|Q8IZL8\|PE | 9,3914 | 26544000 | 10 | 6 | 0,69255071 | 0,12106228 | 20,64746 | 20,46702 | 20,46374 | 20,37263 | 19,9894 | 20,70269 | 21,34624 | 20,39676 |
| sp\|Q8TEX9\|IPO4_HUMAN;tr\|H0YN14\|H0YN14_HUMAN;tr\|H0YMR4\|H0YMR4_HUMAN;tr\|H0Y | sp\|Q8TEX9\|I | 26,08 | 74376000 | 24 | 10 | 0,69419323 | -0,06734276 | 22,22885 | 22,32743 | 22,24589 | 22,40395 | 21,98995 | 22,29561 | 21,994 | 22,65718 |
| sp\|P62280\|RS11_HUMAN;tr\|M0QZC5\|M0QZC5_HUMAN;tr\|M0R1H5\|M0R1H5_HUMAN;tr\|M( | sp\|P62280\|R | 69,912 | 664810000 | 37 | 14 | 0,69939545 | -0,07404041 | 25,43396 | 25,89412 | 25,56485 | 25,08371 | 25,45779 | 25,43094 | 25,22133 | 25,57043 |
| sp\|P61619\|S61A1_HUMAN;tr\|B4DR61\|B4DR61_HUMAN;tr\|H7C1Q9\|H7C1Q9_HUMAN;tr\|C9J) | sp\|P61619\|S | 74,553 | 35289000 | 9 | 3 | 0,70517278 | 0,15238142 | 20,42777 | 21,77133 | 21,06763 | 20,94149 | 20,93948 | 21,24632 | 21,92933 | 20,70261 |
| sp\|Q01581\|HMCS1_HUMAN;tr\|D6RIW1\|D6RIW1_HUMAN | sp\|Q01581\|H | 9,5958 | 27829000 | 3 | 4 | 0,70797968 | -0,04666948 | 20,50659 | 20,96176 | 20,78101 | 20,79685 | 20,6677 | 20,92289 | 20,6802 | 20,58875 |
| tr\|G3V4W0\|G3V4W0_HUMAN;tr\|B4DY08\|B4DY08_HUMAN;tr\|B2R5W2\|B2R5W2_HUMAN;tr\| | tr\|G3V4W0\|( | 84,011 | 580710000 | 35 | 12 | 0,70813194 | 0,19563532 | 24,659 | 24,20672 | 25,3017 | 25,08618 | 23,90454 | 25,31721 | 25,97373 | 24,84067 |
| tr\|E9PEX6\|E9PEX6_HUMAN;sp\|P09622\|DLDH_HUMAN | tr\|E9PEX6\|E9 | 29,249 | 88409000 | 20 | 6 | 0,70950166 | 0,04832077 | 22,30324 | 22,16898 | 22,40691 | 22,58857 | 22,42206 | 22,57013 | 22,49636 | 22,17243 |
| sp\|Q8TDN6\|BRX1_HUMAN | sp\|Q8TDN6\| | 2,9518 | 9222900 | 4 | 2 | 0,71056674 | -0,11900345 | 19,51138 | 19,21966 | 19,57941 | 19,252 | 18,74678 | 19,16362 | 19,90442 | NaN |
| sp\|Q13011\|ECH1_HUMAN;tr\|M0QZW4\|M0QZW4_HUMAN;tr\|M0R248\|M0R248_HUMAN | sp\|Q13011\|E | 2,6158 | 12539000 | 5 | 2 | 0,71409183 | -0,0483222 | 19,81397 | 19,73587 | 19,68274 | 19,93431 | 19,84382 | 19,58401 | 19,53123 | 20,01454 |
| sp\|P61006\|RAB8A_HUMAN | sp\|P61006\|R | 30,677 | 351160000 | 26 | 5 | 0,71454235 | -0,11796331 | 23,95952 | 24,54904 | 24,47091 | 24,67761 | 24,1715 | 25,06336 | 24,09233 | 23,85804 |
| sp\|P23381\|SYWC_HUMAN;tr\|H0YJP3\|H0YJP3_HUMAN;tr\|G3V3H8\|G3V3H8_HUMAN;tr\|G3V3 | sp\|P23381\|S | 4,4775 | 22479000 | 7 | 4 | 0,7155321 | -0,06584549 | 20,41318 | 20,55805 | 20,76077 | 20,75369 | 20,24505 | 20,60892 | NaN | 20,81276 |
| sp\|P42167\|LAP2B_HUMAN;tr\|G5E972\|G5E972_HUMAN;tr\|H0YJH7\|H0YJH7_HUMAN | sp\|P42167\|L | 13,445 | 61568000 | 13 | 5 | 0,71615763 | 0,18481684 | 21,29317 | 21,50722 | 22,25263 | 22,39132 | 21,30647 | 22,04537 | 22,78586 | NaN |
| sp\|O43143\|DHX15_HUMAN | sp\|O43143\|D | 31,012 | 254590000 | 35 | 12 | 0,71781215 | 0,04616308 | 24,0258 | 23,98815 | 24,23067 | 23,72151 | 24,10534 | 23,91816 | 23,94824 | 24,17904 |
| sp\|P11021\|GRP78_HUMAN | sp\|P11021\|G | 323,31 | 931180000 | 99 | 23 | 0,71917747 | 0,01792431 | 25,79875 | 25,87572 | 25,94592 | 25,9844 | 25,89696 | 25,99162 | 25,89904 | 25,88888 |
| sp\|O00231\|PSD11_HUMAN;tr\|J3QRY4\|J3QRY4_HUMAN | sp\|O00231\|P | 18,015 | 71450000 | 18 | 9 | 0,72645874 | -0,04733515 | 22,3396 | 22,05577 | 22,19895 | 22,15962 | 22,1053 | 22,11245 | 21,89556 | 22,45129 |
| sp\|P22061\|PIMT_HUMAN;tr\|A0A0A0MRJ6\|A0A0A0MRJ6_HUMAN;tr\|H7BY58\|H7BY58_HUMA | sp\|P22061\|P | 35,367 | 204210000 | 31 | 5 | 0,72773243 | -0,08762932 | 23,40717 | 23,93085 | 23,68239 | 23,86041 | 23,92443 | 23,72713 | 23,01512 | 23,86363 |
| tr\|V9H019\|V9H019_HUMAN;tr\|E9PHA6\|E9PHA6_HUMAN;sp\|P43246\|MSH2_HUMAN;tr\|A0A( | tr\|V9H019\|V | 13,982 | 27855000 | 9 | 7 | 0,72823311 | 0,07061434 | 20,6778 | 20,71373 | 21,07352 | 20,91576 | 21,02039 | 20,66154 | 20,62872 | 21,35262 |
| tr\|A0A087WW66\|A0A087WW66_HUMAN;sp\|Q99460\|PSMD1_HUMAN;tr\|H7C378\|H7C378_H | tr\|A0A087W\ | 32,542 | 56395000 | 19 | 7 | 0,72948643 | 0,05679274 | 21,57473 | 21,78979 | 22,05364 | 21,89093 | 21,77884 | 21,6387 | 21,91528 | 22,20344 |
| sp\|P21333\|FLNA_HUMAN;tr\|Q5HY54\|Q5HY54_HUMAN;tr\|Q60FE5\|Q60FE5_HUMAN;tr\|A0A0 | sp\|P21333\|F | 319,21 | 793130000 | 146 | 51 | 0,73288112 | -0,06081009 | 25,7619 | 25,79694 | 25,66998 | 25,37657 | 25,26585 | 25,44922 | 25,78024 | 25,86684 |
| sp\|O14776\|TCRG1_HUMAN;tr\|G3V220\|G3V220_HUMAN | sp\|O14776\|T | 16,974 | 40706000 | 13 | 6 | 0,74621884 | -0,0928793 | 21,23719 | 21,45558 | 21,59478 | 21,24743 | 20,77523 | 21,1483 | 21,22712 | 22,01281 |
| sp\|Q13595\|TRA2A_HUMAN | sp\|Q13595\|T | 4,1706 | 9730200 | 8 | 2 | 0,75321208 | 0,13005447 | 18,88748 | 18,67252 | 19,75427 | 19,04014 | 18,57223 | 19,54518 | 19,94364 | 18,81359 |
| sp\|P54886\|P5CS_HUMAN | sp\|P54886\|P | 210,62 | 261950000 | 44 | 15 | 0,76238981 | -0,04766941 | 23,7102 | 24,32704 | 24,07969 | 24,13406 | 24,10205 | 23,86948 | 23,89892 | 24,18986 |
| tr\|F2Z2Q9\|F2Z2Q9_HUMAN;tr\|G5E9R5\|G5E9R5_HUMAN;tr\|D3YTI2\|D3YTI2_HUMAN;sp\|P246 | tr\|F2Z2Q9\|F | 27,507 | 34651000 | 6 | 2 | 0,76321988 | 0,07427597 | 21,13752 | 20,82664 | NaN | 21,3899 | NaN | 21,27219 | 20,87897 | 21,42573 |
| sp\|Q8WXF1\|PSPC1_HUMAN;tr\|X6RDA4\|X6RDA4_HUMAN | sp\|Q8WXF1\| | 11,799 | 35457000 | 8 | 5 | 0,76513205 | -0,09650501 | 21,46822 | NaN | 21,43354 | 21,1187 | 21,72213 | 21,14694 | 20,59632 | 21,5092 |
| sp\|P12081\|SYHC_HUMAN;tr\|B3KWE1\|B3KWE1_HUMAN;tr\|E7ETE2\|E7ETE2_HUMAN;tr\|B4E1( | sp\|P12081\|S | 19,478 | 62223000 | 16 | 4 | 0,76820493 | 0,08202934 | 21,24278 | 22,38432 | 22,13887 | 22,12931 | 22,18211 | 21,88494 | 21,92249 | 22,23386 |
| sp\|Q06203\|PUR1_HUMAN | sp\|Q06203\|P | 3,4701 | 26627000 | 8 | 3 | 0,77516584 | -0,05828619 | 20,62774 | NaN | 20,78835 | 21,13602 | 20,98482 | 20,78875 | 20,44263 | 20,95347 |
| sp\|Q08211\|DHX9_HUMAN | sp\|Q08211\|D | 323,31 | 640080000 | 62 | 24 | 0,77847202 | -0,06515646 | 25,35525 | 25,29309 | 25,44291 | 25,12605 | 24,77294 | 25,15151 | 25,79556 | 25,23666 |
| sp\|P30153\|2AAA_HUMAN;tr\|B3KQV6\|B3KQV6_HUMAN;tr\|C9J9C1\|C9J9C1_HUMAN;sp\|P3015 | sp\|P30153\|2 | 143,49 | 331410000 | 39 | 13 | 0,78038128 | 0,0410533 | 24,12849 | 24,30062 | 24,58606 | 24,46905 | 24,53843 | 24,41745 | 24,13006 | 24,5625 |
| sp\|P36871\|PGM1_HUMAN | sp\|P36871\|P | 2,6432 | 4699300 | 2 | 2 | 0,78359171 | 0,04398044 | NaN | 18,34863 | 18,36098 | 18,23547 | 18,25179 | 18,57163 | 18,05574 | 18,55686 |
| sp\|Q15393\|SF3B3_HUMAN;tr\|J3QRB2\|J3QRB2_HUMAN;tr\|J3QL37\|J3QL37_HUMAN;tr\|H3BM | sp\|Q15393\|S | 50,35 | 216740000 | 45 | 16 | 0,788038 | -0,03494883 | 23,80361 | 23,94306 | 23,95038 | 23,61953 | 23,62879 | 23,62333 | 23,95704 | 23,96763 |
| sp\|P46783\|RS10_HUMAN;tr\|F6U211\|F6U211_HUMAN;tr\|S4R435\|S4R435_HUMAN;sp\|Q9NQ3 | sp\|P46783\|R | 123,61 | 455950000 | 31 | 6 | 0,79246301 | -0,04757214 | 24,98316 | 24,55818 | 24,80302 | 24,78801 | 24,77997 | 25,13426 | 24,44828 | 24,57957 |
| sp\|Q16850\|CP51A_HUMAN;tr\|A0A0C4DFL7\|A0A0C4DFL7_HUMAN;tr\|H7C0D0\|H7C0D0_HUM. | sp\|Q16850\|C | 10,675 | 35096000 | 9 | 5 | 0,79258901 | 0,03637838 | 21,11184 | 21,11946 | 21,0283 | 21,33554 | 21,29821 | 20,99983 | 20,98815 | 21,45448 |
| tr\|B4DXW1\|B4DXW1_HUMAN;sp\|P61158\|ARP3_HUMAN;tr\|A0A0A0MTI9\|A0A0A0MTI9_HUM | tr\|B4DXW1\|E | 7,2066 | 64382000 | 11 | 6 | 0,79595837 | -0,04396582 | 21,86737 | 22,0496 | 22,06487 | 22,19818 | 21,66027 | 22,28153 | 21,85041 | 22,21193 |
| tr\|A0A087WTT1\|A0A087WTT1_HUMAN;sp\|P11940\|PABP1_HUMAN;tr\|E7EQV3\|E7EQV3_HUN | tr\|A0A087WT | 32,577 | 337340000 | 31 | 9 | 0,79768441 | 0,02975082 | 24,53611 | 24,30215 | 24,58663 | 24,24336 | 24,45406 | 24,25911 | 24,46625 | 24,60784 |
| sp\|P12270\|TPR_HUMAN;tr\|Q5SWX9\|Q5SWX9_HUMAN | sp\|P12270\|T | 41,646 | 84915000 | 24 | 16 | 0,80370712 | -0,05482483 | 22,71257 | 22,15437 | 22,81925 | 22,15054 | 22,17228 | 22,25655 | 22,54675 | 22,64185 |
| tr\|A0A0J9YYL3\|A0A0J9YYL3_HUMAN;tr\|A0A0J9YXJ8\|A0A0J9YXJ8_HUMAN;tr\|A0A0J9YVP6\|A0 | tr\|A0A0J9YYL | 29,684 | 39842000 | 8 | 6 | 0,80485071 | 0,06166458 | 20,86627 | 21,32367 | 21,29109 | 21,60246 | 21,21193 | 21,0351 | 21,87094 | 21,21217 |
| sp\|Q00839\|HNRPU_HUMAN;tr\|Q5RI18\|Q5RI18_HUMAN | sp\|Q00839\|H | 323,31 | 1270600000 | 66 | 20 | 0,80538118 | -0,0725956 | 26,28732 | 26,17356 | 26,50611 | 26,12118 | 25,59431 | 26,13792 | 26,90288 | 26,16267 |
| tr\|A0A0R4J2E8\|A0A0R4J2E8_HUMAN;sp\|P43243\|MATR3_HUMAN;tr\|A8MXP9\|A8MXP9_HUM | tr\|A0A0R4J2E | 44,336 | 90983000 | 18 | 10 | 0,80732898 | 0,09055519 | 22,18029 | 22,00607 | 22,66082 | 22,23491 | 21,66049 | 22,14942 | 23,22425 | 22,41015 |
| sp\|Q7Z2W4\|ZCCHV_HUMAN;tr\|C9J6P4\|C9J6P4_HUMAN | sp\|Q7Z2W4\| | 10,784 | 19767000 | 5 | 4 | 0,80794505 | -0,03290002 | 20,58105 | NaN | 20,26888 | 20,40291 | 20,38269 | 20,1437 | 20,55412 | 20,45834 |
| tr\|H3BT71\|H3BT71_HUMAN;sp\|P38159\|RBMX_HUMAN;tr\|H0Y6E7\|H0Y6E7_HUMAN;tr\|H3BR | tr\|H3BT71\|H | 39,792 | 173420000 | 24 | 6 | 0,80856229 | 0,13053799 | 22,84086 | 22,80166 | 23,56407 | 23,38311 | 22,23178 | 23,73624 | 24,3686 | 22,77523 |
| sp\|P08708\|RS17_HUMAN;tr\|H0YN88\|H0YN88_HUMAN;tr\|A0A075B716\|A0A075B716_HUMAI | sp\|P08708\|R | 10,278 | 125060000 | 12 | 4 | 0,81753439 | -0,06614876 | 23,5518 | 22,56584 | 22,82939 | 22,92868 | 23,2387 | 23,16671 | 22,69732 | 22,50838 |
| tr\|B8ZZU8\|B8ZZU8_HUMAN;sp\|Q15370\|ELOB_HUMAN;tr\|I3L0M9\|I3L0M9_HUMAN;tr\|A0A0B | tr\|B8ZZU8\|B | 15,726 | 47016000 | 13 | 4 | 0,81953538 | -0,08800602 | 21,15512 | 21,24951 | 21,84881 | 21,80715 | 21,60372 | 21,87222 | 21,75528 | 20,47734 |
| sp\|Q12906\|ILF3_HUMAN;tr\|K7EKJ9\|K7EKJ9_HUMAN;tr\|K7ER69\|K7ER69_HUMAN;tr\|K7EQR9\| | sp\|Q12906\|I | 89,617 | 363380000 | 43 | 14 | 0,82310683 | -0,05857277 | 24,80843 | 24,30654 | 24,67352 | 24,18715 | 23,99723 | 24,34482 | 24,97767 | 24,42163 |
| sp\|P61586\|RHOA_HUMAN;tr\|C9JRM1\|C9JRM1_HUMAN;tr\|C9JNR4\|C9JNR4_HUMAN;tr\|C9JX2 | sp\|P61586\|R | 11,155 | 33889000 | 6 | 2 | 0,82664097 | 0,14829334 | 20,4958 | NaN | 20,56941 | 21,1061 | 20,36025 | 22,44905 | 20,36656 | 20,31241 |
| tr\|G5E9W3\|G5E9W3_HUMAN;sp\|Q9UKF6\|CPSF3_HUMAN | tr\|G5E9W3\|( | 1,8797 | 7718300 | 2 | 2 | 0,83098285 | -0,02691968 | 19,28697 | 19,36818 | NaN | 19,1818 | 19,39452 | 19,3148 | 19,04687 | NaN |
| tr\|E5RI99\|E5RI99_HUMAN;sp\|P62888\|RL30_HUMAN;tr\|A0A0B4J213\|A0A0B4J213_HUMAN;tr | tr\|E5RI99\|E5 | 78,718 | 111850000 | 12 | 4 | 0,83256751 | -0,03728533 | 22,69356 | 22,68942 | 23,11601 | 22,69136 | 22,56923 | 22,99319 | 22,97956 | 22,49924 |
| sp\|P08559\|ODPA_HUMAN;tr\|Q5JPU0\|Q5JPU0_HUMAN;tr\|Q5JPT9\|Q5JPT9_HUMAN;tr\|Q5JPU | sp\|P08559\|O | 8,0388 | 114450000 | 18 | 6 | 0,83442408 | -0,03149796 | 22,64203 | 22,89698 | 22,79794 | 23,16129 | 23,0071 | 22,96608 | 22,80886 | 22,59021 |
| tr\|F2Z393\|F2Z393_HUMAN;sp\|P37837\|TALDO_HUMAN | tr\|F2Z393\|F2 | 16,319 | 169490000 | 14 | 5 | 0,83609157 | 0,13702742 | 23,12901 | 24,0253 | 22,05639 | 24,31341 | 23,14718 | 23,90288 | NaN | 23,5041 |
| tr\|B4DY09\|B4DY09_HUMAN;sp\|Q12905\|ILF2_HUMAN;tr\|X6R6Z1\|X6R6Z1_HUMAN;tr\|A0A0A( | tr\|B4DY09\|B | 59,585 | 203340000 | 32 | 8 | 0,83787865 | 0,04769516 | 23,54439 | 23,51147 | 23,92162 | 23,46787 | 23,11626 | 23,82172 | 24,04051 | 23,65764 |
| sp\|P60891\|PRPS1_HUMAN;tr\|B1ALA9\|B1ALA9_HUMAN;tr\|A0A0B4J207\|A0A0B4J207_HUMAN | sp\|P60891\|P | 36,823 | 81797000 | 14 | 2 | 0,85736471 | 0,04654741 | 22,42668 | 22,09484 | 22,44223 | 22,6804 | 22,38055 | 22,86081 | 21,88024 | 22,70872 |

| | | | | | | | | | | | | | | | |
|---|---|---|---|---|---|---|---|---|---|---|---|---|---|---|---|
| tr\|F5H1S8\|F5H1S8_HUMAN;tr\|F5GX14\|F5GX14_HUMAN;sp\|Q14165\|MLEC_HUMAN;tr\|H0YG | tr\|F5H1S8\|F5 | 4,2111 | 10183000 | 5 | 2 | 0,85777637 | 0,03393126 | 19,39236 | 19,35204 | 19,58448 | 19,31754 | 19,76801 | 18,96079 | 19,52241 | 19,53094 |
| sp\|P63092\|GNAS2_HUMAN;sp\|Q5JWF2\|GNAS1_HUMAN;tr\|A0A087WTB6\|A0A087WTB6_HUM | sp\|P63092\|G | 3,4451 | 10823000 | 4 | 3 | 0,86025774 | 0,09989977 | 18,50421 | 19,8903 | 19,72059 | NaN | 19,79986 | 20,07535 | 19,47508 | 18,53609 |
| sp\|Q13151\|ROA0_HUMAN | sp\|Q13151\|R | 13,581 | 123840000 | 15 | 5 | 0,86183289 | -0,08686829 | 22,36228 | 22,22647 | 23,49167 | 22,82196 | 22,10847 | 23,11095 | 23,46176 | 21,87373 |
| sp\|Q13247\|SRSF6_HUMAN;tr\|A0A0D9SEM4\|A0A0D9SEM4_HUMAN;sp\|Q08170\|SRSF4_HUM | sp\|Q13247\|S | 15,459 | 57956000 | 9 | 2 | 0,8627858 | 0,0321722 | 21,63139 | 21,59099 | 22,03181 | 21,79432 | 21,6446 | 21,84598 | 22,18365 | 21,50295 |
| sp\|P62854\|RS26_HUMAN;sp\|Q5JNZ5\|RS26L_HUMAN | sp\|P62854\|R | 3,8877 | 226860000 | 10 | 2 | 0,8644041 | 0,09948826 | 22,23002 | 24,39185 | 24,62901 | 23,99921 | 24,0559 | 23,76729 | 23,63334 | 24,19152 |
| sp\|Q6P2Q9\|PRP8_HUMAN;tr\|I3L0J9\|I3L0J9_HUMAN;tr\|I3L3Z8\|I3L3Z8_HUMAN | sp\|Q6P2Q9\| | 49,856 | 73359000 | 23 | 11 | 0,86594539 | -0,03290415 | 22,08845 | 22,32858 | 22,11839 | 21,87128 | 21,69166 | 21,982 | 22,46834 | 22,13308 |
| sp\|Q07065\|CKAP4_HUMAN | sp\|Q07065\|C | 11,946 | 29357000 | 14 | 5 | 0,86662668 | -0,00758457 | 20,92976 | 20,82974 | 20,9512 | 20,92172 | 20,9011 | 20,8226 | 20,89045 | 20,98794 |
| sp\|P54727\|RD23B_HUMAN;tr\|Q5W0S5\|Q5W0S5_HUMAN;tr\|Q5W0S4\|Q5W0S4_HUMAN | sp\|P54727\|R | 13,253 | 42216000 | 5 | 3 | 0,8690407 | 0,04854027 | 21,40416 | 20,8053 | NaN | 21,8064 | 21,16337 | 21,35326 | 21,31766 | 21,71436 |
| sp\|O75439\|MPPB_HUMAN;tr\|G3V0E4\|G3V0E4_HUMAN;tr\|F8WAZ6\|F8WAZ6_HUMAN;tr\|F8V | sp\|O75439\|M | 10,087 | 27743000 | 9 | 4 | 0,87072684 | 0,0267005 | 21,0448 | 20,84698 | 20,99106 | 20,61935 | 21,24696 | 20,79265 | 20,66284 | 20,90654 |
| sp\|P13489\|RINI_HUMAN;tr\|H0YCR7\|H0YCR7_HUMAN;tr\|E9PMJ3\|E9PMJ3_HUMAN;tr\|E9PLZ3 | sp\|P13489\|R | 43,267 | 34117000 | 16 | 5 | 0,87154538 | 0,02747631 | 20,77965 | 20,98461 | 21,05186 | 21,45498 | 21,12912 | 21,25954 | 20,87313 | 21,11921 |
| sp\|Q8WUA2\|PPIL4_HUMAN | sp\|Q8WUA2\| | 4,5755 | 10064000 | 5 | 3 | 0,87301611 | -0,02110243 | 19,48792 | 19,28604 | 19,38934 | 19,43831 | 19,40786 | 19,24488 | 19,16268 | 19,70178 |
| tr\|C9JLU1\|C9JLU1_HUMAN;sp\|P52434\|RPAB3_HUMAN | tr\|C9JLU1\|C9 | 14,056 | 42017000 | 10 | 3 | 0,8749092 | 0,04672352 | NaN | 21,15047 | 21,6356 | 21,34727 | 21,50964 | 21,99828 | 21,16282 | 21,02729 |
| tr\|M0R0F0\|M0R0F0_HUMAN;sp\|P46782\|RS5_HUMAN;tr\|M0R0R2\|M0R0R2_HUMAN;tr\|M0Q | tr\|M0R0F0\|M | 54,622 | 470460000 | 13 | 7 | 0,87751129 | -0,03529406 | 24,72312 | 25,25961 | 25,08201 | 24,61785 | 24,8274 | 25,33987 | 24,78122 | 24,59292 |
| sp\|Q04760\|LGUL_HUMAN | sp\|Q04760\|L | 7,0221 | 82145000 | 9 | 3 | 0,87790537 | 0,02723932 | 22,47719 | 22,19352 | 22,40795 | 22,9456 | 22,5154 | 22,59391 | 22,37008 | 22,65384 |
| tr\|G3V1C3\|G3V1C3_HUMAN;sp\|Q9BZZ5\|API5_HUMAN;tr\|E9PQK6\|E9PQK6_HUMAN;tr\|H0YEF | tr\|G3V1C3\|G | 7,6606 | 76751000 | 11 | 7 | 0,88089795 | -0,03243113 | 22,07078 | 22,20934 | 22,71507 | 22,31838 | 22,08654 | 22,73568 | 22,07467 | 22,28695 |
| sp\|Q9H0S4\|DDX47_HUMAN;tr\|F5H1N9\|F5H1N9_HUMAN | sp\|Q9H0S4\|D | 4,2277 | 5405300 | 4 | 3 | 0,8814272 | -0,02789688 | NaN | 18,37984 | 18,573 | 18,70675 | NaN | 18,58053 | 18,74895 | 18,24642 |
| sp\|P50395\|GDIB_HUMAN;tr\|Q5SX91\|Q5SX91_HUMAN;tr\|Q5SX87\|Q5SX87_HUMAN;tr\|Q5SX8 | sp\|P50395\|G | 25,766 | 107650000 | 20 | 4 | 0,88231966 | -0,04518843 | 22,957 | 22,9107 | 23,00208 | 22,52564 | 22,37849 | 22,87098 | 22,41939 | 23,54581 |
| sp\|P54136\|SYRC_HUMAN;tr\|E5RH09\|E5RH09_HUMAN;tr\|E5RJM9\|E5RJM9_HUMAN | sp\|P54136\|S | 21,755 | 105780000 | 25 | 12 | 0,88657192 | -0,01426411 | 22,99226 | 22,61758 | 22,92078 | 22,88653 | 22,71777 | 22,85055 | 22,83101 | 22,96076 |
| sp\|Q92841\|DDX17_HUMAN;tr\|H3BLZ8\|H3BLZ8_HUMAN;tr\|A0A0U1RQJ0\|A0A0U1RQJ0_HUM | sp\|Q92841\|D | 26,256 | 50096000 | 15 | 5 | 0,89291029 | 0,08454943 | 19,78681 | 21,61742 | 22,15419 | 22,33194 | 21,39273 | 21,30028 | 21,54213 | 21,99341 |
| sp\|Q9Y5B9\|SP16H_HUMAN | sp\|Q9Y5B9\|S | 27,378 | 66768000 | 20 | 10 | 0,90181618 | -0,05501413 | 21,82054 | 21,96969 | 22,18013 | 21,87849 | 21,29771 | 21,48207 | 23,1426 | 21,70641 |
| sp\|P53621\|COPA_HUMAN | sp\|P53621\|C | 14,631 | 50398000 | 23 | 11 | 0,90738445 | -0,02660751 | 22,1062 | 21,65211 | 21,5255 | 21,53146 | 21,4491 | 21,34066 | 21,84506 | 22,07401 |
| sp\|P60866\|RS20_HUMAN;tr\|E5RJX2\|E5RJX2_HUMAN;tr\|E5RIP1\|E5RIP1_HUMAN;tr\|G3XAN0\| | sp\|P60866\|R | 254,75 | 389950000 | 14 | 4 | 0,91025009 | -0,02446508 | 24,54975 | 24,72775 | 24,93112 | 24,26019 | 24,93939 | 24,68244 | 24,21104 | 24,53807 |
| tr\|F8W6I7\|F8W6I7_HUMAN;sp\|P09651\|ROA1_HUMAN;tr\|F8VZ49\|F8VZ49_HUMAN;tr\|F8VTQ | tr\|F8W6I7\|F | 237,52 | 970900000 | 63 | 15 | 0,91108497 | 0,03852797 | 25,8149 | 25,50588 | 26,05007 | 25,88816 | 25,30758 | 26,10554 | 26,61344 | 25,38657 |
| sp\|P42285\|SK2L2_HUMAN | sp\|P42285\|S | 3,4012 | 23330000 | 6 | 3 | 0,9121795 | 0,01348241 | 20,78061 | 20,65334 | NaN | 20,6644 | 20,60711 | 20,5417 | 20,73529 | 20,96762 |
| sp\|P16615\|AT2A2_HUMAN;tr\|H7C5W9\|H7C5W9_HUMAN;sp\|O14983\|AT2A1_HUMAN;sp\|Q9 | sp\|P16615\|A | 79,598 | 165810000 | 24 | 14 | 0,91327859 | 0,03349352 | 23,02837 | 23,14246 | 23,88064 | 23,08618 | 23,55344 | 23,19268 | 23,75562 | 22,76987 |
| sp\|Q92621\|NU205_HUMAN;tr\|U3KPX2\|U3KPX2_HUMAN;tr\|U3KQ47\|U3KQ47_HUMAN;tr\|U3 | sp\|Q92621\|N | 11,311 | 23199000 | 11 | 7 | 0,91380118 | -0,00850964 | 20,5559 | 20,61369 | 20,48434 | 20,64201 | 20,39111 | 20,68594 | 20,53222 | 20,65264 |
| tr\|J3KTA4\|J3KTA4_HUMAN;sp\|P17844\|DDX5_HUMAN;tr\|J3QSF1\|J3QSF1_HUMAN;tr\|J3KRZ1\| | tr\|J3KTA4\|J3 | 66,808 | 619820000 | 69 | 12 | 0,9168989 | 0,01343727 | 25,11874 | 25,15943 | 25,51226 | 25,13762 | 25,0918 | 25,25176 | 25,46566 | 25,17257 |
| sp\|Q96RL7\|VP13A_HUMAN;tr\|H0Y7P8\|H0Y7P8_HUMAN | sp\|Q96RL7\|V | 12,882 | 33363000 | 16 | 9 | 0,92109773 | 0,03409243 | 20,81167 | 21,16153 | 20,37401 | 21,77692 | 21,15956 | 20,97225 | 21,40826 | 20,72042 |
| sp\|P24539\|AT5F1_HUMAN;tr\|Q5QNZ2\|Q5QNZ2_HUMAN | sp\|P24539\|A | 35,898 | 55004000 | 15 | 5 | 0,92183065 | -0,01769686 | 21,97053 | 21,75756 | 21,93769 | 21,67008 | 21,5552 | 22,25046 | 21,8418 | 21,6176 |
| sp\|Q9BVP2\|GNL3_HUMAN | sp\|Q9BVP2\|G | 12,779 | 81100000 | 15 | 8 | 0,92442291 | 0,01325178 | 22,49975 | 22,31332 | 22,52986 | 22,27694 | 22,38717 | 22,10236 | 22,5413 | 22,64205 |
| sp\|Q5VYK3\|ECM29_HUMAN;tr\|J3KN16\|J3KN16_HUMAN;tr\|R4GMY1\|R4GMY1_HUMAN | sp\|Q5VYK3\|E | 6,9008 | 12704000 | 11 | 6 | 0,93132973 | 0,01784134 | 20,03832 | 19,30541 | 19,654 | 19,63456 | 19,44447 | 20,04394 | 19,65901 | 19,55624 |
| tr\|F8W726\|F8W726_HUMAN;sp\|Q14157\|UBP2L_HUMAN;tr\|H0Y5H6\|H0Y5H6_HUMAN;tr\|H7 | tr\|F8W726\|F | 11,972 | 37125000 | 13 | 5 | 0,93455125 | -0,02633238 | 20,63803 | 21,85167 | 21,58486 | 21,37311 | 21,51523 | 20,89896 | 21,28337 | 21,64478 |
| tr\|A0A0J9YXB3\|A0A0J9YXB3_HUMAN;sp\|P61224\|RAP1B_HUMAN;sp\|A6NIZ1\|RP1BL_HUMAN; | tr\|A0A0J9YXB | 18,169 | 72371000 | 17 | 5 | 0,94890021 | 0,02404499 | 21,72617 | 22,05417 | 22,28427 | 21,96353 | 21,49678 | 22,77282 | 21,40914 | 22,44559 |
| sp\|P49721\|PSB2_HUMAN;tr\|A0A087WVV1\|A0A087WVV1_HUMAN | sp\|P49721\|P | 40,094 | 68796000 | 13 | 4 | 0,95049717 | -0,01096582 | 22,42368 | 22,06313 | 22,03882 | 22,12364 | 22,48785 | 22,01699 | 21,82908 | 22,27147 |
| sp\|O75396\|SC22B_HUMAN;tr\|A0A087X1A9\|A0A087X1A9_HUMAN | sp\|O75396\|S | 18,706 | 48361000 | 11 | 5 | 0,96286928 | 0,00739241 | 21,38822 | 21,61993 | 21,48335 | 21,73099 | 21,93009 | 21,38195 | 21,58288 | 21,35714 |
| sp\|P49792\|RBP2_HUMAN;sp\|Q7Z3J3\|RGPD4_HUMAN;sp\|A6NKT7\|RGPD3_HUMAN;tr\|J3KNE | sp\|P49792\|R | 11,109 | 28534000 | 14 | 7 | 0,9766062 | 0,00496769 | 20,66258 | 20,86831 | 20,94027 | 20,6778 | 20,39435 | 20,86204 | 21,10194 | 20,81049 |
| sp\|P51991\|ROA3_HUMAN;tr\|H7C1J8\|H7C1J8_HUMAN | sp\|P51991\|R | 137,69 | 313080000 | 33 | 10 | 0,98585009 | 0,00791454 | 24,03414 | 23,82852 | 24,34908 | 23,99213 | 23,18696 | 24,32113 | 25,0857 | 23,64174 |
| sp\|Q9H9A6\|LRC40_HUMAN | sp\|Q9H9A6\|L | 5,2626 | 16544000 | 5 | 4 | 0,99428915 | 0,0008378 | 20,14607 | 19,97141 | 19,96254 | 20,11959 | 19,96367 | 19,98565 | 19,90317 | 20,35046 |
