## Supplementary material for "Subcellular localization and mitotic interactome analyses identify SIRT4 as a centrosomally localized and microtubule associated protein": Table S2

| Primary antibodies | Supplier | Species | Dilution | Reference |
|---|---|---|---|---|
| Flag M2 | Sigmal-Aldrich | mouse | 1:500 | F3165 |
| eGFP | Roche | mouse | 1:2.000 | 11814460001 |
| SIRT4 | Proteintech | mouse | 1:40.000 | 66543-1-Ig |
| SIRT3 | Cell Signaling Technology | rabbit | 1:500 | 5490 |
| HDAC6 | Santa Cruz | rabbit | 1:1000 | sc-11420 |
| HDAC6 | Cell Signaling Technology | rabbit | 1:1000 | 7558 |
| acetyl. Tubulin (K40) | Abcam | mouse | 1:500 | ab24610 |
| $\alpha$-Tubulin | Abcam | rabbit | 1:1000 | ab52866 |
| OPA1 | BD | mouse | 1:1000 | 612607 |
| OPA1 | | rabbit | 1:1000 | Barrera M et al. FEBS Lett.2016; 590:3309–22. https://doi.org/10.1002/1873-3468.12384 |
| ATP5A1 | Proteintech | rabbit | 1:1000 | 14676-1-AP |
| ANT2 | Cell Signaling Technology | rabbit | 1:1000 | 14671 |
| $\gamma$-Tubulin | Sigma-Aldrich | mouse | 1:1000 | T6557 |
| GCP2 | GeneTex | rabbit | 1:1000 | GTX102281 |
| GCP3 | Santa Cruz | mouse | 1:1000 | sc-373758 |
| Pericentrin | Abcam | rabbit | 1:1000 | ab4448 |
| GAPDH | Santa Cruz | mouse | 1:1000 | sc-47724 |
| Na+ / K+ ATPase | Sigma-Aldrich | mouse | 1:1000 | A276 |
| CDK1 | BD | mouse | 1:500 | 610037 |
