## Supplementary material for "Subcellular localization and mitotic interactome analyses identify SIRT4 as a centrosomally localized and microtubule associated protein": Table S3

| Primary antibodies | Supplier | Species | Dilution | Reference |
|---|---|---|---|---|
| eGFP | Roche | mouse | 1:1000 | 11814460001 |
| HDAC6 | Cell Signaling Technology | rabbit | 1:200 | 7558 |
| α-Tubulin | Santa Cruz | rat | 1:500 | sc-53029 |
| α-Tubulin | Acris | rat | 1:500 | SM 568P |
| γ-Tubulin | Sigma-Aldrich | mouse | 1:500 | T6557 |
| Pericentrin | Abcam | rabbit | 1:1000 | ab4448 |
| Pericentrin | Abcam | mouse | 1:1000 | ab28144 |
| SIRT4 | Santa Cruz | rabbit | 1:500 | sc-135053 |
| SIRT4 | Sigma Aldrich | mouse | 1:500 | SAB1407208 |
| TUBGCP2 | Santa Cruz | mouse | 1:500 | sc-377117 |
| TUBGCP3 | Santa Cruz | mouse | 1:500 | sc-373758 |
| SIRT3 | Cell Signaling Technology | rabbit | 1:500 | 5490 |
| MTC02 | Abcam | mouse | 1:500 | ab3298 |
| TACC3 | Santa Cruz | rabbit | 1:500 | sc-22773 |

| Secondary antibodies | Supplier | Species | Species reactivity | Dilution | Reference |
|---|---|---|---|---|---|
| Alexa Fluor 633 | Invitrogen - Thermo Fisher Scientific | goat IgG | anti-rat | 1:1000 | A-21094 |
| Alexa Fluor 488 | Invitrogen - Thermo Fisher Scientific | goat IgG | anti-rabbit | 1:1000 | A-11034 |
| Alexa Fluor 546 | Invitrogen - Thermo Fisher Scientific | goat IgG | anti-rabbit | 1:1000 | A-11035 |
| Alexa Fluor 488 | Invitrogen - Thermo Fisher Scientific | goat IgG | anti-mouse | 1:1000 | A-11029 |
| Alexa Fluor 546 | Invitrogen - Thermo Fisher Scientific | goat IgG | anti-mouse | 1:1000 | A-11003 |
