## Supplementary figures and images for "Subcellular localization and mitotic interactome analyses identify SIRT4 as a centrosomally localized and microtubule associated protein"

### Suppl. Movie 1

#### Slide 1
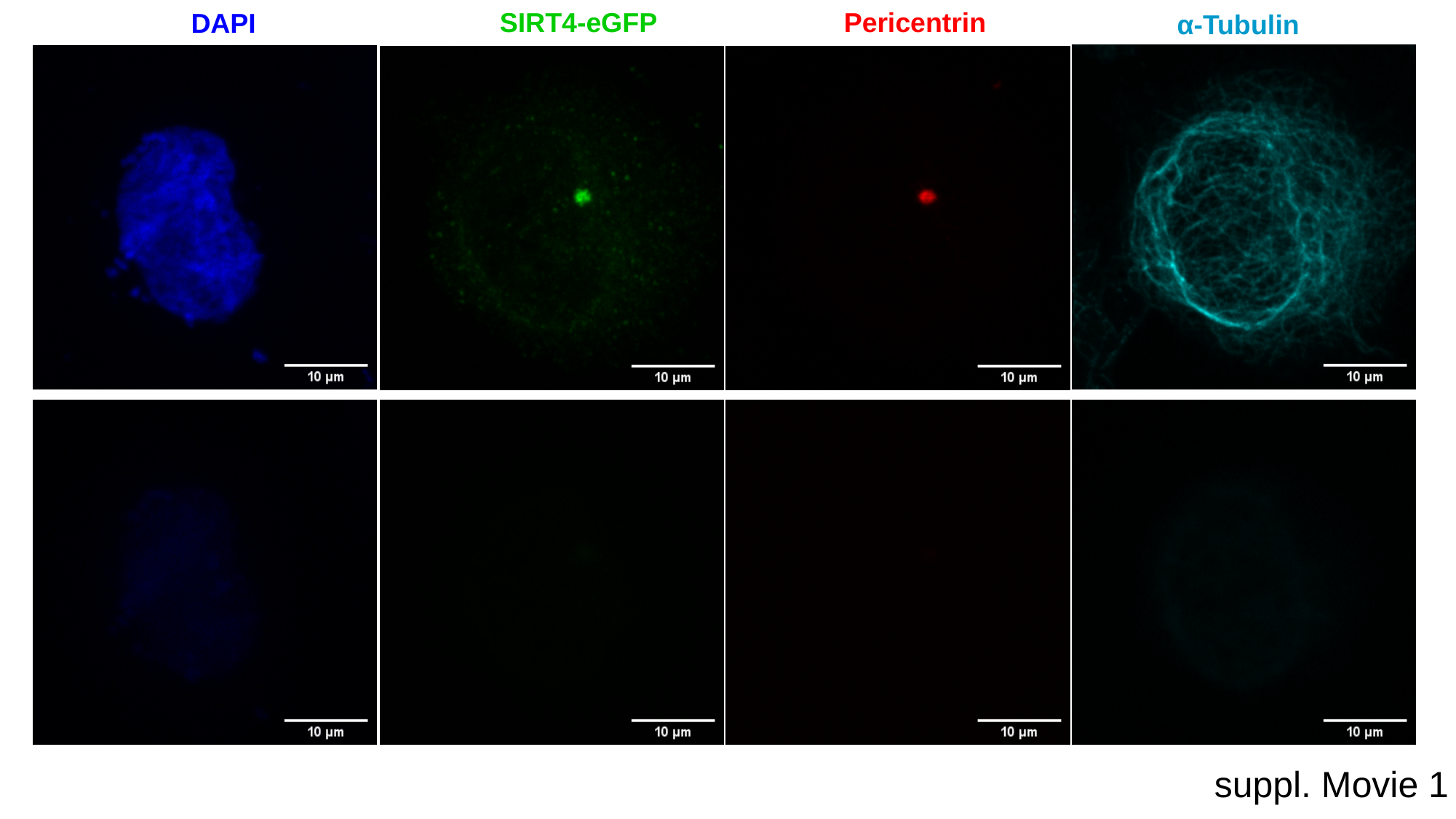

SIRT4-eGFP
Pericentrin
DAPI
α-Tubulin
suppl. Movie 1

### Suppl. Movie 2

#### Slide 1
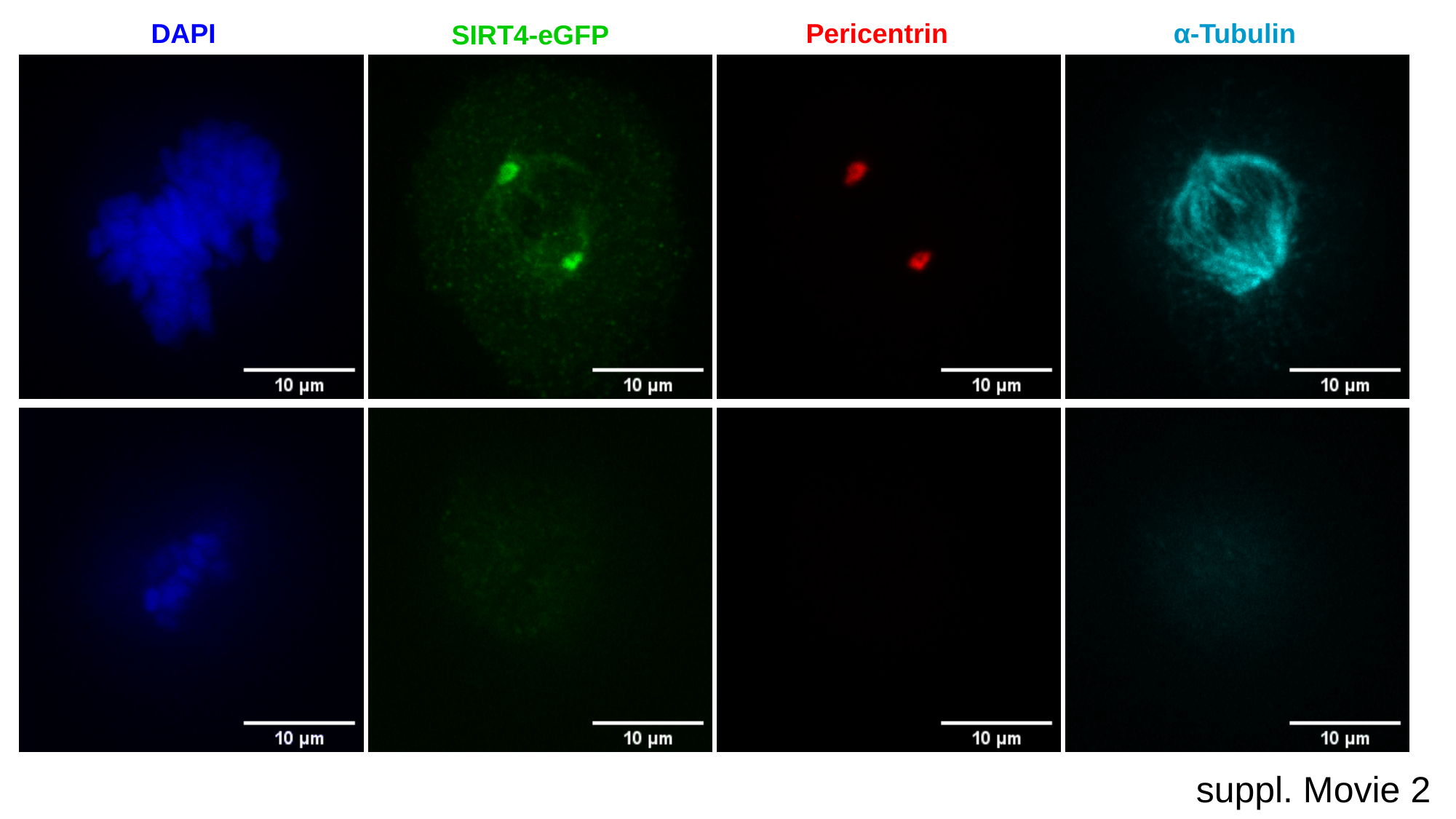

DAPI
Pericentrin
α-Tubulin
SIRT4-eGFP
suppl. Movie 2

### Suppl. Movie 3

#### Slide 1
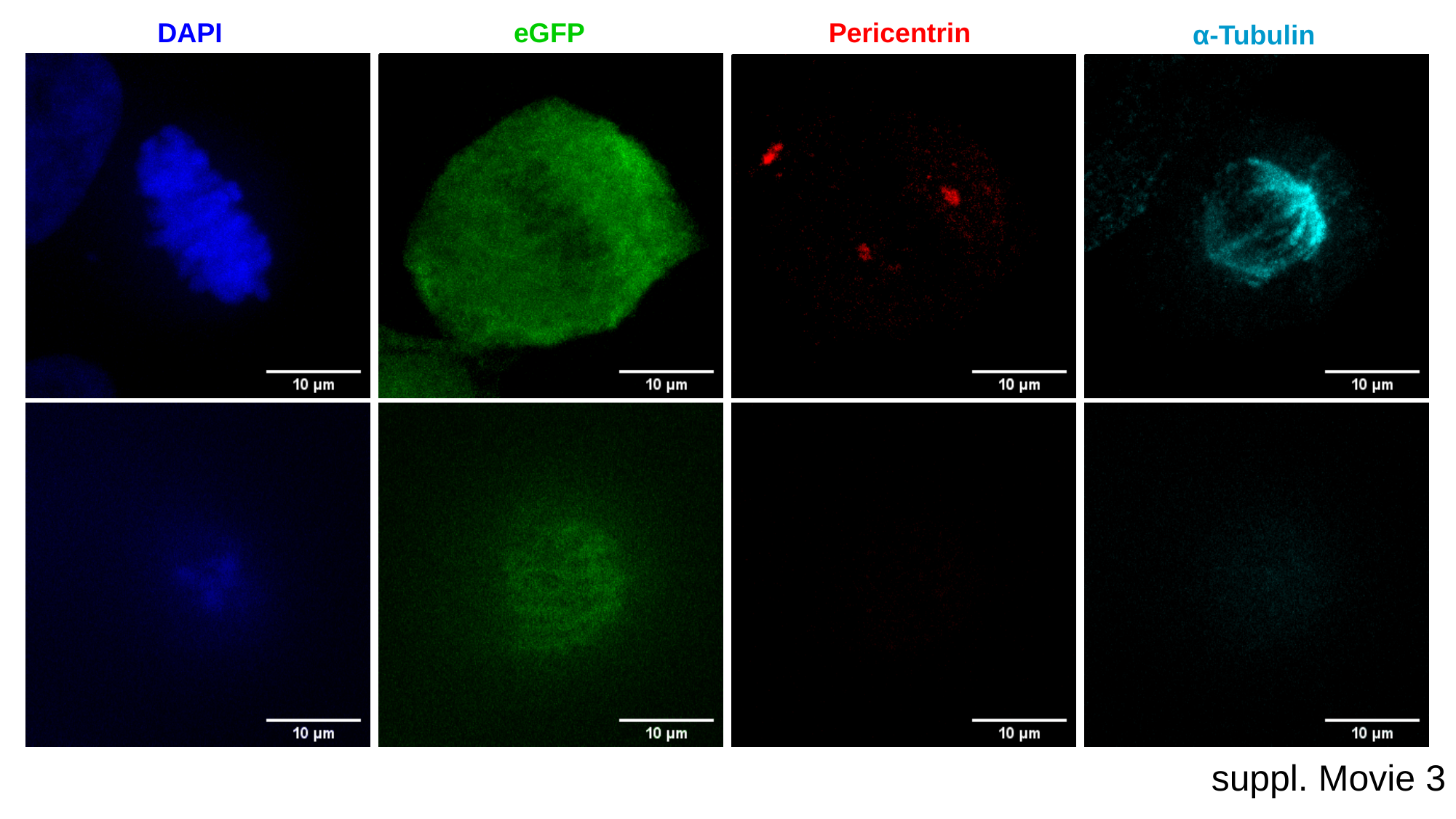

DAPI
eGFP
Pericentrin
α-Tubulin
suppl. Movie 3

### Suppl. Movie 4

#### Slide 1
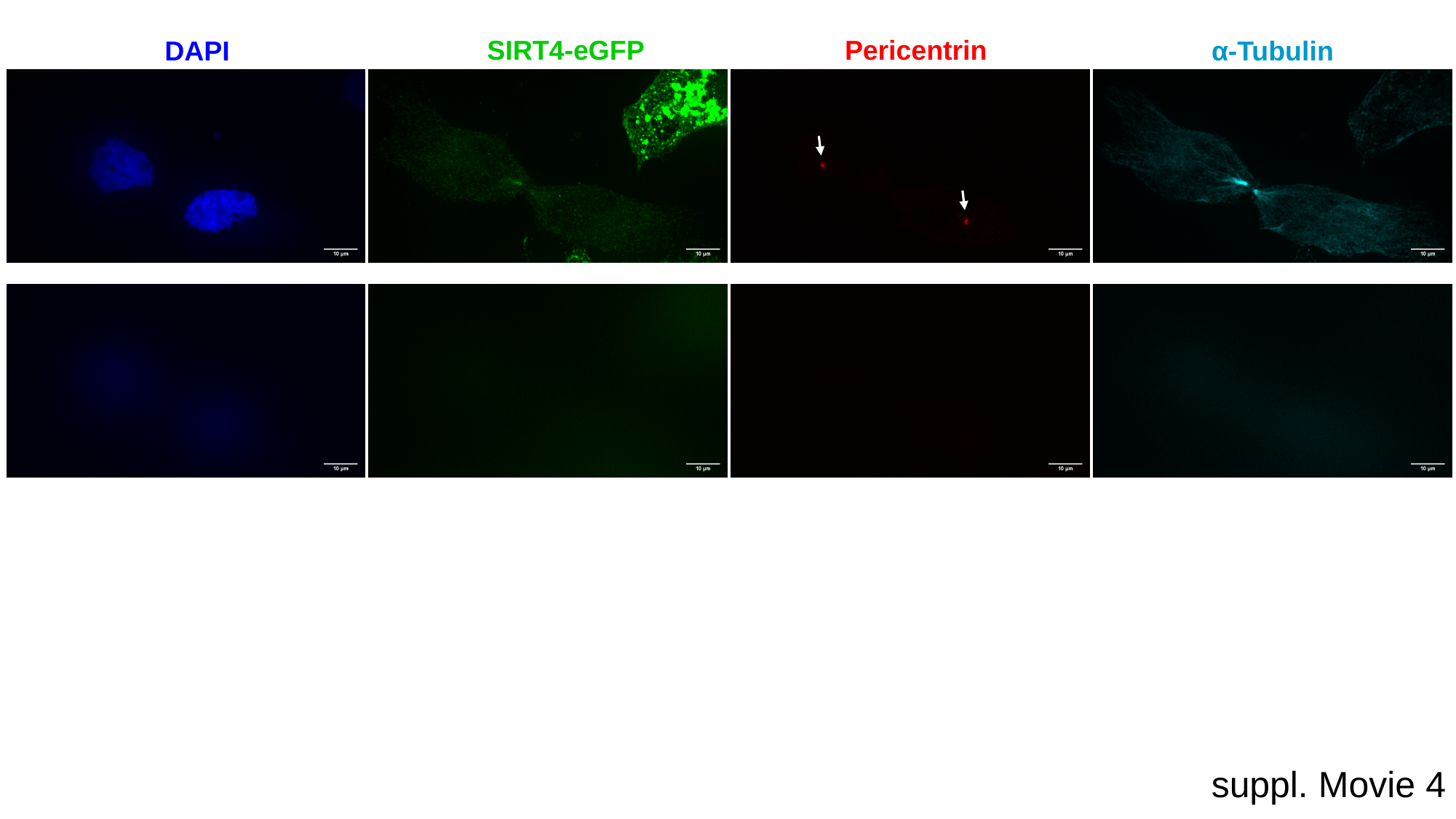

SIRT4-eGFP
Pericentrin
α-Tubulin
DAPI
suppl. Movie 4

### Suppl. Movie 5

#### Slide 1
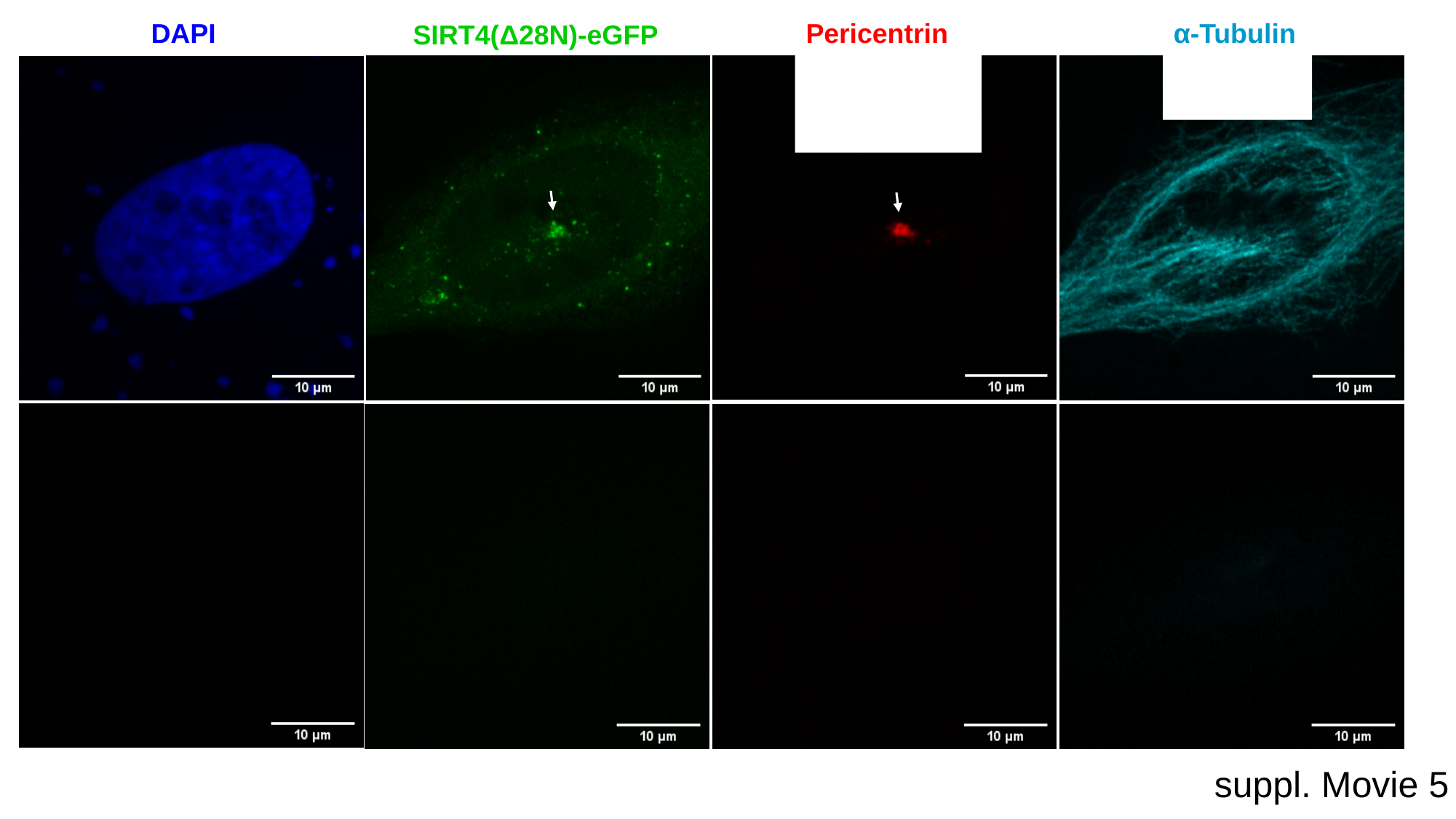

DAPI
Pericentrin
α-Tubulin
SIRT4(Δ28N)-eGFP
suppl. Movie 5

### Suppl. Movie 6

#### Slide 1
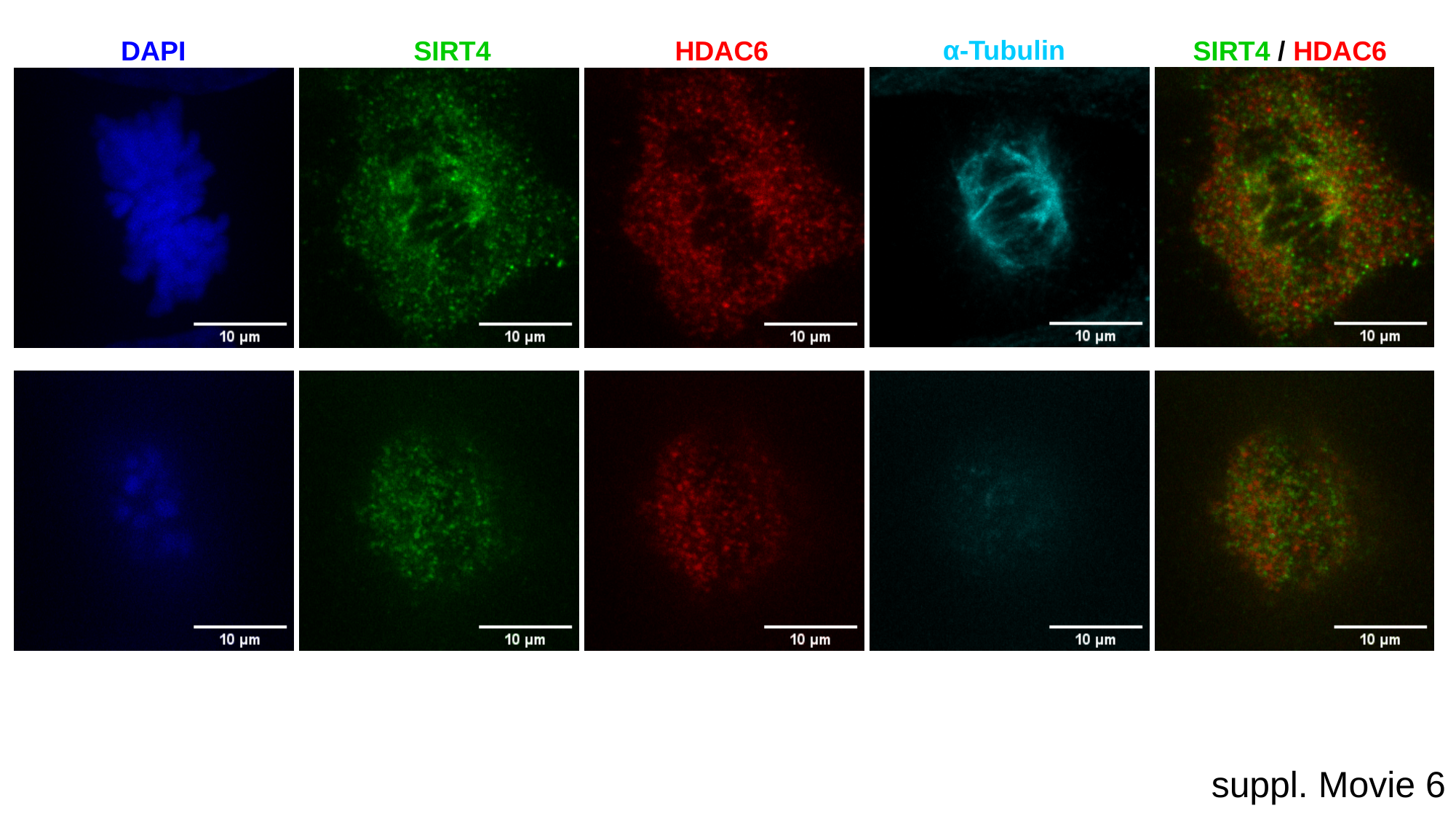

α-Tubulin
DAPI
SIRT4
HDAC6
SIRT4 / HDAC6
suppl. Movie 6
